# Inhibition of Topoisomerase 2 catalytic activity impacts the integrity of heterochromatin and repetitive DNA and leads to interlinks between clustered repeats

**DOI:** 10.1101/2023.08.01.551420

**Authors:** Michalis Amoiridis, Karen Meaburn, John Verigos, William H Gittens, Tao Ye, Matthew J Neale, Evi Soutoglou

**Affiliations:** Genome Damage and Stability Centre, Sussex University, School of Life Sciences, University of Sussex, Brighton, BN1 9RH, UK; Institut de Génétique et de Biologie Moléculaire et Cellulaire (IGBMC), Illkirch, France; Institut National de la Santé et de la Recherche Médicale (INSERM), U1258, Illkirch, France; Centre National de Recherche Scientifique (CNRS), UMR7104, Illkirch, France; Université de Strasbourg, Illkirch, France

## Abstract

DNA replication and transcription generate DNA supercoiling, which can cause topological stress and intertwining of daughter chromatin fibers, posing challenges to the completion of DNA replication and chromosome segregation. Type II topoisomerases (Top2s) are enzymes that relieve DNA supercoiling and decatenate braided sister chromatids. How Top2 complexes deal with the topological challenges in different chromatin contexts, and whether all chromosomal contexts are subjected equally to torsional stress and require Top2 activity is unknown. Here we show that catalytic inhibition of the Top2 complex in interphase has a profound effect on the stability of heterochromatin and repetitive DNA elements. Mechanistically, we find that catalytically inactive Top2 is trapped around heterochromatin leading to DNA breaks and unresolved catenates, which necessitate the recruitment of the structure specific endonuclease, Ercc1-XPF, in an Slx4- and SUMO-dependent manner. Our data are consistent with a model in which Top2 complex resolves not only catenates between sister chromatids but also inter-chromosomal catenates between clustered repetitive elements.

## Introduction

The physiological processes of DNA replication or transcription impose a physical strain on the backbone of DNA, which can lead to topological stress. Unresolved topological stress can hinder faithful completion of replication and sister chromatid decatenation and therefore pose a threat to genomic integrity. Topoisomerases are enzymes that modify the topology of the DNA, and are associated with replication, transcription, and recombination^1^. Mammalian topoisomerases are divided into two classes based on their structure and catalytic mechanism: Type I, such as Topoisomerase 1 (Top1), cut one strand of the DNA, whereas Type II, such as Topoisomerase 2 (Top2), cuts both DNA strands.

Top2 enzymes introduce a double strand break in one DNA segment and form a transient covalent bond with the free DNA overhangs. Immediately after, they pass a second DNA segment through the cut and instantly re-ligate the first segment, thus resolving issues of DNA supercoiling, knotting and catenation^2^. There are two distinct Top2 proteins, Top2α and Top2β, which despite having high similarity at the N-terminal domain, have divergent functions in the cell^3^. Top2α acts during replication, where it is required for proper chromosome compaction and segregation, as well as the decatenation of the sister chromatids at mitosis. Fitting with these roles, Top2α has a cell cycle dependent expression, where its levels increase in S and G2 phase, and thus, is considered a proliferation marker, and it is overexpressed in metastatic and highly proliferating tumors^4^. On the other hand, Top2β is linked with transcription^4^, and its expression remains steady throughout the cell cycle, although it is also overexpressed in tumors^5^.

Due to the importance of preserving genomic integrity for continuous cell proliferation, drugs which can induce topological stress have been efficiently used in clinics to target cancer cells. Nevertheless, off-target side effects of these drugs highlight the critical need to better understand their mechanism of action and for elucidating how to limit their toxicity to only cancer cells. A fairly common approach in chemotherapies is the use of topoisomerase targeting drugs, of which there are two types, the poisons and the catalytic inhibitors. Top2 poisons stabilize the covalent bond between the DNA and the Top2 protein forming DNA-Top2 cleavage complexes (Top2ccs). As a result, unrepaired DSBs are formed, causing major genomic instability, and ultimately leading to cell death^6^. In the clinical setting, Top2 poisons, such as Etoposide, are routinely used for the treatment of both solid tumors and hematologic cancers. However, even though Etoposide can efficiently target cancer cells, its use is correlated with cardiotoxicity and development of secondary leukemia-associated malignancies, due to distinct chromosomal translocations^4^. On the contrary, catalytic inhibitors, such as ICRF-193, inhibit the ATPase activity of Top2, keeping it in a closed “clamp” form^4^ and are proposed to only inactivate Top2, without creating Top2ccs. It remains debated whether Top2 catalytic inhibitors are capable of inducing DSBs in cells, what effect the catalytic inactivation of Top2 might have for genomic integrity.

The genome is non-randomly arranged within the nuclear space^7^. The 3D genome organization corelates with numerous functional features such as gene activity, chromatin structure and replication timing. Although the genome wide distribution of Top2 is emerging, what genomic contexts require most Top2 activity, what parts of the genome are affected most by Top2 poisons or inhibitors, and the consequences of Top2 deregulation for human health are currently unknown. Elucidating these questions could provide vital information for the therapeutic use of Top2 drugs.

In this study, we demonstrate that catalytic inhibition of Top2 induces damage at heterochromatin and repetitive DNA regions and affects the progression of the cell cycle in a Top2α-dependent manner. Moreover, whole genome sequencing reveals the genomic hotspots where the damage is induced and demonstrates that inhibition of Top2s leads to the formation of DSBs, genomic rearrangements and abnormalities associated with repetitive DNA. Our data also demonstrate that the catalytic activity of Top2α is necessary for DNA damage induction and that it is the trapping of the Top2α enzyme around chromatin that leads to the induction of the DNA damage. Most importantly, inhibition of Top2 leads to inter-chromosomal catenates of pericentric repeats from different chromosomes, which are resolved by the structure specific endonuclease Ercc1-XPF, in an Slx4 and Sumo-dependent manner.

Altogether, our results shed light into the mechanism of action of Top2 catalytic inhibitors and broaden our understanding of how they affect genomic stability and can be exploited for therapeutic purposes.

## Results

### Inhibition of Top2 catalytic activity leads to DNA damage at heterochromatin in a cell cycle-dependent manner

To study the role of Top2 activity in protecting chromatin from high torsional stress, which can lead to DNA damage, we identified the chromosomal contexts which are most affected by the catalytic inhibition of Top2. To this end, we used mouse NIH3T3 cells in which heterochromatin can be easily visualized in DAPI dense regions, known as chromocenters, and treated them with the Top2 catalytic inhibitor ICRF-193. We found that ICRF-193 mainly induces DNA damage at heterochromatin, exemplified by γH2AX strictly colocalizing with the DAPI-dense regions in 52% of treated NIH3T3 cells (Figure 1A and 1B). Interestingly, inducing DNA damage at heterochromatin was specific to ICRF-193 as the Top2 poison etoposide induced DNA damage in both euchromatin and heterochromatin at 62.1% of cells, and only 11.3% of cells were found with damage enriched at heterochromatin (Figure 1A and 1B). The DNA damage induced by Top2 inhibition is not permanent, as it was repaired within 6h after removal of ICRF-193 (Figure 1C). The induction of damage by ICRF-193 and the ability of cells to repair this damage after the release from the drug were validated by western blot analysis (Figure S1A). Importantly, to verify that the γH2AX foci we observed co-localizing with DAPI-dense regions did indeed indicate DNA damage in heterochromatin, we performed Chromatin IP (ChIP) and analyzed the enrichment of γH2AX at major satellite repeats, which are abundant in heterochromatin. As expected, an over 6-fold increase in γH2AX was observed in the treated condition (ICRF-193) compared to the mock (DMSO) (Figure S1B).

**Figure 1:**
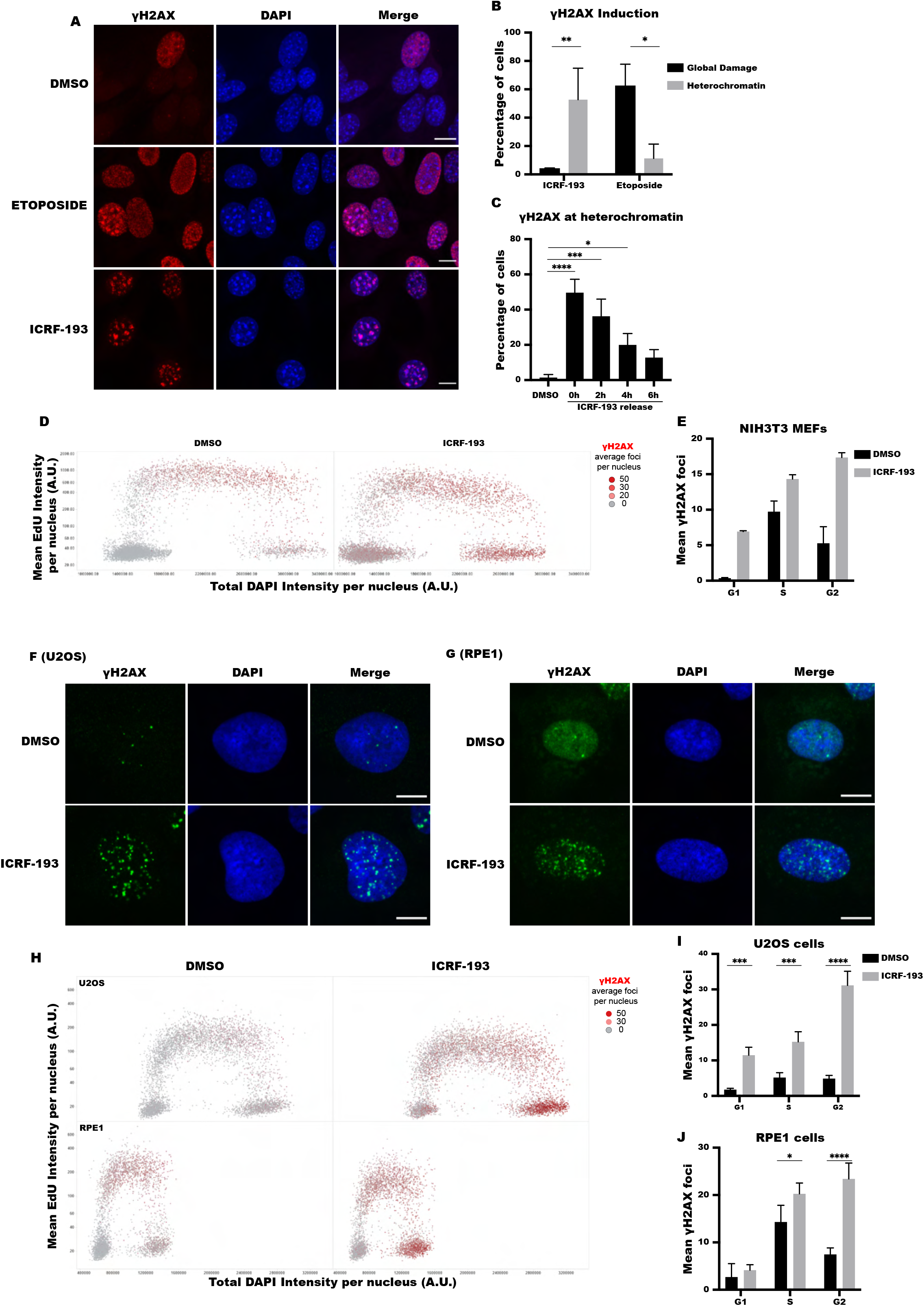
ICRF-193 induces DNA damage at heterochromatin on a cell cycle basis. A) Confocal microscopy images of γH2AX (red) in NIH3T3 cells treated for 4h with 5μM ICRF-193 or 5μΜ of etoposide. Maximum projection of z-stacks is shown and 4’,6-Diamidino-2-Phenylindole, Dihydrochloride (DAPI; blue) was used to visualize DNA. Scale bars = 10μM. B) Quantification of the percentage of cells with γH2AX staining localizing with heterochromatin or both with euchromatin and heterochromatin (global DNA damage). C) Quantification of the percentage of NIH3T3 cells positive for γH2AX at heterochromatin upon treatment with 5μm ICRF-193 for 4h and released in drug free medium for 0, 2, 4 or 6h. D) Quantitative Image-Based Cytometry (QIBC) plots of γH2AX levels in NIH3T3 cells treated with DMSO or ICRF-193 for 4h. The cell cycle stages were identified based on EdU and DAPI intensity. Each dot represents a single cell, and the color of the dots represents the number of γH2AX foci per cell. At least 1000 cells were analyzed per condition. A.U.: Arbitrary Units. One of the two independent biological replicates performed is shown. E) Quantification of the mean number of γH2AX foci from 2 biological replicates. At least 1000 cells were analyzed per condition, per replicate. F-G) Representative confocal microscopy images of γH2AX (green) and DAPI (blue) in (F) U2OS cells or (G) RPE1 cells treated with 5μM ICRF-193 or DMSO for 4h. Maximum projection of z-stacks is shown. Scale bars = 10μM. H) QIBC plots for γH2AX levels in U2OS or RPE1 cells treated with DMSO or ICRF-193 for 4h. The cell cycle staging was performed based on EdU and DAPI intensity. Each dot represents a single cell, and the color of the dots represents the number of γH2AX foci per cell. At least 2000 cells were analyzed per condition. I-J) Quantification of the mean γH2AX foci number per nuclei in (I) U2OS cells and (J) RPE1 cells. All graphs represent mean values, and the error bars represent the standard deviation from at least 50 cells per condition, and 3 independent replicates unless stated otherwise. Statistical significance presented by student *t*-test in 1B, one-way ANOVA test in 1C, and two-way ANOVA in 1I and 1J: (*) p<0.05, (**) p<0.005, (***) p<0.001, (****) p<0.0001

We then sought to understand whether the induction of DNA damage depended on the stage of the cell cycle, given that DNA damage is observed only in 50% of cells. To this end, we stained ΝΙΗ3T3 cells with EdU and phosphorylated H3S10 (pH3S10), as markers of S and G2/M^8^ respectively (Figure S1C) and quantified the percentage of γH2AX positive cells in each phase of the cell cycle. We found that cell cycle stage correlated with ICRF-193 induced DNA damage, as only 25% of G1 cells (EdU^-^, pH3S10^-^) showed damage, while about 50% of the S phase cells (EdU^+^ pH3S10^-^), and nearly 100% of cells in G2 (EdU^-^, pH3S10^+^), had damaged heterochromatin (Figure S1D). Using distinct EdU patterns to differentiate between early (EdU signal covering euchromatin; Figure S1C,), mid (EdU signal co-localizing with heterochromatin; Figure S1C,), and late (EdU signal located at the periphery of the nucleus and the nucleolus; Figure S1C) replicating cells^9^, we found that the γH2AX signal colocalized with heterochromatin in mid and late S phase suggesting that DNA damage at heterochromatin was enriched during and after replication (Figure S1E).

For further validation of DNA damage induction based on a cell cycle stage, we used Quantitative Image-Based Cytometry (QIBC), which involves a high-content automated microscopy approach and allows the unbiased analysis of individual asynchronous cells in large cell populations (>1000 cells per condition). The cell cycle phase of each cell is classified based on DNA content and EdU incorporation. In accordance with the previous quantification by confocal microscopy, QIBC analysis for γH2AX foci revealed that ICRF-193-induced DNA damage in S and G2 phases of the cell cycle (Figure 1D and 1E). Interestingly, the distribution of γH2AX perfectly correlated with the expression pattern of Top2α (Figure S1F). These data were validated in cells synchronized in the G1/S with thymidine or in G2/M with the Cdk1 inhibitor, RO-3306 (Figure S1G-H). Despite the different spatial organization of heterochromatin in human cells, similar to our observations in mouse cells, the induction of DNA damage by ICRF-193 is most evident in S and G2 phases of the cell cycle in human bone osteosarcoma epithelia (U2OS) and human retinal pigment epithelial-1 (RPE1) cells (Figure 1F-1J). Therefore, when taken together our data demonstrate that inhibition of the catalytic activity of Top2 leads to DNA damage at heterochromatin in mid/late S and G2 phases of the cell cycle, when Top2α is expressed.

To decipher whether the induction of DNA damage requires DNA replication, EdU was supplemented into the media 15 minutes prior to ICRF-193 treatment. By adding the EdU before ICRF-193, we could ensure we analyzed the population of G2 cells in which replication had already finished before ICRF-193 was added (the EdU^-^ G2 cells (based on DAPI content)) (Figure S1I). Interestingly, ICRF-193 induces γH2AX foci formation at 92.6% of the EdU^-^ G2 population (p=4.39*10^-7^), revealing that active replication is not necessary during the treatment for DNA damage to be induced, and that the DNA damage is not necessarily passed on to G2 cells from the previous S phase (Figure S1J and S1K).

To examine how ICRF-193 affects cell cycle progression, we performed cell cycle analysis in the presence or absence of ICRF-193. As shown previously^10^, we also observed an increase in the percentage of G2 cells (Figure S1L). Interestingly, we also detected a significant decrease in the number of early replicating cells, with an accompanying increase in both mid and late replicating cells (Figure S1L & S1M). These results suggest that ICRF-193 decreases the speed of replication in mid and late replicating domains. After observing the increase in replication time for heterochromatin during ICRF-193 treatment, we sought to uncover whether cells would delay the transition from S to G2 phase. To this end, we quantified the percentage of cells that are simultaneously positive for the S-phase marker EdU and the G2 marker pH3S10. There was a significant increase in the percentage of cells with this dual mark, after treatment with ICRF-193 (Figure S1N). This observation indicates that even if the S phase is delayed, cells progress to G2 before completing replication.

These results altogether suggest that Top2 activity protects replicating and post replicating heterochromatin from DNA damage, which can delay DNA replication and activate the G2-M checkpoint.

### ICRF-193-induced DNA damage depends on Top2α

Bisdioxopiperazines, a class of Top2 catalytic inhibitors which include ICRF-193, interact in a non-competitive way with the N-terminal site of the Top2 homodimers, and block ATP hydrolysis and enzyme turnover^11^. Since Top2α and Top2β isozymes show an extremely high degree of homology at their N-terminal domains^12^, potentially ICRF-193 could effectively inhibit the function of both Top2 isozymes. However, given that γH2AX foci formation by ICRF-193 depends on the phase of the cell cycle, and that Top2α and Top2β exhibit different cell cycle expression patterns^13^, we sought to investigate whether the two enzymes contribute equally to the induction of DNA damage. To this end, Top2α and Top2β were individually depleted by siRNA in NIH3T3 cells. As Top2α is expressed from mid S-G2, and Top2β throughout the cell cycle, we depleted Top2α in G2-arrested cells and Top2β in an asynchronous population. Depletion of Top2α led to a 3.2-fold reduction of γH2AX positive ICRF-193 treated cells, when compared to the scramble (Scr) control (Figure 2A & 2B). On the contrary, treatment with ICRF-193 after depletion of Top2β led to a smaller reduction of γH2AX foci (1.67-fold reduction; Figure 2C and 2D). siTop2α efficiently depleted Top2α and had no effect on Top2β expression (Figure S2A-S2C). Conversely, while siTop2β efficiently depleted Top2β without affecting the progression of the cell cycle, it has a small but significant off-target downregulation of Top2α levels (1.28-fold reduction, Figure S2D-S2G), suggesting that the reduction in γH2AX foci in siTop2β cells, is due to the off-target reduction of Top2a. To further test the involvement of Top2β in ICRF-193 dependent DNA damage induction, we used Top2β knock out ΜEFs. QIBC analysis showed that γH2AX foci can still be efficiently formed even in the complete absence of Top2β (Figure S2H-S2J). The involvement of Top2α in ICRF-193-induced DNA damage was also further confirmed in HCT116 OsTIR1^14^ expressing cells, in which endogenous Top2α was fused to an auxin inducible degron tag (Top2αAID) to induce its rapid depletion. After ensuring that Top2α is efficiently depleted upon 1h treatment with Indole-3-acetic acid (IAA) (Figure S2K), the induction of DNA damage in these cells was verified by immunofluorecence and QIBC analysis of the mean number of γH2AX or 53BP1 foci and corroborated that Top2α depletion prevents the induction of ICRF-193-induced DNA damage (Figure 2E-2G). Altogether, these results suggest that the ICRF-193-induced DNA damage primarily depends on Top2α, and that the minor effect of Top2β depletion by siRNA on the reduction of ICRF-193-induced DNA damage (Figure 2C & 2D) can be explained by the off-target depletion of Top2α.

**Figure 2:**
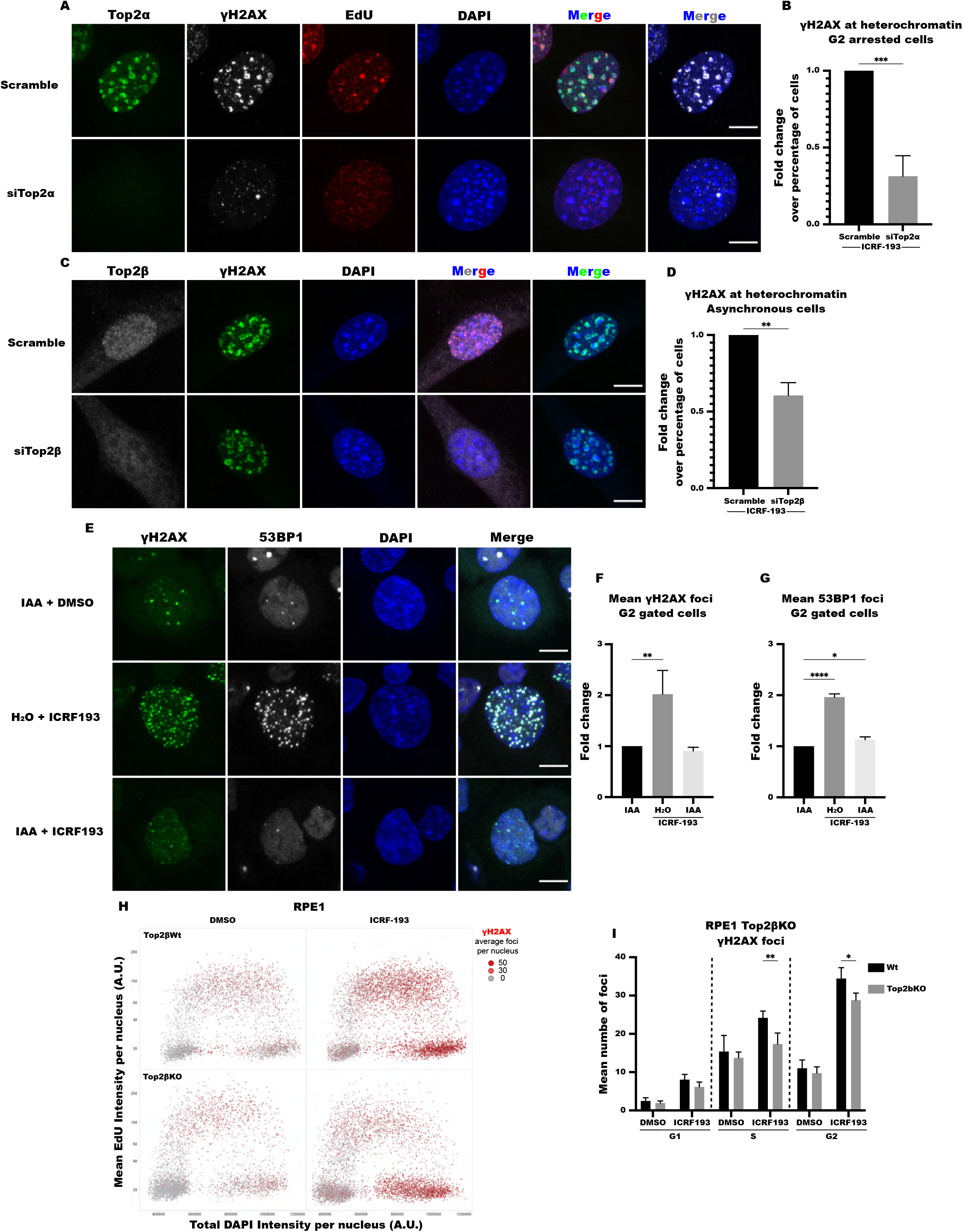
ICRF-193 induces damage by mainly targeting Top2α. A) Representative confocal microscopy images of Top2α (green), γH2AX (white), EdU (red) and DAPI (blue) in G2-arrested NIH3T3 cells transfected with scramble siRNA or siTop2α. Maximum projection of z-stacks is shown. Scale bars = 10μM. B) Quantification of the mid/late S or G2 phase NIH3T3 cells with γH2AX at heterochromatin at the indicated conditions. C) Representative confocal microscopy images of Top2β (white), γH2AX (green), EdU (red) and DAPI (blue) in NIH3T3 cells transfected with scramble siRNA or siTop2β. Maximum projection of z-stacks is shown. Scale bars = 10μM. D) Quantification of NIH3T3 cells with γH2AX at heterochromatin at the indicated conditions. E) Representative confocal microscopy images of γH2AX (green), 53BP1 (white) and DAPI (blue) in HCT116 OsTIR1/Top2αAID cells. Cells were treated with H_2_O or IAA prior to the addition of DMSO or ICRF-193. Maximum projection of z-stacks shown. Scale bars = 10μM. F) Quantification of the mean number of γH2AX foci for G2 gated cells, based on their DAPI intensity levels. The fold change of γH2AX levels in comparison to the control (IAA) condition are depicted. At least 800 cells were analyzed per condition, per replicate. G) Quantification of the mean number of 53BP1 foci for G2 gated cells, based on their DAPI intensity levels. The fold change of 53BP1 levels in comparison to the control (IAA) condition is shown. At least 800 cells were analyzed per condition, per replicate. H) QIBC plots of γH2AX foci number in Top2β wt and knock out (KO) RPE1 cells treated with DMSO or ICRF-193. The cell cycle staging was performed based on EdU and DAPI intensity. Each dot represents a single cell, and the colour of the dots represents the number of γH2AX foci per cell. At least 1000 cells were analyzed per condition, per replicate. I) Quantification of the mean number of γH2AX foci per cell in different stages of the cell cycle. All graphs represent average values, and the error bars represent the standard deviation from at least 50 cells per condition and 3 independent replicates unless stated otherwise; for the QIBC analysis at least 1000 cells were used per condition and 3 independent replicates unless stated otherwise. Statistical significance presented by student *t*-test in 2B and 2D, one-way ANOVA test in 2F and 2G, and two-way ANOVA test in 2I: (*) p<0.05, (**) p<0.005, (***) p<0.001, (****) p<0.0001

Next, after demonstrating that the targeting of Top2α by ICRF-193 is necessary for the induction of DNA damage, we explored whether the over-expression of Top2α is sufficient to trigger DNA damage in the presence of ICRF-193. To this end, we assessed if treatment with ICRF-193 induces DNA in G1 NIH3T3 cells, where the Top2α and DNA damage levels in heterochromatin are physiologically low. Interestingly, ectopic overexpression of Top2α-YFP, led to a very small increase (7.7%) in G1 cells which showed DNA damage at heterochromatin (Figure S2L).

When taken together, these results suggest that ICRF-193-induced DNA damage depends on Top2α, but its presence alone is not sufficient, and other factors may contribute to the induction of DNA damage, such as collision of the trapped Top2 proteins with the replication machinery and/or inhibition of sister chromatid decatenation^15^.

### ICRF-193 induces DSBs at repetitive sequences

Top2 enzymes bind to the genome regardless of chromatin status^16-19^. The main cause of DNA damage induction by Top2 poisons in G1 cells is the collision of Top2ccs with the transcription machineries in euchromatin^20,21^. Our results also show that the trapping of the Top2 enzymes by etoposide can induce damage anywhere in the nucleus (Figure 1A). Therefore, it is unexpected that treatment with ICRF-193 shows such a concentrated response at the heterochromatic domains of mouse cells (Figure 1A). To explore the possibility that due to the low sensitivity of immunofluorescence we may have failed to detect γH2AX enrichment in non-chromocenter regions of the genome after ICRF treatment, we performed γH2AX ChIP-seq in NIH3T3 MEFs treated with ICRF-193 or DMSO. To validate our approach, we first visualized the pericentric region of chromosome 9, in which since it locates to chromocenters, γH2AX enrichment is expected (Figure 3A). Indeed, broad γH2AX peaks spanning megabases were detected in cells treated with ICRF-193 compared to DMSO (Figure 3A).

**Figure 3:**
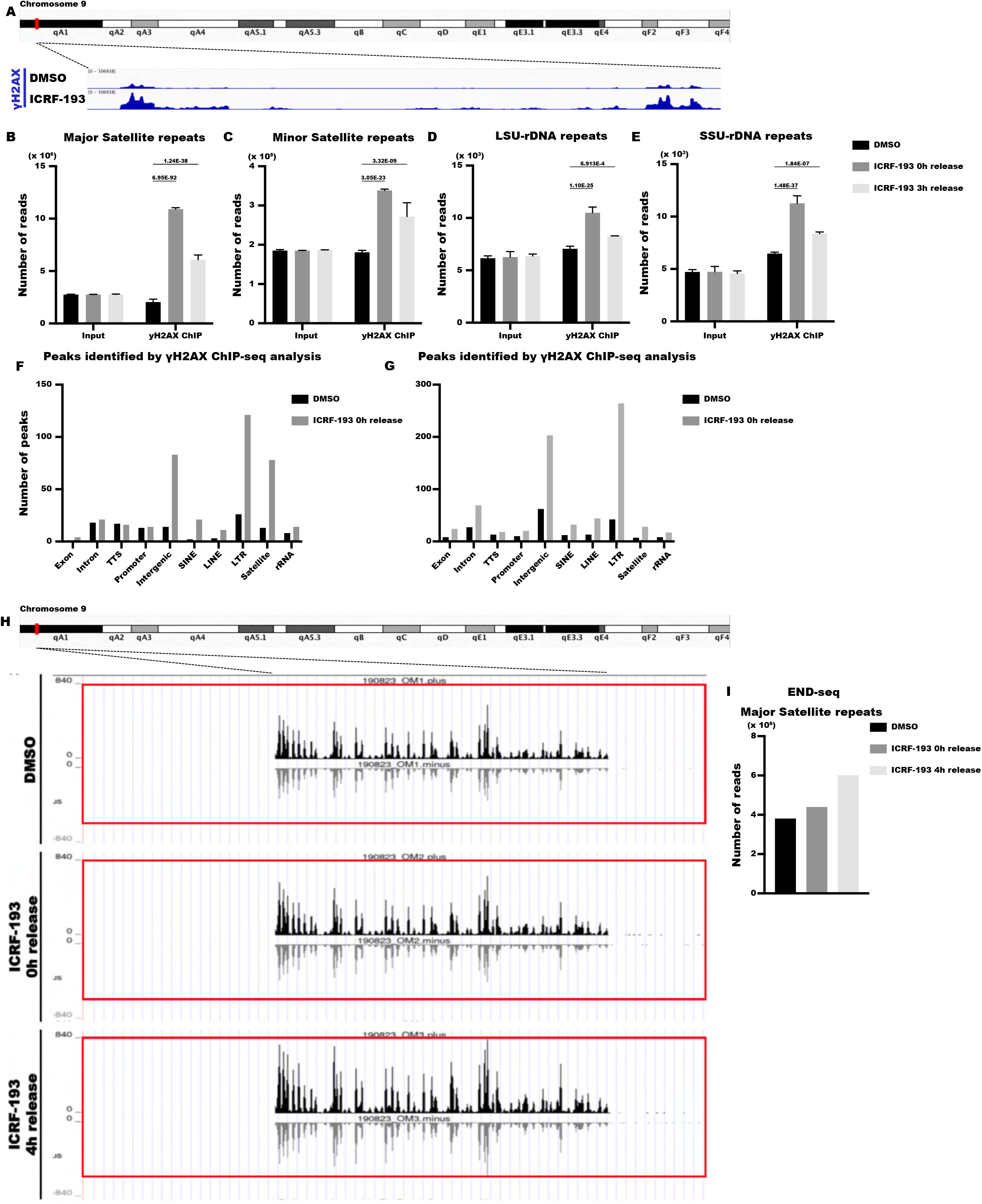
ICRF-193 induces DSBs in repetitive sequences. A) Representative image of the pericentric region from mouse chromosome 9, of mouse mm9 genomic assembly, showing the increase in number of reads with γH2AX signal after treatment of NIH3T3 cells ICRF-193. B-D) Quantification of the increase in number of reads after γH2AX-ChIPseq analysis in NIH 3T3 cells mapped to Major (B) and Minor (C) Satellite repeats, or to Long (D) and Small (E) subunits of ribosomal RNA repeats. Graphs represent average values, and the error bars represent the standard deviation from 2 biological replicates. Statistical significance presented by comparison with DE-Seq-2 analysis. F-G) Quantification of the number of peaks identified for the indicated genomic sequences in NIH3T3 cells treated with DMSO (black) or ICRF-193 (grey) in the first (F) and second (G) biological replicates. Peak calling was performed by MACS2 and they were annotated by HOMER to general categories of genomic regions such as Exons, Introns, Promoters and transcription termination sites (TTS) and repetitive regions such as SINEs, LINEs, LTRs, Satellite and rRNA repeats. H) Representative image of the pericentric region from chromosome 9 of mouse mm9 genomic assembly, of NIH3T3 cells treated with ICRF-193, showing the increase in number of reads by END-seq analysis. (I) Quantification of the number of reads after END-seq analysis, mapped to Major Satellite repeats. Data were analyzed by DE-Seq-2.

Next, we examined whether ICRF-193 can induce DNA damage only in major satellite repeats, which are abundant in pericentric regions, or if it has a broader effect on repetitive sequences. DESeq2 analysis on data from two biological replicates showed that ICRF-193 leads to a 5.3-fold enrichment of reads in major satellite repeats, compared to the DMSO control (Figure 3B). This observation is in accordance with our previous results and verifies the robust DNA damage induction we detect at heterochromatin by microscopy. We also assessed other nuclear compartments that contain tandem repeats, such as centromeric minor satellite repeats, or small subunit (SSU) and large subunit (LSU) rDNA, which are found in the nucleolus. Interestingly, we also detected a significant increase in the number of reads for these regions, after treatment with ICRF-193 (Figure 3C-3E). Additionally, we observed an enrichment in the number of reads for several other classes of repetitive sequences, such as Long Terminal Repeats (LTRs) (Table S1). Thus, we found an enrichment of reads in many classes of repetitive sites, showing that ICRF-193 can induce damage generally in repetitive DNA regions. Interestingly, our ChIP-seq analysis also validates that γH2AX levels are reduced after ICRF-193 is removed from the culturing media (Figure 3B-3E), showing that cells can repair the induced damage at heterochromatin.

To gain a fuller picture of the DNA damage hotspots and to identify if there is any preference for euchromatin or heterochromatin, we plotted the number of γH2AX peaks according to their position in the genome. In promoters, introns, and exons, we did not detect a consistent increase in the number of peaks after treatment with ICRF-193 in either biological replicate (Figure 3F & 3G), thus providing no evidence for targeted DNA damage induction in euchromatin. Even though the mapping of peaks on repetitive sequences is not entirely accurate, since the reads are randomly mapped in each region that has the same sequence, we detected a consistent increase of γH2AX peaks in heterochromatic regions, such as SINEs, LINEs and LTRs (Figure 3F and 3G). Altogether, these data suggest that the vast majority of the DNA damage caused by ICRF-193 is induced in repetitive sequences and not in euchromatin.

While γH2AX is a strong indicator of the DNA damage response (DDR) after the formation of DSBs, it is also tightly linked to the responses of replication stress^22-24^. To decipher whether ICRF-193 treatment leads to induction of DSBs, we first examined the recruitment of DSB repair factors 53BP1, BRCA1 and RPA32 at damaged heterochromatin (Figure S3A-S3D). Similar to γH2AX, 53BP1 and BRCA1 exerted peak enrichment (26 & 28.2% of cells respectively) after ICRF-193 treatment, and the percentage of cells with colocalization of these factors at heterochromatin decreased after release from the drug. Interestingly, there was a small but significant increase in the recruitment of the ssDNA binding protein RPA32 2h after drug release (Figure S3A and S3D). Similar results for 53BP1 foci were obtained in human U2OS and RPE1 cells (Figure S3E). Comparable with the γH2AX foci pattern, 53BP1 foci are induced in a cell cycle manner, with the highest signal increase found in G2 phase (4.3-fold for U2OS & 3.27-fold for RPE1; Figure S3F and S3G). These results suggest DSBs are indeed being formed.

To directly prove DSB formation, and to accurately map the breaks on the genome, we performed END-seq. Consistent with our γH2AX ChIP-seq data, we failed to detect any DBSs in euchromatic regions of the genome, yet we detected an increase in DSBs at the pericentric region of chromosome 9 (Figure 3H). When compared to DMSO treatment, ICRF-193 led to an increase in the number of reads at major satellite repeats (Figure 3I). Surprisingly, the biggest read enrichment was detected 4h after the drug release suggesting that DSBs are detected better when Top2 is removed from chromatin (Figure 3H-I).

We next aimed to decipher the major kinases involved in the DDR after ICFR-193 treatment. We found that for up to two hours after the release of the drug the induction of γH2AX primarily relies on ATM (Figure S3H). Subsequently, ATR is important for the phosphorylation of histone H2AX, until the repair is finished (Figure S3I). These two kinases seem to be the only enzymes responsible for the initiation of DDR, since combinational treatment of ATM inhibitor and ATR inhibitor completely blocked the induction of γH2AX by ICRF-193 (Figure S3J).

To understand which pathways are involved in repairing ICRF-193-induced DNA damage, we followed the γH2AX kinetics upon drug release in the presence of inhibitors of core DSB repair factors. Inhibition of DNAPK, which favors the non-homologous end joining (NHEJ) repair pathway, showed a significant delay in the repair after 4 or 6 hours of drug release (Figure S3K). Conversely, inhibition of Rad51, the core molecule for the homologous recombination (HR) repair pathway, did not have a significant impact in the repair kinetics of ICRF-193 (Figure S3L). This is an unexpected result, since ICRF-193 damage is induced majorly in S and G2 phases of the cell cycle, where HR is usually the predominant pathway choice for DSB repair in the repetitive sequences of heterochromatin^25^. Similarly, inhibition of Rad52, which drives the repair for single strand annealing (SSA) and break induced replication (BIR), did not significantly alter the repair kinetics of ICRF-193 (Figure S3M). These results indicate that cells use NHEJ to repair damage by ICRF-193. To further validate our finding, we used Ku80 knock-out (Ku80KO) MEFs, where the NHEJ pathway is completely abolished, and found that cells are incapable of repairing ICRF-193 dependent damage, even after 6 hours of drug release (Figure S3N).

These results demonstrate that ICRF-193 leads to the induction of DSBs in heterochromatin, which are repaired by NHEJ.

### ICRF-193-induced DNA damage in interphase jeopardizes genome integrity

DNA damage can be lethal for cells since it can severely affect genomic stability. DSBs are the most toxic type of lesions, which can enhance the chances of occurring translocations and impair proper chromosome segregation. Therefore, we next sought to investigate the consequences of the ICRF-193-induced DNA damage, observed in heterochromatin during interphase, on genomic instability. We performed fluorescence *in situ* hybridization (FISH) to visualize major satellite repeats in metaphase spreads from NIH3T3 cells treated with ICRF-193 while in interphase and in which the drug had been removed to allow progression to mitosis (Figure S4A). We found several types of chromosomal abnormalities associated with heterochromatin malformations (Figure 4A). These include: i) chromocenter malformations where the pericentromeric regions have lost their round shape, or DNA breaks where the whole pericentromeric region was detached from the rest of the chromosome arm; ii) whole chromosome translocations, where two telocentric mouse chromosomes are conjugated at their centromeric sites giving rise to a single metacentric chromosome; iii) heterochromatin aggregates, where single indistinct masses formed from the chromocenters of different chromosomes; iv) heterochromatin DNA bridges forming between two neighboring chromocenters. The last two types of abnormalities are indicative of failed decatenation by Top2α at the pericentric region. Quantification of the metaphases revealed a significant increase for all the pericentric malformations by ICRF-193 (Fold increase over DMSO: 7.4 for malformations, 2.9 for Translocations, 4.3 for aggregates, 3.7 for bridges; Figure 4B).

**Figure 4:**
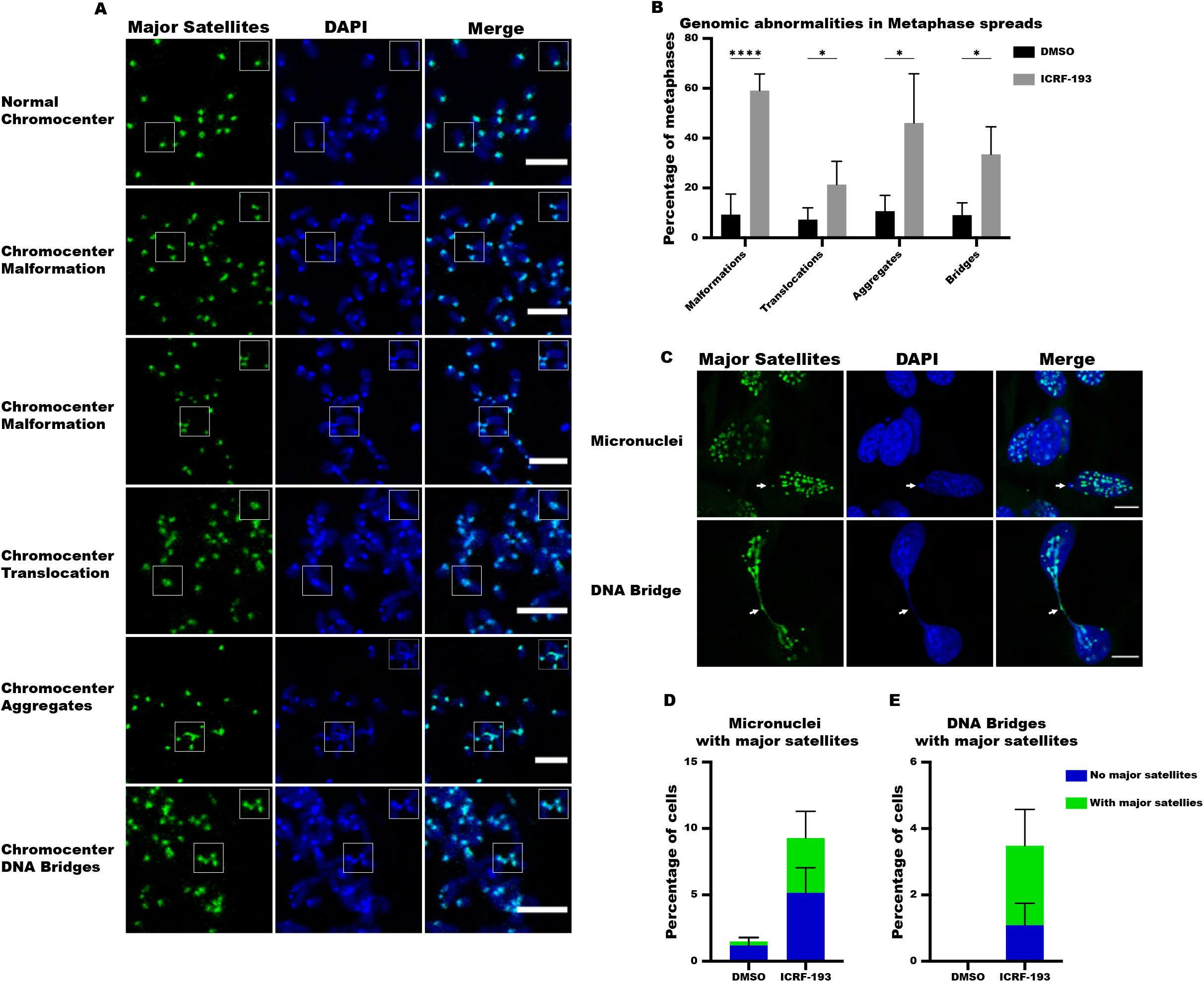
ICRF-193 induces genomic instability at heterochromatin. A) Representative confocal microscopy images of metaphase chromosome spreads from NIH3T3 cells. FISH was utilized to visualize Major Satellite repeats (green). DNA is visualized with DAPI (blue); examples of the different classes of genomic abnormalities detected are shown. Scale bars = 10μM B) Quantification of the abnormalities induced by DMSO or ICRF-193. C) Representative confocal microscopy images of micronuclei and DNA bridges formed in NIH 3T3 cells treated with DMSO or ICRF-193; Major Satellite repeats (green), and DNA (blue). Scale bars = 10μM D) Quantification of the percentage of cells with micronuclei containing Major Satellites DNA (green bar) or not (blue bar). At least 500 cells were counted per condition. E) Quantification of the percentage of cells with DNA bridges that contain Major Satellites DNA (green bar) or not (blue bar). At least 500 cells were counted per condition. All graph bars represent average values, and the error bars in the graphs represent the standard deviation from at least 50 metaphases per condition, and 3 independent replicates unless stated otherwise. Statistical significance presented by student t-test in 4B: (*) p<0.05, (**) p<0.005, (***) p<0.001, (****) p<0.0001

To investigate whether the cells carry these genomic aberrations to the next cell cycle, we quantified the percentage of cells with micronuclei that contain pieces of heterochromatin, and/or with DNA bridges that still connect two daughter nuclei in the next G1 phase (Figure 4C). We observed a 6.2-fold increase in the total number of micronuclei in general (significance: p<0.0001), and a 13.9-fold increase in the micronuclei containing major satellite repeat heterochromatin (Figure 4D). Furthermore, even though it is a rare event to find DAPI DNA bridges in the control conditions, a large increase was observed after releasing ICRF-193 (significance: p<0.01), and the majority (69%) were positive for major satellite repeat heterochromatin (Figure 4E).

Altogether, these results reveal that ICRF-193 induces damage at heterochromatin during interphase, which can lead to a significant level of genomic instability that is passed on to daughter cells.

### ICRF-193 has a differential effect in the dynamics of the Top2s in chromatin

To gain further insights into the mechanism by which Top2 inhibition leads to DNA damage, we investigated the impact of ICRF-193 on Top2α and Top2β localization and dynamics on heterochromatin. In accordance with previous reports^17,18^, we found that both Top2α and Top2β are physiologically enriched at sites of heterochromatin (Figure 5A). Interestingly, after treatment with ICRF-193 we observed that Top2α stays bound on DNA, and the number of cells with Top2α at heterochromatin is increased (Figure 5A and 5B). On the contrary, the percentage of cells with Top2β is reduced with approximately 50% of cells having lost the expression of Top2β after ICRF-193 treatment, compared to the control (Figure 5A, C).

**Figure 5:**
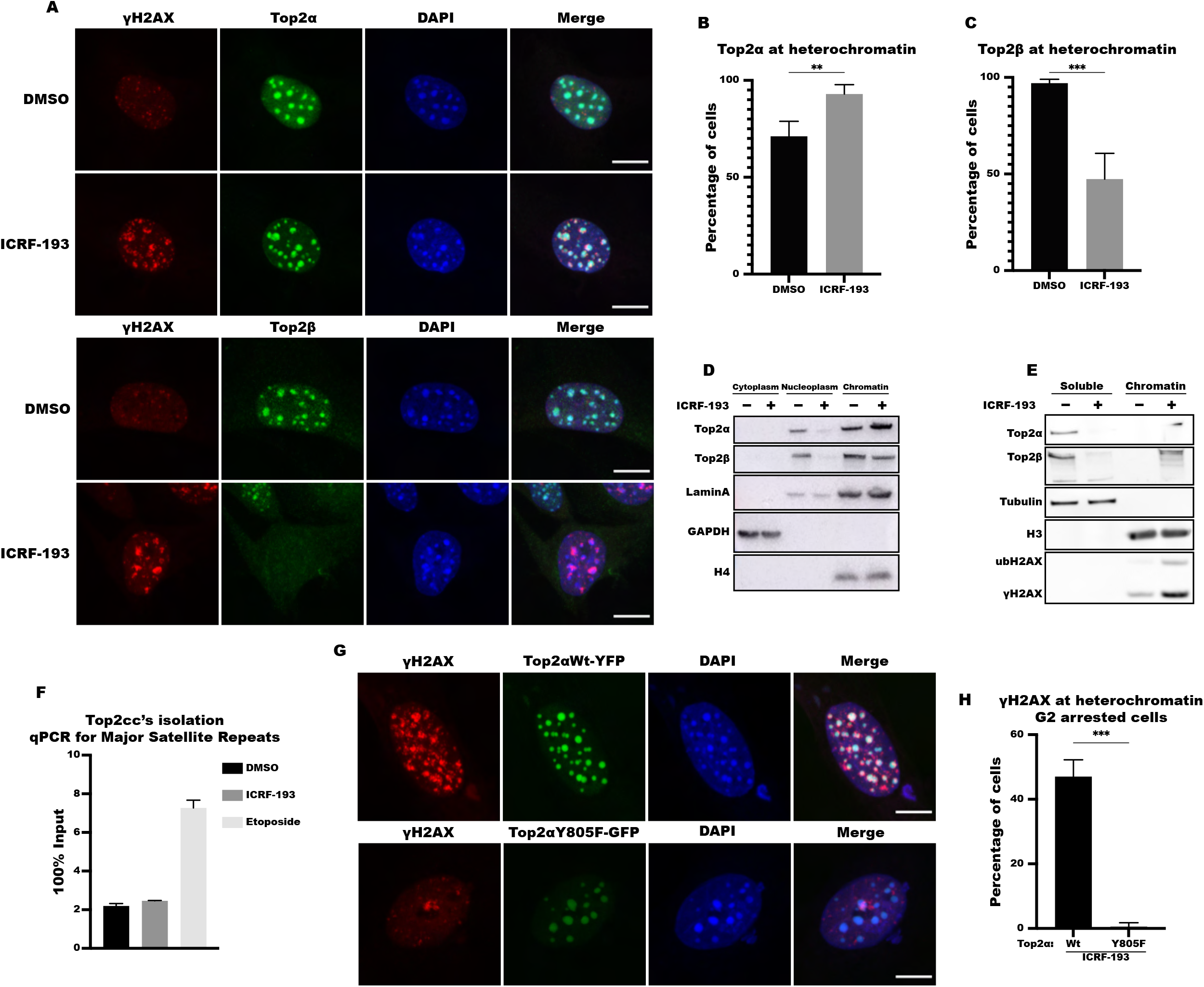
ICRF-193 affects the localization of Topoisomerase 2 α & β. A) Representative confocal microscopy images of γH2AX (red), Top2α (upper panel; green), Top2β (lower panel; green) and DAPI (blue) in NIH3T3 cells, treated with DMSO or ICRF-193. Maximum projection of z-stacks is shown. Scale bars = 10μM B-C) Quantification of NIH3T3 cells with (B) Top2α or (C) Top2β at heterochromatin. D-E) Western blot analysis of Top2α and Top2β in chromatin fractionations from NIH3T3 cells treated with DMSO or ICRF-193, with (D) 200mM or (E) 500mM NaCl containing buffer. F) Quantification of Top2-cc’s at Major Satellite repeats in NIH3T3 cells treated with DMSO and ICRF-193 or etoposide. G) Representative confocal microscopy images of γH2AX (red), Top2αWt-YFP (top; green), Top2αY805F-GFP (bottom; green) and DAPI (blue) in G2-arrested NIH3T3 cells, transfected with Top2αWt or Top2αY805F and treated with DMSO or ICRF-193. Maximum projection of z-stacks is shown. Scale bars = 10μM H) Quantification of the percentage of Top2αWt or Top2αY805F G2-arrested cells, after ICRF-193 treatment, with γH2AX at heterochromatin. I) Western blot analysis of GFP from chromatin fractionation of lysates from NIH3T3 cells transfected with Top2αWt-YFP or Top2αY805F-GFP and treated with DMSO as control (-) or ICRF-193 (+). For Western blots Lamin A, GAPDH, Tubulin, H3 & H4 were used as fraction specific loading controls. All graphs represent average values, and the error bars in the graphs represent the standard deviation from at least 50 cells per condition, and 3 independent replicates unless stated otherwise. Statistical significance presented by student *t*-test in 5B, 5C, and 5H: (*) p<0.05, (**) p<0.005, (***) p<0.001, (****) p<0.0001

To further corroborate these findings, we performed chromatin fractionation assays. The efficiency of these assays in separating the various protein fractions was verified based on the levels of lamin A or histones H3 and H4 found on chromatin, and GAPDH or Tubulin found in the soluble fractions (Figure 5D & 5E). When the fractionation was performed with a low salt containing buffer, we observed that ICRF-193 leads to an increase of Top2α on chromatin, and a downregulation of Top2β in both the nucleoplasm and on chromatin, as detected by immunofluorescence (Figure 5A & 5D). To assess to what extent ICRF-193 can trap the Top2 enzymes on the chromatin, we performed the fractionation assays with high salt containing buffer, in which all proteins that are not covalently linked or trapped on the DNA are removed and partition to the soluble fraction. Top2α and Top2β were found in the soluble fraction, in untreated conditions however, upon treatment with ICRF-193, the vast majority of Top2α stayed bound to chromatin (Figure 5E). This is also the case for the remaining Top2β fraction, which does not get degraded and remains bound to chromatin in the presence of ICRF-193 (Figure 5E). The induction of DNA damage was verified by γH2AX and ubH2AX (Figure 5E).

While Top2 poisons trap the Top2 enzymes when they are covalently linked to the DNA, Top2 catalytic inhibitors, such as ICRF-193, are reported to trap them in the closed clamp form around chromatin^26^. Since we find that ICRF-193 keeps the Top2 enzymes tightly linked to chromatin, even in high-salt conditions, we sought to test whether ICRF-193 induces Top2ccs using CC-seq^16^. This technique is powerful because it can detect even transient Top2ccs, which are not stabilized by chemical substances. Indeed, while treatment with etoposide led to the expected increase of Top2cc signal at major satellite repeats (NIH3T3 cells), satellite II, satellite III and centromeric repeats (U2OS cells), no increase was observed upon ICRF-193 treatment (Figure 5F, Figure S5A & S5B). Therefore, we conclude that ICRF-193 traps Top2 on heterochromatin in the closed clamp form, without inducing Top2ccs. To further support these findings, we performed FRAP analysis to determine how ICRF-193 affects the dynamics of Top2a in heterochromatin in the presence of ICRF-193. Indeed, we observed that ICRF-193 drastically decreases the mobility of Top2α fused to YFP, suggesting that catalytic inhibition of Top2α leads to its avid binding to chromatin (Video 1 and 2; Figure S5C).

To establish if the catalytic activity of Top2 is important for the induction of DNA damage by ICRF-193, we followed the formation of γH2AX foci at heterochromatin in G2-arrested NIH3T3 cells treated with ICRF-193, expressing either Top2α Wt or Top2αY805F, a dominant negative mutant which lacks the catalytic activity^27^. Interestingly, unlike control cells over-expressing Top2αWt, we failed to detect γH2AX foci at heterochromatin in cells over-expressing the mutated form of the enzyme (Figure 5G & 5H), suggesting that the catalytic activity of Top2α is necessary for the induction of damage by ICRF-193.

These results demonstrate that ICRF-193 leads to the prompt degradation of Top2β and the trapping of Top2α on heterochromatin, and that the catalytic activity of Top2α is necessary for DNA damage induction. Our results once again reveal the differential effect ICRF-193 has on the two Top2 isozymes.

### Sumoylation has an important role in the cellular response to treatment with ICRF-193

Sumoylation was shown to drive the removal of DNA-protein crosslinks (DPCs)^28-30^. By regulating the ubiquitination of proteins covalently trapped on the DNA, it can catalyze their proteasome-dependent degradation^28,30^. This was also shown to be the case for Top2ccs stabilized by etoposide, where the trapped Top2 enzymes are SUMOylated and subsequently ubiquitinated in order to be removed by proteasome-dependent degradation from the DNA^31^. In addition, inhibition of sumoylation or ubiquitination was reported to reduce the induction of γH2AX by etoposide^30,32^. Whether sumoylation plays a role in the removal of trapped Top2 from the DNA or in the induction of DDR and repair by ICRF-193 remains unknown.

To elucidate if sumoylation affects the induction of γH2AX, we treated NIH3T3 cells with the sumoylation inhibitor, ML-792 (Sumoi), or with DMSO as a control, prior to the addition of ICRF-193. We observed a significant reduction of γH2AX foci for all phases of the cell cycle (Figure 6A). Noticeably, the highest effect was found in G2 phase (15.8-fold reduction), where we also observed the highest induction of damage by ICRF-193 (Figure 1E, 1I, 1J, S1D, S1H, 2I and S2J). For this reason, we also tested Sumoi in NIH3T3 cells synchronized in G2 and confirmed the reduction of γH2AX foci (8.5-fold decrease; Figure S6A and S6B). To further validate this observation, we overexpressed Sumo1, Sumo2 and Sumo3 proteins, fused to RFP, in G2 arrested NIH3T3 cells. We observed an ICRF-193-dependent recruitment at heterochromatin in 5.1% of cells for Sumo1, 32% for Sumo2 and 36.6% for Sumo3. The recruitment was abolished in Sumoi treated cells (Figure S6C & S6D). Therefore, our data suggest sumoylation has an important role in the induction of γH2AX by ICRF-193.

**Figure 6:**
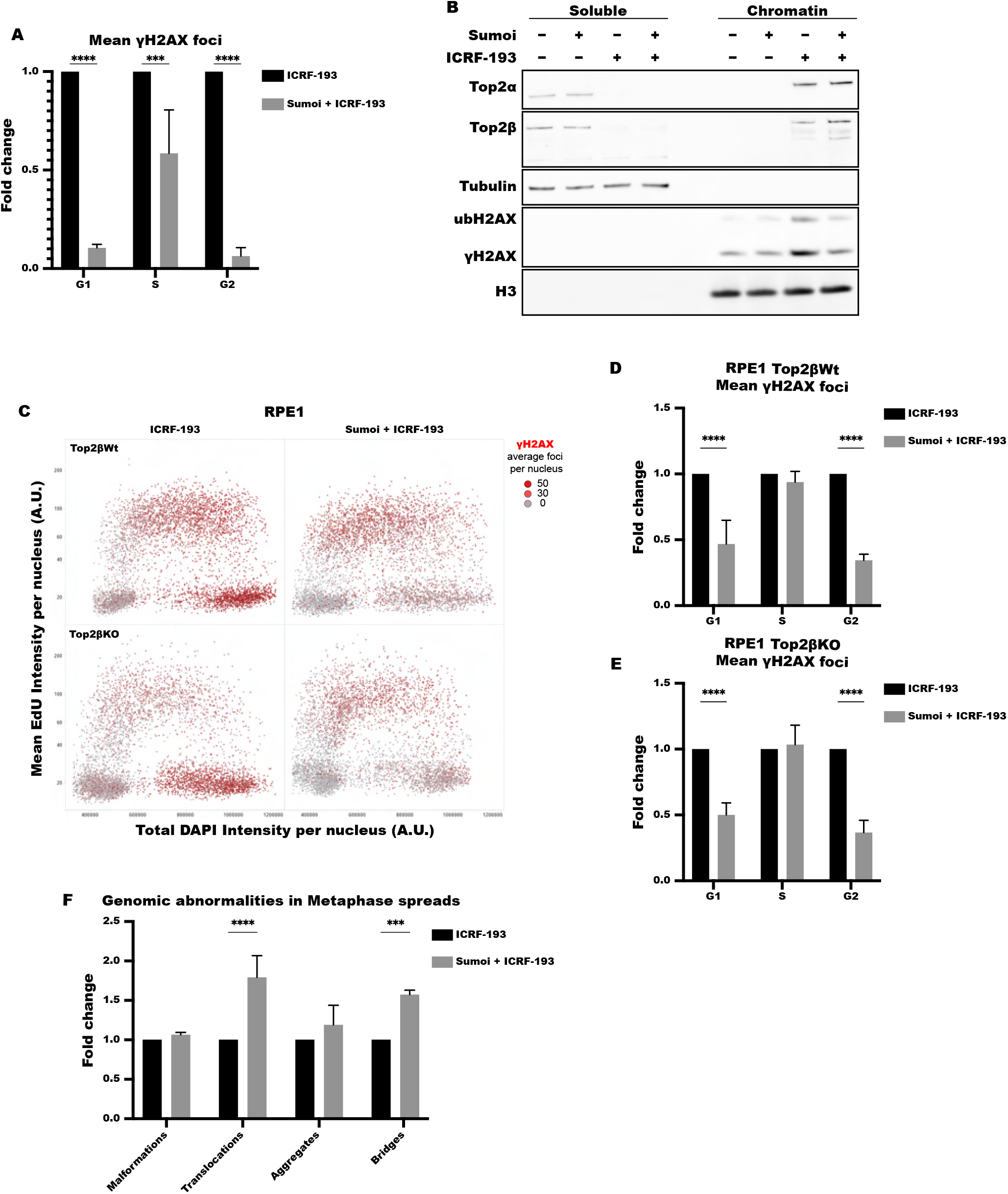
Sumoylation is necessary for ICRF-193 dependent DDR activation. A) Quantification of the γH2AX foci, in asynchronous NIH3T3 cells treated with DMSO or 5μΜ of ICRF-193, with pre-treatment with DMSO or ML-792 sumo inhibitor (Sumoi). B) Western blot analysis of Top2α and Top2β upon chromatin fractionations of lysates from NIH3T3 cells treated with ICRF-193, with high-salt (500mM NaCl) containing buffer. Cells were pre-treated with DMSO or Sumoi, prior to addition of ICRF-193. Tubulin & H3 were used as loading controls. C) QIBC plots of γH2AX in asynchronous Top2βWt and Top2βKO RPE1 cells pre-treated with DMSO or Sumoi and subsequently with ICRF-193. The cell cycle staging was performed based on EdU and DAPI intensity. Each dot represents a single cell, and the color of the dots represents the number of γH2AX foci per cell. D-E) Quantification of the mean number of γH2ΑΧ foci, in asynchronous (D) RPE1 Top2βWt or (E) RPE1 Top2βKO cells pre-treated with DMSO or Sumoi. F) Quantification of chromosomal abnormalities, as classified in Fig. 4A, induced by ICRF-193 after pre-treatment with DMSO or Sumoi. Graphs represent average values, and the error bars represent the standard deviation from at least 50 metaphases per condition, and 3 biological replicates. QIBC graphs in this figure represent average values, and the error bars represent the standard deviation from at least 1000 cells per condition, and 3 biological replicates unless stated otherwise. A.U.: Arbitrary Units. Statistical significance presented by = two-way ANOVA in 6A, 6D, 6E, 6F: (*) p<0.05, (**) p<0.005, (***) p<0.001, (****) p<0.0001

The effect on γH2AX was not due to changes in the levels of Top2α (Figure S6E & S6F). Furthermore, we confirmed that both Top2 isozymes remain trapped on chromatin by fractionation assay, with a slight increase in the levels of Top2α on chromatin upon pre-treatment with Sumoi (Figure 6B). Finally, FRAP analysis did not show any difference in the mobility of Top2α with or without the presence of Sumoi, thus further supporting our findings (Video 3; Figure S6G). Altogether, our results show that sumoylation has an important role in the induction of DNA damage response by ICRF-193, without affecting the localization of the Top2 enzymes.

Recently, sumoylation was shown to have a role in the degradation of Top2ccs, stabilized by topoisomerase poisons^30^. In agreement with this, we observed by both Western blot and microscopy analysis, that Sumoi prevents the degradation of Top2β in ICRF-193 treated NIH3T3 MEFs (Figure 6B, S6H and S6I). Hence, to test whether Sumoi prevents the ICRF-193 dependent γH2AX formation by inhibiting the degradation of trapped Top2s by the proteasome, and thus not revealing DSBs left behind by the cleaved DNA protein crosslinks, we used the proteasome inhibitor MG132. The proteasome inhibition led to a slight but significant increase of Top2α foci and an evident increase in the number of Top2β foci in G2 arrested NIH3T3 MEFs (Figure S6J & S6K). Furthermore, the impact of proteasome inhibition led to a significant decrease of γH2AX foci (1.5-fold) induced by ICRF-193 (Figure S6L), however it was not as potent as that of Sumoi (Figure S6B).

Since there was a significant rescue of the Top2β levels, we wanted to explore further whether the inhibition of ICRF-193-dependent Top2β degradation by Sumoi is responsible for the reduction of γH2AX foci. To this end, we pre-treated both Top2β Wt and KO RPE1 cells with the Sumoi, and subsequently treated them with ICRF-193 to compare how stabilization of Top2β would affect the induction of γH2AX. Treatment with Sumoi reduced the γH2AX induction in a similar way, both in the presence (2.9-fold decrease in G2 cells) or absence (2.7-fold decrease in G2 cells) of Top2β (Figure 6C, 6D, 6E and S6M). Hence, we conclude that Top2β degradation by ICRF-193 does not contribute to the γH2AX formation initiated by sumoylation signaling.

So far, we have shown that Sumoi reduces the γH2AX levels induced by ICRF-193, without reducing the trapping of Top2α around chromatin. Thus, sumoylation either reduces the levels of DNA damage or has an impact on the response and signaling to DNA damage, which might augment genomic instability. To investigate which scenario holds true, we performed metaphase spreads to check the genomic stability induced by ICRF-193, in the presence or absence of Sumoi. If Sumoi affects the induction of DNA damage, the combination of the two drugs should decrease the aberrations we found on metaphase chromosomes. However, if sumoylation plays a role in the DDR induced by ICRF-193, pre-treatment with Sumoi should increase the genomic instability even further, since the cells will carry severe damage and normally continue their cell cycle, as they will have no means to detect it. We found that pre-treatment with Sumoi led to an increase in both chromosomal translocations and chromocenter bridges induced by ICRF-193 (Figure 6F). These observations suggest that Sumoylation dependent mechanisms protect against genomic instability caused by Top2 inhibition.

### Ercc1-XPF endonuclease safeguards genomic stability by resolving inter-chromosomal DNA bridges

To understand the mechanism by which sumoylation protects against ICRF-193-induced genomic instability, we searched for DNA repair factors that are regulated at sites of damage in a SUMO-dependent manner. Ercc1 works in a complex with XPF to form a structure-specific endonuclease that takes part in Nucleotide excision repair (NER), DSB and interstrand crosslink (ICL) repair^33^. Its activity is regulated by the E3 SUMO ligase, Slx4^34^. Importantly, Slx4 SUMOylates the XPF subunit of the Ercc1-XPF complex^35^. The Ercc1-XPF complex also has a role in the repair of Top1 induced damage^36^. Thus, we hypothesized that Ercc1-XPF may also have a role in facilitating DNA repair at heterochromatin after damage induction by ICRF-193, possibly in a SUMO dependent manner.

Interestingly, Ercc1 is efficiently recruited to heterochromatin in cells treated with ICRF-193 (Figure 7A & 7B). Since the recruitment of Ercc1-XPF can be facilitated by different pathways, we aimed to elucidate the mechanism which regulates this process. For this reason, we used siRNAs to downregulate Slx4 (Figure S7A). In the presence of ICRF-193, the downregulation of Slx4 abolished the recruitment of Ercc1 to damaged heterochromatin, almost to the same extent as the downregulation of Ercc1 itself (Figure 7C). Therefore, we propose that Ercc1 is recruited to heterochromatin, in an Slx4-dependent manner. Since Slx4 exerts E3 SUMO ligase activity and SUMOylates the Ercc1-XPF complex^35^, and we have already determined the importance of sumoylation in the DNA repair and genomic stability of cells upon treatment with ICRF-193, we examined the recruitment of Ercc1 in the presence of both ICRF-193 and SUMOi. Indeed, inhibition of sumoylation prevented the recruitment of Ercc1 at heterochromatin upon treatment with ICRF-193 (Figure S7B).

**Figure 7:**
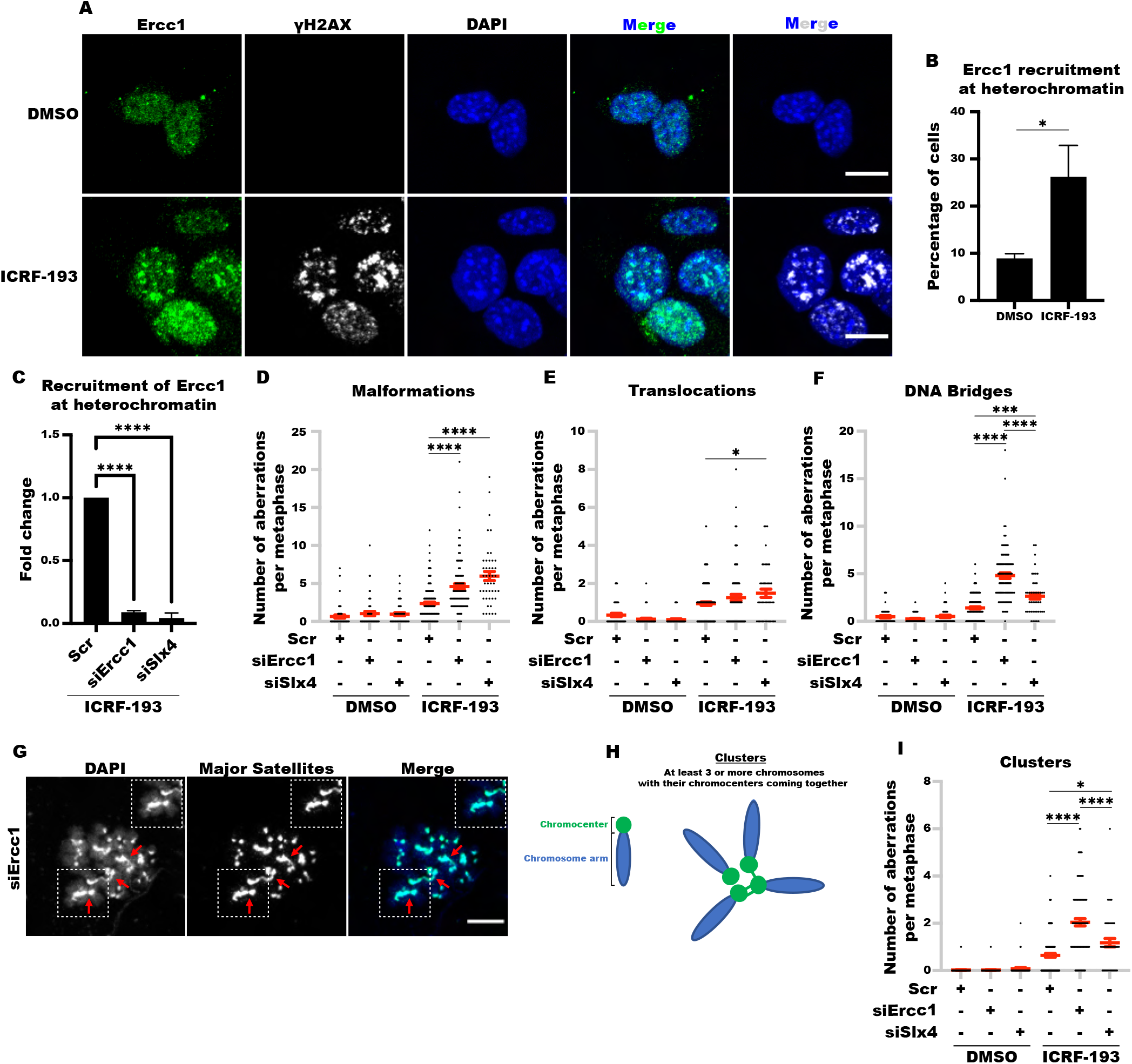
Ercc1-XPF endonuclease safeguards genomic stability by resolving inter-chromosomal DNA bridges. A) Representative confocal microscopy images of γH2AX (white), Ercc1 (green) and DAPI (blue) in NIH3T3 MEFs, treated with DMSO or 5μΜ of ICRF-193 for 4h. Maximum projection of z-stacks is shown. Scale bars = 10μM B) Quantification of cells with Ercc1 at heterochromatin. C) Quantification of asynchronous NIH3T3 cells with Ercc1 at heterochromatin, after scramble siRNA (Scr), siErcc1 or siSlx4. D-F) Quantification of (D) malformations, (E) translocations, and (F) DNA bridges per metaphase in cells transfected with the indicated siRNAs. The red lines indicate the mean values, and the error bars the standard error of the mean (±SEM). G) Representative confocal microscopy images of metaphase chromosome spreads from NIH3T3 cells transfected with siErcc1; Major Satellite repeats (green), DNA (blue); the location of clusters is indicated by the red arrows. Scale bars = 10μM H) A cartoon to represent the classification of cluster patterns. I) Quantification of chromosome clusters per metaphase after transfection with the indicated siRNAs. The red lines indicate the mean values, and the error bars the standard error of the mean (±SEM). Graphs represent average values, and the error bars in the graphs represent the standard deviation from at least 50 cells per condition, and 3 independent replicates unless stated otherwise. For the metaphase spreads in 7E, 7F, 7G, 7K and 7L, for the DMSO condition 1 biological replicate (n) is presented, with at least 50 metaphases counted for Scr, siErcc1 and siSlx4; for the ICRF-193 condition 3 biological replicates are presented, with metaphases counted per condition: n=1 Scr=50, siErcc1=22 and siSlx4=5; n=2 Scr=43, siErcc1=36 and siSlx4=19; n=3 Scr=35, siErcc1=35 and siSlx4=21. Statistical significance presented by student t-test in 7B, and One-way ANOVA in 7C. Statistical significance presented by One-way ANOVA in 7D, 7E, 7F, and 7I, only between ICRF-193 treated where 3 biological replicates were performed: (*) p<0.05, (**) p<0.005, (***) p<0.001, (****) p<0.0001

To decipher the potential role of Ercc1 at damaged heterochromatin, we assessed whether depletion of Ercc1 affects the genomic stability of cells treated with ICRF-193. To this end, and similarly to figure 4A, we quantified the number of malformations, DNA bridges and translocations in the presence of DMSO or ICRF-193, while depleting Ercc1 or Slx4. Indeed, treating NIH3T3 cells with ICRF-193 in the absence of Ercc1 or Slx4, further increased the number of Malformations or translocations, which are indicative of DBSs (Figure 7D and 7E). In addition, knock down of Ercc1 or Slx4 also led to an increase of DNA bridges in the presence of ICRF-193 when compared to Scr (Figure 7F). Therefore, these results indicate that Ercc1 and Slx4 have a role in preserving ICRF-193-induced genomic stability by resolving DNA bridges and preventing chromosome breaks.

The observed increase of DNA bridges related to heterochromatin in the absence of Ercc1 or Slx4 indicates elevated catenation of pericentromeric heterochromatin among clustered chromosomes. Hence, to unravel if there is an overall increase of inter-connected chromosomes at the level of mitosis, after damage induction by ICRF-193 in the absence of Ercc1 or Slx4, we quantified the number of chromosome clusters between 3 or more chromosomes with inter-chromosomal DNA bridges per metaphase (Figure 7G & 7H). In the presence of ICRF-193, the number of clusters significantly increased for both siErcc1 and siSlx4 over the control (Figure 7I), thus indicating that there are more chromosome clusters with unresolved inter-chromosomal DNA bridges in the absence of these two proteins. Overall, these results reveal a novel role of the Ercc1-XPF complex and Slx4 in the repair of DNA lesions resulting from Top2 trapped around chromatin, where they catalyze the resolution of inter-chromosomal DNA bridges forming between clustered regions of heterochromatin.

In conclusion, our study has uncovered that inhibition of Topoisomerase 2 activity has a multifaced impact on the integrity of heterochromatin and repetitive DNA and sheds light onto the distinct cellular outcomes of Top2 poisons and inhibitors.

## Discussion

Repetitive DNA is often packaged in heterochromatin to maintain its integrity, which is necessary for cell viability and fitness. Centromeric and pericentromeric repeats are essential for proper chromosome segregation, rDNA for growth, and LINEs and other retroelements for the silencing of transposable elements^37-40^. There has been a substantial progress in sequencing and bioinformatic approaches to aid the precise mapping of repetitive elements^40,41^. Nevertheless, it remains unclear whether their repetitive nature, which makes them prone to the formation of secondary structures, their chromatin status, and their organization to form clusters inside the nucleus leads to elevated topological stress, which can impede DNA replication or chromosome segregation and can jeopardize repeat integrity. Here, we have discovered that repetitive DNA is uniquely sensitive to the inhibition of Topoisomerase 2 activity by the catalytic inhibitor ICRF-193. We demonstrated using microscopy and ChIP-sequencing that ICRF-193 leads to the induction of DNA damage at repetitive regions (Figure 3B-3G, Table S1), and we failed to detect a consistent induction of DNA damage at any other region of the genome (Figure 3F and 3G). Thus, we propose ICRF-193 preferentially induces DNA damage at repetitive sequences, without significantly affecting euchromatin.

Top2 poisons, such as etoposide, induce DNA damage throughout the nucleus (Figure 1A and 1B), with a preference for CTCF binding regions and active promoters/genes^16,19,21^. This prompts the questions of why is there such a striking difference between poisons and inhibitors, and why does treatment with ICRF-193 lead to DNA damage only in repetitive DNA? One highly likely explanation for the difference between poisons and inhibitors could be that the DNA damage induced by etoposide and ICRF-193 depends on distinct Top2 isozymes. Although DNA damage induced by etoposide has been shown to depend on both Top2s^21^, etoposide induces DNA damage in all stages of the cell cycle and particularly in G1 where primarily Top2β is expressed, but Top2α is not, suggesting that the trapping of Top2β has a major role in the genome instability induced by Top2 poisons. In agreement, DNA breaks induced by etoposide correlate with sites of Top2β binding in the genome^19^ and depend on RNA pol II transcription^21^. On the other hand, our data demonstrate that ICRF-193 induced DNA damage depends on Top2α, and perfectly correlates with Top2α expression during the cell cycle. Thus, since the Top2 enzymes are naturally enriched at heterochromatin and repetitive sequences^17,18^, a plausible explanation for the specific action of ICRF-193 in these regions is because these genomic regions are either more compacted or tangled and they require elevated levels of Top2 activity to resolve the high topological stress. Therefore, the catalytic inhibition of Top2 activity could result in unresolved torsional stress of pericentric heterochromatin and subsequent occurrence of DNA breaks.

ICRF-193 does not induce DNA damage simply by inhibiting Top2 activity, since the depletion of Top2α rescues the induction of γH2AX foci in the presence of ICRF-193 (Figure 2A, 2B, 2E and 2F). Therefore, it is more probable that ICRF-193 induces DNA damage by trapping Top2α around the DNA. Nonetheless, the trapping of Top2α on chromatin is necessary but not sufficient to induce the formation of γH2AX foci at heterochromatin, as the ectopic overexpression of Top2α in G1 did not lead to a robust DNA damage induction in the presence of ICRF-193 (Figure S2L); instead, the Top2 trapping needs to interfere with DNA transactions occurring in S and G2 phases of the cell cycle.

Overall, our results point to the possibility that the mechanism by which ICRF-193 induces DNA damage in S and G2 phases are distinct. In xenopus egg extracts, the trapping of Top2α by ICRF-193 during S phase delayed the activation of replication origin clusters and replication fork progression, while also affecting the spacing of nucleosomes^42,43^. In accordance with these observations, our results in NIH 3T3 cells show a significant decrease in the percentage of early replicating cells and a non-significant increase in mid and late replicating cells (Figure S1M). Altogether, these observations show that the trapped Top2s by ICRF-193 act as physical barriers, obstructing the progression of the replisome, possibly leading to DNA damage by head on collisions.

The mechanism by which ICRF-193 induces damage preferentially at heterochromatin in G2 cells, could be due to decatenation defects of tangled repeats of the same or different chromosomes which cluster together in the nucleus^44^. The trapping of Top2 enzymes around chromatin in a closed clump can potentially affect the DNA topology leading to chromatin unwinding, thus causing the formation of secondary structures, such as inter-chromosomal full or semi-catenates. In accordance with this hypothesis, previous work demonstrated the stabilization of inter-chromosomal linkages between the rDNA loci of human chromosomes, which also cluster together physiologically, upon Top2 inhibition by ICRF-193^45^. Consequently, this hypothesis could explain the necessity of structure-specific endonucleases, such as Ercc1, in preserving genomic stability and preventing the formation of inter-chromosomal DNA bridges and chromosome breaks (Figure 7D-7F).

Another important aspect of this study is the establishment of DNA damage induction by ICRF-193 outside of mitosis. The initial assumptions were that ICRF-193 primarily acts by inhibiting the enzymatic activity of the Top2 enzymes, hence it would damage cells due to defective decatenation and thus, uneven chromosome segregation to daughter cells^15^. Here, we present an unbiased analysis of γH2AX induction in unperturbed replicating cells based on their cell cycle profile, where we show damage at heterochromatin can occur in all phases of the cell cycle, before mitosis begins (Figure 1D, 1E, 1H,1I and 1J).

We demonstrated that the catalytic activity of Top2α is necessary for the induction of γH2AX. This observation suggests that DNA damage is induced at the end of the Top2 catalytic cycle when both DNA strands are trapped inside the closed Top2 clump. Another possibility is that γH2AX marks the lesions induced by the Top2s themselves which are not promptly ligated due to the subsequent trapping into a clumped form.

The distinction between Topoisomerase poisons and catalytic inhibitors relies predominantly on the ability of a drug to stabilize topoisomerase cleavage complexes. ICRF-193 has long been perceived to act as a catalytic inhibitor of Top2 enzymes, since it does not increase the number of Top2ccs^26^. While accurate, this distinction initially led to the notion that the catalytic inhibitors are less cytotoxic than the Top2 poisons and they simply act by inactivating the Top2 enzymes^46^. Our work provides strong evidence that the absence of Top2 activity leads to a different cellular response than that of the catalytic inhibition of the enzyme. Furthermore, it is evident that there are several similarities between Top2 poisons and catalytic inhibitors. Drugs from both categories have been shown to induce the proteasome dependent degradation of the trapped Top2 enzymes, by utilizing a SUMO/ubiquitin related signaling pathway^30,47,48^. Moreover, the inhibition of sumoylation or ubiquitination leads to a decrease of γH2AX induction in either case^30,32^. Finally, treatment with both types of drugs can stimulate the formation of DSBs and lead to genomic instability^32^. These similarities could suggest that catalytic inhibitors may also act as “non-canonical poisons”, even though the mechanism of Top2 inhibition differs.

Given that ICRF-193 has such an effect on genomic stability, it could prove to be an effective drug for use in treating cancers. The clinical relevance is highlighted by the fact that ICRF-193 can efficiently induce DNA damage in replicating human cells (Figure 1H-1J). Furthermore, since we observed that cells commence the G2 phase without first completing DNA replication (Figure S1N), it is highly probable that they will move into mitosis with defects and therefore carry genomic instability into daughter cells. Similar to mouse cells, our experiments with different human cell lines validated the effects of ICRF-193, including cell cycle dependent induction of DNA damage and the recruitment of DSB repair factors, the absence of Top2ccs stabilization and the importance of the Top2α enzyme in the formation of γH2AX foci. ICRF-193 has yet to be used in the clinical setting for cancer treatment due to its extremely low aqueous solubility, however recent studies have shown that pre-drugs can be used to overcome this obstacle^49,50^. Thus, our findings can be used to analyze and predict the responses of human cells to treatment with ICRF-193. Importantly, the use of Top2 poisons, such as etoposide, in clinic was shown to be efficient in targeting cancer cells, but unfortunately to also to have off-target effects which can lead to secondary malignancies^51^. Consequently, the use of ICRF-193 in clinic could prove to be beneficial over etoposide since its effects are restrained to heterochromatin, suggesting that the risk of DSB formation and subsequent translocations involving potentially oncogenic genes in euchromatin is dramatically decreased, essentially diminishing the possibility of secondary malignancies such as AML to occur after treatment. In addition, we found that DSBs induced in heterochromatin by ICRF-193 are predominantly repaired by the NHEJ repair pathway. These observations are in agreement with previous screens performed on DT40 cells^52^. As a result, treatment with ICRF-193 could have an even greater effect on cancer cells which have lost key molecules of the NHEJ pathway, such as 53BP1 or DNAPK, or other important DNA repair factors presented in this study, such as Slx4, and the Ercc1-XPF complex.

## Figure legends

**Figure S1:**
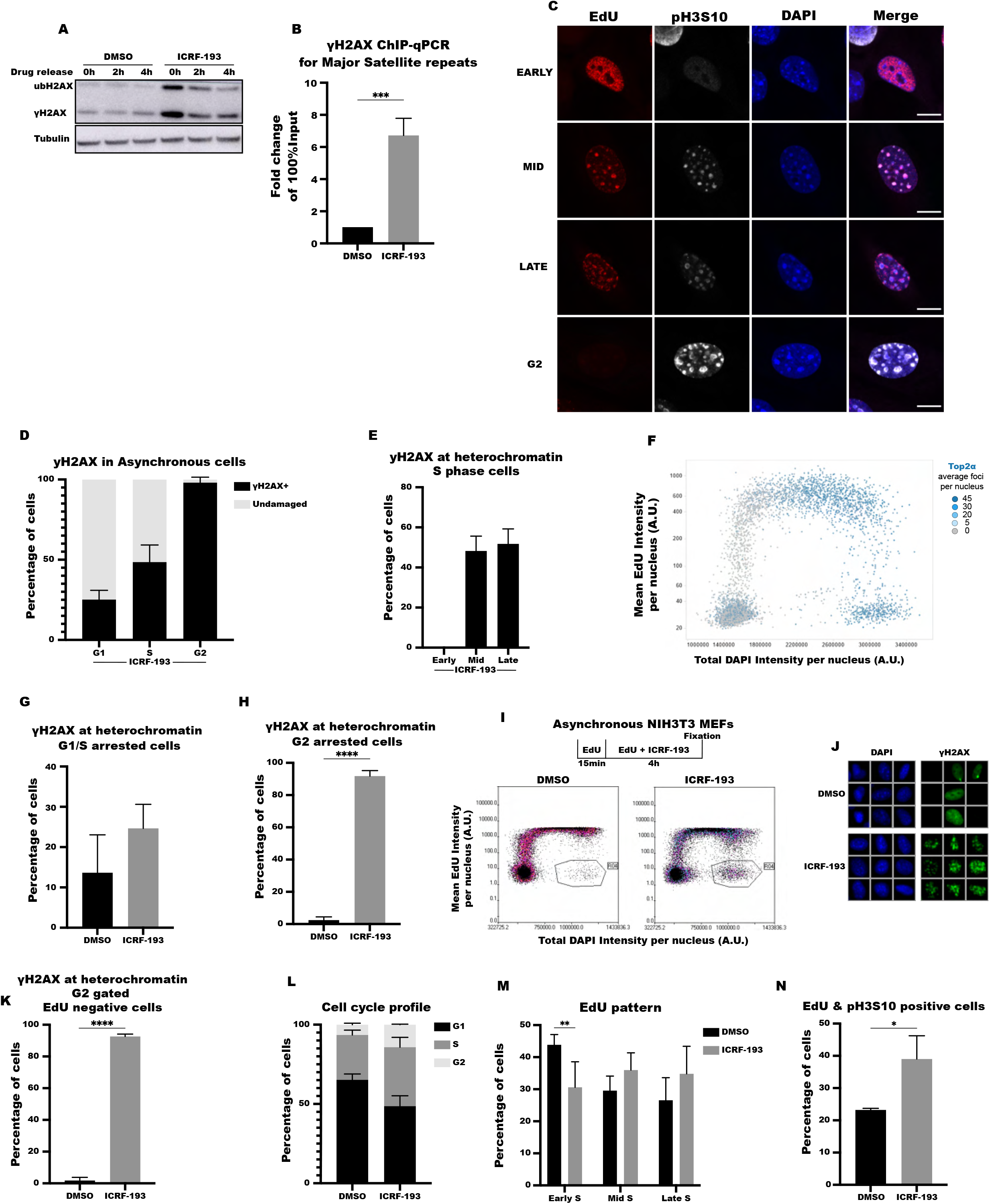
ICRF-193 induces damage at heterochromatin on a cell cycle basis. A) Western blot analysis of γH2AX, ubiquitinated H2AX (ubH2AX) and tubulin of cells treated with ICRF-193 and released in drug free medium for the indicated time. B) Chromatin immunoprecipitation analysis of γH2AX levels at pericentromeric heterochromatin over input (compared to no-antibody level) from 3 independent biological experiments, after treatment with ICRF-193. C) Representative confocal microscopy images of 5-Ethynyl-2′-deoxyuridine (EdU; red), DAPI (blue) and pH3S10 (white) in NIH3T3 cells for early, mid and late S (EdU positive) or G2 phase cells (EdU negative and pH3S10 positive cells). Maximum projection of z-stacks is shown. Scale bars = 10μM D) Quantification of NIH 3T3 cells with γH2AX foci localizing at heterochromatin regions in G1, S or G2. E) Quantification of NIH 3T3 cells with γH2AX foci localizing at heterochromatin regions in early, mid, or late S phase cells. F) QIBC-derived cell cycle plot of Top2α levels in NIH3T3 cells treated only with DMSO. Each dot represents a single cell, and the color of the dots represents the number of Top2α foci per cell in G1, S and G2 phases of the cell cycle. At least 1000 cells were analyzed per condition (See Materials and methods for details). A.U.: Arbitrary Units G) Quantification of cells with γH2AX foci localizing at heterochromatin in NIH3T3 cells synchronized in G1/S border by double thymidine block. H) Quantification of cells with γH2AX foci localizing at heterochromatin in NIH3T3 MEFs synchronized in G2 by RO-3306. I) Visual representation of the order of drug treatment applied in NIH3T3 cells (upper panel), to select the EdU negative and G2 positive cell population using QIBC (lower panel). J) Representative wide field images with the use of QIBC of γH2AX (green), and DAPI (blue) in NIH3T3 cells, after a 15min treatment with EdU and a subsequent 4h treatment of EdU+ICRF-193. The cells depicted are shown from the EdU negative G2 population as gated in Figure S1I. K) Quantification of the EdU negative G2 phase cells with γH2AX at heterochromatin, from Figure S1I. L) Analysis of the cell cycle profiles of NIH3T3 cells with or without ICRF-193 treatment. The data for these results were acquired from 3 biological replicates, using QIBC. At least 1500 cells were analyzed per condition. M) Quantification of S phase NIH3T3 cells based on their EdU patterns, by confocal microscopy N) Quantification of the NIH3T3 cells that are simultaneously positive for EdU and pH3S10 signal, using confocal microscopy. All graphs represent average values, and the error bars in the graphs represent the standard deviation from at least 50 cells per condition, and 3 independent replicates unless stated otherwise. Statistical significance presented by student t-test in S1B, S1G, S1H, S1K, and S1N and two-way ANOVA in S1M: (*) p<0.05, (**) p<0.005, (***) p<0.001, (****) p<0.0001

**Figure S2:**
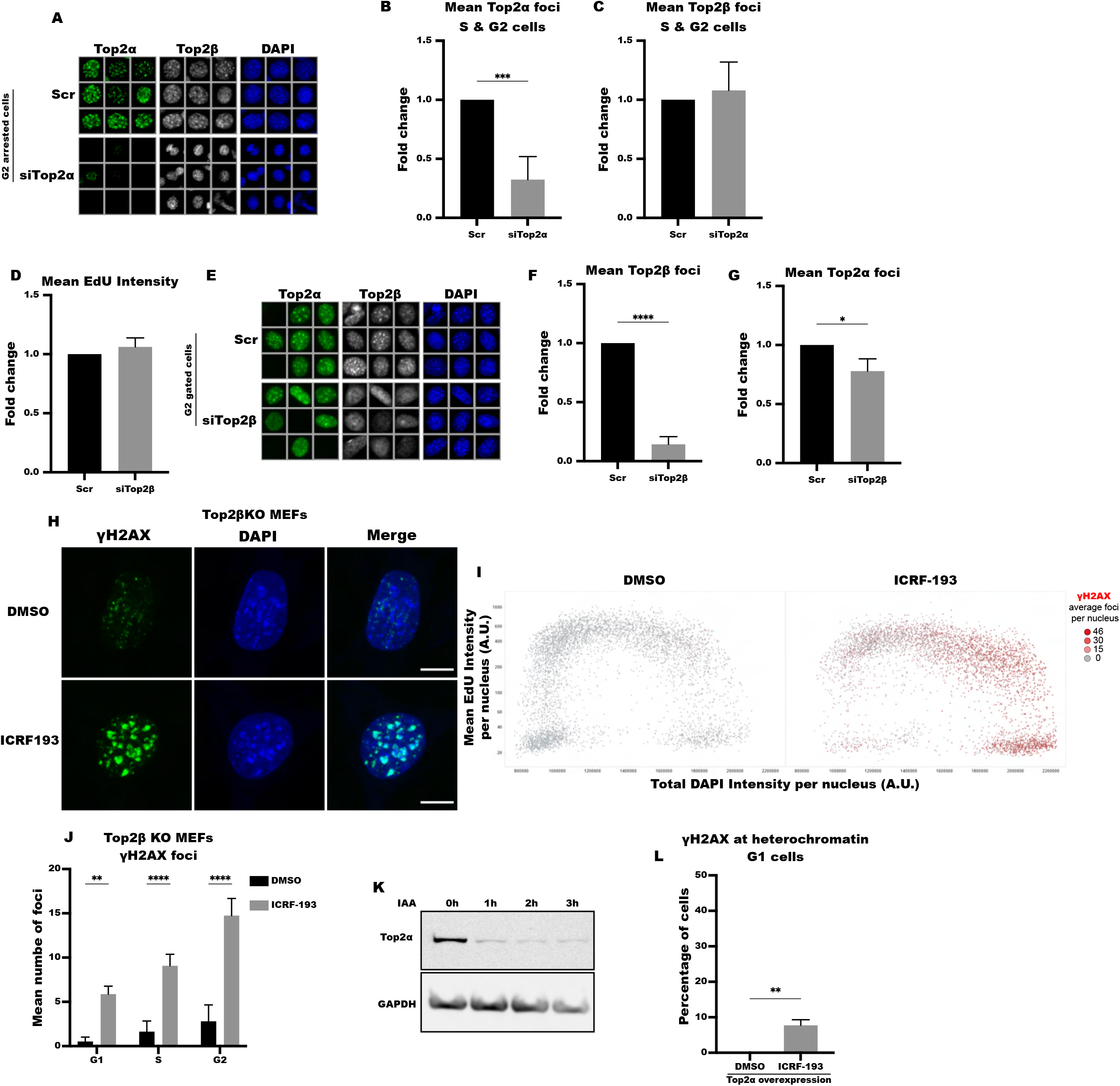
ICRF-193 induces damage by mainly targeting Top2α. A) Representative wide field images acquired using QIBC of Top2α (green), Top2β (white) and DAPI (blue) in NIH3T3 cells, after transfection with non-targeting (Scr) or Top2α siRNA. The cells were synchronized in G2 phase, with RO-3306. Scale bars for the indicative size of cells can be found in fig. 2A. B) Quantification of the Top2α foci presented in fig. S2A. Fold change of the Top2α foci in comparison to the control (Scr) condition is depicted. At least 320 cells were analyzed per condition. C) Quantification of the Top2β foci in Top2α knockdown and control conditions presented in fig. S2A, by the ScanR software. The graph bars display the fold change of the Top2β foci in comparison to the control (Scr) condition. D) Quantification of the EdU signal for comparison of the cell cycle progression, by the ScanR software. The graph bars display the fold change of EdU in comparison to the control (Scr) condition. E) Representative wide field images using QIBC of Top2α (green), Top2β (white) and DAPI (blue) in NIH3T3 cells, after transfection with non-targeting (Scr) or Top2β siRNA. The cells represent only G2 phase cells (based on the absence of EdU signal and their DAPI intensity), where Top2α expression is high. F) Quantification of the Top2β foci presented in fig. S2E. Fold change of the Top2β foci in comparison to the control (Scr) condition is shown. G) Quantification of the Top2α foci presented in fig. S2E after Top2β knockdown. H) Representative confocal microscopy images of γH2AX (green) and DAPI (blue) in Top2β KO MEFs treated with DMSO or ICRF-193 obtained by a confocal microscope. Maximum projection of z-stacks shown. Scale bars = 10μM I) QIBC plots of γH2AX levels in Top2β KO MEFs treated with DMSO or ICRF-193 as shown in fig. S2H. The cell cycle staging was performed based on EdU and DAPI intensity. Every dot represents a single cell, and the colour of the dots represents the number of γH2AX foci per cell. J) Quantification of the γH2AX foci presented in fig. S2H. K) Western blot analysis of Top2α in HCT116 OsTIR1/Top2αAID cells. Cells were treated with IAA for 0, 1, 2 or 3h. GAPDH levels were used as a protein-loading control. L) Quantification of the number of cells with γH2AX at heterochromatin in NIH3T3 cells overexpressing human Top2α-YFP and treated with DMSO or ICRF-193. Only G1 cells were selected for quantification, by choosing nuclei negative for EdU and pH3S10. The positive cells where counted based on the colocalization of the γH2AX foci with the Top2α-YFP foci forming the heterochromatin domains. Data presented from 3 biological replicates with 23-56 cells counted per replicate. Graph bars represent average values, and the error bars in the graphs represent the standard deviation from at least 1000 cells per condition, and 3 independent replicates unless stated otherwise. Statistical significance presented by student t-test in S2B, S2C, S2D, S2F, S2G and S2L, and two-way ANOVA in S2J: (*) p<0.05, (**) p<0.005, (***) p<0.001, (****) p<0.0001

**Figure S3:**
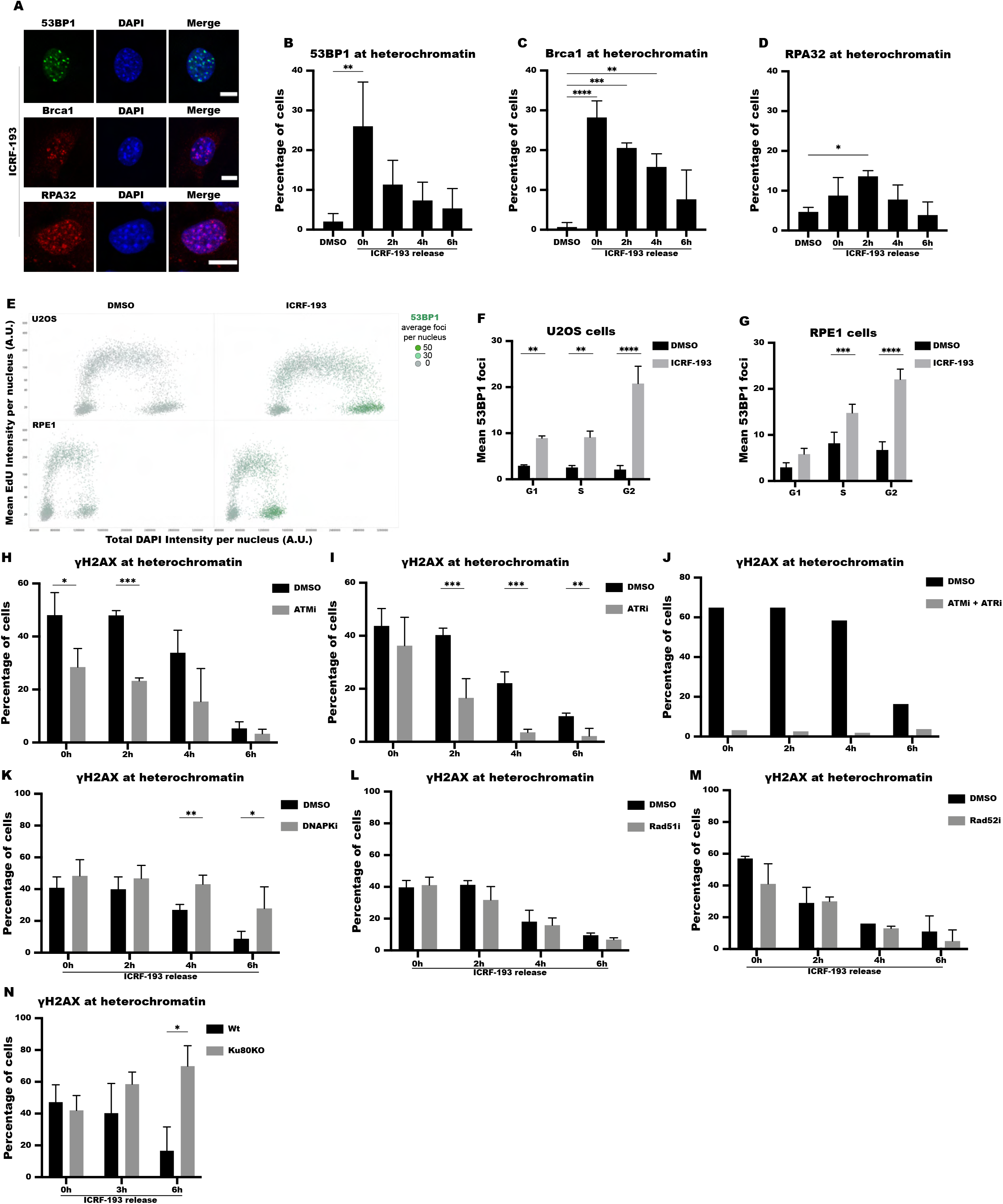
ICRF-193 induces DSBs in repetitive sequences. A) Representative confocal microscopy images of 53BP1 (green), Brca1 (red), RPA32 (red) and DAPI (blue) in NIH3T3 cells, treated with ICRF-193. Maximum projection of z-stacks is shown. Scale bars = 10μM. B-D) Quantification of number of cells with 53BP1 (B), Brca1 (C) and RPA32 (D) at heterochromatin, as shown in fig. S3A. E) QIBC plots of 53BP1 foci in U2OS (Upper panel) or RPE1 (Lower panel) cells treated with DMSO or ICRF-193. At least 1000 cells were counted per condition. The cell cycle staging was performed based on EdU and DAPI intensity. Each dot represents a single cell, and the color of the dots represents the number of 53BP1 foci per cell. F) Quantification of the mean number of γH2AX foci in U2OS cells. G) Quantification of the mean number γH2AX foci in RPE1 cells. H) Quantification of γH2AX induction in NIH3T3 cells treated with DMSO or ICRF-193 cells and pre-treated with or without ATMi. I) Quantification of γH2AX induction in NIH3T3 cells treated with DMSO or ICRF-193 cells and pre-treated overnight with or without ATRi. J) Quantification of γH2AX induction in NIH3T3 cells from one biological replicate, treated with DMSO or ICRF-193 cells and pre-treated with both ATMi & ATRi. K) Quantification of γH2AX induction in NIH3T3 cells pre-treated with DMSO or DNAPKi, and subsequently treated with ICRF-193. ICRF-193 was then removed from the culturing medium for the indicated time points. L) Quantification of γH2AX induction in NIH3T3 cells pre-treated with DMSO or Rad51i, and subsequently treated with ICRF-193. ICRF-193 was then removed from the culturing medium for the indicated time points. M) Quantification of γH2AX induction in NIH3T3 cells from 2 biological replicates, pre-treated with DMSO or Rad52i, and subsequently treated with ICRF-193. ICRF-193 was then removed from the culturing medium for the indicated time points. N) Quantification of γH2AX induction in in Wt or Ku80KO MEFs treated with ICRF-193, which was subsequently removed for the indicated time points. Graphs represent average values, and the error bars in the graphs represent the standard deviation from at least 50 cells per condition, and 3 independent replicates unless stated otherwise. Statistical significance presented by one-way ANOVA for S3B, S3C, S3D, two-way ANOVA for S3F and S3G and student *t*-test for S3H, S3I, S3K, S3L, S3N: (*) p<0.05, (**) p<0.005, (***) p<0.001, (****) p<0.0001

**Figure S4:**
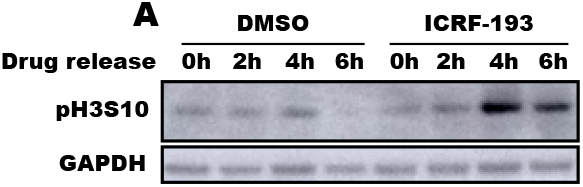
ICRF-193 induces genomic instability at heterochromatin. A) Western blot analysis of pH3S10, and GAPDH of cells treated with DMSO or ICRF-193 and released in drug free medium for the indicated time.

**Figure S5:**
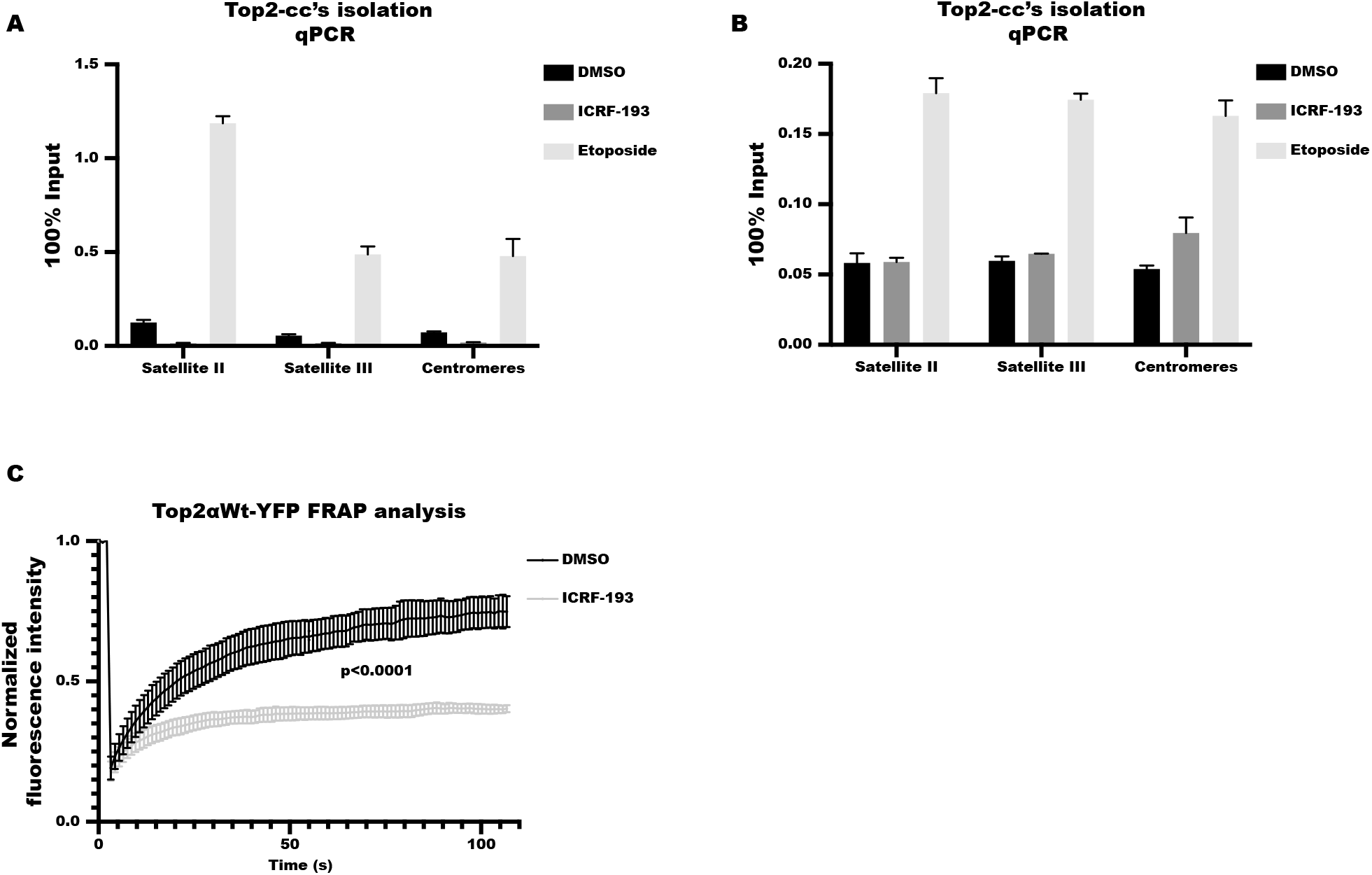
ICRF-193 affects the localization of Topoisomerase 2 α & β. A-B) Quantification of Top2-cc’s at Satellite II, Satellite III and centromeric repeats in U2OS cells treated with DMSO or ICRF-193 or etoposide, with (A) representing the first and (B) the second biological replicate. The cells were synchronized in G2 phase, with RO-3306. Graphs represent average values, and the error bars represent technical replicates from 1 biological replicate. C) FRAP analysis of Top2α Wt mobility in NIH3T3 cells transfected with Top2αWt and treated with DMSO or ICRF-193. Graph curves for FRAP results represent average values, and the error bars represent the standard error of the mean from at least 20 cells per condition, and 3 biological replicates. Statistical significance is presented by one-way ANOVA test in S5C.

**Figure S6:**
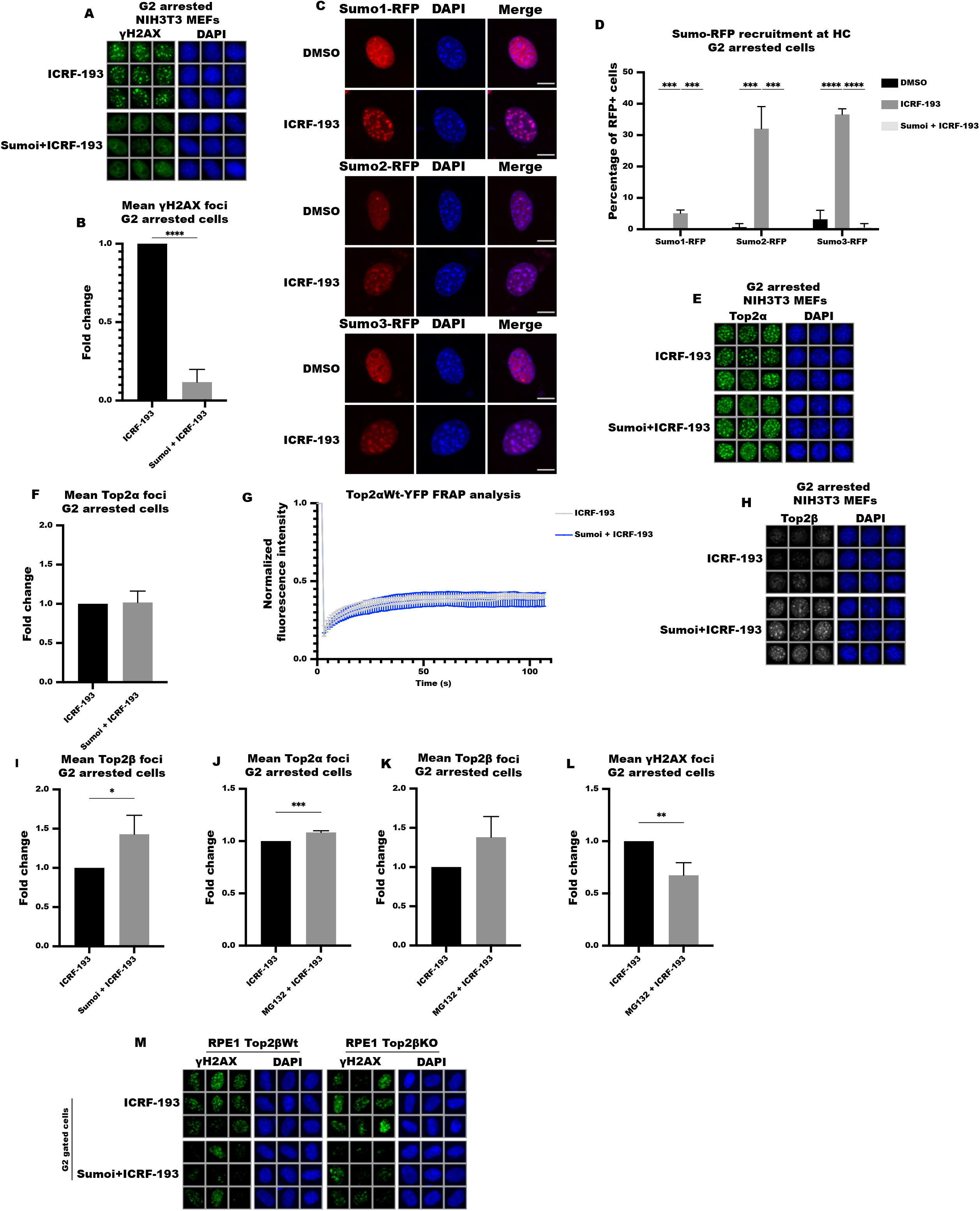
Sumoylation is necessary for ICRF-193 dependent DDR activation. A) Representative wide field images with the use of QIBC of γH2AX (green) and DAPI (blue) in NIH3T3 cells synchronized in G2 phase, treated with ICRF-193 and pre-treated with DMSO or ML-792 sumo inhibitor (Sumoi). The cells were synchronized in G2 phase, with RO-3306. B) Quantification of the γH2AX foci, in G2 arrested NIH3T3 cells treated with ICRF193, and pre-treated with DMSO or Sumoi. C) Representative confocal microscopy images of RFP-Sumo1, RFP-Sumo2 and RFP-Sumo3 (red) and DAPI (blue) in G2-arrested NIH3T3 cells, transiently transfected with Sumo-1, 2 or 3-RFP and treated with DMSO or ICRF-193. Maximum projection of z-stacks shown. Scale bars = 10μM. D) Quantification of cells with Sumo1, 2 or 3 at heterochromatin, as presented in fig. S6C. Graph bars represent average values, and the error bars in the graphs represent the standard deviation from at least 50 cells per condition, and 3 independent replicates. E) Representative wide field images with the use of QIBC of Top2α (green) and DAPI (blue) in NIH3T3 cells synchronized in G2 phase, pre-treated with DMSO or Sumoi prior to the addition of ICRF-193. The cells were synchronized in G2 phase, with RO-3306. F) Quantification of the mean number of Top2α foci, in G2 arrested NIH3T3 cells pre-treated with DMSO or Sumoi. G) FRAP analysis of Top2α Wt mobility in NIH3T3 cells transfected with Top2αwt-YFP and treated with ICRF-193, after pre-treatment with DMSO or Sumoi. Graph curves represent average values, and the error bars represent the standard error of the mean from at least 20 cells per condition, and 3 biological replicates. H) Representative wide field images with the use of QIBC of Top2β (white) and DAPI (blue) in NIH3T3 MEFs synchronized in G2 phase, pre-treated with DMSO or Sumoi prior to addition of ICRF-193. The cells were synchronized in G2 phase, with RO-3306. I) Quantification of the Top2β foci, in G2 arrested NIH3T3 cells pre-treated with DMSO or Sumoi, prior to addition of ICRF-193. J-L) Quantification of the Top2α (J), Top2β (K) or γH2AX (L) foci, by the ScanR software, in G2 arrested NIH3T3 cells pre-treated with DMSO or MG132, prior to addition of ICRF-193. M) Representative wide field images with the use of QIBC of γH2AX (green) and DAPI (blue) in G2-gated RPE1 Top2βWt and Top2βKO cells, pre-treated with DMSO or Sumoi prior to addition of ICRF-193. Graphs represent average values, and the error bars in the graphs represent the standard deviation from at least 450 cells per condition, and 3 independent replicates unless stated otherwise. Statistical significance presented by student *t*-test in S6B, S6F, S6I, S6J, S6K, S6L and one-way ANOVA in S6D, S6G: (*) p<0.05, (**) p<0.005, (***) p<0.001, (****) p<0.0001

**Figure S7:**
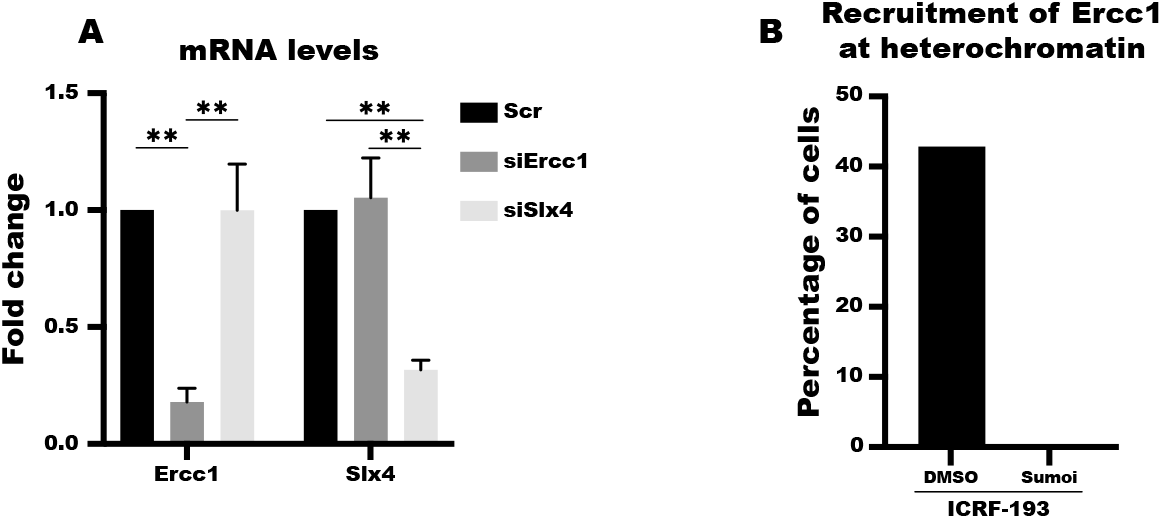
Ercc1-XPF endonuclease safeguards genomic stability by resolving inter-chromosomal DNA bridges. A) Quantification of Erccc1 or Slx4 mRNA levels by qRT-PCR analysis after transfection of the indicated siRNAs. Graph bars represent average values, and the error bars in the graphs represent the standard error of the mean (SEM) from 3 independent replicates. Statistical significance presented by one-way ANOVA in S7A: (*) p<0.05, (**) p<0.005, (***) p<0.001, (****) p<0.0001 B) Quantification of cells with Ercc1 at heterochromatin as shown in fig. 7A, in the presence of ICRF-193 with or without pre-treatment with Sumoi. Data presented from a single biological replicate.

## Materials & Methods

**Table 1.**

| siRNA |  |  |  |
| --- | --- | --- | --- |
| Target | Reference | Company | Type |
| Non-targeting | D-001810-10 | Dharmacon | ON-TARGETplus SMARTpool |
| siTop2α | L-043916-01-0005 | Dharmacon | ON-TARGETplus SMARTpool |
| siTop2β | L-042134-00-0005 | Dharmacon | ON-TARGETplus SMARTpool |
| siErcc1 | L-045170-01-0005 | Dharmacon | ON-TARGETplus SMARTpool |
| siSlx4 | L-057081-01-0005 | Dharmacon | ON-TARGETplus SMARTpool |

**Table 2.**
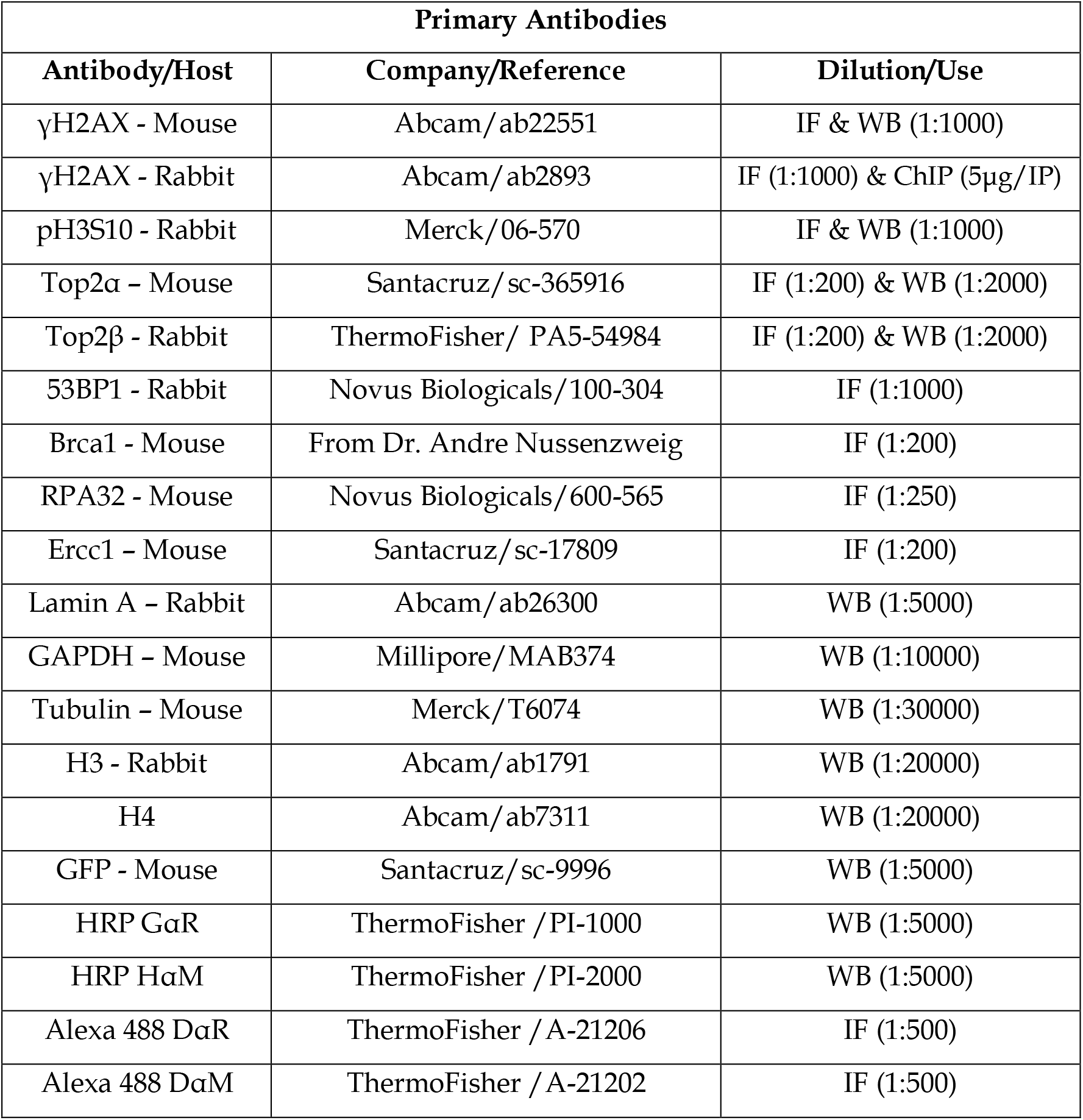
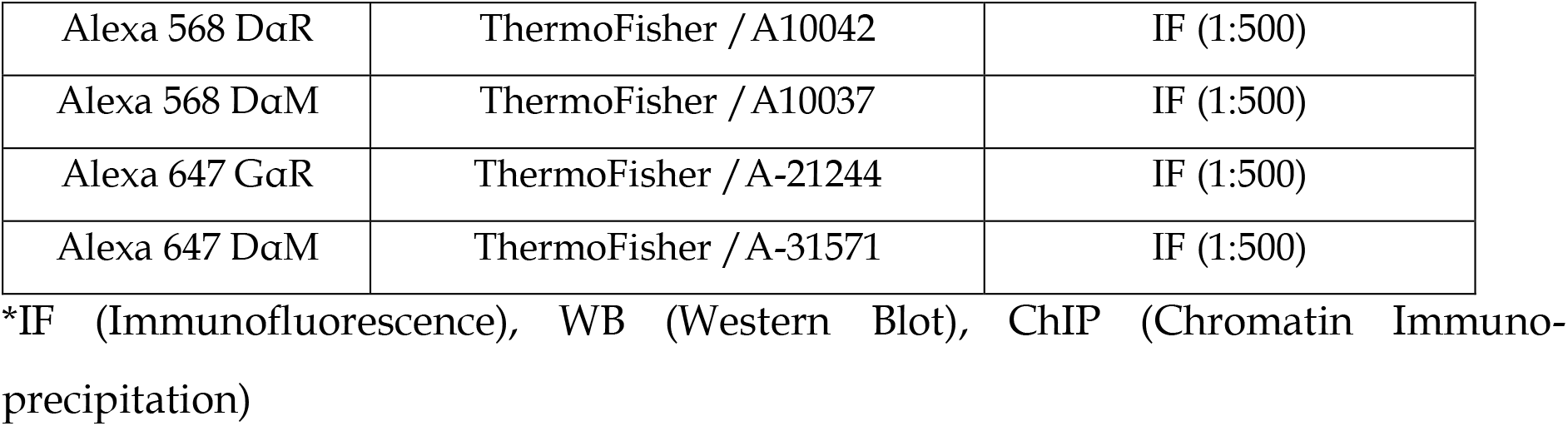

**Table 3.**

| Plasmids used |  |
| --- | --- |
| Top2αWt-YFP | CMVp-humanTop2αWt-YFP |
| Top2αY805F-GFP | Gift from Dr. Andy Porter |
| Major Satellite repeats | Gift from Dr. Maria-Elena Torres Padilla |
| pmRFP-Sumo1 | Gift from Dr. Steve Jackson |
| pmRFP-Sumo2 | Gift from Dr. Steve Jackson |
| pmRFP-Sumo3 | Gift from Dr. Steve Jackson |

**Table 4.**
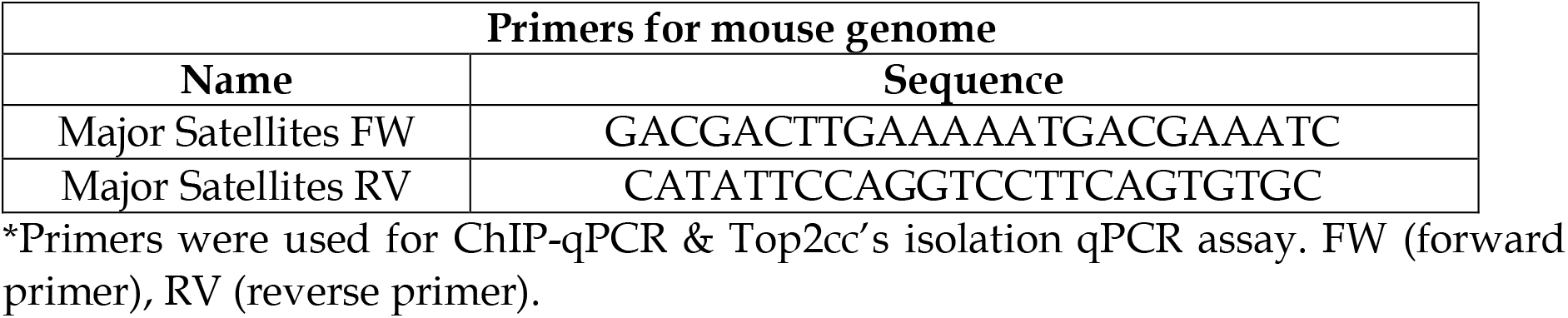

**Table 5.**
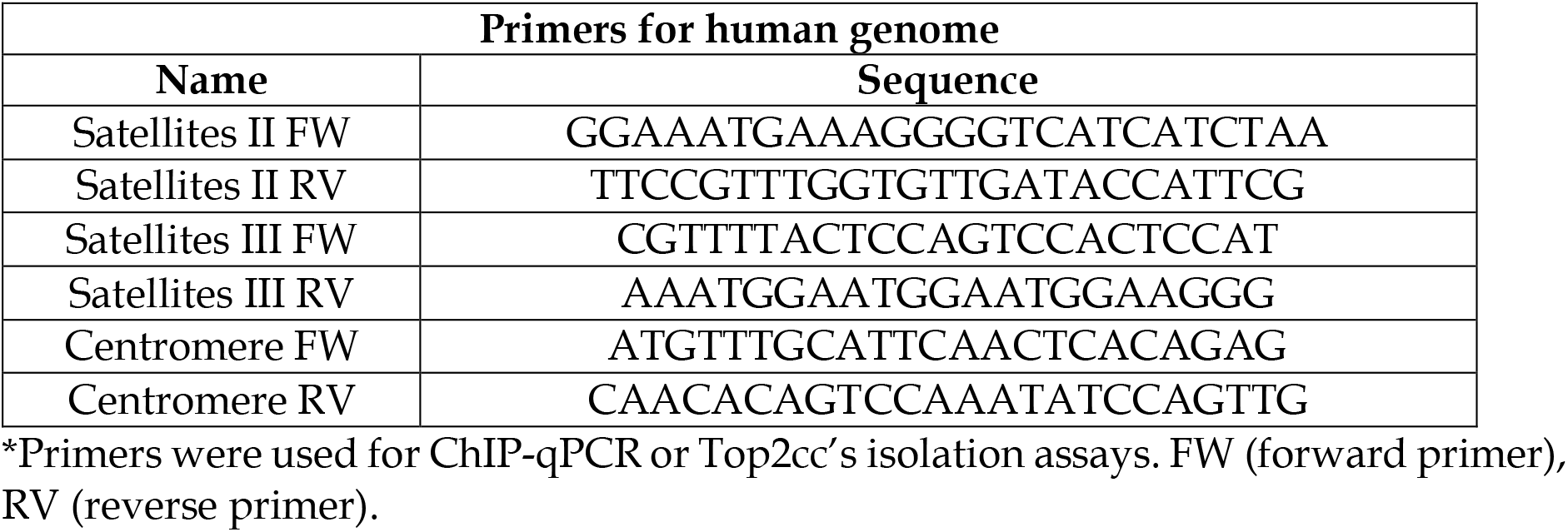

### Cell culture

All cell lines, unless stated otherwise, were obtained from the cell banks of Genome Damage and Stability Centre (GDSC) or Institut de Génétique et de Biologie Moléculaire et Cellulaire (IGBMC) and cultured in a humidified incubating chamber with 5% CO_2_ at 37 °C and were passaged before they reached 90% confluency. Human U2OS and mouse embryonic fibroblast (MEFs) cell lines were grown in Dulbecco’s Modified Eagle Medium (DMEM, Gibco), supplemented with 10% Foetal Calf Serum (FCS), 100 U/ml Penicillin (Gibco) and 100 U/ml Streptomycin (Pen/Strep) and 2mM L-Glutamine (Gibco). The Ku80 knock-out (Ku80KO) MEFs were a kind gift from Dr. Andre Nussenzweig’s lab^53^. Human RPE1-hTERT cells were grown in DMEM/F12 (Gibco) supplemented with 10% FCS, 100 U/ml Pen/Strep, 2 mM L-Glutamine and 125 μg/ml hygromycin. NIH3T3 MEFs were cultured in DMEM supplemented with 10% Newborn Calf Serum (NCS), 100 U/ml Pen/Strep and 2 mM L-Glutamine. HCT116 cells expressing OsTIR1 and Top2α-mAID where a kind gift from Dr. Christian Thomas Friberg Nielsen^14^; they were grown in DMEM, supplemented with 10% FCS, 100 U/ml Pen/Strep, 2 mM L-Glutamine and were selected with 1 μg/ml Puromycin, 7.5 μg/ml blasticidin and 125 μg/ml hygromycin. Selection antibiotics where excluded for experimental cell treatments. Depletion of Top2α was achieved by addition of 500μM of indole acetic acid (IAA) (Merck; I5148-2G) for 1h, prior to the addition of ICRF-193.

### Cell treatments

Unless stated otherwise, cells were treated with 5μM of ICRF-193 (Merck; I4659-1MG) for 4h. For direct comparison with ICRF-193, cells were treated with 5 μM etoposide (Merck; E1383) for 4h for the immunofluorescence assay, and with 50 μM for 30’ for the Top2cc’s isolation assay. For synchronization of cells in G1/S border, double thymidine (Sigma; T1895) block was used, cells were treated with 2 mM thymidine for 18h, released in drug-free media for 9h and treated with 2 mM thymidine for 16h prior to addition of ICRF-193. For synchronization of cells in G2, cells were treated with 10 μM of the CDK1 inhibitor, RO-3306 (Calbiochem; 217699) for 24h before fixation/collection of cells. The sumoylation inhibitor, 10μM of ML-70292 (Axon Medchem; 3109), or DMSO as negative control, were added for 90min prior to addition of ICRF-193. The proteasome inhibitor, 20μM of MG132 (BIO-TECHNE; 1748), or DMSO as negative control, were added for 1h prior to addition of ICRF-193. For the inhibition of ATM, 20 μM of Ku-60019 (Selleckchem; S1570), or DMSO were used for 1h prior to addition of ICRF-193. For the inhibition of ATR, 0.5 μM of VE821 (Calbiochem; 504972), or DMSO were added to cell media overnight prior to addition of ICRF-193. For the simultaneous inhibition of ATM and ATR, VE821 (or DMSO) was first added in cell media overnight, and ATM (or DMSO) was then added 1h before addition of ICRF-193. For the inhibition of DNAPK, 20 μM of Nu7026 (Merck; N1537-5MG), or DMSO was added to cell media 1h prior to addition of ICRF-193. For the inhibition of Rad51, 20 μM of B02 (Calbiochem; 553525), or DMSO was added to cell media overnight prior to addition of ICRF-193. For the inhibition of Rad52, 20 μM of (-)-Epigallocatechin (Merck; E3768), or H_2_O was added to cell media 1h prior to addition of ICRF-193.

### siRNA transfection by RNAimax

siRNAs (Table 1) where reverse transfected, at a final concentration of 40 nM, with Optimem (Gibco) and Lipofectamine RNAimax reagent (ThermoFisher), according to the manufacturer’s instructions. Cell treatments were performed 48h after the transfection for the downregulation of Top2α and 72h after all other siRNAs.

### Plasmid DNA transfections with Lipofectamine 2000

Plasmid DNA (1-2 μg) was transiently transfected with Optimem (Gibco) and Lipofectamine 2000 (ThermoFisher) according to the manufacturer’s instructions, for 16h before treatments.

### Staining for Ethynyl-2’-deoxyuridine (EdU) Click-iT

For the detection of replicating cells, EdU (25 μΜ) was added to the culturing medium 30min before fixation. Subsequently, Click-iT EdU Alexa Fluor 594 Imaging Kit, or Pacific blue, (Invitrogen) was used, according to manufacturer’s instructions. After this step, the protocol for immunofluorescence staining was followed.

### Immunofluorescence

Protocol was followed according to previous publication^44^. In brief, cells were cultured on glass coverslips and fixed with 4% paraformaldehyde (PFA) in 1X PBS for 10min at room temperature (RT). For the visualization of RPA32, Brca1 and Ercc1 proteins pre-extraction was performed prior to RT fixation, with a 30 second incubation with ice cold 0.1% Triton in 1X PBS and a subsequent 4% PFA in 1X PBS for 10 min on ice. Next, cells were permeabilized in 0.5% Triton in 1X PBS for 10min, blocked in 5% Bovine Serum Albumin (BSA) for 1h, then incubated with the primary antibody and secondary antibody (**Table 2**) in 5% BSA/1x PBS/0.1% Tween), for 1h each at RT. Finally, nuclei were counterstained with 1 mg/ml of 4’,6-Diamidino-2-Phenylindole, Dihydrochloride (DAPI) for 10mins.

Images were captured and quantified with a confocal laser scanning microscope (TCS SP8; Leica or LSM 880; Zeiss), using a x63 objective for at least 50 cells per biological replicate. Brightness & contrast of each colour of the confocal microscope images was adjusted accordingly with the ImageJ/Fiji software (version 2.1.0/1.53c).

For quantitative image-based cytometry (QIBC)^54^, images were captured on a ScanR High-Content Screening Station, which includes an inverted fluorescence Olympus IX83 microscope and a CMOS camera, A 40x NA 0.6 LUCPLFLN dry objective was used, to acquire at least 1000 cells per biological replicate in identical settings for all samples within the same experiment and non-saturating conditions. Image analysis was performed using the ScanR software (Version 3.0.1). Nuclei identification and segmentation was performed based on DAPI intensity by an integrated intensity-based object detection module. Background correction was applied and foci segmentation with the use of a spot-detection module. Fluorescence intensities were quantified and portrayed as arbitrary units (A.U.). Cell cycle analysis was based on detection of cell Total DAPI intensity containing 2C or 4C DNA (G1 or G2 phase cells respectively) and levels of Mean EdU Intensity (S phase cells). The colour-coded scatter plots from asynchronous populations were created with the Spotfire software (version 7.0.1; TIBCO 7.11). A similar number of cells was used to compare sample conditions within the same experiment. Single cells were selected based on object area/Total DAPI intensity or circularity/ Total DAPI intensity.

### Fluorescence *In-Situ* Hybridization for interphase cells and metaphase spreads

#### Preparation of metaphase spreads for hybridization with FISH probe

Protocol was followed according to previous publication^44^. In brief, after 4h of ICRF-193 treatment, cells were washed once with PBS and colcemid (0.02 μg/ml, 15210040; ThermoFisher) was added to ICRF-193 free growing media for 4h. The cells were trypsinized and the cell pellet was resuspended in pre-warmed solution of 0.06M KCl and incubated at 37 °C for 30 mins. Next, the cell pellet was fixed in 3:1 ethanol (EtOH): glacial acetic acid solution, three times and left in the fixative solution overnight at 4°C. The following day, cell metaphases were spread on ice-cold wet glass slides and left to air dry.

#### Preparation of interphase cells for hybridization with FISH probe

After treatment with ICRF-193 (5μM) for 4h, cells were released in drug free culturing medium for 6h and fixed with 4% PFA. Subsequently, they were washed with PBS, permeabilized with 0.5% Triton.

#### Preparation of FISH probe for hybridization

For the hybridization step, metaphase spreads or interphase cells were treated with RNase A (100 μg/ml in 2X SSC) at 37 °C for 1h, dehydrated in a series of sequential EtOH washes (70%, 85%, and 100% EtOH) and left to dry. In parallel, the FISH probe was prepared by nick translation from a plasmid containing the major satellite repeats (**Table 3**). For each coverslip 300 μg of probe DNA and 9μg of salmon sperm DNA was resuspended in 50% formamide, 2X SSC and 10% dextran sulfate buffer). Finally, the probe and metaphase spreads or cells, were co-denatured for 5min at 85 °C, and left to hybridize at 37 °C Overnight.

The next day, the samples were washed twice for 20 min in 2XSSC at 42°C and subsequently incubated with fluorescein anti-biotin antibody (Vector Laboratories, SP-3040; Dilution 1:100) for at least 45min. Next, they were washed with 2X SSC, incubated with DAPI and mounted with Prolong Gold reagent (Molecular Probes).

Both metaphase spreads and interphase cells were observed and quantified with a confocal laser scanning microscope (LSM 880; Zeiss), using a x63 objective.

### Western Blot

#### Fractionation of soluble fraction in high salt buffer

This protocol was followed according to previous publication^55^. Briefly, after treatment with ICRF-193 cells were trypsinized, harvested and then lysed in TEB150 buffer, containing 50 mM HEPES pH 7.4, 500 mM NaCl, 2mM MgCl2, 5mM EGTA pH 8, 1mM dithiothreitol (DTT), 10% glycerol, 0.5% Triton X-100 and Protease Inhibitor Cocktail (PIC; Calbiochem), for 45min on ice. The chromatin fraction was then pelleted by centrifugation, washed twice in TEB150 buffer and resuspended in 1X Laemmli Sample buffer (Bio-Rad; #1610747), supplemented with 5% β-mercaptoethanol (Merck; M6250). Subsequently, chromatin was briefly incubated at 95 °C and sheared by vortexing.

#### Fractionation of nuclear soluble fraction in low salt buffer

Protocol was followed as in previous publication^56^. In brief, after treatment with ICRF-193, cells were trypsinized and harvested. Cytosolic protein fraction was collected with incubation in hypotonic buffer (10mM HEPES pH 7, 0.3 M sucrose, 0.5% Triton, 50 nM NaCl and PIC) for 10min on ice. After centrifugation (1500 g x 5’), supernatant was collected and the remaining pellet was incubated in the nuclear buffer (10 mM HEPES pH 7, 1 mM EDTA, 0.5% NP-40, 200 mM NaCl and PIC) for 10min on ice. The supernatant containing the nuclear fraction was collected by centrifugation (13000 g x 2’) and the remaining pellet was resuspended in lysis buffer (10 mM HEPES pH 7, 1 mM EDTA, 1% NP-40, 500 mM NaCl, Benzonase [Millipore] and PIC) and sonicated at 10-20% amplitude for three times. The chromatin protein fraction was then collected by centrifugation at 13000 g x 2min and collection of the supernatant.

#### Western Blot analysis

Equal amounts of protein samples were supplemented with β-mercaptoethanol (Sigma; M6250) and diluted in 1X Laemmli Sample buffer (Bio-Rad; #1610747), incubated at 95 °C for 10min. Protein samples where run on SDS-Page gels (4-12% NuPAGE Bis-Tris; Invitrogen), in Tris/Glycine/SDS buffer at 80-140V for 2h, then transferred on nitrocellulose membrane (0.45 μm; GE Healthcare Lifesciences, 10600062) for 90 min at 400 mA, in ice cold Tris/Glycine buffer, containing 10% methanol. Next, the membranes were blocked for 1h in 5% milk/PBS-T (Tween-20) for 1h. Primary antibodies (**Table 2**) were then incubated on the membranes overnight at 4°C, in 5% milk/PBS-T. After 3 washes in PBS-T x 5min each, the membranes were incubated for 1h in secondary antibody (**Table 2**) in 5% milk/PBS-T and washed again for 3 times x 5’ each. Protein levels were detected using chemiluminescent Clarity Western ECL reagent (Merck; GERPN2236) and developed using ImageQuant LAS 4000.

### Fluorescence recovery after photobleaching (FRAP)

The FRAP experiments were carried out in a Zeiss LSM880 confocal microscope, containing a FRAP unit and a 40x 1.3NA oil immersion objective lens. NIH3T3 MEFs were seeded in sterile cell culture chambers on coverglass (Sarstedt; 94.6190.402). Top2αWt plasmid was transfected for 16h, followed by treatment with DMSO or ICRF-193 for 4h, with 1h pretreatment with DMSO or Sumo inhibitor (ML-792). During all FRAP measurements, cells were kept at 37°C in a stage incubator. Protein fluorescence was excited with a laser line at 488 nm, 65.0 intensity and 500 ms interval. The recovery of fluorescence was monitored with the same frequency and exposure time for a total of 100 scans (3 before photobleaching). At least 20 cells were counted per condition, and 3 biological replicates were performed. FRAP kinetics raw data were acquired with a custom-made fiji plug-in macro, created by Dr. Alex Herbert. For the data normalization, three ROI’s were created: 1) The photobleached area, which was detected by detection of significant shifts in the intensity of signal using a standard score (The number of standard deviations from the mean). 2) The whole area of nucleus for the detection of loss of fluorescent signal due to photobleaching over time. 3) Background area outside the cell nucleus. FRAP curves were generated by background subtraction, correction of photobleaching overtime and normalization to pre-bleached values. The plot curves of the normalized intensity signal of the photobleached area were created over time after the raw values were analyzed online using the easyFRAP tool with double normalization^57^. Info: (https://easyfrap.vmnet.upatras.gr/easyfrap_web_manual_appendix.pdf)

### Quantitative PCR (qPCR/qRT-PCR) analysis

Isolation of mRNA transcripts was performed using the RNeasy Kit (QIAGEN, Cat No. 74104) according to manufacturer’s instructions. Subsequently, RevertAid First Strand cDNA synthesis Kit (ThermoFisher Scientific, Ref. K1622) was used according to manufacturer’s instructions to generate cDNA.

Analysis by qPCR was performed using the Luna Universal qPCR Master Mix (NEB; M3003X) and analyzed on an AriaMx Real-Time PCR instrument (G8830A). The corresponding Ct values were calculated with the Agilent AriaMx software (version 1.71).

### Chromatin Immunoprecipitation (ChIP) assay

After treatment with ICRF-193, cells were fixed in 1% PFA for 10min at RT, followed by the addition of 0.125 M glycine for 10min, on ice, to stop the cross-linking. Then they were washed twice with pre-cooled 1X PBS and collected by scraping. After centrifugation at 600g x 5’, they were initially lysed in 50 mM Tris-HCl pH 8, 2 mM EDTA pH 8, 10% glycerol, 0.1% NP-40 and PIC for 5’ at 4 °C. Subsequently they sample was centrifuged at 800g x 5’, and further lysed in 50mM Tris-HCl pH 8, 10 mM EDTA pH 8, 1% SDS and PIC. Next, the chromatin was sonicated with E220 Focused-ultrasonicator (Covaris) to create genomic DNA fragments with a size ranging between 300-800 bp. 1% of input chromatin was saved and 70 μg of chromatin pre-clearing was done by mixing with pre-blocked Sepharose beads (1:1 vol. mix of Protein A and protein G beads; Invitrogen) for 1h at 4°C. Subsequently, 5 μg of antibody were used to immunoprecipitate the chromatin fragments in rotation at 4°C overnight. The following day, protein A/G Sepharose beads, pre-blocked with BSA, were used to pull down the antibody for 4h at 4°C. The beads were washed first twice for 5min with 20 mM Tris pH8, 2 mM EDTA, 0.5% NP-40, 0.1% SDS, 150 mM NaCl and PIC, then they were washed for 5min with 20mM Tris pH8, 2 mM EDTA, 0.5% NP-40, 0.1% SDS, 500 mM NaCl and PIC, and finally for 5min with 10 mM Tris pH8, 1 mM EDTA, 0.5% NP-40, 0.5% sodium deoxycholate, 250 mM LiCl and PIC at 4 °C. The beads were then eluted twice for 10min at RT in TE (Tris-HCl 10 mM, EDTA 1 mM), 0.1 M NaHCO_3_ and 1% SDS. The eluted DNA samples (and their corresponding 1% Inputs) were then treated with 50 μg/ml of RNase A in the presence of 0.2 M NaCl for 30min at 37 °C, and followed by treatment with 0.07 mg/ml of Proteinase K. The chromatin fragments were then left to reverse crosslink overnight at 65°C with shaking and the DNA was then purified with the QIAquick PCR purification kit (QIAGEN; 28104), according to manufactures instructions, and resuspended in elution buffer.

After sample purification, qPCR analysis was performed using the primers from **tables 4 and 5**. The corresponding Ct values from samples and inputs were used to calculate the precipitated percentage over input, with the formula 100×2^C_t(adjustedinput)_-C_t(antibodyofinterest)_^. No antibody controls were used to subtract background antibody noise signal from the samples.

### Statistical analysis and figure preparation

Statistical analysis was performed using the GraphPad Prism software. Statistical significance was determined with the respective tests as: (*) p<0.05, (**) p<0.005, (***) p<0.001, (****) p<0.0001; Not significant differences are not marked on graphs. For figure preparation, the Adobe Illustrator and ImageJ (Fiji) were used.

### ChIP-seq analysis

The ChIP-seq libraries were sequenced on an Illumina Hiseq 4000 sequencer as paired-end 100 bases by the GenomEast platform, a member of the ‘France Genomique’ consortium (ANR-10-INBS-0009). ChIP samples were purified with Agencourt AMPure XP beads (Beckman Coulter) and their concentration was quantified with Qubit (Invitrogen). For the preparation of the ChIP-seq libraries, a total of 10 ng of dsDNA was purified using the MicroPlex Library Preparation kit v3 (C05010001, Diagenode s.a., Seraing, Belgium), according to manufacturer’s instructions. RTA and bcl2fastq were performed using base calling and image analysis. Mapping of sequence reads to the reference genome (mm10) was performed with Bowtie 2 ^58^, using the parameters: -q -N 1 -X 1000. Bedtools was used for the removal of reads overlapping with ENCODE hg38 blacklisted region V2 ^59^. Bigwig files were generated using bamCoverage from deepTools software^60^ with RPKM normalization. The peak calling was performed by MACS2 analysis and peak annotation by HOMER, using the default settings.

#### Repeats analysis

The analysis of repetitive sequences was done according to a previous publication^61^. Reads that were simultaneously mapped for more than one genomic site of the mouse genome, or not mapped at all were aligned to RepBase56 v21.08 (rodent) repetitive sequences. Reads that were mapped uniquely to a single repeat family were annotated with their corresponding family name. The read counts from the two annotation steps per repeat family were added, and the sums of read counts were used to calculate the proportion of each repeat type in every analyzed stage. Two biological replicates were performed, and independent libraries were created, having each library sequenced with at least two technical replicates. Repetitive elements analysis was performed until level 2, showing high reproducibility among the biological replicates.

The implemented process in DESeq2 Bioconductor package was used to identify significantly differential repeat sequences. Data analysis was performed in collaboration with the GenomEast platform from IGBMC. Reads were mapped to the mm9 mouse genome assembly using Bowtie. Since we were interested in repetitive sequences, multiple mapped reads were taken into consideration for the analysis. The peak calling was done by MACS2 analysis and peak annotation by HOMER, using the default settings. Data normalization and comparisons were calculated by DESeq2’s median of ratios for all sequence tracks, using the Bam Coverage.

##### END-seq

The END-seq was performed according to the original publication ^62,63^. Briefly, 15 million NIH3T3 cells were harvested, followed by a PBS wash, resuspended in cell suspension buffer and embedded in agarose before they were transferred into plug molds (Biorad; #1703591). After a 30min incubation at 4°C, the solidified plugs were treated with Proteinase K (QIAGEN; 158918) solution at 50°C for 1h and then transferred at 37 °C for 7h. After consecutive washes in wash buffer (10 mM Tris pH8 and 50 mM EDTA) and subsequently TE buffer (10 mM Tris pH8 and 1 mM EDTA), the plugs were embedded in RNase A solution at 37 °C for 1h while shaking. The DSB ends were then processed with blunting enzymes (ExoVII and ExoT enzymes; NEB, M0379 and M0265 respectively), and for A-tailing with Klenow Fragment (3’->5’ exo–) (NEB; M0212). A biotinylated primer (ENDseq-adaptor-1, 5′-Phos-GATCGGAAGAGCGTCGTGTAGGGAAAGAGTGUU[Biotin-dT]U[Biotin-dT]UUACACTCTTTCCCTACACGACGCTCTTCCGATC∗T-3′[∗phosphorothioate bond]) was then ligated at the free DSB ends with the quick ligase enzyme (NEB; M2200). After the primer ligation, the agarose plugs were dissolved with β-agarase (NEB; M0392) enzyme according to manufacturer’s instructions. The recovered DNA was sonicated (Covaris) to be fragmented between 150-200 bp, and the biotinylated fragments were isolated with streptavidin beads (Invitrogen; 65-001). The newly generated DSBs by the sonication step were also processed with T4 DNA polymerase (15 U) (NEB; M0203), Klenow fragment (5 U) & T4 polynucleotide kinase (15 U) (NEB; M0201). The A-tailing was performed with Klenow exo-fragment (15 U) and then the ends were ligated to a second primer: (5′-Phos-GATCGGAAGAGCACACGTCUUUUUUUUAGACGTGTGCTCTTCCGATC∗T-3′ [∗phosphorothioate bond]). The libraries were then prepared by digesting the hairpin structures on both primer adaptors using Uracil-Specific Excision Reagent (USER) enzyme (NEB; M5505) and a following PCR amplification for 16 cycles with Truseq barcoded primers. Finally, they were quantified with qPCR.

Libraries were sequenced in collaboration with Dr. Andre Nussenzweig’s lab with an Illumina Hiseq 2500, as a 1×50 bases single end reads. Data analysis was performed in collaboration with the Galaxeast genome platform from IGBMC, similar to ChIp-seq analysis.

#### Top2cc isolation assay

The Top2cc isolation assay was performed using a cc-seq protocol adapted from^16^. In brief, NIH3T3 or U2OS cells were treated with DMSO, ICRF-193 (5 μM for 4h) or Etoposide (50 μM for 30’). Since they have a slower cell cycle, the U2OS cells were first synchronized in G2 phase with RO-3306 for 20h prior to addition of DMSO and ICRF-193, or 23.5h prior to addition of etoposide. Subsequently, the cells were fixed in ice cold 70% EtOH for 10 min, collected by scraping and pelleted by centrifugation. After removing the supernatant, the cells were lysed in 2% SDS, 0.5M Tris pH 8.0 and 10 mM EDTA buffer at 65 °C for 10 min. After cooling the samples down on ice, Phenol-Chloroform-isoamyl alcohol (25:24:1; Sigma) was supplemented and the samples were mixed with pipetting. The three phases were separated by centrifugation at 2×10^5^ g for 20 min and 500 μl of the aqueous phase were transferred in a new tube, and the nucleic acids were precipitated with 1 ml EtOH, washed with 70% ice-cold EtOH and resuspended in TE buffer. Samples were then sonicated to an average of 300-400 bp size fragment with a Covaris sonicator (10% duty cycle, 75W intensity power incidence, 200 cycles for 20 min). 5% of input was stored and the rest of the samples were then supplemented with a final concentration of 0.3M NaCl, 0.1% N-Lauroylsarcosine salt and 0.2% Triton-X100 and passed through Miniprep (QIAGEN) silica-fibre membrane spin columns in order to bind only the Top2cc’s and not DNA. Six steps of washes were performed with 10mM Tris, 0.3M NaCl and 1mM EDTA buffer and the Top2cc-bound DNA was eluted in 500 μl of 1mM EDTA, 0.5% SDS and 10mM Tris buffer. The samples were incubated at 50°C for 1h with 1 mg/ml proteinase K, prior to precipitation overnight at -20°C with a final concentration of 70% ethanol, 75 mM sodium acetate and 50 μg/ml glycogen. Samples were centrifuged at 2.1×10^4^ g for 1 hour at 60°C, prior to washing once with 1 ml 70% ethanol, and resuspension in 100 μL TE. Finally, qPCR was performed to assess the enrichment of signal in repetitive sequences.

## Supplemental items

- **<u>FRAP Videos: 1) DMSO, 2) ICRF-193, 3) Sumoi+ICRF-193</u>**
- **<u>Table S1 for γH2AX ChIP-seq repeat analysis</u>**
- **<u>Table S2 for END-seq repeat analysis</u>**

