## Supplementary material for "Inhibition of Topoisomerase 2 catalytic activity impacts the integrity of heterochromatin and repetitive DNA and leads to interlinks between clustered repeats": Table S1

### Table S1: Repetitive sequences from ChIP-seq analysis

#### Normalized values of γH2AX ChIP-seq reads by DESeq2 analysis

| 2 Biological replicates | Average Input values | | | Average γH2AX ChIP values | | |
|---|---|---|---|---|---|---|
| **Repeat family name** | **DMSO** | **ICRF-193 0h release** | **ICRF-193 3h release** | **DMSO** | **ICRF-193 0h release** | **ICRF-193 3h release** |
| (A)n | 20942.8937 | 21043.4664 | 20599.4211 | 19680.8514 | 21017.2281 | 20422.6509 |
| (AAATG)n | 172.01108 | 177.88007 | 180.057365 | 146.782527 | 146.124639 | 166.206668 |
| (AACTG)n | 171.34841 | 174.859044 | 168.705795 | 151.150894 | 131.054688 | 162.269801 |
| (AAGTG)n | 25.5914321 | 25.30651 | 28.2404287 | 27.5130127 | 33.127386 | 38.5002162 |
| (AATAG)n | 225.51591 | 251.676903 | 248.079715 | 207.049737 | 218.92443 | 224.605535 |
| (AATTG)n | 68.8778291 | 63.8380584 | 56.5136994 | 77.8217388 | 60.1545982 | 72.1221618 |
| (ACATG)n | 112.806974 | 112.98127 | 113.093082 | 116.375608 | 114.734082 | 134.770517 |
| (ACCG)n | 33.4878993 | 34.1231348 | 42.4263268 | 46.8326161 | 48.8427185 | 54.3170921 |
| (ACCTG)n | 66.4903301 | 61.2254928 | 64.0571419 | 90.0691838 | 76.8799418 | 94.3611471 |
| (ACGTG)n | 9.12953656 | 7.67366524 | 10.6804759 | 11.591762 | 10.8269278 | 15.161747 |
| (ACTAG)n | 75.9218427 | 96.4910811 | 75.5472074 | 77.9535359 | 91.5451558 | 83.0572272 |
| (ACTCG)n | 7.89646722 | 4.08176697 | 11.4172535 | 9.89258565 | 8.64525512 | 17.2685011 |
| (ACTG)n | 595.332393 | 598.375686 | 611.915503 | 653.306044 | 690.666989 | 631.823875 |
| (ACTTG)n | 9.95577698 | 14.8573606 | 6.1426993 | 6.19934066 | 6.58512559 | 8.12402874 |
| (AGATG)n | 65.6840116 | 59.5924622 | 81.4129152 | 59.5261896 | 66.4164225 | 58.790528 |
| (AGCTG)n | 235.760033 | 226.698355 | 239.135529 | 290.53177 | 341.170991 | 309.041933 |
| (AGGGGG)n | 1097.36883 | 1144.50603 | 1155.02974 | 1428.93579 | 1433.10036 | 1485.9682 |
| (AGGTG)n | 316.476271 | 322.779964 | 317.469966 | 378.556147 | 434.89782 | 421.711785 |
| (AGTAG)n | 97.4271598 | 93.3890516 | 74.3363871 | 97.9018023 | 102.25054 | 102.83073 |
| (AGTTG)n | 53.7276426 | 47.1843439 | 52.1072902 | 43.007574 | 30.0973528 | 43.6024343 |
| (ATAAG)n | 68.0778019 | 58.8576592 | 82.4781115 | 56.1278368 | 53.8927738 | 53.7673502 |
| (ATAGG)n | 92.8692068 | 105.14418 | 97.2365779 | 113.272147 | 109.784906 | 123.246189 |
| (ATATG)n | 380.783041 | 450.292555 | 370.379718 | 349.847917 | 360.079229 | 386.750188 |
| (ATCTG)n | 34.642329 | 34.7758209 | 40.8870879 | 39.6945591 | 28.7239331 | 27.32803 |
| (ATG)n | 5794.49653 | 5854.77243 | 5799.62162 | 5435.44716 | 5471.41698 | 5370.13664 |
| (ATGGTG)n | 3294.33776 | 3238.82118 | 3229.70544 | 3393.78933 | 3391.94547 | 3379.05997 |
| (ATGTG)n | 367.33147 | 406.291245 | 340.778516 | 352.544616 | 359.794814 | 388.32696 |
| (ATTAG)n | 69.6254299 | 72.164612 | 84.9183374 | 94.5574893 | 75.1631672 | 97.4093046 |
| (ATTG)n | 2310.0265 | 2307.70524 | 2320.78923 | 2151.91234 | 2335.48315 | 2182.47168 |
| (ATTTG)n | 260.144609 | 243.922132 | 220.69101 | 229.523539 | 249.627056 | 228.466901 |
| (C)n | 686.693292 | 712.827877 | 711.1161 | 843.473073 | 940.150007 | 945.240444 |
| (CA)n | 287137.849 | 286907.892 | 284875.833 | 282208.39 | 287569.994 | 287556.265 |
| (CAA)n | 15247.2734 | 15597.1931 | 15202.2176 | 14142.8947 | 13777.5718 | 13801.7594 |
| (CAAA)n | 36057.2389 | 36617.4073 | 36507.5339 | 33718.3403 | 34985.8537 | 33770.9801 |
| (CAAAA)n | 28507.8514 | 28790.5954 | 28753.2599 | 25920.461 | 26971.606 | 25940.6171 |
| (CAAAAA)n | 11869.2615 | 11965.6326 | 11821.9754 | 10720.7077 | 10898.4172 | 10542.2722 |
| (CAAAC)n | 4386.96042 | 4404.87951 | 4360.61055 | 4022.07941 | 4106.71052 | 4215.92006 |

| | | | | | | |
|---|---|---|---|---|---|---|
| (CAAAG)n | 729.848623 | 708.908017 | 727.120393 | 678.476056 | 678.506749 | 627.959461 |
| (CAAAT)n | 335.240211 | 308.657912 | 308.704247 | 323.526332 | 323.215656 | 328.693829 |
| (CAACC)n | 221.075393 | 278.861985 | 228.36582 | 235.765548 | 265.806677 | 244.227544 |
| (CAACG)n | 13.7336247 | 11.7554322 | 14.1858981 | 16.8135077 | 6.58512559 | 13.1735533 |
| (CAACT)n | 59.1841843 | 66.1229656 | 62.8862917 | 51.8836861 | 67.668299 | 45.590628 |
| (CAAG)n | 1711.18363 | 1711.86101 | 1667.93241 | 1704.93115 | 1878.72728 | 1836.99981 |
| (CAAGA)n | 316.461593 | 315.18963 | 275.322795 | 362.098664 | 338.689734 | 337.828668 |
| (CAAGC)n | 130.751488 | 127.02252 | 118.400479 | 105.970995 | 123.116807 | 126.712355 |
| (CAAGG)n | 258.596981 | 241.881553 | 236.523966 | 252.478069 | 261.889397 | 308.011856 |
| (CAAGT)n | 2.67584444 | 9.14306897 | 9.81233083 | 9.08566637 | 2.86838257 | 9.14447234 |
| (CAAT)n | 2259.09214 | 2216.60059 | 2265.24003 | 2227.82535 | 2360.08622 | 2251.52239 |
| (CAATA)n | 261.594199 | 293.880949 | 264.435976 | 257.909369 | 256.09186 | 237.282039 |
| (CAATC)n | 44.5718928 | 45.2248691 | 53.942106 | 53.0708354 | 50.6609826 | 57.3323161 |
| (CAATG)n | 34.7209687 | 29.5514991 | 33.7120342 | 37.956504 | 38.3390919 | 33.5033852 |
| (CAATT)n | 64.3261674 | 65.1434305 | 70.6010719 | 66.6177874 | 64.3964004 | 70.9172909 |
| (CACAA)n | 443.613483 | 388.332259 | 436.222413 | 440.925994 | 443.159613 | 461.61752 |
| (CACAC)n | 910.706662 | 913.074131 | 882.144374 | 1073.71012 | 1051.79389 | 1038.83449 |
| (CACAG)n | 1377.93143 | 1388.67127 | 1409.61342 | 1776.32835 | 2181.94081 | 2182.50461 |
| (CACAT)n | 304.723317 | 270.044753 | 282.160138 | 254.491577 | 276.169317 | 300.035781 |
| (CACCAT)n | 3405.49382 | 3397.19054 | 3469.31717 | 3521.78751 | 3770.94933 | 3693.54663 |
| (CACCC)n | 1790.31135 | 1770.31446 | 1736.48303 | 2370.27937 | 2547.79874 | 2482.39903 |
| (CACCG)n | 29.6051988 | 25.7961764 | 22.2290969 | 24.3317948 | 21.1287965 | 40.0111216 |
| (CACCT)n | 275.996785 | 263.513946 | 283.92927 | 294.818589 | 334.667912 | 330.882671 |
| (CACG)n | 2098.65576 | 2105.66476 | 2011.20754 | 2402.78562 | 2433.97379 | 2481.99724 |
| (CACGC)n | 34.0520076 | 31.674196 | 23.8011777 | 41.3083977 | 46.984347 | 45.6433215 |
| (CACGT)n | 7.79161434 | 17.959289 | 17.757004 | 12.2280056 | 11.1502291 | 15.2012671 |
| (CACTA)n | 193.942624 | 160.981724 | 187.962068 | 173.527499 | 191.775674 | 173.481507 |
| (CACTC)n | 201.294905 | 210.941451 | 233.232016 | 279.366697 | 267.88747 | 285.815438 |
| (CACTG)n | 438.137019 | 440.00683 | 433.028988 | 619.377044 | 620.04704 | 689.304145 |
| (CACTT)n | 11.7267413 | 12.897987 | 7.97751507 | 17.1937378 | 10.8269278 | 8.13720211 |
| (CAG)n | 5523.15521 | 5497.38554 | 5469.34232 | 6762.92214 | 6847.10652 | 7031.50867 |
| (CAGA)n | 14975.3282 | 14796.778 | 14942.8771 | 15238.3694 | 15353.6258 | 15286.311 |
| (CAGAA)n | 365.868773 | 324.90256 | 340.328023 | 345.976904 | 353.129473 | 337.337714 |
| (CAGAC)n | 217.783014 | 245.309622 | 225.49865 | 288.17312 | 598.755983 | 455.201997 |
| (CAGAG)n | 746.93282 | 734.215033 | 787.008147 | 988.882125 | 976.389472 | 1033.22864 |
| (CAGAGA)n | 12282.0084 | 12059.754 | 12253.8122 | 12581.181 | 12444.6296 | 12842.7881 |
| (CAGAT)n | 27.6245287 | 26.6124893 | 25.5374679 | 25.4724849 | 39.0458554 | 27.4465904 |
| (CAGC)n | 585.376616 | 596.172794 | 619.186918 | 778.24013 | 815.558901 | 782.799389 |
| (CAGCC)n | 721.308884 | 680.337267 | 709.290576 | 900.826937 | 933.162586 | 945.566239 |
| (CAGCG)n | 30.7596285 | 32.6535287 | 32.613996 | 37.8711662 | 52.5594615 | 38.5792565 |
| (CAGCT)n | 306.389428 | 312.657158 | 290.611696 | 423.12352 | 383.833321 | 365.120225 |
| (CAGG)n | 2605.95476 | 2649.34363 | 2655.10592 | 3389.63149 | 3555.535 | 3636.70733 |

| | | | | | | |
|---|---|---|---|---|---|---|
| (CAGGA)n | 389.767881 | 353.311099 | 412.280575 | 450.686782 | 481.499315 | 519.97028 |
| (CAGGC)n | 263.391377 | 240.002879 | 256.024388 | 318.16093 | 418.273966 | 398.42601 |
| (CAGGG)n | 476.963501 | 464.415113 | 482.808835 | 660.060569 | 752.295886 | 688.66573 |
| (CAGGT)n | 49.8187288 | 38.6947704 | 43.2616301 | 27.1948909 | 58.0944687 | 34.9945305 |
| (CAGT)n | 507.97897 | 554.537395 | 549.331916 | 557.495507 | 595.93076 | 565.870482 |
| (CAGTA)n | 373.010825 | 364.08639 | 394.385075 | 425.683656 | 703.492664 | 601.733962 |
| (CAGTC)n | 49.4118999 | 53.388403 | 71.4035332 | 55.064904 | 47.186105 | 51.7001162 |
| (CAGTG)n | 544.739327 | 583.845477 | 599.843576 | 778.077035 | 827.538047 | 798.761173 |
| (CAGTT)n | 141.716998 | 167.34941 | 180.352941 | 177.794879 | 126.934429 | 170.890878 |
| (CAT)n | 5751.36793 | 5590.61217 | 5599.98606 | 5306.76151 | 5179.99662 | 5240.34319 |
| (CATA)n | 19840.8102 | 19750.2247 | 19601.4841 | 18447.1043 | 19513.7503 | 18861.5069 |
| (CATAA)n | 172.08972 | 169.144955 | 171.671491 | 165.202292 | 183.655478 | 185.058528 |
| (CATAC)n | 186.486014 | 174.206965 | 187.021111 | 174.543972 | 169.150694 | 184.521959 |
| (CATAG)n | 121.884084 | 151.512414 | 136.223166 | 125.825855 | 152.830087 | 152.045605 |
| (CATAT)n | 422.173699 | 388.494976 | 413.043066 | 355.589758 | 337.920978 | 328.548922 |
| (CATATA)n | 10088.5778 | 10091.0691 | 10240.5635 | 9304.465 | 9634.55507 | 9483.55275 |
| (CATCC)n | 310.927459 | 366.291001 | 299.851458 | 401.658612 | 397.791772 | 417.422777 |
| (CATCG)n | 13.0646636 | 7.34701866 | 10.5491084 | 9.89258565 | 15.3920313 | 2.04088721 |
| (CATCT)n | 90.7113355 | 85.5522654 | 69.4466426 | 55.0454647 | 70.6582248 | 67.9481737 |
| (CATG)n | 3048.41853 | 3137.75816 | 3050.39478 | 3399.11206 | 3404.60644 | 3491.56161 |
| (CATGC)n | 113.895347 | 148.574011 | 138.02514 | 165.838536 | 154.890217 | 179.824576 |
| (CATGT)n | 126.790148 | 142.042293 | 122.905413 | 151.445786 | 121.884374 | 121.715524 |
| (CATTA)n | 63.3358324 | 63.6742292 | 66.1946627 | 52.2250375 | 70.3349236 | 61.8255121 |
| (CATTC)n | 74.4591457 | 73.4698831 | 74.3363871 | 93.4945565 | 94.6152964 | 72.5006498 |
| (CATTG)n | 48.2448876 | 35.5923361 | 41.5581818 | 50.1845097 | 28.8855837 | 37.5192927 |
| (CATTT)n | 229.319447 | 171.75833 | 217.07777 | 171.851552 | 169.111807 | 171.315473 |
| (CCA)n | 8229.53172 | 8360.77781 | 8125.11004 | 9358.48701 | 9655.71176 | 9611.13916 |
| (CCAA)n | 4866.61969 | 5039.50194 | 4911.50742 | 4867.80306 | 5146.0397 | 5082.47133 |
| (CCCA)n | 1830.17011 | 1859.86203 | 1755.36162 | 2259.98646 | 2420.74521 | 2481.18403 |
| (CCCCAA)n | 211.696831 | 230.207731 | 249.823133 | 294.44215 | 297.318167 | 299.002164 |
| (CCCCAG)n | 1458.28593 | 1505.32727 | 1490.32953 | 2016.72304 | 2149.82404 | 2085.30608 |
| (CCCCCA)n | 2775.85935 | 2910.24734 | 2834.89342 | 3661.91417 | 3925.21239 | 3991.18584 |
| (CCCCCG)n | 263.535549 | 250.452737 | 243.337759 | 298.344949 | 320.083523 | 312.830847 |
| (CCCCCT)n | 1143.01022 | 1156.34439 | 1137.19993 | 1391.64634 | 1471.44189 | 1474.38813 |
| (CCCCG)n | 1411.53624 | 1413.08472 | 1339.6949 | 1287.58786 | 1196.16409 | 1207.79341 |
| (CCCG)n | 225.771227 | 238.208347 | 224.811135 | 217.217776 | 220.074817 | 187.132349 |
| (CCCGA)n | 7.94889365 | 7.02037208 | 4.43925101 | 3.14233925 | 8.28184649 | 4.05542768 |
| (CCCGAA)n | 21.0921969 | 35.9189827 | 22.9658745 | 47.1275083 | 43.2274966 | 54.6692334 |
| (CCCGG)n | 78.466621 | 86.0419317 | 62.7220824 | 52.0543618 | 68.7597459 | 53.7541768 |
| (CCCTA)n | 30.8644813 | 30.0413678 | 23.8997033 | 38.5927476 | 46.4592878 | 49.7119225 |
| (CCCTAA)n | 1848.29078 | 1963.37358 | 1858.32757 | 2101.09917 | 2202.4911 | 2198.35796 |
| (CCCTAG)n | 182.051788 | 198.206889 | 214.130659 | 260.473783 | 270.230794 | 251.317956 |

| | | | | | | |
|---|---|---|---|---|---|---|
| (CCCTG)n | 637.936723 | 678.867357 | 689.060505 | 976.355925 | 1959.56094 | 1547.45169 |
| (CCG)n | 2293.50222 | 2375.7174 | 2341.93823 | 2292.65153 | 2233.45381 | 2193.72694 |
| (CCGAA)n | 16.3570427 | 13.0614115 | 12.7358921 | 25.984512 | 8.96855633 | 15.2144405 |
| (CCGAG)n | 44.6767457 | 42.6129106 | 41.492498 | 29.7630948 | 28.3605245 | 22.3180256 |
| (CCGCG)n | 161.36642 | 193.147003 | 169.494 | 152.508719 | 121.359315 | 137.022178 |
| (CCGG)n | 12.4743421 | 12.897987 | 7.9446732 | 17.7057649 | 17.0887522 | 10.1517426 |
| (CCGGA)n | 12.4743421 | 12.7349672 | 11.4500954 | 6.7502464 | 13.5336598 | 13.1867266 |
| (CCGGG)n | 80.6832101 | 87.8381844 | 73.3040327 | 58.6571621 | 64.558051 | 48.6783055 |
| (CCGTA)n | 5.19440956 | 2.93880746 | 1.8019739 | 3.69324499 | 8.12019588 | 11.1458394 |
| (CCTA)n | 788.894926 | 685.317261 | 705.92365 | 800.058249 | 827.559932 | 833.436495 |
| (CCTAA)n | 53.1373211 | 43.9190922 | 47.7994067 | 49.1680361 | 45.6109273 | 55.7160238 |
| (CCTAG)n | 73.8363197 | 84.2466908 | 68.1280041 | 79.1406852 | 77.0816998 | 119.714157 |
| (CCTAT)n | 81.4376262 | 68.899158 | 97.0909539 | 96.5050987 | 95.1804631 | 106.761011 |
| (CCTCG)n | 71.5012471 | 89.144366 | 72.26455 | 98.669843 | 90.0903004 | 84.0644974 |
| (CCTG)n | 2828.98408 | 2886.3272 | 2884.73182 | 3811.02833 | 3839.19918 | 3972.60708 |
| (CCTTG)n | 143.881161 | 155.431161 | 181.826497 | 193.785818 | 180.988854 | 184.614173 |
| (CG)n | 661.081414 | 662.377472 | 665.472799 | 637.280504 | 746.721382 | 753.7137 |
| (CGA)n | 227.870381 | 257.799958 | 234.564911 | 222.288285 | 261.948947 | 243.855643 |
| (CGAA)n | 77.5549256 | 85.2258212 | 70.8966487 | 73.3448042 | 82.7583038 | 75.4961138 |
| (CGAAA)n | 33.9733679 | 29.7147212 | 32.7453635 | 51.4181182 | 39.8340548 | 36.5383692 |
| (CGAAG)n | 5.86337067 | 9.63273531 | 7.9446732 | 5.94332712 | 11.6752883 | 11.1194927 |
| (CGAAT)n | 1.33792222 | 1.30618162 | 0.90098695 | 2.16474429 | 0 | 1.02044361 |
| (CGAG)n | 244.777901 | 236.902469 | 263.534989 | 274.57164 | 265.584254 | 296.112088 |
| (CGAGA)n | 19.3542611 | 21.796324 | 17.7241622 | 17.4265217 | 15.3920313 | 25.3266629 |
| (CGAGG)n | 70.8847124 | 80.0008923 | 62.8041871 | 76.7393667 | 106.088827 | 80.1276301 |
| (CGAT)n | 37.3443867 | 38.5309413 | 24.6364809 | 30.5700141 | 34.2990476 | 50.6269791 |
| (CGATA)n | 9.82471089 | 14.3675931 | 9.64812149 | 9.34167991 | 5.93852315 | 10.1253958 |
| (CGATG)n | 16.4094691 | 14.693835 | 14.9883595 | 25.4336062 | 25.8555505 | 14.2203436 |
| (CGCGG)n | 157.674027 | 190.207285 | 153.039213 | 133.445129 | 113.320555 | 106.991545 |
| (CGCTG)n | 30.8644813 | 34.9392453 | 29.1742576 | 35.9159763 | 45.7725779 | 34.4711353 |
| (CGG)n | 2237.93179 | 2323.71254 | 2248.04783 | 2193.29092 | 2097.49128 | 1982.58123 |
| (CGGA)n | 181.120695 | 182.859761 | 180.451467 | 205.206905 | 250.818772 | 258.395194 |
| (CGGAA)n | 10.5460984 | 9.9593819 | 10.1314568 | 13.7565063 | 12.84695 | 9.14447234 |
| (CGGAG)n | 40.1775104 | 49.3068384 | 41.492498 | 43.985169 | 47.3477556 | 23.2726023 |
| (CGGG)n | 198.087457 | 216.820382 | 237.457794 | 226.637213 | 256.616919 | 225.194797 |
| (CGGGA)n | 21.8203997 | 27.7554488 | 29.075732 | 24.3706734 | 32.4005687 | 20.2507917 |
| (CGGGG)n | 1446.15131 | 1493.25106 | 1440.60111 | 1258.73647 | 1293.1616 | 1217.44505 |
| (CGGGGG)n | 218.418946 | 219.595461 | 228.701367 | 292.129959 | 240.153495 | 269.307453 |
| (CGGT)n | 54.3441772 | 54.8580091 | 48.4705005 | 59.5261896 | 67.5880842 | 67.9086536 |
| (CGGTG)n | 41.8698351 | 31.3475494 | 30.4764752 | 50.2233883 | 66.1745571 | 42.5424705 |
| (CGTAG)n | 0.66896111 | 1.95927243 | 0 | 0.80691928 | 0 | 0 |
| (CGTG)n | 2407.28851 | 2456.52803 | 2348.52061 | 2804.2304 | 2896.84684 | 2907.44879 |

| | | | | | | |
|---|---|---|---|---|---|---|
| (CGTTG)n | 16.4356823 | 20.572057 | 7.87898946 | 17.4497513 | 15.7153325 | 19.2962149 |
| (CTA)n | 1415.70257 | 1370.38423 | 1483.4736 | 1409.262 | 1376.44496 | 1383.28591 |
| (CTAA)n | 755.111866 | 701.316064 | 765.640066 | 649.686766 | 739.186406 | 658.223675 |
| (CTAAA)n | 190.316289 | 218.287863 | 176.86394 | 170.047598 | 155.315618 | 178.478338 |
| (CTAAG)n | 24.2797231 | 16.0002189 | 18.5266235 | 21.7792399 | 24.5222383 | 24.2798726 |
| (CTAAT)n | 139.743144 | 122.287156 | 125.509849 | 115.805263 | 120.915691 | 102.027648 |
| (CTACT)n | 71.8750475 | 74.776267 | 72.600097 | 64.7090566 | 67.2246756 | 73.8864081 |
| (CTAG)n | 71.5012471 | 71.8378643 | 100.422874 | 81.2665508 | 85.3635463 | 101.412039 |
| (CTAGG)n | 93.4196844 | 83.592993 | 87.8511914 | 90.1815415 | 134.933082 | 129.15149 |
| (CTAGGG)n | 251.179167 | 271.024187 | 265.534014 | 324.833907 | 350.100051 | 373.988056 |
| (CTAGT)n | 43.2339706 | 61.8783812 | 49.502855 | 45.0481018 | 56.316312 | 47.6578619 |
| (CTATA)n | 414.146166 | 374.209904 | 379.725134 | 354.860596 | 364.847311 | 406.773986 |
| (CTATG)n | 189.58127 | 165.06511 | 168.36312 | 180.572637 | 209.99476 | 199.644186 |
| (CTATT)n | 296.157889 | 325.718772 | 286.590096 | 275.999154 | 256.192739 | 268.155275 |
| (CTCA)n | 1351.46081 | 1453.57068 | 1463.11929 | 1603.36692 | 1631.64408 | 1573.64072 |
| (CTCCG)n | 76.7480831 | 93.8794262 | 84.1158761 | 98.669843 | 92.5952743 | 96.2703006 |
| (CTCG)n | 301.77171 | 334.944463 | 300.499003 | 351.885142 | 349.009214 | 369.731981 |
| (CTCGG)n | 95.3741413 | 112.98127 | 89.5546396 | 77.9146572 | 57.4478662 | 63.8795727 |
| (CTCTG)n | 726.5164 | 772.174696 | 757.894609 | 926.016389 | 956.452391 | 1002.2566 |
| (CTG)n | 5151.31821 | 5180.97376 | 5161.69334 | 6377.54508 | 6331.34705 | 6565.75363 |
| (CTGGGG)n | 1432.65465 | 1412.59444 | 1454.76129 | 1976.28318 | 2326.51032 | 2199.32217 |
| (CTGTG)n | 2065.63393 | 2117.33342 | 2147.2693 | 2689.94227 | 3381.31114 | 3036.72444 |
| (CTTA)n | 372.538987 | 337.63864 | 395.220378 | 356.059116 | 357.069249 | 341.709302 |
| (CTTAA)n | 13.7336247 | 13.5510778 | 15.0212013 | 8.62009849 | 8.28184649 | 7.10358513 |
| (CTTAG)n | 35.954038 | 18.9392288 | 26.6683479 | 30.9113655 | 33.8140958 | 19.2698682 |
| (CTTAT)n | 74.2819444 | 71.0210455 | 65.5564107 | 54.2191061 | 55.4278441 | 73.0503917 |
| (CTTCG)n | 5.27304922 | 5.5511707 | 3.50542219 | 5.47775923 | 6.10017376 | 8.11085536 |
| (CTTG)n | 1824.81108 | 1771.37308 | 1709.77471 | 1768.75553 | 1871.15219 | 1905.69887 |
| (CTTTA)n | 15.7667212 | 16.979754 | 7.97751507 | 13.5663913 | 12.645192 | 15.1749204 |
| (CTTTG)n | 720.26717 | 689.969091 | 710.172978 | 745.838655 | 731.085654 | 703.165761 |
| (G)n | 745.031314 | 712.664149 | 659.778429 | 883.078504 | 892.720815 | 954.47713 |
| (GA)n | 63861.5618 | 62453.8569 | 64153.175 | 69224.1979 | 66108.603 | 69381.631 |
| (GAA)n | 11435.7454 | 10803.3183 | 11363.6609 | 10020.616 | 9734.17543 | 9792.5876 |
| (GAAA)n | 22730.7143 | 22030.9712 | 22793.8742 | 19069.8967 | 19068.0207 | 19064.2559 |
| (GAAAA)n | 5750.12385 | 5619.83136 | 5725.4637 | 5214.74705 | 5304.3374 | 5235.9432 |
| (GAATG)n | 91.0982424 | 87.3483157 | 84.2144017 | 90.9420016 | 108.148956 | 84.1962312 |
| (GACTG)n | 77.2660562 | 76.0823475 | 96.2042235 | 115.692905 | 119.945788 | 122.617407 |
| (GAGAA)n | 5287.91526 | 4458.46715 | 4385.99362 | 6780.39958 | 5256.69696 | 5814.23765 |
| (GAGTG)n | 132.712761 | 148.083333 | 129.752048 | 176.19669 | 186.382264 | 151.933631 |
| (GATTG)n | 39.5415778 | 43.1024757 | 36.1194182 | 55.5380524 | 31.2690145 | 35.4652321 |
| (GCATG)n | 76.5908038 | 96.1643334 | 97.203736 | 117.477419 | 97.685437 | 102.353442 |
| (GCCTAA)n | 89.0127195 | 97.7971616 | 84.148718 | 106.397684 | 105.037487 | 145.916356 |

| | | | | | | |
|---|---|---|---|---|---|---|
| (GCCTG)n | 226.401393 | 240.24832 | 249.921659 | 320.170648 | 318.00273 | 328.292041 |
| (GCGTG)n | 37.3705999 | 45.7149402 | 33.4492992 | 46.5301433 | 36.844129 | 52.7073865 |
| (GCTG)n | 540.555174 | 528.578907 | 566.751209 | 708.204551 | 774.514908 | 707.843384 |
| (GCTTG)n | 111.134571 | 119.349259 | 110.324438 | 144.485985 | 170.727093 | 148.938167 |
| (GGA)n | 13364.0766 | 13253.3235 | 13326.7901 | 15571.9174 | 15842.0566 | 16096.6948 |
| (GGAA)n | 13841.2472 | 13694.6285 | 13930.7551 | 14428.4973 | 13979.9891 | 14108.9576 |
| (GGAAA)n | 2254.61335 | 2287.54043 | 2305.12761 | 2145.36737 | 2204.6916 | 2172.81394 |
| (GGAGA)n | 2148.55313 | 2146.23091 | 2008.61736 | 2577.33668 | 2433.75259 | 2758.54965 |
| (GGAGAA)n | 1718.68061 | 1625.32739 | 1635.89594 | 1793.57282 | 1737.89639 | 1703.3683 |
| (GGATG)n | 431.978488 | 401.067227 | 446.238923 | 522.728289 | 494.79172 | 524.575449 |
| (GGCTG)n | 652.66645 | 641.153639 | 566.187933 | 818.889055 | 805.439957 | 861.406481 |
| (GGGA)n | 7097.5861 | 7009.4039 | 7035.61099 | 8257.97311 | 8603.42376 | 8626.32841 |
| (GGGAA)n | 1225.16976 | 1185.89285 | 1230.09362 | 1430.59988 | 1474.85539 | 1463.47941 |
| (GGGAGA)n | 3464.56529 | 3477.92371 | 3507.34918 | 4020.68515 | 3897.61451 | 4085.38841 |
| (GGGGA)n | 1517.66723 | 1472.67495 | 1449.47744 | 1760.12027 | 1795.66328 | 1837.49381 |
| (GGGTG)n | 1724.78567 | 1782.39553 | 1725.08436 | 2219.82682 | 2395.5156 | 2312.34417 |
| (GGTTG)n | 273.734061 | 264.248142 | 276.092415 | 301.192396 | 300.288039 | 286.496914 |
| (GTATG)n | 222.584225 | 229.961988 | 217.945915 | 218.362256 | 224.357947 | 261.496045 |
| (GTCTG)n | 174.817991 | 175.349318 | 148.173018 | 191.101467 | 176.382422 | 183.554209 |
| (GTGTG)n | 1036.58511 | 1001.40229 | 1049.7921 | 1166.26975 | 1228.13804 | 1188.14151 |
| (GTTTG)n | 4053.71294 | 4059.65246 | 4076.83619 | 3794.94961 | 3875.28249 | 3814.97234 |
| (T)n | 21555.0943 | 21337.8398 | 20855.1342 | 20190.2835 | 21053.7422 | 20327.3068 |
| (TA)n | 33237.8027 | 32813.6935 | 33005.3032 | 28643.2655 | 30104.6466 | 28739.4505 |
| (TAA)n | 6974.32477 | 6716.65539 | 6923.41736 | 6208.30099 | 6395.17166 | 6288.81261 |
| (TAAA)n | 12125.7866 | 11444.3069 | 11922.1576 | 10577.4288 | 10813.5605 | 10684.0404 |
| (TAAAA)n | 3938.87601 | 3766.00673 | 3874.18847 | 3500.05045 | 3514.88057 | 3613.95507 |
| (TAAAAA)n | 1116.40224 | 1102.87184 | 1072.1975 | 1011.44078 | 989.48134 | 1015.13729 |
| (TAAAG)n | 43.0242648 | 49.633485 | 32.5483123 | 37.2349226 | 36.9656722 | 36.4856757 |
| (TAAG)n | 406.276436 | 398.209322 | 414.877882 | 394.201945 | 350.302419 | 368.606151 |
| (TAATG)n | 94.9285166 | 105.634251 | 96.565484 | 83.516633 | 80.4751416 | 85.1771547 |
| (TACGG)n | 9.85092411 | 6.36748362 | 4.50493474 | 10.6995049 | 9.45350816 | 11.1326661 |
| (TACTG)n | 313.236319 | 326.207932 | 317.918295 | 412.516934 | 825.276159 | 619.196524 |
| (TAG)n | 1453.27711 | 1429.16129 | 1409.22644 | 1346.51999 | 1336.04452 | 1334.20936 |
| (TAGA)n | 26612.5215 | 26776.4181 | 26577.4243 | 24476.3824 | 24957.6042 | 23977.5798 |
| (TAGG)n | 736.347402 | 723.601953 | 739.457218 | 719.470122 | 780.251063 | 773.865691 |
| (TAGGG)n | 57.7676224 | 64.8170875 | 54.9416185 | 69.4187752 | 79.3034799 | 82.6458057 |
| (TAGTG)n | 327.455412 | 329.474499 | 335.970876 | 338.76867 | 336.223646 | 354.227727 |
| (TATAA)n | 1525.55898 | 1496.34358 | 1456.73027 | 1239.75394 | 1322.10918 | 1280.47799 |
| (TATAG)n | 376.421163 | 397.311702 | 354.750942 | 328.371638 | 366.826005 | 328.256061 |
| (TATATG)n | 10424.6956 | 10446.5094 | 10517.1857 | 9500.76022 | 9679.43979 | 9517.82933 |
| (TATCG)n | 5.86337067 | 5.87761493 | 4.47209288 | 7.72784136 | 8.12019588 | 12.1926298 |
| (TATG)n | 19067.3271 | 18921.9678 | 18980.3282 | 17948.8903 | 18502.6995 | 18180.8577 |

| | | | | | | |
|---|---|---|---|---|---|---|
| (TATTG)n | 260.767959 | 253.472751 | 246.434822 | 239.167691 | 250.85949 | 268.63915 |
| (TC)n | 64627.4987 | 64114.6258 | 65006.1015 | 70133.6724 | 67498.0036 | 70470.4606 |
| (TCC)n | 12977.728 | 12925.3162 | 13268.1302 | 15222.3914 | 15369.9524 | 15677.8908 |
| (TCCA)n | 12310.0686 | 12104.1603 | 12266.0477 | 13739.6053 | 14433.1017 | 14380.4077 |
| (TCCC)n | 6880.29694 | 6770.62893 | 6837.58811 | 8143.72434 | 8237.50688 | 8413.79036 |
| (TCCCC)n | 1390.94996 | 1460.91871 | 1428.41703 | 1767.93296 | 1810.45126 | 1833.94861 |
| (TCCCG)n | 39.6071109 | 29.8781457 | 33.3836155 | 28.4441484 | 37.8340863 | 43.5365674 |
| (TCCG)n | 257.580433 | 226.779258 | 220.911611 | 276.356155 | 344.040595 | 308.23885 |
| (TCCGG)n | 10.3888191 | 5.71439281 | 13.1863856 | 18.8929142 | 16.8869942 | 15.2144405 |
| (TCCTG)n | 371.882608 | 394.781455 | 419.69265 | 461.832416 | 476.145402 | 473.293341 |
| (TCG)n | 248.805298 | 220.248349 | 210.625237 | 241.604098 | 224.821624 | 233.108051 |
| (TCGGG)n | 8.59164155 | 10.122604 | 9.77948897 | 14.3927499 | 28.2389813 | 13.1208598 |
| (TCGTG)n | 2.64963122 | 3.10202958 | 2.70296085 | 8.44942279 | 4.40345286 | 7.1167585 |
| (TCTA)n | 26800.6223 | 26792.2574 | 26941.9634 | 24496.5037 | 24516.2938 | 23895.1985 |
| (TCTCC)n | 2259.13722 | 2179.37421 | 2238.43102 | 2711.31279 | 2609.59051 | 2783.34633 |
| (TCTCCC)n | 3469.65432 | 3544.37524 | 3392.58053 | 4123.07905 | 3916.03719 | 4103.67736 |
| (TCTCG)n | 16.4356823 | 11.7554322 | 10.5491084 | 12.9495871 | 21.6137483 | 19.2435214 |
| (TCTCTG)n | 12276.1776 | 12261.6369 | 12174.0163 | 12716.3992 | 12612.8925 | 12888.211 |
| (TCTG)n | 15133.7078 | 15143.6371 | 15017.891 | 16192.7507 | 16352.7575 | 16367.3205 |
| (TCTTG)n | 294.955752 | 274.455595 | 251.95569 | 280.11481 | 227.772053 | 253.898951 |
| (TG)n | 288689.227 | 289608.601 | 286511.047 | 283617.22 | 286762.417 | 289187.201 |
| (TGAA)n | 6681.63961 | 6716.50056 | 6498.98645 | 6507.59321 | 6792.15274 | 6684.2467 |
| (TGAG)n | 1482.22435 | 1456.91684 | 1438.43851 | 1568.41289 | 1633.17976 | 1620.19251 |
| (TGG)n | 8509.87256 | 8474.41024 | 8512.23082 | 9694.62537 | 10014.4084 | 10097.4322 |
| (TGGA)n | 12572.9662 | 12534.2906 | 12635.7762 | 14016.4981 | 14436.8409 | 14562.2165 |
| (TGGG)n | 1876.60995 | 1837.00709 | 1835.76293 | 2328.85103 | 2406.86697 | 2635.23454 |
| (TGGGGG)n | 2660.33826 | 2601.18238 | 2565.73191 | 3293.62656 | 3476.60225 | 3481.88411 |
| (TTA)n | 6998.95837 | 6794.53237 | 6699.35943 | 6071.87331 | 6397.69608 | 6138.8079 |
| (TTAA)n | 3500.29756 | 3492.37585 | 3424.13429 | 3165.81985 | 3211.26292 | 3204.37713 |
| (TTAAA)n | 249.368883 | 230.860417 | 255.548181 | 212.481037 | 219.651247 | 203.103765 |
| (TTAAG)n | 23.7418281 | 27.9186709 | 25.6359935 | 28.8708377 | 24.8455395 | 20.2507917 |
| (TTACG)n | 11.1364199 | 9.30629109 | 7.14221185 | 6.83558424 | 6.90842681 | 5.06269791 |
| (TTAG)n | 664.741302 | 683.928659 | 737.014828 | 649.481002 | 739.772848 | 672.476952 |
| (TTAGG)n | 47.2477372 | 40.3273963 | 45.8660653 | 48.423225 | 32.6023267 | 44.557011 |
| (TTAGGC)n | 74.0722387 | 77.3893385 | 85.7136705 | 93.7429895 | 92.7569249 | 87.1258283 |
| (TTAGGG)n | 3094.18417 | 3113.92045 | 3000.80485 | 6064.12517 | 5569.43845 | 6115.25207 |
| (TTATA)n | 1505.28569 | 1447.85254 | 1537.80548 | 1248.60682 | 1284.03139 | 1196.87456 |
| (TTATG)n | 140.228088 | 148.08394 | 145.871288 | 111.960781 | 121.86371 | 124.724161 |
| (TTC)n | 11306.6637 | 10845.5228 | 11117.2347 | 9812.4796 | 9754.22856 | 9797.12995 |
| (TTCA)n | 6544.11725 | 6415.67772 | 6308.55844 | 6415.36735 | 6469.06473 | 6287.58189 |
| (TTCC)n | 14050.555 | 13790.7158 | 14026.611 | 14539.2838 | 14466.0492 | 14597.5616 |
| (TTCCC)n | 1249.26494 | 1240.83207 | 1278.92754 | 1428.37254 | 1397.75119 | 1481.58393 |

| | | | | | | |
|---|---|---|---|---|---|---|
| (TTCCG)n | 5.91579711 | 3.59189827 | 7.07652812 | 5.39242139 | 5.73676514 | 7.1167585 |
| (TTCG)n | 87.0844758 | 92.8990817 | 95.3453711 | 71.8125132 | 110.12826 | 80.5127049 |
| (TTCGG)n | 5.27304922 | 5.22452412 | 11.4829373 | 9.8072478 | 9.45350816 | 7.10358513 |
| (TTCGGG)n | 41.1222344 | 54.0414939 | 35.5140081 | 49.5482661 | 50.3777888 | 29.3820906 |
| (TTCTC)n | 5039.19699 | 4308.42657 | 4361.15296 | 6676.15099 | 5240.03117 | 5601.996 |
| (TTCTCC)n | 1618.87224 | 1551.61288 | 1618.36667 | 1618.03133 | 1600.13137 | 1616.87784 |
| (TTCTG)n | 313.446025 | 294.208 | 305.370162 | 302.21645 | 265.766569 | 268.810404 |
| (TTG)n | 15536.3034 | 15677.929 | 15590.2815 | 14272.7774 | 14424.1994 | 14040.41 |
| (TTGG)n | 5142.80049 | 5170.77005 | 5140.35874 | 5181.50172 | 5906.51778 | 5633.5849 |
| (TTGGGG)n | 279.610538 | 257.962371 | 235.228876 | 320.341324 | 333.91921 | 334.717691 |
| (TTGTG)n | 431.086714 | 461.719722 | 469.146242 | 446.035382 | 505.819795 | 483.027076 |
| (TTTA)n | 12315.7595 | 12177.7812 | 12085.9451 | 10967.3709 | 11496.5485 | 11058.1765 |
| (TTTAA)n | 244.935705 | 247.02396 | 225.228787 | 199.768024 | 229.973169 | 195.628279 |
| (TTTAG)n | 154.971446 | 177.71705 | 154.913999 | 123.106415 | 129.984516 | 133.671033 |
| (TTTC)n | 22844.5693 | 22511.7086 | 22751.1465 | 19126.225 | 19456.9235 | 19011.9042 |
| (TTTCC)n | 2375.58573 | 2292.60173 | 2395.42832 | 2252.05237 | 2255.3532 | 2362.21771 |
| (TTTCG)n | 28.3721294 | 19.4286927 | 31.6801672 | 18.8929142 | 18.9070163 | 15.2012671 |
| (TTTG)n | 36631.4896 | 36605.3164 | 36361.3642 | 33659.5266 | 34780.9648 | 34158.3287 |
| (TTTTA)n | 4083.02507 | 3936.61937 | 3976.35756 | 3479.93909 | 3888.7335 | 3636.10234 |
| (TTTTC)n | 5888.59356 | 5728.97408 | 5739.2998 | 5074.76881 | 5386.97111 | 5130.4423 |
| (TTTTG)n | 29099.4531 | 29276.9819 | 29174.4596 | 26517.4112 | 27175.1566 | 26405.3952 |
| (TTTTTA)n | 1145.59537 | 1132.25901 | 1120.95861 | 1035.21029 | 1015.63831 | 1021.55635 |
| (TTTTTG)n | 12103.7644 | 12170.2922 | 12059.1376 | 10994.6019 | 11099.2404 | 10835.0965 |
| 4.5SRNA | 2871.11291 | 2818.16477 | 2760.69553 | 2930.75412 | 3179.5059 | 3132.7353 |
| 5S | 3197.97957 | 3266.66016 | 3173.23884 | 3341.23383 | 3497.16344 | 3493.16473 |
| 7SK | 3799.7536 | 3722.33733 | 3710.34174 | 3996.30213 | 4118.68295 | 4104.82954 |
| 7SLRNA | 1710.93461 | 1724.92303 | 1742.61643 | 1858.71615 | 1982.28924 | 1967.07649 |
| A-rich | 52368.8614 | 51935.8739 | 52040.3722 | 47036.1399 | 48011.1187 | 47253.2575 |
| AmnSINE1 | 5442.03319 | 5427.83187 | 5560.88453 | 5610.77861 | 5628.86745 | 5615.45049 |
| AmnSINE2 | 1559.39185 | 1541.24554 | 1607.08078 | 1673.02606 | 1618.93812 | 1671.88299 |
| Arthur1 | 6402.18687 | 6335.92591 | 6317.05927 | 5947.76607 | 6163.33584 | 5896.32533 |
| Arthur1A | 669.843966 | 675.600992 | 679.107514 | 691.999779 | 748.638083 | 648.019239 |
| Arthur1B | 3657.49975 | 3697.92884 | 3659.10756 | 3600.02408 | 3570.99513 | 3438.60442 |
| Arthur1C | 832.075946 | 792.992194 | 788.460317 | 795.841118 | 828.427736 | 745.826792 |
| AT_rich | 328727.029 | 326208.432 | 327586.503 | 289587.131 | 297161.115 | 285932.494 |
| B1_Mm | 133894.724 | 132636.691 | 130318.174 | 142867.875 | 151691.166 | 149277.371 |
| B1_Mur1 | 152674.901 | 153782.358 | 150326.528 | 160715.749 | 177245.482 | 171281.904 |
| B1_Mur2 | 125623.153 | 125857.046 | 123047.28 | 131358.743 | 142533.155 | 138200.099 |
| B1_Mur3 | 96256.5082 | 97036.8885 | 94666.24 | 103479.012 | 111840.788 | 108736.109 |
| B1_Mur4 | 153240.207 | 154598.289 | 150959.585 | 164198.399 | 184958.676 | 176622.439 |
| B1_Mus1 | 331810.627 | 331549.282 | 327140.301 | 359679.078 | 391468.319 | 383163.397 |
| B1_Mus2 | 237376.1 | 237138.152 | 232035.318 | 257523.033 | 279977.621 | 271865.879 |

| | | | | | | |
|---|---|---|---|---|---|---|
| B1F | 172680.055 | 172749.692 | 169761.511 | 181389.978 | 194485.284 | 187942.683 |
| B1F1 | 75855.0981 | 76462.3288 | 74806.0417 | 80704.0085 | 85861.2124 | 82811.202 |
| B1F2 | 63123.3098 | 63102.6925 | 61946.4621 | 66647.8451 | 72005.9737 | 70084.324 |
| B2_Mm1a | 57353.4254 | 56654.8755 | 55523.5747 | 58724.4468 | 64411.0706 | 62154.9861 |
| B2_Mm1t | 85137.6035 | 84545.8241 | 82739.1168 | 87015.7458 | 95637.8216 | 92208.3115 |
| B2_Mm2 | 371417.931 | 369900.545 | 362751.927 | 380164.29 | 415380.181 | 400469.964 |
| B3 | 735917.978 | 736647.774 | 725063.657 | 764413.788 | 820226.593 | 798661.478 |
| B3A | 422290.077 | 422178.901 | 418350.482 | 439980.68 | 462059.089 | 455390.135 |
| B4 | 318911.635 | 321027.284 | 316639.529 | 342992.955 | 352595.02 | 351959.353 |
| B4A | 571634.181 | 575313.028 | 571840.063 | 616642.968 | 636079.87 | 631647.485 |
| BC1_Mm | 18476.0346 | 18362.6305 | 18279.0061 | 18898.0027 | 20054.1151 | 19468.1561 |
| BGLII | 17294.6352 | 17174.0286 | 17265.0385 | 16767.9042 | 15663.7629 | 16324.7016 |
| BGLII_A | 1781.14983 | 1787.78196 | 1764.3058 | 1854.37859 | 1730.13838 | 1923.37879 |
| BGLII_B | 8475.96263 | 8453.26003 | 8643.84837 | 8909.4027 | 8776.01533 | 9091.03764 |
| BGLII_B2 | 2499.46255 | 2637.42427 | 2469.63334 | 2555.23553 | 2577.98119 | 2730.63894 |
| BGLII_C | 438.970075 | 407.923871 | 471.072455 | 437.419074 | 450.713421 | 463.928461 |
| BGLII_Mur | 5115.29392 | 5132.56465 | 5298.02279 | 5377.81503 | 5527.10247 | 5509.14347 |
| BGLII_Mus | 4324.94783 | 4240.14084 | 4397.24181 | 4383.13275 | 4100.8686 | 4243.49131 |
| BLACKJACK | 5040.35612 | 5076.80699 | 5095.50644 | 4577.12617 | 4510.8656 | 4476.85014 |
| C-rich | 42467.4271 | 42614.066 | 42166.7734 | 49581.6944 | 49998.8746 | 51545.172 |
| C573_MM | 753.688489 | 761.562831 | 788.016952 | 706.178696 | 583.386447 | 684.133013 |
| Chap1_Mam | 832.240565 | 833.808042 | 922.158988 | 826.752483 | 870.120162 | 861.633475 |
| Charlie1 | 6816.14902 | 6837.89222 | 6903.66999 | 6422.46983 | 6882.604 | 6560.81962 |
| Charlie10 | 2040.75078 | 2090.31702 | 2067.07586 | 1928.56443 | 1959.10337 | 1933.55029 |
| Charlie10a | 325.559673 | 305.474171 | 307.51914 | 311.806563 | 321.982002 | 273.152106 |
| Charlie10b | 240.397149 | 238.125117 | 239.405392 | 233.597014 | 237.426709 | 250.959228 |
| Charlie11 | 313.596489 | 288.656627 | 322.883016 | 296.358461 | 277.179328 | 273.952142 |
| Charlie12 | 4783.5301 | 4816.31235 | 4681.95929 | 4284.75633 | 4583.76322 | 4317.17707 |
| Charlie13a | 1212.78716 | 1247.69144 | 1253.91121 | 1116.46168 | 1142.69306 | 1148.69684 |
| Charlie13b | 791.754263 | 798.540734 | 748.736951 | 750.695819 | 760.699276 | 727.021039 |
| Charlie14a | 351.584147 | 335.923796 | 394.385075 | 302.984491 | 330.567707 | 311.593042 |
| Charlie15a | 1041.67362 | 1006.95477 | 1017.24162 | 1037.58459 | 1034.46389 | 1050.55996 |
| Charlie15b | 1667.406 | 1669.41163 | 1683.77033 | 1503.56775 | 1545.63821 | 1493.85255 |
| Charlie16a | 1060.51725 | 1088.42355 | 1126.88071 | 1054.54506 | 1089.99322 | 1128.1233 |
| Charlie17 | 859.143706 | 794.37857 | 792.019966 | 846.975715 | 819.09455 | 771.015134 |
| Charlie17a | 284.509787 | 334.372377 | 322.56389 | 313.164388 | 330.385392 | 358.13166 |
| Charlie17b | 826.691229 | 897.970521 | 903.862258 | 856.286096 | 834.388754 | 862.433511 |
| Charlie18a | 1226.93967 | 1234.79285 | 1189.08662 | 1244.80122 | 1225.39303 | 1177.19986 |
| Charlie19a | 1531.57125 | 1483.85648 | 1465.91365 | 1453.2325 | 1567.06903 | 1565.18736 |
| Charlie1a | 18654.7201 | 18560.2604 | 18439.8265 | 17207.624 | 18213.6371 | 17358.8179 |
| Charlie1b | 5196.26443 | 5209.29836 | 5185.13389 | 4920.4263 | 5266.97667 | 4949.53446 |
| Charlie1b_Mars | 17.1308567 | 24.3267726 | 36.0865763 | 22.7179562 | 30.9056058 | 20.2903118 |

| | | | | | | |
|---|---|---|---|---|---|---|
| Charlie20a | 1078.56609 | 1048.83116 | 981.277123 | 996.850819 | 1029.59676 | 994.13611 |
| Charlie21a | 1094.20909 | 1064.91309 | 1135.78276 | 990.569443 | 986.833548 | 929.381001 |
| Charlie22a | 1042.99215 | 1041.32183 | 943.70057 | 1021.75247 | 1057.51244 | 920.052102 |
| Charlie23a | 790.062462 | 839.846957 | 841.384325 | 818.702242 | 838.385639 | 807.056455 |
| Charlie24 | 2383.61222 | 2406.97241 | 2494.07493 | 2252.55254 | 2282.1361 | 2206.37307 |
| Charlie25 | 2115.94547 | 2161.90894 | 2203.18408 | 1899.37975 | 1905.02767 | 1807.44037 |
| Charlie26a | 550.379885 | 489.068734 | 504.465363 | 532.550698 | 501.173256 | 465.001598 |
| Charlie29a | 1985.38324 | 2119.05018 | 1922.58673 | 1878.15899 | 1947.79393 | 1875.08861 |
| Charlie2a | 3144.48313 | 3161.67415 | 3073.6491 | 2877.4571 | 2941.75711 | 2800.17097 |
| Charlie2b | 5168.71382 | 5215.09245 | 5069.14576 | 4539.49206 | 4910.13351 | 4807.18003 |
| Charlie4 | 749.721381 | 769.969579 | 754.624044 | 702.319053 | 728.299317 | 698.402511 |
| Charlie4a | 6503.44326 | 6487.5262 | 6357.29232 | 6184.12141 | 6437.77078 | 6258.52255 |
| Charlie4z | 1854.24484 | 1923.37465 | 1923.36781 | 1972.60033 | 2065.7592 | 2073.02478 |
| Charlie5 | 7351.95069 | 7306.46899 | 7196.15151 | 6769.77705 | 7011.68732 | 6761.09967 |
| Charlie6 | 1011.9641 | 968.666539 | 1036.27513 | 1020.91046 | 905.124752 | 883.168178 |
| Charlie7 | 5360.33343 | 5465.79133 | 5478.18581 | 4943.13765 | 5032.96712 | 5055.15038 |
| Charlie7a | 449.5686 | 451.354308 | 443.855088 | 426.106555 | 495.335612 | 398.808037 |
| Charlie8 | 4199.61566 | 4283.49129 | 4105.78056 | 4359.78234 | 4355.26557 | 4392.84443 |
| Charlie9 | 1022.52382 | 1058.70923 | 1043.18962 | 985.21211 | 985.540954 | 976.587921 |
| Cheshire | 6116.18608 | 6159.51514 | 6105.54017 | 5725.12788 | 5890.88632 | 5755.07697 |
| Cheshire_Mars | 16.9211509 | 12.5715428 | 12.4824498 | 8.91499068 | 15.7153325 | 10.164916 |
| Cheshire_Mars_ | 40.0464443 | 37.8780528 | 38.1512852 | 32.7812175 | 41.8941843 | 28.3879938 |
| CR1_Mam | 1520.81544 | 1511.69586 | 1455.52658 | 1495.99922 | 1540.06065 | 1482.0217 |
| CT-rich | 68415.6262 | 67636.6642 | 68772.3557 | 72674.8237 | 72088.575 | 73526.4018 |
| CYRA11_Mm | 3001.1451 | 3195.71911 | 3076.47414 | 3066.27816 | 3051.19744 | 3084.52465 |
| DNA1_Mam | 208.069971 | 200.655828 | 198.945249 | 195.981861 | 199.491133 | 203.712787 |
| ERV3-16A3_I-int | 7292.81579 | 7247.37095 | 7154.24849 | 7000.34078 | 6966.38749 | 6875.08184 |
| ERV3-16A3_LTR | 1752.90405 | 1744.67837 | 1776.52056 | 1823.94417 | 1774.45794 | 1817.6281 |
| ERVB2_1-I_MM-int | 925.738365 | 965.400984 | 956.159471 | 843.371598 | 770.718561 | 801.424255 |
| ERVB2_1A-I_MM-int | 2988.58583 | 2922.32537 | 2931.59307 | 2786.29418 | 2588.38394 | 2569.33738 |
| ERVB3_1-I_MM-int | 2011.28504 | 1963.04754 | 2025.32343 | 1712.62293 | 1672.30888 | 1721.21289 |
| ERVB3_1-LTR_MM | 1158.79686 | 1145.40335 | 1177.29881 | 1115.66993 | 1173.11371 | 1219.34713 |
| ERVB4_1-I_MM-int | 6757.13418 | 6681.24378 | 6768.84044 | 6866.27652 | 6373.93771 | 6743.0565 |
| ERVB4_1-LTR_MM | 93.8003001 | 96.6546068 | 84.2872137 | 81.7321187 | 78.5158911 | 91.7507579 |
| ERVB4_1B-I_MM-int | 12935.4246 | 12795.3633 | 12891.193 | 11631.3969 | 11488.4045 | 11525.7327 |
| ERVB4_1B-LTR_MM | 6112.50522 | 6041.96496 | 6270.58779 | 6499.02959 | 6180.23496 | 6699.08825 |
| ERVB4_1C-LTR_Mm | 4302.89256 | 4198.02175 | 4169.97836 | 4509.75415 | 4399.90354 | 4568.61004 |
| ERVB4_2-I_MM-int | 9105.64481 | 9140.9516 | 9219.64725 | 9757.79076 | 9634.00925 | 9878.64894 |
| ERVB4_2-LTR_MM | 820.22443 | 790.706376 | 790.853445 | 862.862365 | 897.084771 | 929.87805 |
| ERVB4_3-I_MM-int | 2921.58854 | 2856.94166 | 2925.99442 | 2931.95264 | 2755.69387 | 2965.27107 |
| ERVB4_3-LTR_MM | 1083.6352 | 1176.18063 | 1087.44859 | 990.864823 | 829.476023 | 929.506148 |

| | | | | | | |
|---|---|---|---|---|---|---|
| ERVB5_1-I_MM-int | 4474.79197 | 4424.88343 | 4552.01885 | 4388.65696 | 4383.31857 | 4459.63384 |
| ERVB5_1-LTR_MM | 1263.35978 | 1218.71274 | 1260.88705 | 1287.08342 | 1206.94969 | 1207.58568 |
| ERVB5_2-LTR_MM | 2363.42962 | 2287.38297 | 2267.48537 | 2275.82947 | 2241.8171 | 2292.91976 |
| ERVB7_1-LTR_MM | 4246.36325 | 4212.63245 | 4299.08936 | 4687.76665 | 4457.62972 | 4694.9966 |
| ERVB7_2-LTR_MM | 2736.54687 | 2735.55192 | 2791.38547 | 2999.38969 | 2944.053 | 3175.34718 |
| ERVB7_2B-LTR_MM | 3133.41644 | 3084.3737 | 3177.85159 | 3418.3577 | 3401.09146 | 3531.54993 |
| ERVB7_3-LTR_MM | 2246.59735 | 2247.70624 | 2238.39818 | 2228.48152 | 2129.26163 | 2318.32192 |
| ERVB7_4-LTR_MM | 1295.0374 | 1226.38681 | 1233.14142 | 1367.01158 | 1296.57448 | 1272.71269 |
| ERVL-B4-int | 11656.1728 | 11724.9091 | 11651.6712 | 11170.1628 | 10851.9792 | 10919.8019 |
| ERVL-E-int | 7851.81405 | 7734.80392 | 7878.57605 | 7580.04698 | 7643.02008 | 7567.73104 |
| ERVL-int | 5855.84592 | 5969.63814 | 5910.79652 | 5520.44914 | 5428.20526 | 5498.21194 |
| ETnERV-int | 20029.4331 | 20110.6624 | 20240.7623 | 19819.8782 | 18744.8918 | 19829.4109 |
| ETnERV2-int | 19673.0467 | 19410.468 | 19642.7312 | 17335.6212 | 16609.3046 | 16501.104 |
| ETnERV3-int | 11896.0341 | 11761.3075 | 11789.5114 | 10273.7193 | 15572.0458 | 12527.2495 |
| Eulor1 | 303.464034 | 294.045587 | 291.03864 | 274.152532 | 224.862953 | 235.59988 |
| Eulor10 | 165.091841 | 155.593776 | 134.749611 | 139.093564 | 120.955799 | 153.019941 |
| Eulor11 | 289.756623 | 264.003916 | 244.304429 | 270.641821 | 259.525409 | 251.920391 |
| Eulor12 | 162.547063 | 166.043026 | 160.72828 | 140.110038 | 158.808718 | 175.35114 |
| Eulor2A | 250.588845 | 226.615428 | 275.08361 | 209.909043 | 233.99316 | 243.842469 |
| Eulor2B | 215.487261 | 211.186183 | 197.347454 | 203.41481 | 194.684164 | 228.568749 |
| Eulor2C | 100.227779 | 116.409946 | 95.8029928 | 106.358806 | 96.3521247 | 101.412039 |
| Eulor3 | 146.727915 | 150.450964 | 153.61891 | 138.83755 | 135.377926 | 145.96905 |
| Eulor4 | 153.11555 | 144.410329 | 142.562917 | 118.501473 | 115.299248 | 133.072137 |
| Eulor5A | 420.612964 | 401.637796 | 414.713673 | 391.459275 | 375.975655 | 367.931262 |
| Eulor5B | 163.268451 | 172.084066 | 176.110742 | 159.685655 | 149.234887 | 155.192562 |
| Eulor6A | 104.372612 | 95.8382939 | 100.202274 | 99.6474379 | 85.1617883 | 92.8173083 |
| Eulor6B | 335.089747 | 291.595637 | 289.579341 | 294.527487 | 298.874512 | 273.715022 |
| Eulor6C | 65.4019575 | 67.5923693 | 54.7117255 | 49.9749553 | 51.7311548 | 61.8386854 |
| Eulor6D | 253.4356 | 261.391148 | 244.435797 | 239.462583 | 234.355347 | 226.09668 |
| Eulor6E | 88.3699716 | 97.6337371 | 96.4341166 | 102.619101 | 95.7869581 | 85.190328 |
| Eulor7 | 22.836948 | 19.9185614 | 17.7898459 | 12.1426678 | 7.91843787 | 15.267134 |
| Eulor8 | 632.919514 | 624.661427 | 644.93073 | 559.869806 | 541.753571 | 550.372814 |
| Eulor9A | 314.495601 | 309.554926 | 296.083301 | 295.714637 | 281.461238 | 247.460129 |
| Eulor9B | 64.1164617 | 55.184251 | 53.7122129 | 48.9120225 | 60.3563562 | 52.7073865 |
| Eulor9C | 289.704196 | 310.53608 | 267.237462 | 256.353848 | 315.619909 | 262.111654 |
| FAM | 317.728739 | 300.412262 | 275.813259 | 319.076417 | 353.677028 | 380.933561 |
| FordPrefect | 2728.05956 | 2660.60757 | 2769.73607 | 2857.61789 | 3495.04193 | 3126.01375 |
| FordPrefect_a | 373.673494 | 337.882259 | 340.067452 | 353.813312 | 374.864765 | 396.556376 |
| G-rich | 43352.4931 | 43583.8774 | 43522.3812 | 51506.487 | 51815.894 | 52825.0455 |
| GA-rich | 67908.0221 | 66612.8146 | 67643.2046 | 71349.4827 | 71500.2333 | 72154.1263 |
| GC_rich | 17109.6543 | 17941.9403 | 17068.2642 | 17056.6889 | 16810.8704 | 16129.5606 |
| GSAT_MM | 2759019.42 | 2747142.67 | 2770792.3 | 2042989.43 | 10889452.3 | 6079181.55 |

| | | | | | | |
|---|---|---|---|---|---|---|
| HAL1 | 22242.5677 | 22490.6875 | 22145.1598 | 21202.9281 | 22353.1498 | 21578.3753 |
| HAL1_SS | 230.900104 | 245.309318 | 244.581421 | 204.807235 | 217.267816 | 206.33635 |
| HAL1b | 3918.47689 | 3895.2372 | 3882.89564 | 3418.12822 | 3399.58072 | 3462.08121 |
| HAL1M8 | 4368.8581 | 4282.92163 | 4430.01506 | 4065.02916 | 4127.17083 | 4094.55009 |
| HAL1ME | 5215.62655 | 5263.74721 | 5074.67159 | 4636.20293 | 4825.93979 | 4689.7074 |
| hAT-N1_Mam | 1000.65781 | 989.075475 | 972.909834 | 910.567799 | 903.691782 | 905.091002 |
| Helitron1Na_Mam | 414.336473 | 398.04782 | 372.090295 | 366.715952 | 368.037774 | 373.96171 |
| Helitron1Nb_Mam | 547.520548 | 558.374632 | 545.136815 | 510.814127 | 539.553676 | 484.093626 |
| Helitron2Na_Mam | 395.978839 | 387.107689 | 389.183333 | 395.835223 | 364.341085 | 347.433716 |
| Helitron3Na_Mam | 1089.03355 | 1106.95503 | 1070.69327 | 1056.62495 | 1049.83525 | 1082.8914 |
| HERV16-int | 1905.72339 | 1872.68012 | 1956.57079 | 1951.63229 | 2007.45992 | 1841.23003 |
| HERVL32-int | 2.00688333 | 7.34701866 | 7.07652812 | 2.25008213 | 4.56510347 | 0 |
| HERVL40-int | 1298.3497 | 1356.9992 | 1345.45559 | 1286.5676 | 1233.12793 | 1322.12212 |
| HERVL74-int | 1089.44615 | 1126.05677 | 1137.95313 | 1076.80979 | 1040.68377 | 998.902407 |
| HUERS-P3b-int | 12.3694892 | 21.5511873 | 16.8231752 | 18.5126841 | 13.6953104 | 17.2158076 |
| HY1 | 103.834717 | 122.614612 | 114.993582 | 131.831291 | 144.084564 | 133.829113 |
| HY3 | 34.878248 | 36.4088514 | 32.5811541 | 35.8770976 | 22.1388075 | 28.4011671 |
| HY4 | 9.82471089 | 20.8980965 | 13.2520693 | 8.44942279 | 7.27183543 | 6.06996815 |
| HY5 | 1.33792222 | 6.69392785 | 6.20838304 | 2.33541998 | 5.41346392 | 2.01454047 |
| IAP-d-int | 11201.4014 | 11139.0177 | 11581.1538 | 12041.112 | 11351.7637 | 11762.0961 |
| IAP1-MM_I-int | 7603.79515 | 7716.11145 | 7730.30947 | 7716.93032 | 7290.75719 | 7861.7232 |
| IAP1-MM_LTR | 2506.48716 | 2452.28294 | 2526.78312 | 2656.85381 | 2495.52125 | 2709.64435 |
| IAPA_MM-int | 403.075279 | 406.863028 | 379.152566 | 408.125337 | 453.784782 | 437.456208 |
| IAPEy-int | 11756.9925 | 11719.2776 | 12307.9857 | 12027.8077 | 10774.7267 | 12004.181 |
| IAPEY_LTR | 4334.5167 | 4377.69635 | 4484.2171 | 4841.99068 | 4400.5076 | 4914.97062 |
| IAPEY2_LTR | 7889.75925 | 7846.40357 | 8131.05505 | 8571.53961 | 8088.08724 | 8623.08315 |
| IAPEY3-int | 22886.2179 | 22825.0828 | 23277.4117 | 23467.6206 | 22786.0561 | 23947.6911 |
| IAPEY3_LTR | 3804.26332 | 3821.03656 | 3905.54888 | 4199.30211 | 4060.10597 | 4230.95617 |
| IAPEY3C_LTR | 2162.37116 | 2176.11422 | 2212.55801 | 2312.09535 | 2317.66575 | 2453.06304 |
| IAPEY4_I-int | 9774.55506 | 9802.02026 | 10047.1149 | 10119.5723 | 12694.3484 | 11603.057 |
| IAPEY4_LTR | 707.844207 | 756.091956 | 706.04356 | 670.593188 | 741.74971 | 698.139043 |
| IAPEY5_I-int | 1764.22292 | 1790.14726 | 1886.23462 | 1851.32159 | 1666.02456 | 1804.30305 |
| IAPEY5_LTR | 1331.97391 | 1282.38768 | 1360.82163 | 1415.63678 | 1312.04551 | 1439.46301 |
| IAPEz-int | 206413.378 | 206486.752 | 208177.452 | 218621.469 | 262259.465 | 244484.484 |
| IAPLTR1_Mm | 7963.70147 | 7971.05259 | 8106.29496 | 8556.78888 | 10246.9363 | 9503.66679 |
| IAPLTR1a_Mm | 9223.66979 | 9260.2153 | 9454.36084 | 10166.6329 | 13198.4034 | 12021.0695 |
| IAPLTR2_Mm | 8367.68897 | 8448.04674 | 8406.17863 | 9547.60578 | 9248.55637 | 9681.55026 |
| IAPLTR2a | 3954.19292 | 4005.60884 | 3972.7223 | 4411.33653 | 4089.65821 | 4392.18272 |
| IAPLTR2a2_Mm | 9999.88171 | 9780.54795 | 10251.5389 | 10314.5026 | 10191.9181 | 10709.0039 |
| IAPLTR2b | 6153.9179 | 6091.03294 | 6118.36746 | 6608.5492 | 6173.52829 | 6602.10758 |
| IAPLTR3 | 1197.99085 | 1170.87389 | 1234.75563 | 1348.44058 | 1265.76793 | 1424.44567 |
| IAPLTR3-int | 10268.5163 | 10120.9682 | 10431.1095 | 10434.0897 | 10907.6774 | 10869.1775 |

| | | | | | | |
|---|---|---|---|---|---|---|
| IAPLTR4 | 2677.92522 | 2629.83444 | 2740.68173 | 2771.68527 | 2592.21795 | 2767.63484 |
| IAPLTR4_I | 1686.58306 | 1697.57665 | 1737.17054 | 1879.514 | 1792.67519 | 1789.39818 |
| ID | 19524.3119 | 19592.0386 | 19057.4813 | 20577.1053 | 22122.3198 | 21660.9655 |
| ID_B1 | 463797.551 | 466204.572 | 456606.072 | 501254.412 | 539486.815 | 528760.415 |
| ID2 | 17075.4251 | 17334.0461 | 16773.0649 | 17874.1002 | 18993.3412 | 18492.8708 |
| ID4 | 75382.3764 | 75273.9817 | 73812.0637 | 79252.871 | 85005.7856 | 82679.2597 |
| ID4_ | 80997.9322 | 81548.7635 | 79905.7446 | 84411.962 | 92705.0404 | 89094.0587 |
| IMPB_01 | 84494.0608 | 84504.2087 | 84666.0481 | 86708.9317 | 86566.2557 | 87643.5858 |
| Kanga1 | 1910.2955 | 1878.88297 | 1812.7856 | 1730.21255 | 1744.11872 | 1772.88361 |
| Kanga11a | 1167.572 | 1174.71082 | 1180.01106 | 1093.02924 | 1119.68828 | 1134.02861 |
| Kanga1a | 1132.26123 | 1200.26125 | 1137.50047 | 1114.98343 | 1071.89324 | 1065.84331 |
| Kanga1b | 680.049292 | 732.256064 | 728.251273 | 699.29714 | 703.83663 | 659.629194 |
| Kanga1c | 1131.6447 | 1120.83346 | 1096.58983 | 968.832872 | 1011.47611 | 964.747433 |
| Kanga1d | 666.17831 | 664.580363 | 646.962597 | 628.376884 | 616.855967 | 627.781621 |
| Kanga2_a | 2320.79593 | 2291.29929 | 2228.88639 | 2095.67545 | 2167.38014 | 2129.64927 |
| L1_Mm | 52680.4841 | 53255.4621 | 54360.6749 | 47912.7996 | 43408.1839 | 45924.2904 |
| L1_Mur1 | 84052.6518 | 83282.691 | 85134.824 | 73714.9413 | 66843.4189 | 70676.2258 |
| L1_Mur2 | 214493.477 | 210534.802 | 216850.018 | 189301.13 | 172436.123 | 180670.011 |
| L1_Mur3 | 253479.481 | 251861.342 | 252564.34 | 219817.846 | 210868.999 | 210389.274 |
| L1_Mus1 | 433419.745 | 428634.221 | 436414.315 | 424445.191 | 404998.277 | 419714.743 |
| L1_Mus2 | 311787.621 | 307698.79 | 314523.967 | 298134.641 | 281839.132 | 295327.559 |
| L1_Mus3 | 643796.157 | 635634.268 | 650154.953 | 605063.605 | 562698.917 | 590669.101 |
| L1_Mus4 | 184189.185 | 182208.855 | 186367.172 | 179795.732 | 168623.951 | 176806.503 |
| L1_Rod | 89882.5419 | 88895.6578 | 89337.8932 | 77232.8395 | 75372.5123 | 73924.3462 |
| L1M | 1572.81825 | 1566.22571 | 1617.98402 | 1514.30235 | 1490.93718 | 1472.13696 |
| L1M1 | 3623.19767 | 3591.722 | 3551.72311 | 3338.04882 | 3305.48926 | 3165.14274 |
| L1M2 | 120694.415 | 119924.399 | 120328.161 | 105622.677 | 103698.887 | 101414.98 |
| L1M2a | 1415.92538 | 1410.38477 | 1394.33445 | 1360.62971 | 1350.46726 | 1391.88064 |
| L1M2a1 | 51.9828914 | 48.8171721 | 45.8003816 | 35.9624355 | 41.6109905 | 46.7032852 |
| L1M2b | 62.2668577 | 75.5928835 | 65.3922014 | 89.9644066 | 95.4636569 | 76.0853758 |
| L1M2c | 2362.9085 | 2284.92989 | 2443.14346 | 2006.73559 | 2092.56338 | 2057.0336 |
| L1M3 | 41145.8095 | 40799.1741 | 41269.421 | 36734.6737 | 37374.4049 | 35795.2243 |
| L1M3a | 725.53288 | 796.744683 | 757.317712 | 638.863045 | 625.441671 | 638.035211 |
| L1M3b | 1258.74259 | 1258.87398 | 1260.19241 | 1051.39135 | 1147.17917 | 1107.17786 |
| L1M3c | 2356.63673 | 2308.27783 | 2299.86515 | 2048.7229 | 1944.58098 | 1939.21543 |
| L1M3d | 1115.94456 | 1095.36312 | 1036.10379 | 962.9986 | 1005.27811 | 932.290838 |
| L1M3de | 1155.65547 | 1173.73078 | 1195.01368 | 1112.08146 | 1098.33706 | 1076.06751 |
| L1M3e | 4501.32131 | 4581.93699 | 4286.68748 | 4377.41841 | 4345.7689 | 4274.84232 |
| L1M3f | 610.830167 | 599.516925 | 578.356221 | 577.051685 | 554.178172 | 542.917087 |
| L1M4 | 83025.2209 | 82796.9788 | 81939.9574 | 75385.6498 | 78029.8182 | 75144.3965 |
| L1M4a1 | 4013.65548 | 3908.70525 | 3739.2133 | 3594.77813 | 3570.32908 | 3517.77699 |
| L1M4a2 | 4472.20315 | 4588.95949 | 4439.14204 | 4066.87578 | 4122.94908 | 4064.87514 |

| | | | | | | |
|---|---|---|---|---|---|---|
| L1M4b | 15466.4106 | 15320.5379 | 15225.2894 | 14037.2707 | 14194.8978 | 13897.2894 |
| L1M4c | 14561.1149 | 14310.1581 | 14346.9959 | 12794.5034 | 12791.6894 | 12436.6905 |
| L1M5 | 94713.1027 | 94768.82 | 93355.276 | 85187.0192 | 88445.5821 | 84734.7771 |
| L1M6 | 1986.70858 | 1949.00751 | 1901.05228 | 1757.62127 | 1751.41061 | 1640.57199 |
| L1M6B | 239.649549 | 267.595713 | 221.279999 | 213.183179 | 251.727293 | 213.86453 |
| L1M7 | 1300.33719 | 1333.24351 | 1288.69774 | 1279.69693 | 1191.83963 | 1143.64378 |
| L1M8 | 1386.07692 | 1412.83503 | 1392.75027 | 1258.38705 | 1348.48918 | 1244.36471 |
| L1MA10 | 4622.44574 | 4678.26839 | 4626.22017 | 4201.72092 | 4049.70322 | 4080.67884 |
| L1MA4 | 54236.9191 | 53988.1766 | 54155.8364 | 48876.3499 | 48857.4489 | 47963.5498 |
| L1MA4A | 19782.1241 | 20041.0913 | 19770.2971 | 17768.6719 | 17845.8242 | 17628.2226 |
| L1MA5 | 32196.6004 | 31976.199 | 32164.3504 | 29340.856 | 29130.2649 | 28471.3742 |
| L1MA5A | 9641.86013 | 9638.25516 | 9783.13782 | 8850.20735 | 9303.85666 | 8730.36146 |
| L1MA6 | 46924.1121 | 46192.6375 | 46303.6755 | 42062.4744 | 41522.8541 | 40894.7634 |
| L1MA7 | 20771.1831 | 20777.5134 | 20950.3711 | 18827.9229 | 18792.6888 | 18261.0259 |
| L1MA8 | 25647.2757 | 25653.6589 | 25196.4891 | 22935.009 | 23008.7211 | 22302.7931 |
| L1MA9 | 29725.0812 | 29531.4235 | 29675.985 | 26512.7707 | 25925.502 | 25707.6661 |
| L1MB1 | 16971.2308 | 16852.5579 | 16752.3679 | 15208.5143 | 15449.8005 | 15301.2604 |
| L1MB2 | 20225.5116 | 20382.8159 | 19968.3807 | 19627.9072 | 20315.9603 | 19862.8129 |
| L1MB3 | 37898.3001 | 38200.5514 | 37513.0607 | 36934.7752 | 39303.4462 | 37305.4866 |
| L1MB4 | 17680.5025 | 17819.0282 | 17584.8895 | 16924.4095 | 18061.0653 | 16952.6524 |
| L1MB5 | 26286.9037 | 26323.9711 | 25903.1783 | 25604.5882 | 26820.7667 | 25596.8953 |
| L1MB7 | 49377.6234 | 49360.5314 | 48290.997 | 46438.5972 | 49060.0023 | 47049.1141 |
| L1MB8 | 40865.2454 | 41145.7075 | 40453.9679 | 38796.5943 | 41196.6516 | 38687.7079 |
| L1MC | 31381.5563 | 31308.75 | 31082.659 | 27689.227 | 28238.4878 | 27405.1762 |
| L1MC1 | 42200.3203 | 41669.6377 | 41767.8902 | 39018.2051 | 38765.6871 | 38234.8225 |
| L1MC2 | 16263.0898 | 16223.0823 | 16331.8899 | 15242.2632 | 15137.8007 | 14834.1687 |
| L1MC3 | 38985.6177 | 38643.0728 | 38736.8217 | 36179.2241 | 35839.9479 | 35788.066 |
| L1MC4 | 35167.3996 | 35100.9071 | 34838.2623 | 33414.6039 | 35300.1761 | 33694.5609 |
| L1MC4a | 19483.8791 | 19246.4802 | 18985.3602 | 18270.3344 | 19119.5987 | 18376.526 |
| L1MC5 | 21540.7232 | 21444.9588 | 20986.8561 | 19818.2936 | 20742.4328 | 19852.8795 |
| L1MC5a | 14617.7207 | 14787.6339 | 14526.3586 | 14044.0494 | 14688.9639 | 13907.3895 |
| L1MCa | 23387.1198 | 23274.7703 | 23474.5962 | 21262.9411 | 20955.6381 | 20715.6915 |
| L1MCb | 5076.42864 | 5311.66443 | 5199.78734 | 4700.78923 | 4532.69567 | 4478.51864 |
| L1MCc | 3244.62493 | 3240.37513 | 3293.81897 | 2993.91426 | 2953.34669 | 2875.64073 |
| L1MD | 32431.2553 | 32700.1232 | 32094.2407 | 28985.1974 | 30034.4185 | 28764.4923 |
| L1Md_A | 254286.176 | 251829.315 | 257524.926 | 251487.618 | 228527.615 | 244617.897 |
| L1Md_F | 95116.8224 | 95477.318 | 96797.0413 | 91177.4938 | 82487.9587 | 87205.2635 |
| L1Md_F2 | 1292547.55 | 1276865.53 | 1308203.84 | 1235878.77 | 1139429.09 | 1203376.84 |
| L1Md_F3 | 233058.427 | 230150.553 | 235363.729 | 222588.147 | 204356.672 | 217629.596 |
| L1Md_Gf | 18701.4268 | 19013.756 | 18876.7835 | 18894.9813 | 16891.041 | 17844.8678 |
| L1Md_T | 452617.194 | 444717.911 | 456943.02 | 437886.064 | 401730.376 | 426024.279 |
| L1MD1 | 16351.3188 | 16453.0524 | 16302.6115 | 15161.4569 | 15723.1239 | 15160.3444 |

| | | | | | | |
|---|---|---|---|---|---|---|
| L1MD2 | 20751.2715 | 20786.8222 | 20793.4773 | 19450.0966 | 20278.8524 | 19414.1587 |
| L1MD3 | 13276.8532 | 13605.0846 | 13106.8244 | 12420.0449 | 12643.4383 | 12318.2405 |
| L1MDa | 22878.6564 | 22754.1119 | 22864.8532 | 20885.3294 | 20928.5951 | 20373.3038 |
| L1MDb | 3501.9967 | 3316.86209 | 3392.63476 | 3135.15361 | 3189.89042 | 3095.50278 |
| L1ME1 | 38403.0113 | 38053.1038 | 37727.0647 | 34949.9918 | 36165.7536 | 34586.875 |
| L1ME2 | 18434.6775 | 18746.2241 | 18724.5182 | 16950.6561 | 17632.1127 | 16813.376 |
| L1ME2z | 9643.44026 | 9610.33892 | 9629.37914 | 9003.25414 | 9516.64142 | 8715.44342 |
| L1ME3 | 9777.53708 | 10026.5096 | 9970.92295 | 9402.24356 | 9631.6818 | 9221.83128 |
| L1ME3A | 13497.3687 | 13689.6631 | 13084.9495 | 12623.9438 | 12793.6674 | 12532.2687 |
| L1ME3B | 4472.87474 | 4514.10455 | 4490.64888 | 4159.32587 | 4454.6119 | 4111.34484 |
| L1ME3C | 2003.78701 | 2022.4786 | 1938.30754 | 1849.73428 | 1896.36358 | 1848.10308 |
| L1ME3Cz | 5924.4417 | 5953.79892 | 5918.4227 | 5498.13794 | 5601.84817 | 5472.63853 |
| L1ME3D | 2062.32268 | 2129.41802 | 1990.40274 | 1904.06575 | 2067.87704 | 1977.07369 |
| L1ME3E | 2024.34865 | 1949.98603 | 2031.8531 | 1834.01929 | 1868.74993 | 1787.18958 |
| L1ME3F | 1931.14496 | 1981.33358 | 1958.05644 | 1804.60843 | 1964.80003 | 1826.42043 |
| L1ME3G | 6182.48872 | 6075.67539 | 6021.09867 | 5785.33347 | 5847.37502 | 5670.69935 |
| L1ME4a | 8774.94001 | 8690.81498 | 8491.88581 | 7846.3493 | 8342.05158 | 7848.2735 |
| L1ME4b | 10162.2096 | 10353.0421 | 10174.8636 | 9810.10103 | 10329.1089 | 9793.48595 |
| L1ME4c | 5456.70996 | 5462.11691 | 5405.46148 | 5046.20239 | 5292.12851 | 5125.22457 |
| L1ME5 | 1066.2894 | 1151.11895 | 1122.16231 | 1068.85296 | 1111.78745 | 1051.57381 |
| L1MEa | 1129.64411 | 1115.60762 | 1117.6995 | 988.544564 | 1026.38441 | 1043.79534 |
| L1MEb | 2190.99389 | 2203.62342 | 2140.0679 | 1956.13625 | 1996.65426 | 1951.72422 |
| L1MEc | 37401.7045 | 37069.4857 | 36672.8885 | 33616.6579 | 34710.8032 | 33547.5628 |
| L1MEd | 18322.2895 | 18246.6199 | 18462.1141 | 16623.3721 | 17275.6967 | 16513.8246 |
| L1MEf | 20084.4143 | 20001.2576 | 19777.1803 | 17819.2963 | 18136.062 | 17312.605 |
| L1MEg | 20733.7301 | 20715.7071 | 20374.4907 | 18368.1964 | 18744.5012 | 18114.9733 |
| L1MEg1 | 585.868376 | 578.212392 | 575.803213 | 585.959095 | 560.742023 | 493.992028 |
| L1MEg2 | 739.325223 | 685.724204 | 679.313858 | 684.349694 | 668.710496 | 631.333413 |
| L1MEh | 1163.06018 | 1207.28132 | 1142.11322 | 1057.33089 | 1004.166 | 1002.30625 |
| L1MEi | 2819.47498 | 2890.56987 | 2905.49016 | 2687.59682 | 2686.77614 | 2577.44165 |
| L1MEj | 1135.84248 | 1132.6704 | 1119.39366 | 1045.70831 | 1047.29139 | 983.408279 |
| L1P5 | 613.880336 | 630.866194 | 559.782499 | 581.827302 | 564.116022 | 567.289174 |
| L1PB4 | 1171.02113 | 1127.20054 | 1169.10782 | 1085.61573 | 1011.71919 | 1017.49434 |
| L1VL1 | 39111.2739 | 38616.7944 | 39603.6271 | 35153.3788 | 33056.4391 | 34448.8492 |
| L1VL2 | 95931.2112 | 95076.3416 | 96354.6639 | 87310.9647 | 84409.4883 | 85392.658 |
| L1VL4 | 129186.337 | 127444.764 | 130159.061 | 115398.678 | 109526.861 | 111389.816 |
| L2 | 108598.444 | 108472.246 | 107574.179 | 109463.323 | 112152.374 | 111111.05 |
| L2a | 137265.249 | 137809.3 | 136138.759 | 138803.732 | 144312.323 | 141747.233 |
| L2b | 68284.7379 | 69120.5889 | 68251.6424 | 73502.3798 | 76734.3058 | 76484.4273 |
| L2c | 60975.0901 | 61234.0952 | 60196.7431 | 62967.6814 | 65335.077 | 64119.1536 |
| L3 | 46547.5126 | 46479.4541 | 46728.089 | 45960.8679 | 47145.4183 | 46989.7404 |
| L3b | 9890.10656 | 9852.79357 | 9781.11894 | 9922.28615 | 10127.4361 | 10111.1586 |

| | | | | | | |
|---|---|---|---|---|---|---|
| L4_A_Mam | 3877.22359 | 3995.16151 | 3975.60806 | 3803.90018 | 3783.8996 | 3701.84191 |
| L4_B_Mam | 2099.35723 | 2146.07154 | 2027.82221 | 2056.81202 | 2134.21386 | 2062.35623 |
| L4_C_Mam | 3438.4627 | 3384.12933 | 3384.56305 | 3254.16663 | 3325.66759 | 3286.0486 |
| L5 | 713.235215 | 723.438933 | 745.630595 | 707.46353 | 719.491801 | 698.731845 |
| LFSINE_Vert | 5444.57325 | 5259.18022 | 5564.34998 | 5569.10233 | 5511.8502 | 5697.53692 |
| Looper | 1183.6019 | 1156.26257 | 1187.70942 | 1139.28014 | 1357.3763 | 1225.40746 |
| LSU-rRNA_Hsa | 6153.03609 | 6252.42768 | 6362.51201 | 7059.55264 | 10486.1874 | 8278.39509 |
| LTR10_RN | 3542.98525 | 3513.27576 | 3624.18191 | 3778.19589 | 3812.68186 | 3608.79407 |
| LTR101_Mam | 1090.27291 | 1118.05636 | 1162.89944 | 1093.8869 | 1013.21233 | 1065.71512 |
| LTR102_Mam | 1304.69854 | 1398.55097 | 1413.47081 | 1403.23052 | 1347.88329 | 1341.91184 |
| LTR103_Mam | 353.859978 | 387.351409 | 349.354313 | 359.76043 | 366.179402 | 340.734965 |
| LTR103b_Mam | 303.024177 | 297.473454 | 309.171161 | 285.146929 | 285.946126 | 317.768398 |
| LTR104_Mam | 1484.1332 | 1528.34978 | 1545.87159 | 1526.46873 | 1591.42901 | 1531.77314 |
| LTR105_Mam | 821.228396 | 804.25715 | 779.154871 | 821.845069 | 892.15748 | 796.878366 |
| LTR106_Mam | 1007.1178 | 1001.15664 | 971.905993 | 899.410306 | 866.808156 | 845.793792 |
| LTR107_Mam | 566.022355 | 528.579211 | 554.048159 | 552.076066 | 517.37293 | 490.420475 |
| LTR108a_Mam | 237.983437 | 216.412023 | 261.249679 | 230.376917 | 220.155642 | 237.719808 |
| LTR108b_Mam | 125.97649 | 129.960822 | 140.432524 | 137.68928 | 130.651172 | 147.90455 |
| LTR108c_Mam | 114.433242 | 108.899301 | 106.377815 | 113.279728 | 95.3822211 | 96.283474 |
| LTR108d_Mam | 190.939114 | 194.288546 | 161.915551 | 199.779883 | 164.425161 | 150.580806 |
| LTR108e_Mam | 220.669088 | 223.840551 | 204.989422 | 179.261272 | 178.342893 | 191.796798 |
| LTR109A2 | 184.924756 | 168.0024 | 173.473465 | 165.392408 | 170.727093 | 166.285708 |
| LTR16A | 9614.93916 | 9845.85825 | 9760.05853 | 10223.4667 | 10115.9559 | 10362.8545 |
| LTR16A1 | 3989.23526 | 4068.87532 | 4140.24299 | 4301.15647 | 4063.29765 | 4240.0682 |
| LTR16A2 | 2585.71292 | 2645.42549 | 2572.38862 | 2608.19352 | 2519.58409 | 2603.90683 |
| LTR16B | 1813.56928 | 1730.88135 | 1866.24653 | 1776.11452 | 1731.77433 | 1766.42503 |
| LTR16B1 | 2045.14464 | 2056.35448 | 2124.0286 | 2161.33227 | 2160.59203 | 2119.13526 |
| LTR16B2 | 2656.17874 | 2621.26113 | 2608.97712 | 2676.57259 | 2669.82837 | 2664.68909 |
| LTR16C | 8379.62017 | 8404.77082 | 8580.49299 | 8961.86712 | 8666.20548 | 8702.86876 |
| LTR16D | 1177.47482 | 1271.77166 | 1159.49471 | 1267.85722 | 1196.02311 | 1224.66012 |
| LTR16D1 | 537.13802 | 519.273122 | 548.339532 | 562.620544 | 609.504527 | 542.857807 |
| LTR16D2 | 354.12211 | 366.454223 | 325.431061 | 360.784484 | 322.587276 | 335.48784 |
| LTR16E1 | 3176.52773 | 3119.63828 | 3108.1165 | 3396.02328 | 3218.71157 | 3248.62811 |
| LTR16E2 | 2659.78568 | 2769.59102 | 2693.37928 | 2662.0324 | 2619.85288 | 2698.44227 |
| LTR23-int | 26.9031411 | 37.5514062 | 40.6900367 | 34.9848405 | 30.7840626 | 34.497482 |
| LTR27C | 41.3057269 | 30.5308318 | 41.5253399 | 37.0642469 | 24.6838889 | 32.4170747 |
| LTR28 | 50.0546477 | 55.0208265 | 53.9092641 | 55.8329446 | 71.8298865 | 60.8050685 |
| LTR28B | 386.692023 | 432.413663 | 408.446734 | 438.990243 | 387.671607 | 412.587073 |
| LTR29 | 188.276377 | 176.655499 | 213.032621 | 188.339357 | 171.98019 | 218.887708 |
| LTR31 | 1219.20783 | 1252.26156 | 1214.68977 | 1251.3111 | 1242.33958 | 1292.78918 |
| LTR32 | 33.33062 | 27.5922267 | 40.0189428 | 26.9621069 | 37.3691882 | 32.4302481 |
| LTR33 | 9222.41889 | 9266.01282 | 9277.40678 | 9833.71563 | 9811.12765 | 9819.97393 |

| | | | | | | |
|---|---|---|---|---|---|---|
| LTR33A | 3345.64015 | 3303.96775 | 3444.92484 | 3477.73644 | 3565.94385 | 3704.14931 |
| LTR33A_ | 3019.12946 | 3085.67675 | 3098.52477 | 3266.65309 | 3292.07287 | 3180.17276 |
| LTR33B | 1589.11658 | 1586.96281 | 1596.27823 | 1572.99839 | 1511.35555 | 1491.59735 |
| LTR33C | 1442.09927 | 1472.75464 | 1461.07533 | 1494.66083 | 1526.76947 | 1556.0069 |
| LTR34 | 20.987344 | 27.2653777 | 18.6251491 | 19.0635899 | 28.5622825 | 17.2553277 |
| LTR36 | 25.1059635 | 17.4694203 | 17.7241622 | 21.8645777 | 19.2704249 | 16.2480575 |
| LTR37-int | 1574.89172 | 1597.40973 | 1533.86102 | 1433.98588 | 1479.21995 | 1353.50253 |
| LTR37A | 2956.26494 | 2991.14332 | 2874.22268 | 2736.19121 | 2645.93146 | 2612.63732 |
| LTR37B | 2187.81999 | 2173.66295 | 2105.81397 | 1915.88223 | 1907.19173 | 1863.29472 |
| LTR39 | 35.5472091 | 38.3681239 | 45.063604 | 38.0418419 | 45.6510347 | 44.6228779 |
| LTR40a | 4923.28531 | 4871.7472 | 4920.78715 | 4677.32646 | 4552.95849 | 4669.68007 |
| LTR40A1 | 572.856139 | 628.661583 | 625.838666 | 571.32598 | 566.680546 | 566.574764 |
| LTR40b | 3716.41446 | 3779.97345 | 3804.57049 | 3655.58108 | 3730.4079 | 3649.52923 |
| LTR40c | 1622.43986 | 1645.82108 | 1565.10711 | 1710.77301 | 1731.02929 | 1685.39552 |
| LTR41 | 3249.26152 | 3146.00573 | 3205.60868 | 3509.43529 | 3469.47079 | 3411.69086 |
| LTR41B | 1774.02194 | 1837.17183 | 1744.36915 | 1855.71319 | 1831.74415 | 1906.61746 |
| LTR44 | 9.26060265 | 13.2248359 | 7.14221185 | 3.86392068 | 8.12019588 | 4.06860106 |
| LTR45 | 139.198433 | 129.634681 | 101.807196 | 137.033597 | 144.306375 | 128.661029 |
| LTR45C | 102.365728 | 81.4704984 | 86.8188369 | 96.46622 | 117.279163 | 109.012672 |
| LTR48 | 3401.85962 | 3389.68324 | 3349.42239 | 3507.86412 | 3551.33819 | 3478.07898 |
| LTR48B | 1917.78566 | 1974.47774 | 1987.33355 | 1940.10264 | 2022.20779 | 1967.03697 |
| LTR50 | 3198.30776 | 3271.39543 | 3178.26708 | 3283.39214 | 3339.06149 | 3345.88896 |
| LTR52 | 1382.29226 | 1329.24365 | 1312.18475 | 1321.03283 | 1268.71897 | 1318.31344 |
| LTR52-int | 390.889282 | 362.45427 | 342.981721 | 375.243132 | 348.200961 | 364.047088 |
| LTR53 | 1487.71392 | 1567.04202 | 1529.3347 | 1479.18143 | 1520.34599 | 1524.51501 |
| LTR53B | 575.26356 | 636.090415 | 610.16992 | 636.446077 | 625.744308 | 588.181921 |
| LTR55 | 1027.2983 | 1041.97401 | 1084.74563 | 1029.37981 | 983.98583 | 1001.90446 |
| LTR58 | 41.3581533 | 38.5311436 | 35.5140081 | 41.649749 | 46.1359865 | 32.4170747 |
| LTR59 | 26.9555675 | 35.3476042 | 29.1414157 | 34.1779212 | 28.0773307 | 32.4961149 |
| LTR64 | 32.0975507 | 19.9185614 | 30.9762315 | 21.6550234 | 26.703911 | 28.3353003 |
| LTR65 | 1010.40336 | 998.872545 | 951.089096 | 990.189701 | 998.647991 | 957.577981 |
| LTR67B | 3617.85175 | 3597.60376 | 3432.22179 | 3728.07349 | 3779.6523 | 3795.85652 |
| LTR68 | 571.05267 | 520.416081 | 516.652236 | 481.431262 | 468.389835 | 470.390091 |
| LTR69 | 522.125191 | 498.94438 | 511.607575 | 472.632908 | 457.359929 | 497.731294 |
| LTR72_RN | 5908.28859 | 5913.1467 | 5819.23419 | 5916.68073 | 5732.53701 | 5745.85247 |
| LTR73 | 884.957426 | 927.195981 | 958.336962 | 993.188384 | 964.411547 | 980.577482 |
| LTR75 | 569.714748 | 528.577794 | 519.486565 | 598.986439 | 590.232881 | 560.471863 |
| LTR75_1 | 42.4863698 | 32.6533263 | 31.0090733 | 36.4280033 | 27.0272122 | 22.331199 |
| LTR75B | 427.650163 | 453.883948 | 432.764089 | 391.800627 | 446.393235 | 457.058457 |
| LTR77 | 121.910297 | 107.267079 | 106.016554 | 116.526844 | 114.087479 | 109.549241 |
| LTR78 | 3755.67923 | 3748.29804 | 3824.34727 | 3743.57514 | 3811.794 | 3601.15391 |
| LTR78B | 2412.11594 | 2445.99393 | 2416.84566 | 2336.85763 | 2421.26783 | 2407.35386 |

| | | | | | | |
|---|---|---|---|---|---|---|
| LTR79 | 2246.74781 | 2270.07485 | 2287.15 | 2243.14166 | 2242.60346 | 2280.03248 |
| LTR80A | 739.548035 | 741.233786 | 724.762271 | 716.735033 | 578.659082 | 647.834812 |
| LTR80B | 749.268417 | 694.54012 | 709.046427 | 651.12186 | 534.34136 | 660.132829 |
| LTR81 | 388.450405 | 359.189018 | 355.635508 | 418.165369 | 372.764528 | 411.652256 |
| LTR81A | 1024.79389 | 1071.76903 | 1073.56044 | 1017.28788 | 1045.20938 | 1010.63751 |
| LTR81B | 1242.98216 | 1247.20036 | 1233.33134 | 1214.64603 | 1169.88192 | 1150.3851 |
| LTR81C | 618.247981 | 626.131438 | 595.967601 | 593.547558 | 662.023881 | 572.147684 |
| LTR82A | 2396.57203 | 2344.60477 | 2338.54406 | 2308.59552 | 2336.97934 | 2252.82301 |
| LTR82B | 1841.522 | 1866.31365 | 1819.89713 | 1862.17991 | 1814.65296 | 1828.68526 |
| LTR83 | 959.639908 | 974.544154 | 942.555434 | 885.789387 | 891.673139 | 911.213664 |
| LTR84a | 812.151285 | 811.196113 | 764.117248 | 806.431567 | 829.862537 | 766.436311 |
| LTR84b | 1155.75351 | 1165.48655 | 1190.1261 | 1145.56531 | 1183.13361 | 1133.98909 |
| LTR85a | 1303.30085 | 1202.95513 | 1259.02372 | 1234.93565 | 1254.58003 | 1204.42909 |
| LTR85b | 718.043766 | 716.092825 | 669.468685 | 699.626633 | 668.889759 | 636.13923 |
| LTR85c | 710.867638 | 684.988895 | 693.579696 | 713.158424 | 691.757825 | 668.444332 |
| LTR86A1 | 515.908466 | 546.702531 | 555.580269 | 548.060909 | 536.966046 | 542.857807 |
| LTR86A2 | 378.565928 | 366.455741 | 374.112869 | 357.549226 | 336.162264 | 377.71769 |
| LTR86B1 | 274.330674 | 255.024879 | 309.072635 | 286.846105 | 262.291692 | 259.586892 |
| LTR86B2 | 220.150591 | 224.982195 | 228.372949 | 203.112337 | 201.289343 | 224.447454 |
| LTR86C | 488.972296 | 454.211404 | 538.731381 | 459.112976 | 474.752539 | 470.350571 |
| LTR87 | 1352.69912 | 1322.79577 | 1284.4227 | 1333.43531 | 1438.7794 | 1359.76302 |
| LTR88a | 318.023899 | 333.719488 | 327.322267 | 342.306888 | 370.683734 | 354.797229 |
| LTR88b | 1027.08178 | 960.911263 | 1025.65538 | 1017.75675 | 974.511047 | 1027.85332 |
| LTR88c | 666.184077 | 633.968426 | 680.672466 | 679.309995 | 665.942382 | 593.896208 |
| LTR89 | 829.314647 | 848.828423 | 858.216793 | 829.758747 | 791.077991 | 857.288233 |
| LTR9 | 60.7716561 | 51.9192017 | 69.0946748 | 51.7789089 | 60.5581142 | 60.857762 |
| LTR90A | 527.373075 | 536.66083 | 537.356351 | 497.926648 | 551.611816 | 479.626777 |
| LTR90B | 492.074367 | 454.700463 | 469.303323 | 453.685466 | 449.725906 | 415.523257 |
| LTR91 | 301.135777 | 296.983383 | 352.374236 | 312.69882 | 274.594139 | 267.180939 |
| LTR9A1 | 39.6595373 | 49.6330803 | 46.9969453 | 49.3387117 | 52.9228701 | 53.7410035 |
| LTR9D | 26.8245015 | 26.4492672 | 40.5915111 | 42.7126819 | 47.5094062 | 46.6901118 |
| LTRIS_Mm | 5226.84161 | 5189.62281 | 5472.27454 | 5632.9898 | 5839.95182 | 6096.05062 |
| LTRIS_Mus | 9597.29924 | 9457.44205 | 9503.63787 | 10529.7889 | 11905.8988 | 11720.1029 |
| LTRIS2 | 13292.0265 | 13209.5008 | 13364.0382 | 13863.7875 | 13262.7755 | 13769.4037 |
| LTRIS3 | 1525.17784 | 1457.24611 | 1424.5268 | 1524.67994 | 1721.95986 | 1598.19065 |
| LTRIS4 | 1107.52907 | 1145.24164 | 1080.78259 | 1215.13104 | 1106.35516 | 1155.32619 |
| LTRIS4A | 1464.33331 | 1457.81658 | 1580.4146 | 1569.67779 | 1504.44834 | 1615.6304 |
| LTRIS4B | 624.793419 | 531.27187 | 574.508124 | 596.22812 | 619.13974 | 672.015884 |
| LTRIS5 | 659.671143 | 656.826504 | 631.908553 | 700.953648 | 673.112118 | 670.166011 |
| LTRIS6 | 1017.33518 | 1110.38502 | 1075.23601 | 1103.00765 | 1057.65159 | 1115.00549 |
| Lx | 334383.985 | 331689.206 | 338586.664 | 337812.683 | 316216.059 | 334590.034 |
| Lx10 | 68031.5236 | 67074.6422 | 68379.6041 | 62141.8987 | 59269.3354 | 60161.6262 |

| | | | | | | |
|---|---|---|---|---|---|---|
| Lx2 | 215296.71 | 213200.901 | 219403.725 | 224145.967 | 207577.677 | 225019.46 |
| Lx2A | 17006.7245 | 17084.6497 | 17526.2005 | 18064.51 | 16690.8851 | 18209.4388 |
| Lx2A1 | 26786.7267 | 26254.5712 | 27114.6383 | 28831.939 | 26531.0952 | 28655.0795 |
| Lx2B | 77770.1529 | 76521.4233 | 78609.628 | 78600.5099 | 72555.8874 | 78574.0842 |
| Lx2B2 | 131534.023 | 129781.748 | 133779.725 | 134639.37 | 122680.078 | 133040.819 |
| Lx3_Mus | 177781.049 | 175688.765 | 180648.123 | 179532.458 | 165526.04 | 178701.716 |
| Lx3A | 106922.057 | 105232.745 | 109015.869 | 109993.282 | 99207.0364 | 109030.942 |
| Lx3B | 66877.53 | 66094.3234 | 68101.6706 | 69992.8207 | 63555.7139 | 69272.9584 |
| Lx3C | 79366.0383 | 78324.696 | 80813.1798 | 82146.2069 | 74190.8155 | 80694.2338 |
| Lx4A | 126101.353 | 124398.178 | 127918.197 | 128881.728 | 119345.164 | 127430.808 |
| Lx4B | 64278.7073 | 63764.3998 | 65315.2194 | 66663.5754 | 61663.1298 | 65270.4868 |
| Lx5 | 208471.388 | 206801.056 | 210310.306 | 212050.15 | 198976.596 | 210546.343 |
| Lx5b | 87819.943 | 86578.6791 | 88668.2769 | 85687.8143 | 80228.1887 | 84399.4897 |
| Lx5c | 166472.299 | 164816.013 | 168669.089 | 162749.426 | 164181.239 | 164507.173 |
| Lx6 | 188567.041 | 186111.244 | 190213.024 | 174307.744 | 163230.664 | 169819.349 |
| Lx7 | 343211.075 | 341048.084 | 345688.088 | 314906.547 | 301462.019 | 306302.484 |
| Lx8 | 401223.677 | 399670.098 | 401358.032 | 393203.728 | 385711.556 | 389855.06 |
| Lx8b | 192527.798 | 190540.583 | 192986.278 | 179249.692 | 171301.239 | 175049.461 |
| Lx9 | 362418.004 | 359213.288 | 364242.264 | 334051.467 | 320677.027 | 325863.857 |
| MADE2 | 746.703717 | 676.011678 | 697.709114 | 656.327957 | 677.051283 | 641.797777 |
| Mam_R4 | 837.51309 | 851.930351 | 842.768647 | 770.077043 | 767.384059 | 756.192355 |
| MamGyp-int | 277.623053 | 294.698881 | 281.259151 | 264.259946 | 304.610056 | 349.550103 |
| MamGypLTR1a | 832.804149 | 829.31741 | 832.219539 | 811.944415 | 817.277507 | 768.671259 |
| MamGypLTR1b | 794.548591 | 818.134672 | 770.764668 | 764.681319 | 790.553542 | 807.510444 |
| MamGypLTR1c | 775.961852 | 806.625591 | 723.359364 | 767.858259 | 818.932899 | 789.623287 |
| MamGypLTR1d | 678.035593 | 643.274717 | 676.313156 | 648.053488 | 650.750888 | 671.002027 |
| MamGypLTR2b | 544.837888 | 541.070862 | 532.443057 | 522.786119 | 565.105369 | 521.109284 |
| MamGypLTR2c | 540.811015 | 534.048163 | 599.221745 | 577.687928 | 532.846397 | 533.420474 |
| MamGypLTR3 | 613.237064 | 581.232608 | 585.230734 | 600.08825 | 542.462166 | 590.80853 |
| MamGypLTR3a | 966.276569 | 987.116708 | 933.207853 | 1024.82133 | 977.299215 | 956.850398 |
| MamRep1151 | 1542.10424 | 1642.63501 | 1565.51331 | 1422.4676 | 1395.09006 | 1395.84081 |
| MamRep1161 | 41.4892194 | 37.9595627 | 31.7786928 | 38.6780855 | 40.0358128 | 46.6505917 |
| MamRep137 | 1820.63322 | 1840.92543 | 1917.41999 | 1829.09243 | 1961.1635 | 1807.03859 |
| MamRep1527 | 1848.28344 | 1806.55776 | 1896.79366 | 1792.03956 | 1806.25079 | 1845.01187 |
| MamRep1879 | 491.221914 | 516.089178 | 500.033241 | 510.767668 | 526.563908 | 502.26401 |
| MamRep1894 | 137.545952 | 145.960636 | 143.970788 | 162.525521 | 154.385822 | 161.302051 |
| MamRep38 | 1095.32315 | 1103.36242 | 1125.37928 | 1188.47568 | 1259.56993 | 1192.74363 |
| MamRep4096 | 993.330698 | 1059.44262 | 998.902759 | 950.413594 | 994.671771 | 979.280398 |
| MamRep434 | 2900.16239 | 2841.42868 | 3015.7204 | 2821.36863 | 2869.62275 | 2707.50161 |
| MamRep488 | 142.150041 | 160.981522 | 158.320896 | 141.661768 | 174.483943 | 178.228044 |
| MamRep564 | 693.251313 | 680.009811 | 715.283323 | 689.090711 | 635.702822 | 642.785287 |
| MamRep605 | 3871.47766 | 3803.15494 | 3901.11243 | 3702.01452 | 3632.827 | 3667.69962 |

| | | | | | | |
|---|---|---|---|---|---|---|
| MamSINE1 | 2091.49379 | 2172.44041 | 2028.12924 | 2199.04792 | 2200.12345 | 2209.39842 |
| MamTip1 | 384.101633 | 398.536677 | 405.661669 | 394.535716 | 424.919863 | 415.753791 |
| MamTip2 | 3720.57083 | 3601.35898 | 3675.93227 | 3733.92768 | 3738.61136 | 3672.03168 |
| MamTip3 | 1020.45036 | 988.25987 | 1032.11504 | 1003.33698 | 1029.97656 | 967.611163 |
| MARINER1_EC | 5.19440956 | 6.85714996 | 8.84566015 | 8.91499068 | 3.55509241 | 7.10358513 |
| MARNA | 3224.4979 | 3323.80217 | 3243.54713 | 3100.53005 | 3188.39302 | 3106.66838 |
| MER101B | 102.365728 | 104.001625 | 102.004248 | 109.477915 | 88.9587461 | 81.121727 |
| MER102a | 2251.62819 | 2287.46246 | 2265.63197 | 2308.04792 | 2517.01956 | 2365.76242 |
| MER102b | 3207.87715 | 3142.73805 | 3067.8627 | 3318.00384 | 3828.19116 | 3644.4569 |
| MER102c | 1983.37006 | 1974.47805 | 1962.53503 | 2082.12189 | 2191.61796 | 2124.13868 |
| MER103C | 5394.27534 | 5481.06049 | 5400.33688 | 5280.22231 | 5337.86098 | 5331.70534 |
| MER104 | 1356.13672 | 1404.59039 | 1282.47293 | 1265.05954 | 1269.3449 | 1177.75923 |
| MER105 | 794.12184 | 812.909541 | 808.573278 | 819.346554 | 836.022872 | 856.192289 |
| MER106A | 622.760323 | 651.436936 | 634.510824 | 629.882155 | 663.92236 | 665.580601 |
| MER106B | 801.297444 | 800.909174 | 875.516175 | 796.729097 | 788.353036 | 767.190241 |
| MER110 | 844.439668 | 839.03206 | 868.87372 | 766.989719 | 775.425261 | 772.193658 |
| MER110-int | 273.471405 | 254.126853 | 289.349448 | 256.594213 | 280.855963 | 252.907901 |
| MER110A | 1172.33389 | 1157.81177 | 1096.2707 | 1012.67487 | 1002.81202 | 1085.0539 |
| MER112 | 1689.0298 | 1680.84082 | 1687.72624 | 1577.86692 | 1664.85596 | 1507.68428 |
| MER113 | 3780.55085 | 3880.05613 | 3832.59961 | 3747.55472 | 3950.06281 | 3822.4843 |
| MER113A | 698.866182 | 718.540246 | 712.303371 | 670.817904 | 712.483106 | 676.765961 |
| MER113B | 317.709341 | 389.229273 | 351.02492 | 296.35088 | 329.415488 | 340.682272 |
| MER115 | 3191.94581 | 3338.00866 | 3337.40189 | 3457.03958 | 3535.48309 | 3548.28186 |
| MER117 | 3743.06753 | 3876.30161 | 3859.50498 | 3804.67629 | 3878.21165 | 3937.43825 |
| MER119 | 2545.7404 | 2558.07819 | 2438.42721 | 2433.26932 | 2639.45027 | 2457.85569 |
| MER121 | 3742.01166 | 3784.38278 | 3727.71177 | 3451.97616 | 3551.88513 | 3332.6926 |
| MER123 | 210.562323 | 209.635674 | 230.789625 | 182.946936 | 200.966653 | 180.677306 |
| MER124 | 1420.16144 | 1347.28658 | 1364.25424 | 1339.44783 | 1287.18236 | 1290.85722 |
| MER125 | 490.054902 | 476.579916 | 480.007348 | 447.121544 | 405.246532 | 445.234682 |
| MER126 | 701.082247 | 688.418886 | 652.678352 | 608.188253 | 635.603164 | 630.619004 |
| MER127 | 723.473046 | 737.479475 | 663.734345 | 679.375894 | 681.496064 | 680.330927 |
| MER129 | 378.874195 | 373.066945 | 381.935467 | 369.020562 | 348.40394 | 387.451424 |
| MER130 | 574.338234 | 566.375551 | 557.440799 | 584.60127 | 597.26163 | 610.654487 |
| MER131 | 1250.40679 | 1260.2639 | 1209.01182 | 1206.61193 | 1220.70516 | 1144.06533 |
| MER132 | 87.7010105 | 79.1845794 | 112.389147 | 104.357157 | 76.8799418 | 77.0399526 |
| MER133A | 134.010839 | 120.001237 | 129.74492 | 134.632278 | 139.417971 | 139.767348 |
| MER133B | 84.3562049 | 88.1646286 | 89.1276954 | 64.1505704 | 60.9822944 | 89.1403687 |
| MER134 | 246.679932 | 231.840155 | 243.304917 | 223.553192 | 240.617172 | 214.977187 |
| MER135 | 1118.51293 | 1139.36382 | 1163.94605 | 1170.48261 | 1024.88761 | 1137.70554 |
| MER136 | 140.352863 | 126.205701 | 151.784094 | 129.666546 | 106.330692 | 116.442052 |
| MER2 | 49971.0516 | 50325.3699 | 49909.0761 | 48596.5396 | 50769.2957 | 48453.6353 |
| MER20 | 31747.0843 | 32012.7074 | 31866.1691 | 33156.1082 | 34746.8237 | 34006.3369 |

| | | | | | | |
|---|---|---|---|---|---|---|
| MER20B | 5095.9124 | 5018.35719 | 4991.79938 | 4939.04522 | 5009.10971 | 4867.94911 |
| MER21-int | 640.63878 | 659.764198 | 626.401942 | 623.197808 | 719.088285 | 621.467945 |
| MER21B | 11996.2032 | 12219.3609 | 12096.5995 | 12633.4785 | 12985.863 | 13026.399 |
| MER21C | 17661.2285 | 17863.1957 | 17659.2609 | 18056.4184 | 18950.4358 | 18246.2148 |
| MER2B | 4261.38237 | 4267.0821 | 4232.62547 | 4195.32767 | 4325.37181 | 4117.95747 |
| MER3 | 14230.8662 | 14321.9196 | 14211.1476 | 14245.8168 | 14577.6943 | 14439.7115 |
| MER31-int | 984.432361 | 977.810013 | 934.158103 | 951.193005 | 901.610378 | 841.030542 |
| MER31A | 4854.04049 | 4755.16968 | 4817.91196 | 4988.78361 | 4819.91617 | 4855.69671 |
| MER31B | 3531.71305 | 3518.33605 | 3574.15422 | 3632.69294 | 3540.45354 | 3543.20599 |
| MER33 | 13353.8986 | 13502.4779 | 13205.3239 | 12559.3613 | 13430.4046 | 12851.3058 |
| MER34 | 3813.17948 | 3861.85221 | 3821.32735 | 3702.26295 | 3572.32173 | 3654.98713 |
| MER34-int | 500.967461 | 490.783376 | 524.531226 | 479.755804 | 519.475609 | 514.216472 |
| MER34A | 4089.08452 | 4184.95336 | 4129.71743 | 3966.30246 | 3917.29028 | 3965.23344 |
| MER34A1 | 3132.04601 | 3270.66022 | 3268.19723 | 3061.55708 | 2985.26353 | 3012.7217 |
| MER34B-int | 482.722019 | 457.720679 | 479.467622 | 422.82056 | 445.402667 | 424.759943 |
| MER44A | 3829.09719 | 3945.7681 | 3868.02006 | 3514.27155 | 3826.47927 | 3686.26215 |
| MER44B | 9367.99867 | 9295.89026 | 9217.70398 | 9089.55379 | 9255.56333 | 8953.37351 |
| MER44C | 1326.65 | 1385.9775 | 1276.94277 | 1246.74076 | 1271.44819 | 1222.1617 |
| MER44D | 4712.72875 | 4780.6383 | 4575.19883 | 4491.58135 | 4515.99343 | 4406.43295 |
| MER45A | 3500.59063 | 3583.31424 | 3588.56938 | 3755.72539 | 3850.59922 | 3756.28415 |
| MER45B | 4504.61998 | 4432.55385 | 4436.10569 | 4376.86274 | 4529.24451 | 4289.42798 |
| MER45C | 1733.43026 | 1635.37041 | 1779.7775 | 1876.85239 | 1841.31798 | 1901.08662 |
| MER45R | 1845.49016 | 1805.57651 | 1839.66525 | 1833.2702 | 1812.77697 | 1780.02672 |
| MER46C | 4020.78338 | 4118.34487 | 4017.59815 | 3734.86494 | 3884.73661 | 3817.06287 |
| MER47A | 675.051482 | 723.928903 | 678.065867 | 700.883471 | 730.499212 | 700.979967 |
| MER47B | 2889.76046 | 3033.59352 | 2942.14434 | 2952.622 | 3001.00074 | 2958.92495 |
| MER47C | 48.8808202 | 43.265799 | 33.5314039 | 50.2233883 | 41.2074744 | 41.5747204 |
| MER49 | 26.6672221 | 30.3680144 | 15.1197269 | 28.3199319 | 27.3906208 | 32.4302481 |
| MER4B-int | 248.286801 | 260.575442 | 255.327581 | 273.260275 | 239.001887 | 253.688177 |
| MER4CL34 | 19.6232086 | 15.0206839 | 19.4932942 | 25.006917 | 19.7954842 | 12.1926298 |
| MER50 | 41.3581533 | 49.3066361 | 43.3929975 | 45.0481018 | 33.0872786 | 29.4084374 |
| MER50-int | 89.734107 | 90.6131627 | 76.0398354 | 86.9149858 | 116.027287 | 85.1376345 |
| MER50B | 59.0793314 | 58.9395737 | 57.5296328 | 61.4349203 | 59.3062377 | 57.8359513 |
| MER51-int | 48.4021669 | 49.7967071 | 32.8110472 | 34.5192726 | 53.8927738 | 42.5292971 |
| MER52-int | 1418.59389 | 1369.65246 | 1431.467 | 1406.60943 | 1442.13152 | 1403.88226 |
| MER53 | 7084.18433 | 7029.97778 | 7069.40296 | 6634.04345 | 6788.79757 | 6559.44399 |
| MER54A | 2083.99157 | 2100.52134 | 2142.26894 | 2109.69604 | 2078.27918 | 2194.05529 |
| MER54B | 914.425792 | 939.685811 | 869.920331 | 868.13057 | 903.509467 | 946.81063 |
| MER57-int | 998.381459 | 1050.95386 | 1023.80198 | 952.427102 | 981.498467 | 943.064776 |
| MER57C2 | 2597.28972 | 2573.26017 | 2535.45248 | 2383.46883 | 2417.42832 | 2425.04391 |
| MER57D | 1849.58833 | 1900.76009 | 1885.25433 | 1646.94814 | 1636.86128 | 1624.1167 |
| MER57E1 | 1216.57759 | 1203.77387 | 1191.61391 | 1150.12709 | 1157.45915 | 1165.31985 |

| | | | | | | |
|---|---|---|---|---|---|---|
| MER57E2 | 374.755576 | 388.250648 | 382.40951 | 350.674763 | 337.333926 | 351.278369 |
| MER57E3 | 19.7280614 | 31.8368111 | 15.8893464 | 13.5004928 | 15.9170905 | 13.1867266 |
| MER58A | 19582.5314 | 20009.5947 | 19864.7867 | 19442.0275 | 20017.7445 | 19037.3393 |
| MER58B | 15060.6097 | 15176.4628 | 14839.3844 | 14773.9759 | 15145.7574 | 14899.6185 |
| MER58C | 1818.92674 | 1851.70133 | 1827.03934 | 1690.30892 | 1790.23465 | 1766.54054 |
| MER58D | 1251.20052 | 1284.58935 | 1203.94144 | 1255.34948 | 1266.92137 | 1201.38752 |
| MER5A | 38428.0271 | 38643.99 | 38918.0666 | 39847.9721 | 40270.5273 | 39945.7702 |
| MER5A1 | 17097.0269 | 17298.3694 | 17066.0889 | 17657.3911 | 17882.6184 | 17845.1318 |
| MER5B | 21707.9409 | 21767.7446 | 21605.5525 | 21873.1947 | 22520.9003 | 22336.108 |
| MER5C | 1187.73363 | 1285.16093 | 1230.19214 | 1128.73994 | 1188.60906 | 1123.45885 |
| MER5C1 | 386.567248 | 391.842243 | 364.816715 | 330.582841 | 405.648828 | 376.98352 |
| MER63A | 3338.56258 | 3470.90637 | 3405.97606 | 3373.72492 | 3479.22511 | 3450.42466 |
| MER63B | 3526.57578 | 3548.29885 | 3633.44306 | 3613.80333 | 3685.84892 | 3544.41086 |
| MER63C | 3077.78991 | 3154.08472 | 3166.72775 | 2776.78182 | 2758.22256 | 2687.9029 |
| MER63D | 5114.8268 | 5253.87287 | 5054.80711 | 5003.0673 | 5022.78252 | 4893.53167 |
| MER65-int | 238.954374 | 255.677868 | 227.57978 | 236.46769 | 234.194918 | 283.310436 |
| MER65D | 563.313482 | 571.028292 | 607.957423 | 562.310003 | 528.139697 | 562.539097 |
| MER66-int | 83.2542017 | 89.7976592 | 102.248397 | 95.531294 | 92.3734626 | 83.2021343 |
| MER66A | 17.7473913 | 18.9390264 | 14.0873725 | 17.7911027 | 24.5222383 | 17.2158076 |
| MER66B | 28.1099972 | 34.7762256 | 34.8100723 | 32.0131769 | 31.3905577 | 36.4988491 |
| MER66C | 81.7065737 | 77.0622872 | 74.0408103 | 78.2019689 | 74.2534246 | 74.9990653 |
| MER67A | 1167.80792 | 1148.26226 | 1135.27588 | 1155.83433 | 1221.23083 | 1245.38161 |
| MER67B | 1431.73719 | 1424.6743 | 1406.89187 | 1472.40844 | 1474.33216 | 1456.66869 |
| MER67C | 4012.91942 | 3912.95247 | 4064.41663 | 3619.45176 | 3406.20534 | 3573.70782 |
| MER67D | 2783.26353 | 2819.79558 | 2862.14146 | 2603.16189 | 2218.5048 | 2466.69413 |
| MER68 | 4929.66613 | 4922.76666 | 4993.47152 | 5072.28508 | 5225.85063 | 5051.73641 |
| MER68-int | 506.476953 | 540.824815 | 520.828753 | 481.446911 | 537.835681 | 483.165396 |
| MER68B | 1508.10518 | 1552.92038 | 1524.22715 | 1533.85473 | 1602.43581 | 1689.19102 |
| MER68C | 272.664562 | 230.696588 | 277.523836 | 282.253023 | 262.656321 | 268.691844 |
| MER70-int | 459.780742 | 422.61892 | 458.139512 | 444.018084 | 421.768897 | 416.649087 |
| MER70A | 1357.83534 | 1372.26401 | 1337.36312 | 1290.26464 | 1354.91143 | 1233.41952 |
| MER70B | 1730.23067 | 1715.37302 | 1719.11083 | 1753.04005 | 1623.76941 | 1685.54701 |
| MER72 | 41.7649822 | 32.0002355 | 26.5369804 | 37.3202605 | 35.6323599 | 36.5515426 |
| MER72B | 224.10564 | 236.493402 | 221.312841 | 200.059126 | 211.772916 | 222.531714 |
| MER73 | 952.595371 | 927.196183 | 898.808304 | 934.014917 | 880.94709 | 873.773411 |
| MER74A | 5227.23953 | 5279.83237 | 5175.33085 | 5611.64055 | 5608.66967 | 5524.81544 |
| MER74B | 2296.78411 | 2366.56462 | 2270.66237 | 2402.12945 | 2602.38006 | 2438.4475 |
| MER74C | 759.007673 | 718.378643 | 723.509316 | 731.127783 | 681.030556 | 641.280969 |
| MER76 | 1555.55633 | 1595.86105 | 1636.33931 | 1662.6333 | 1524.28332 | 1716.6903 |
| MER76-int | 123.405499 | 98.4504548 | 119.465675 | 123.125854 | 76.9200492 | 90.2266792 |
| MER77 | 4209.10275 | 4227.97806 | 4279.22335 | 4313.47263 | 4287.67688 | 4245.80579 |
| MER77B | 3703.72937 | 3609.36041 | 3796.24814 | 3731.64154 | 3755.47708 | 3665.06642 |

| | | | | | | |
|---|---|---|---|---|---|---|
| MER81 | 2063.04249 | 2130.40019 | 2159.73966 | 2237.17131 | 2322.45084 | 2313.45682 |
| MER83A-int | 7.25371933 | 4.4082112 | 0.90098695 | 1.5285007 | 1.01001106 | 2.01454047 |
| MER83B-int | 42.5650094 | 35.9187804 | 50.3381582 | 30.5311354 | 37.8541401 | 40.5279301 |
| MER87 | 230.362733 | 195.676035 | 194.571681 | 210.312503 | 217.87309 | 253.46423 |
| MER88 | 370.58453 | 351.515353 | 399.53539 | 331.855328 | 371.916168 | 400.414203 |
| MER89 | 4217.69753 | 4171.4045 | 4279.4111 | 4198.63078 | 3965.87719 | 4030.0615 |
| MER89-int | 685.885926 | 735.602117 | 705.700885 | 674.06881 | 607.30219 | 619.002463 |
| MER90 | 1566.52079 | 1587.04462 | 1566.46789 | 1469.11866 | 1469.70567 | 1513.71473 |
| MER90a | 3565.96794 | 3394.33284 | 3521.03532 | 3437.95227 | 3451.14961 | 3529.12347 |
| MER91A | 1857.81351 | 1816.67915 | 1804.7424 | 2021.82008 | 2122.41581 | 2067.89926 |
| MER91B | 1008.73777 | 991.932267 | 1020.70491 | 1049.10667 | 1116.49476 | 1071.49222 |
| MER91C | 489.326175 | 494.864131 | 438.695481 | 517.362888 | 518.444323 | 499.640448 |
| MER92-int | 1943.78288 | 1942.3965 | 1930.90692 | 1791.39145 | 1871.76113 | 1748.71876 |
| MER92A | 209.38168 | 206.370119 | 214.736069 | 193.817117 | 205.491038 | 191.480637 |
| MER92B | 1576.05873 | 1572.43149 | 1585.78768 | 1475.86084 | 1418.01524 | 1508.8628 |
| MER92C | 377.038222 | 375.351447 | 380.706062 | 380.647413 | 365.936316 | 371.17702 |
| MER92D | 380.39561 | 380.7398 | 410.485729 | 363.348898 | 345.65588 | 339.622308 |
| MER94 | 1993.41759 | 2096.27442 | 2108.7332 | 1994.05387 | 2078.31929 | 2079.2954 |
| MER94B | 393.040862 | 351.842505 | 360.975746 | 321.489594 | 350.485345 | 343.559175 |
| MER95 | 290.832413 | 299.351419 | 291.301375 | 281.092893 | 252.576875 | 256.446521 |
| MER96 | 538.902693 | 457.63836 | 497.520203 | 505.204571 | 511.131159 | 488.547302 |
| MER96B | 2000.72373 | 1866.96694 | 1951.06202 | 1886.89194 | 1867.0131 | 1926.76946 |
| MER97a | 405.567631 | 435.353078 | 397.055194 | 360.384815 | 397.56996 | 381.262895 |
| MER97b | 327.015555 | 330.943093 | 328.875763 | 292.579878 | 309.115609 | 300.167515 |
| MER97c | 497.347941 | 472.169074 | 476.689685 | 425.94725 | 412.517147 | 365.080705 |
| MER97d | 137.237685 | 132.900236 | 153.046342 | 140.156497 | 112.71406 | 115.579689 |
| MER99 | 1189.33368 | 1181.56615 | 1112.7512 | 1078.49711 | 1116.41455 | 1115.89774 |
| MERV1_I-int | 20823.0522 | 20550.9899 | 20744.1318 | 22275.3986 | 21329.0923 | 22852.103 |
| MERV1_LTR | 4790.90545 | 4757.70165 | 4923.25092 | 4959.64869 | 4735.198 | 4838.03606 |
| MERVK26-int | 13651.6151 | 13484.1978 | 13799.3318 | 14229.696 | 13365.5481 | 14313.595 |
| MERVL-int | 97071.7076 | 96655.3537 | 97208.162 | 99513.7123 | 98170.0548 | 100575.736 |
| MERVL_2A-int | 45250.1779 | 45051.3398 | 45746.1371 | 44379.383 | 44312.8549 | 44193.747 |
| MERX | 348.193206 | 359.842311 | 347.552339 | 290.15912 | 301.499808 | 292.998063 |
| MIR | 188145.816 | 189679.146 | 186801.675 | 191562.713 | 198439.906 | 194899.833 |
| MIR3 | 69931.7689 | 70346.8206 | 69333.4896 | 74762.6782 | 78293.0529 | 76601.2938 |
| MIRb | 208521.816 | 211307.15 | 208919.202 | 215899.092 | 223368.56 | 219140.432 |
| MIRc | 102934.516 | 103314.635 | 102923.036 | 108613.068 | 111937.906 | 110258.16 |
| MLT-int | 6205.35608 | 6150.12552 | 6085.92913 | 6081.5995 | 5919.74758 | 6009.05751 |
| MLT1-int | 1878.56546 | 1781.49477 | 1864.36029 | 1715.69607 | 1667.56025 | 1603.01976 |
| MLT1A | 29606.1372 | 29484.5129 | 29541.2539 | 29627.5814 | 30352.6935 | 30116.4668 |
| MLT1A-int | 850.847225 | 866.133811 | 855.661621 | 732.80373 | 690.948351 | 689.429292 |
| MLT1A0 | 64358.5187 | 64223.0463 | 64268.7796 | 63943.1156 | 63508.3282 | 63709.9471 |

| | | | | | | |
|---|---|---|---|---|---|---|
| MLT1A0-int | 1741.04415 | 1774.88407 | 1720.18316 | 1656.80184 | 1628.0726 | 1676.54086 |
| MLT1A1 | 46011.3618 | 46070.6191 | 46125.5945 | 45528.4549 | 46737.0585 | 45821.569 |
| MLT1A1-int | 1162.64077 | 1164.58842 | 1173.67844 | 1168.41885 | 1083.26894 | 1116.92477 |
| MLT1B | 57451.2861 | 57538.4622 | 57597.4209 | 58413.3546 | 58918.9102 | 57915.5648 |
| MLT1B-int | 1145.29339 | 1101.48314 | 1090.49639 | 1049.56796 | 997.982556 | 1067.53206 |
| MLT1C | 50839.3322 | 51015.1 | 50206.5521 | 51822.9939 | 52137.204 | 51902.2147 |
| MLT1C-int | 825.136785 | 864.992167 | 844.070865 | 799.390219 | 802.06779 | 756.972631 |
| MLT1D | 54533.4031 | 54832.212 | 54777.2249 | 55206.4697 | 55674.633 | 55211.6909 |
| MLT1D-int | 1397.77011 | 1448.51069 | 1316.87961 | 1374.84799 | 1327.37921 | 1276.61053 |
| MLT1E | 3294.6801 | 3473.02897 | 3318.77674 | 3376.28408 | 3294.76138 | 3350.33959 |
| MLT1E-int | 55.7869523 | 58.4497051 | 57.5132119 | 60.4184467 | 56.1145539 | 49.6724024 |
| MLT1E1 | 3677.10933 | 3653.35899 | 3840.87983 | 3707.75587 | 3816.13608 | 3705.27514 |
| MLT1E1-int | 82.4279612 | 64.3275223 | 81.2487059 | 74.4698452 | 54.417833 | 59.824145 |
| MLT1E1A | 7677.83384 | 7798.88489 | 7614.29996 | 7635.64812 | 7450.5352 | 7591.2732 |
| MLT1E1A-int | 215.356195 | 210.941451 | 202.811931 | 202.134742 | 189.190485 | 185.463363 |
| MLT1E2 | 12004.0257 | 12205.2366 | 12284.8891 | 12000.5725 | 11875.5079 | 11925.2258 |
| MLT1E2-int | 275.944359 | 232.167105 | 269.546321 | 261.303932 | 196.62275 | 253.372016 |
| MLT1E3 | 6902.31709 | 6768.50937 | 6643.06786 | 6610.90357 | 6619.33569 | 6608.25658 |
| MLT1E3-int | 85.6417007 | 79.0213573 | 97.4007872 | 91.5004879 | 69.3650199 | 67.8954802 |
| MLT1F | 11320.651 | 11607.5127 | 11262.0603 | 11839.0422 | 12213.2411 | 12230.4225 |
| MLT1F-int | 2943.64066 | 2851.55118 | 2875.09579 | 2728.44442 | 2704.32552 | 2626.14581 |
| MLT1F1 | 10527.0524 | 10657.0482 | 10601.1352 | 10844.9762 | 11133.8579 | 11031.6885 |
| MLT1F1-int | 628.086323 | 645.886777 | 650.533702 | 605.755637 | 636.208438 | 634.634912 |
| MLT1F2 | 16472.9344 | 16530.6901 | 16415.3898 | 16721.5874 | 16702.513 | 17055.8545 |
| MLT1F2-int | 599.936482 | 527.353831 | 596.762934 | 514.740156 | 524.059545 | 523.933494 |
| MLT1G | 6081.44467 | 6083.84419 | 6181.68999 | 6188.80508 | 6256.91924 | 6080.27425 |
| MLT1G-int | 277.308494 | 287.188033 | 252.920197 | 282.601954 | 271.280912 | 268.665497 |
| MLT1G1 | 7129.84145 | 7169.73597 | 7338.71163 | 7430.9371 | 7412.48114 | 7446.98277 |
| MLT1G1-int | 456.756786 | 480.496437 | 461.283674 | 466.239174 | 443.704726 | 482.674934 |
| MLT1G3 | 7484.60683 | 7433.01217 | 7376.09113 | 7790.80182 | 8052.96782 | 8202.45108 |
| MLT1G3-int | 547.272046 | 557.39449 | 540.075733 | 488.670306 | 533.796247 | 507.511135 |
| MLT1H | 15138.2931 | 15120.7045 | 15113.0173 | 15252.0615 | 15257.0902 | 15542.6799 |
| MLT1H-int | 2573.88971 | 2555.29967 | 2541.52301 | 2439.30508 | 2337.54206 | 2372.43837 |
| MLT1H1 | 6553.18125 | 6542.70884 | 6509.89529 | 6476.19402 | 6648.84843 | 6580.89562 |
| MLT1H1-int | 397.985722 | 408.495654 | 408.570973 | 348.963728 | 379.389761 | 393.310618 |
| MLT1H2 | 6018.4905 | 6142.6177 | 6086.69658 | 6039.15567 | 6054.71894 | 6086.55804 |
| MLT1H2-int | 474.550313 | 432.005911 | 483.644138 | 457.611495 | 464.934401 | 466.732911 |
| MLT1I | 10570.494 | 10619.828 | 10457.6778 | 10542.271 | 10589.4583 | 10622.4455 |
| MLT1I-int | 461.223517 | 471.518512 | 466.985172 | 434.85466 | 372.178087 | 406.589558 |
| MLT1J | 18184.7215 | 18425.077 | 18024.838 | 18711.1113 | 19520.0443 | 18644.6702 |
| MLT1J-int | 3542.86048 | 3544.6218 | 3547.10259 | 3383.86117 | 3370.0072 | 3355.10235 |
| MLT1J1 | 6251.72305 | 6203.35606 | 6084.87602 | 6323.6578 | 6369.89095 | 6394.85617 |

| | | | | | | |
|---|---|---|---|---|---|---|
| MLT1J1-int | 539.617789 | 478.782099 | 517.684591 | 450.217424 | 510.847965 | 477.467329 |
| MLT1J2 | 6649.94836 | 6582.87271 | 6568.57159 | 6721.58428 | 6713.37126 | 6564.84821 |
| MLT1J2-int | 235.20274 | 246.207748 | 216.800778 | 227.545119 | 246.556916 | 211.863163 |
| MLT1K | 17287.2934 | 17375.0289 | 17245.1006 | 17954.6764 | 18441.5033 | 18154.6621 |
| MLT1K-int | 64.3261674 | 69.8784907 | 66.5887651 | 66.9980174 | 50.499332 | 70.0022343 |
| MLT1L | 9422.77694 | 9328.22129 | 9346.26659 | 9447.42688 | 9465.33383 | 9480.09828 |
| MLT1L-int | 88.186479 | 82.9400033 | 73.2711908 | 59.0141625 | 49.9742728 | 72.9976982 |
| MLT1M | 1409.54351 | 1430.63393 | 1437.89598 | 1520.59509 | 1561.41065 | 1547.02356 |
| MLT1N2 | 3746.69439 | 3759.40403 | 3700.30385 | 3792.98782 | 3975.06512 | 3860.21741 |
| MLT1N2-int | 20.9349176 | 30.2045899 | 26.6026642 | 24.5413491 | 33.2489292 | 22.3443724 |
| MLT1O | 2998.48132 | 3117.92121 | 3115.91618 | 3348.64782 | 3424.78966 | 3382.64469 |
| MLT1O-int | 20.3708093 | 15.3471281 | 16.888859 | 15.4556827 | 15.3920313 | 13.1735533 |
| MLT2B1 | 11187.3128 | 11198.93 | 11273.9029 | 10950.5101 | 10843.9933 | 10972.3117 |
| MLT2B2 | 8670.12439 | 8835.80458 | 8784.13527 | 8304.4843 | 8338.23518 | 8290.56747 |
| MLT2B3 | 10283.2586 | 10239.3304 | 10288.7821 | 10448.3265 | 10262.3436 | 10387.5894 |
| MLT2B4 | 11953.2476 | 11921.0675 | 12173.361 | 11418.1365 | 11341.8533 | 11387.6475 |
| MLT2B5 | 2302.22125 | 2300.19611 | 2252.73403 | 2486.95696 | 2333.32092 | 2221.12644 |
| MLT2C1 | 5304.61778 | 5238.68411 | 5200.4578 | 5148.09661 | 5040.69836 | 4976.29653 |
| MLT2C2 | 3851.34958 | 3959.32222 | 3865.26007 | 3694.43364 | 3706.81546 | 3711.08164 |
| MLT2D | 9566.79441 | 9674.50494 | 9619.02212 | 9350.42394 | 9377.08124 | 9244.84799 |
| MLT2E | 802.136791 | 749.561554 | 734.081974 | 711.016909 | 738.055462 | 774.652554 |
| MLT2F | 3765.04364 | 3705.76786 | 3801.91183 | 3774.47513 | 3833.34637 | 3883.96376 |
| MLTR11A | 18587.8959 | 18642.6321 | 18668.9461 | 19682.0231 | 19745.6498 | 20091.1243 |
| MLTR11B | 23823.6898 | 23906.7957 | 23719.7118 | 25425.431 | 24858.3646 | 25342.2212 |
| MLTR12 | 3226.0125 | 3125.18763 | 3167.67303 | 3171.98459 | 2994.90118 | 3068.92209 |
| MLTR13 | 1601.96472 | 1583.3703 | 1564.42889 | 1507.64172 | 1601.12377 | 1518.51091 |
| MLTR14 | 47105.0786 | 47110.2388 | 47193.4299 | 49266.2214 | 49640.8338 | 49591.5157 |
| MLTR18_MM | 3791.91637 | 3930.01575 | 3774.11973 | 3855.50083 | 3845.74726 | 3716.78325 |
| MLTR18A_MM | 2817.55146 | 2752.28391 | 2879.64999 | 2742.55744 | 2662.73763 | 2755.45233 |
| MLTR18B_MM | 3989.13774 | 3964.87389 | 4020.85013 | 3989.11517 | 3936.05815 | 3890.97209 |
| MLTR18C_MM | 4943.0595 | 4876.39499 | 4849.8877 | 4889.63255 | 4842.94466 | 4758.38925 |
| MLTR18D_MM | 6621.48763 | 6648.42156 | 6552.7814 | 6812.86103 | 6676.83282 | 6647.37546 |
| MLTR25A | 19244.51 | 19262.4811 | 19393.6185 | 19116.3128 | 18216.9739 | 18842.2436 |
| MLTR25C | 6272.9741 | 6086.86187 | 6440.97628 | 5878.08749 | 5316.8634 | 5579.02714 |
| MLTR31A_MM | 3273.50612 | 3183.14676 | 3290.36561 | 3339.27484 | 3352.41405 | 3382.71665 |
| MLTR31C_MM | 2134.04674 | 2008.43766 | 2161.15682 | 2206.83029 | 2001.70188 | 2047.10227 |
| MLTR31D_MM | 2973.5059 | 2930.89635 | 3057.02018 | 3064.70699 | 2951.55214 | 3021.42536 |
| MLTR31E_MM | 1321.97357 | 1279.69158 | 1343.63936 | 1279.23185 | 1411.45016 | 1286.98622 |
| MLTR31F_MM | 3662.07553 | 3656.21507 | 3555.5777 | 3630.48174 | 4643.77195 | 4305.12885 |
| MLTR31FA_MM | 5228.61048 | 5131.74692 | 5287.94277 | 5229.37502 | 4894.12399 | 5205.7302 |
| MLTR32C_MM | 913.723278 | 949.564999 | 943.156515 | 891.135349 | 819.155321 | 864.151651 |
| MLTR73 | 5236.86869 | 5294.44085 | 5252.85137 | 5306.36942 | 5481.91877 | 5525.51363 |

| | | | | | | |
|---|---|---|---|---|---|---|
| MMAR1 | 2575.61402 | 2597.42584 | 2653.21687 | 2395.14972 | 2471.12666 | 2575.12108 |
| MMERGLN-int | 20170.3679 | 20221.2705 | 20236.3295 | 22870.0428 | 25133.8289 | 24986.1927 |
| MMERGLN_LTR | 3657.81326 | 3646.50163 | 3693.3872 | 3688.97959 | 3625.2902 | 3732.03976 |
| MMERVK10C-int | 54322.1907 | 53565.6738 | 54369.6042 | 53139.0126 | 50360.503 | 51882.8065 |
| MMERVK10D3_I-int | 10120.3555 | 10175.7412 | 10435.4508 | 10730.4984 | 10401.1733 | 10767.1073 |
| MMERVK10D3_LTR | 1321.69833 | 1305.08081 | 1372.35599 | 1355.38094 | 1348.78998 | 1414.5574 |
| MMERVK9C_I-int | 25141.7896 | 25374.1511 | 25255.3482 | 22815.9842 | 21637.4331 | 22117.5532 |
| MMERVK9E_I-int | 27984.8984 | 27734.6022 | 28223.6784 | 26039.925 | 25309.6432 | 25760.9628 |
| MMETn-int | 23941.4705 | 23750.6184 | 23620.8873 | 21412.7674 | 21555.6666 | 20751.1862 |
| MMSAT4 | 11558.3121 | 11834.7099 | 11780.0897 | 11180.6223 | 10675.6411 | 10833.7695 |
| MMTV-int | 675.727258 | 724.989948 | 662.678441 | 669.786269 | 624.895948 | 639.63833 |
| MMVL30-int | 12418.4382 | 12287.1143 | 12674.1118 | 12769.5544 | 12578.8566 | 13145.3983 |
| MRLTR33 | 3569.31065 | 3593.27413 | 3611.48599 | 3672.28651 | 3643.16959 | 3469.10881 |
| MT-int | 3143.1473 | 3158.9824 | 3131.60071 | 3223.44586 | 3247.02955 | 3191.08806 |
| MT2_Mm | 18166.3916 | 17832.2643 | 18143.6304 | 17887.182 | 17226.7293 | 17615.1996 |
| MT2A | 64356.1511 | 64535.3733 | 64361.1748 | 65183.2759 | 64103.5157 | 64507.706 |
| MT2B | 39637.6996 | 39797.7979 | 38903.4628 | 41590.298 | 43459.5134 | 43646.1423 |
| MT2B1 | 44299.4578 | 43818.4116 | 43972.1423 | 42768.9288 | 42384.7257 | 42216.5751 |
| MT2B2 | 16731.2263 | 17074.6077 | 16672.6479 | 17617.4704 | 17833.2895 | 18104.667 |
| MT2C_Mm | 16438.4184 | 16422.8535 | 16597.4861 | 16927.5898 | 16564.0315 | 17009.0489 |
| MTA_Mm | 124371.208 | 124606.453 | 125384.198 | 134473.682 | 136191.536 | 138739.955 |
| MTA_Mm-int | 57279.3159 | 57548.6046 | 58357.7467 | 59159.0001 | 63563.7316 | 62811.5579 |
| MTB | 53918.6373 | 54439.4071 | 53841.3314 | 58213.2707 | 57707.7102 | 58732.85 |
| MTB-int | 8999.19297 | 8993.76923 | 9133.27611 | 9117.26645 | 9035.43741 | 9274.97034 |
| MTB_Mm | 39805.8044 | 40028.4193 | 39550.9004 | 42586.5074 | 42390.5499 | 43246.9334 |
| MTB_Mm-int | 5436.08436 | 5595.17916 | 5519.5612 | 5455.52392 | 5494.95048 | 5316.04045 |
| MTC | 187096.292 | 188588.292 | 186717.917 | 199072.522 | 204733.982 | 204852.515 |
| MTC-int | 34410.2079 | 34454.6245 | 33954.0055 | 34576.9752 | 33592.5703 | 34649.2229 |
| MTD | 350787.533 | 351881.91 | 351635.385 | 364417.305 | 370245.905 | 369609.394 |
| MTD-int | 25106.2838 | 25012.6071 | 25155.7863 | 24893.5467 | 24488.8094 | 24735.2096 |
| MTE-int | 42436.7771 | 42255.0503 | 42396.3822 | 41516.6908 | 40441.2155 | 40804.1234 |
| MTE2a | 105831.199 | 105563.066 | 105795.116 | 105760.043 | 106611.772 | 106511.144 |
| MTE2a-int | 4264.07289 | 4309.69522 | 4471.88244 | 4275.84941 | 4054.63174 | 4263.87078 |
| MTE2b | 93475.9064 | 93973.5083 | 93715.4297 | 95834.1147 | 95143.0639 | 95971.2067 |
| MTE2b-int | 7387.75689 | 7439.9485 | 7308.39503 | 7026.1205 | 6821.42117 | 6889.29305 |
| MTEa | 128148.286 | 128667.901 | 128386.741 | 132360.499 | 133172.732 | 133549.179 |
| MTEa-int | 15017.5382 | 14856.3743 | 14814.8229 | 14606.5828 | 14422.2432 | 14453.412 |
| MTEb | 65057.68 | 65590.4131 | 65219.5958 | 67413.4462 | 67544.2747 | 67322.0026 |
| MTEb-int | 6799.69761 | 6853.31874 | 6948.55793 | 6805.06962 | 6814.79899 | 6672.90326 |
| MuLV-int | 4222.29323 | 4358.6748 | 4269.54455 | 4540.10752 | 4294.8229 | 4475.98729 |
| MurERV4-int | 14237.0667 | 14132.7715 | 13755.1041 | 12749.25 | 12785.6287 | 12646.4008 |
| MurERV4_19-int | 11765.0258 | 11746.4507 | 11847.7679 | 10419.4058 | 10011.8178 | 9946.85548 |

| | | | | | | |
|---|---|---|---|---|---|---|
| MuRRS-int | 9661.63694 | 9817.28628 | 9943.23091 | 10634.8268 | 10390.0425 | 10646.8307 |
| MuRRS4-int | 13177.5964 | 13226.8043 | 13394.3347 | 14369.4704 | 14042.4627 | 14364.1136 |
| MurSAT1 | 907.912334 | 964.505083 | 1051.24924 | 1053.77751 | 966.369576 | 1065.90259 |
| MurSatRep1 | 13843.3574 | 13887.219 | 13560.0206 | 13283.5108 | 12697.3846 | 12650.9269 |
| MURVY-int | 5625.2111 | 5468.49178 | 5837.93801 | 6702.96022 | 5582.5297 | 6954.4991 |
| MURVY-LTR | 6127.56314 | 6184.33745 | 6328.68006 | 7103.56776 | 6905.08494 | 7190.01944 |
| MusHAL1 | 42460.7952 | 42158.2903 | 42196.1528 | 36093.3596 | 34467.1984 | 34807.0406 |
| MYSERV-int | 13637.4212 | 13552.4375 | 13628.5762 | 12257.7887 | 11917.5559 | 11917.4113 |
| MYSERV16_I-int | 27892.921 | 27849.3709 | 27679.3955 | 24695.1168 | 23783.8966 | 23477.6105 |
| MYSERV6-int | 35803.1469 | 35689.6438 | 35464.8499 | 31402.3995 | 30696.519 | 30544.1928 |
| nhAT5a_ML | 17.7211781 | 11.4287856 | 10.5491084 | 13.4151549 | 12.6852994 | 22.331199 |
| ORR1A0 | 15781.16 | 15877.9514 | 15956.5247 | 16869.0854 | 16547.6811 | 16990.114 |
| ORR1A0-int | 10705.9681 | 10718.5236 | 10724 | 11051.5457 | 10734.0102 | 10923.4064 |
| ORR1A1 | 28344.8293 | 28234.0428 | 28536.8876 | 30365.9264 | 29982.4004 | 30695.8134 |
| ORR1A1-int | 17715.7556 | 17810.14 | 17981.6885 | 18160.8634 | 17350.8027 | 18099.6722 |
| ORR1A2 | 105435.237 | 105494.584 | 105105.399 | 112460.439 | 112949.498 | 113681.691 |
| ORR1A2-int | 36542.7107 | 36557.103 | 36757.5871 | 37006.5134 | 36201.4407 | 36835.9401 |
| ORR1A3 | 30470.751 | 30710.3222 | 30259.6253 | 32109.3228 | 31929.8443 | 31772.5162 |
| ORR1A3-int | 15966.7023 | 15909.626 | 15968.1235 | 15447.2307 | 15118.2337 | 15472.2926 |
| ORR1A4 | 55877.7842 | 55821.7048 | 55362.4801 | 58579.5328 | 59369.8485 | 59747.4949 |
| ORR1A4-int | 22425.7131 | 22426.7731 | 22438.2219 | 22247.4863 | 22016.8332 | 22280.2207 |
| ORR1B1 | 112718.902 | 113074.36 | 112636.714 | 116904.42 | 117675.99 | 118033.283 |
| ORR1B1-int | 29716.7129 | 29689.2527 | 30068.2777 | 29160.8301 | 27977.9214 | 28784.4483 |
| ORR1B2 | 56388.6566 | 56295.015 | 56668.4268 | 58801.8603 | 59259.7944 | 59401.9632 |
| ORR1B2-int | 10207.6046 | 10193.702 | 10191.5005 | 9850.99752 | 9799.24755 | 9604.69779 |
| ORR1C1 | 51347.7657 | 50850.2926 | 51461.5458 | 52423.4681 | 52449.6128 | 52708.082 |
| ORR1C1-int | 9726.20899 | 9918.26719 | 9806.18453 | 9386.22804 | 9558.06521 | 9381.19881 |
| ORR1C2 | 65210.7306 | 65196.1259 | 64348.6466 | 67350.5093 | 68172.7957 | 68241.3464 |
| ORR1C2-int | 10912.677 | 10906.0291 | 10834.6135 | 10415.1252 | 10280.4412 | 10380.5648 |
| ORR1D-int | 3912.36345 | 3916.95282 | 4013.2503 | 3851.42259 | 3614.54165 | 3764.03628 |
| ORR1D1 | 127115.785 | 127151.628 | 127168.097 | 131094.208 | 133132.254 | 133062.156 |
| ORR1D1-int | 17565.5199 | 17887.1921 | 17675.7272 | 17533.5475 | 17493.78 | 17374.6049 |
| ORR1D2 | 97059.2542 | 96859.6042 | 96604.3011 | 99152.1265 | 99349.5522 | 98708.2359 |
| ORR1D2-int | 12145.0345 | 12088.2542 | 11951.1614 | 11653.9532 | 11526.54 | 11518.4979 |
| ORR1E | 154592.552 | 155675.063 | 154884.47 | 160227.011 | 160129.755 | 160002.495 |
| ORR1E-int | 15785.4128 | 15940.4789 | 15879.6117 | 15422.6363 | 15117.9025 | 15081.6851 |
| ORR1F | 148007.909 | 147958.566 | 147841.637 | 151063.486 | 152781.479 | 151393.11 |
| ORR1F-int | 3748.30231 | 3721.44183 | 3650.03417 | 3511.8124 | 3546.41334 | 3496.32486 |
| ORR1G | 28474.8243 | 28630.7041 | 28576.3534 | 29021.5887 | 29066.4755 | 29159.3114 |
| ORR1G-int | 604.783828 | 608.662018 | 682.204577 | 608.161234 | 617.037671 | 572.822573 |
| ORSL | 1258.20522 | 1208.18137 | 1237.36007 | 1174.33895 | 1206.36447 | 1217.96491 |
| ORSL-2a | 463.617307 | 416.658884 | 403.49347 | 407.954661 | 419.022058 | 444.852654 |

| | | | | | | |
|---|---|---|---|---|---|---|
| ORSL-2b | 1367.17563 | 1274.1396 | 1317.41934 | 1246.90764 | 1253.20661 | 1251.74798 |
| PB1 | 44143.5941 | 44191.4051 | 43355.0281 | 46973.5239 | 51569.6417 | 49635.0777 |
| PB1D10 | 234015.847 | 235614.15 | 231212.53 | 246022.33 | 264673.889 | 256645.135 |
| PB1D11 | 60969.1083 | 61218.8279 | 60548.586 | 63989.6629 | 69530.1987 | 67859.8398 |
| PB1D7 | 89411.5367 | 89966.3523 | 88164.7739 | 95441.6839 | 102901.791 | 100611.893 |
| PB1D9 | 99630.8246 | 100103.306 | 97611.9981 | 106019.886 | 115716.602 | 112076.925 |
| Plat_L3 | 3268.85904 | 3234.00542 | 3066.66893 | 3169.2462 | 3132.90013 | 3130.12187 |
| polypurine | 2995.7672 | 2925.75607 | 2985.67087 | 3099.50649 | 3277.53469 | 3176.04488 |
| polypyrimidine | 2912.66504 | 2765.09775 | 2883.7915 | 2914.89925 | 2925.51305 | 3025.11193 |
| RCHARR1 | 50039.368 | 50202.432 | 49633.2998 | 46613.8005 | 49142.1213 | 47166.8188 |
| Ricksha | 992.701581 | 1015.19779 | 1019.46125 | 998.79794 | 1051.75623 | 999.047314 |
| Ricksha_0 | 682.647021 | 652.416471 | 676.822204 | 591.176562 | 578.517485 | 586.578802 |
| Ricksha_a | 256.865336 | 286.698063 | 258.865845 | 254.196685 | 267.645605 | 229.158011 |
| Ricksha_b | 47.0642447 | 50.6130201 | 57.4146863 | 50.3087262 | 44.4392656 | 43.5102206 |
| Ricksha_c | 2105.65364 | 2053.74565 | 2127.66755 | 2021.25634 | 2172.24849 | 2114.675 |
| RLTR1 | 318.135568 | 362.127825 | 386.196252 | 358.903261 | 336.1416 | 339.661828 |
| RLTR10 | 15116.7458 | 14866.8997 | 15426.9602 | 17086.171 | 17631.368 | 18006.3655 |
| RLTR10-int | 18639.388 | 18653.4902 | 19013.0646 | 21298.0263 | 22273.7162 | 22578.2623 |
| RLTR10A | 6170.13024 | 6221.31879 | 6191.19312 | 7038.93597 | 7021.30605 | 7192.32026 |
| RLTR10B | 1720.45734 | 1777.24766 | 1721.70814 | 1828.57282 | 1799.9288 | 1957.09904 |
| RLTR10B2 | 5870.86399 | 5863.59513 | 5895.21547 | 6239.26981 | 5897.40152 | 6069.60825 |
| RLTR10C | 15000.7969 | 14966.5753 | 15055.2628 | 16112.7848 | 15723.494 | 16289.2886 |
| RLTR10D | 3706.45816 | 3725.93257 | 3723.04416 | 4104.5199 | 3832.77693 | 3986.41954 |
| RLTR10D2 | 1739.75918 | 1806.39262 | 1710.70357 | 1952.54069 | 1785.50484 | 1934.439 |
| RLTR10E | 1710.95453 | 1688.02462 | 1780.72558 | 1824.71697 | 1776.23243 | 1817.24302 |
| RLTR10F | 3275.83752 | 3305.51967 | 3204.2194 | 3572.01469 | 3398.06204 | 3527.54719 |
| RLTR11A | 23982.5691 | 24095.9366 | 23947.8505 | 24629.7045 | 24905.874 | 24884.0303 |
| RLTR11A2 | 24457.5163 | 24516.6818 | 24664.9501 | 25301.4407 | 25822.0633 | 25824.7522 |
| RLTR11B | 8287.03249 | 8241.09553 | 8170.09153 | 8490.14921 | 8646.49388 | 8653.55459 |
| RLTR11C_MM | 7807.38319 | 7665.9886 | 7726.29284 | 8122.53391 | 8395.59428 | 8202.09944 |
| RLTR11D | 4375.67983 | 4470.18829 | 4374.25176 | 4657.17671 | 4776.18611 | 4747.16742 |
| RLTR12A | 6944.80346 | 7040.10433 | 6930.14282 | 7420.51575 | 7064.02986 | 7066.67947 |
| RLTR12B | 7690.96665 | 7582.39359 | 7494.64716 | 7479.6628 | 7526.64822 | 7647.51262 |
| RLTR12B2 | 879.088812 | 839.193866 | 835.135972 | 804.604872 | 860.384681 | 842.245046 |
| RLTR12BD_Mm | 7120.61493 | 7075.0378 | 7001.78184 | 7033.50748 | 6951.2817 | 7087.37363 |
| RLTR12C | 2481.65539 | 2447.05811 | 2442.40668 | 2722.77703 | 2685.01743 | 2844.86885 |
| RLTR12D | 5556.11884 | 5574.36459 | 5667.03461 | 5622.89047 | 5691.09072 | 5770.78442 |
| RLTR12E | 5654.42939 | 5866.93804 | 5710.22839 | 5970.71497 | 6259.72502 | 6367.85393 |
| RLTR12F | 499.420881 | 463.10883 | 448.341438 | 431.141976 | 413.688809 | 413.469196 |
| RLTR12G | 2355.29776 | 2362.81 | 2399.93109 | 2455.62648 | 2448.74233 | 2566.61453 |
| RLTR12H | 1608.09022 | 1557.98036 | 1648.00288 | 1673.30909 | 1635.44226 | 1727.97092 |
| RLTR13A | 2248.78248 | 2298.64499 | 2294.13946 | 1980.60791 | 2023.72219 | 2082.01776 |

| | | | | | | |
|---|---|---|---|---|---|---|
| RLTR13A1 | 2410.37066 | 2545.91348 | 2398.3683 | 2256.40509 | 2209.75866 | 2182.88664 |
| RLTR13A2 | 2575.99306 | 2577.83272 | 2651.07719 | 2307.1751 | 3430.88495 | 3017.86039 |
| RLTR13A3 | 3514.94917 | 3513.76239 | 3552.80969 | 3112.80354 | 3146.53833 | 3124.67714 |
| RLTR13B1 | 8927.07466 | 9105.11018 | 9163.45917 | 8818.83518 | 8593.73144 | 8556.86983 |
| RLTR13B2 | 4922.87166 | 4914.27628 | 5010.17045 | 4881.47471 | 4480.88369 | 4662.76396 |
| RLTR13B3 | 4381.84885 | 4471.32973 | 4455.45616 | 4326.5825 | 4273.84119 | 4334.81393 |
| RLTR13B4 | 4092.37585 | 4140.22382 | 4054.71988 | 3992.77907 | 3683.381 | 3839.83488 |
| RLTR13C1 | 5015.71361 | 5078.60264 | 4907.92485 | 4611.36057 | 4352.35769 | 4572.08987 |
| RLTR13C2 | 11195.3094 | 11020.1567 | 11083.7491 | 10937.7222 | 10042.5927 | 10544.3582 |
| RLTR13C3 | 3462.0289 | 3651.80635 | 3656.38818 | 3419.32295 | 3284.35924 | 3319.97963 |
| RLTR13D | 519.94163 | 517.313546 | 542.001946 | 499.451359 | 484.407805 | 470.297877 |
| RLTR13D1 | 3351.11661 | 3257.02895 | 3265.15873 | 3220.12906 | 3029.70646 | 3273.46381 |
| RLTR13D2 | 4698.39117 | 4641.69986 | 4685.66393 | 4353.88168 | 4218.75365 | 4250.29544 |
| RLTR13D3 | 4995.02195 | 5037.62316 | 4944.05573 | 4639.03191 | 4570.74189 | 4589.28946 |
| RLTR13D3A | 2109.25481 | 2122.72491 | 2025.17067 | 1996.26935 | 2039.2151 | 2044.0374 |
| RLTR13D3A1 | 6328.27663 | 6217.72234 | 6133.12656 | 5823.10793 | 5941.34188 | 5862.45359 |
| RLTR13D4 | 2717.52604 | 2625.83347 | 2687.2087 | 2525.78579 | 2362.00414 | 2544.48447 |
| RLTR13D5 | 9018.54513 | 8745.11578 | 8915.8535 | 8611.75512 | 8254.11923 | 8416.53652 |
| RLTR13D6 | 12174.226 | 12048.2564 | 12193.6967 | 11462.6337 | 11111.5538 | 11338.4818 |
| RLTR13E | 7454.32533 | 7561.66306 | 7456.93211 | 7175.50632 | 7498.12422 | 7222.48616 |
| RLTR13F | 1289.54783 | 1309.15964 | 1318.59732 | 1248.15739 | 1293.46363 | 1338.12647 |
| RLTR13G | 6144.64419 | 6130.04829 | 6084.10641 | 5871.33394 | 5750.91897 | 5742.11979 |
| RLTR14 | 16334.3672 | 16021.5378 | 15999.7215 | 16103.2571 | 16542.1322 | 16081.8116 |
| RLTR14-int | 13620.9509 | 13817.2578 | 13678.6268 | 14134.6141 | 14140.0363 | 14320.1889 |
| RLTR14_RN | 7550.80829 | 7512.76114 | 7453.82296 | 7559.40611 | 7328.93463 | 7435.4554 |
| RLTR15 | 17982.1196 | 17701.64 | 18028.713 | 17549.5564 | 17640.3653 | 17749.168 |
| RLTR16 | 11357.9226 | 11368.4945 | 11062.733 | 12495.996 | 12375.5107 | 12475.9295 |
| RLTR16B_MM | 5734.16263 | 5815.84153 | 5753.23658 | 5775.33427 | 5903.45853 | 6050.63479 |
| RLTR16C_MM | 3381.1407 | 3431.39792 | 3369.51397 | 3439.64105 | 3423.3907 | 3605.83104 |
| RLTR17 | 19686.6183 | 19476.2833 | 19547.1603 | 20479.5534 | 20072.5397 | 20353.6149 |
| RLTR17B_Mm | 77138.0879 | 77742.1691 | 77639.6827 | 74124.594 | 76172.1668 | 74970.7764 |
| RLTR17C_Mm | 1920.23083 | 1854.88112 | 1822.93348 | 1749.98256 | 1810.77701 | 1797.14726 |
| RLTR17D_Mm | 11827.6418 | 11924.1668 | 11837.1816 | 10717.6341 | 10017.4525 | 10203.2422 |
| RLTR18 | 5937.79942 | 6076.09012 | 6106.79809 | 6066.54067 | 6150.74959 | 6156.39256 |
| RLTR18-int | 3548.09421 | 3463.8861 | 3517.63556 | 3200.76153 | 3200.95739 | 3050.37272 |
| RLTR18B | 8242.49992 | 8146.97673 | 8114.90793 | 8355.28085 | 8532.77225 | 8438.82819 |
| RLTR19 | 7110.27015 | 7168.18577 | 7123.79836 | 6763.28428 | 6929.87276 | 6990.49382 |
| RLTR19-int | 17889.2001 | 17762.4545 | 17542.8837 | 17035.075 | 16968.9924 | 16621.2463 |
| RLTR19A | 1101.84971 | 1056.58684 | 1056.4674 | 1082.57817 | 1158.10514 | 1090.63999 |
| RLTR19A2 | 557.666633 | 539.355108 | 502.35852 | 553.092539 | 567.325928 | 543.493176 |
| RLTR19B | 2094.20318 | 2203.62403 | 2135.88209 | 2040.89407 | 2043.55718 | 1946.4244 |
| RLTR19C | 1899.06629 | 2025.98768 | 1991.61572 | 2023.30115 | 1840.20892 | 1878.65407 |

| | | | | | | |
|---|---|---|---|---|---|---|
| RLTR19D | 408.026954 | 416.496067 | 453.831628 | 414.557462 | 421.647354 | 416.754474 |
| RLTR1A2_MM | 3752.282 | 3729.68546 | 3738.23021 | 3874.82044 | 3843.10374 | 3851.96519 |
| RLTR1B | 8478.03557 | 8499.79685 | 8320.99756 | 8834.8167 | 8596.11487 | 8570.92599 |
| RLTR1B-int | 9774.16186 | 9671.73654 | 9911.72424 | 10498.9775 | 10180.6968 | 10247.8752 |
| RLTR1C | 6153.73965 | 6329.07199 | 6317.68606 | 6349.3479 | 5798.94855 | 6036.30552 |
| RLTR1D | 3115.38699 | 3118.41391 | 3111.95034 | 3317.63926 | 3324.85873 | 3395.55124 |
| RLTR1D2_MM | 1855.04487 | 1834.23221 | 1852.2834 | 1986.34596 | 1995.38172 | 2036.28222 |
| RLTR1E_MM | 499.479075 | 498.130496 | 470.744037 | 485.733731 | 454.552318 | 475.094061 |
| RLTR1F_Mm | 548.825442 | 507.027821 | 520.39468 | 506.352842 | 500.567982 | 567.210133 |
| RLTR20A | 2121.71028 | 2151.78654 | 2156.69618 | 2206.71035 | 2287.38364 | 2364.23834 |
| RLTR20A1 | 627.443575 | 646.620467 | 617.973933 | 664.548874 | 677.717329 | 704.8839 |
| RLTR20A2 | 2643.3306 | 2555.1409 | 2548.86443 | 2701.18033 | 2774.66471 | 2792.62607 |
| RLTR20A2B_MM | 3508.72983 | 3455.07089 | 3489.89056 | 3722.4753 | 3693.74609 | 3834.99613 |
| RLTR20A3_MM | 6392.31446 | 6555.77004 | 6477.46842 | 6821.40716 | 6779.66858 | 6790.95148 |
| RLTR20A4 | 28469.0459 | 28139.5618 | 28109.1465 | 27335.2394 | 27770.4175 | 27962.3264 |
| RLTR20B1 | 4647.71475 | 4738.27367 | 4666.42191 | 4949.621 | 4927.50118 | 4986.20358 |
| RLTR20B2 | 6584.6953 | 6683.68745 | 6524.34112 | 7109.5518 | 7016.51792 | 7262.75063 |
| RLTR20B3 | 7429.45005 | 7312.10673 | 7534.56261 | 7735.72983 | 7569.79123 | 7908.46856 |
| RLTR20B3A_MM | 4998.52456 | 4951.90717 | 4972.70948 | 5238.0027 | 4968.54212 | 5270.96259 |
| RLTR20B4_MM | 3071.84003 | 3194.73887 | 3050.93667 | 3272.09906 | 3347.11603 | 3595.09358 |
| RLTR20B5_MM | 6911.34963 | 7043.6132 | 7016.93874 | 7253.71208 | 7116.10855 | 7101.82352 |
| RLTR20C | 1591.89675 | 1585.32765 | 1505.62555 | 1578.98866 | 1584.09457 | 1601.0184 |
| RLTR20C1_MM | 14293.2594 | 14112.1142 | 14026.9237 | 14219.7537 | 14189.4162 | 14389.4828 |
| RLTR20C2_MM | 12863.9511 | 12710.7902 | 12816.1027 | 13274.4518 | 13782.9804 | 13849.0018 |
| RLTR20D | 5241.89953 | 5300.40291 | 5284.29452 | 5771.73822 | 5463.45477 | 5652.09071 |
| RLTR21 | 19212.6353 | 19153.574 | 19309.1012 | 18755.3566 | 21718.9711 | 20516.5657 |
| RLTR22_Mur | 21358.9959 | 20968.3655 | 21452.3718 | 18671.9855 | 17080.6424 | 18158.79 |
| RLTR22_Mus | 7056.63582 | 7032.5872 | 6932.30299 | 6506.99633 | 6176.32622 | 6435.8781 |
| RLTR23 | 12477.2869 | 12571.9424 | 12592.2462 | 13314.2445 | 13062.0939 | 13260.3862 |
| RLTR24 | 10421.6827 | 10326.6783 | 10337.2126 | 9946.28038 | 10118.992 | 10212.9294 |
| RLTR24B_MM | 3047.42243 | 3302.49571 | 3151.62877 | 3109.23879 | 3124.82554 | 2941.19234 |
| RLTR25A | 11827.6706 | 11998.8623 | 12112.4994 | 11592.7592 | 11141.955 | 11588.8447 |
| RLTR25B | 19169.1748 | 18667.6993 | 18990.8169 | 18500.8054 | 18314.3817 | 18614.7151 |
| RLTR26 | 3213.32059 | 3261.02738 | 3252.84761 | 3721.16063 | 3431.75215 | 3840.23009 |
| RLTR26_Mus | 11068.7652 | 11109.7158 | 11294.2216 | 13184.7212 | 12133.4739 | 13398.707 |
| RLTR26B_MM | 4749.00941 | 4832.72589 | 4797.91891 | 5307.31107 | 5159.42871 | 5473.01652 |
| RLTR26C_MM | 4398.44181 | 4279.73708 | 4360.65485 | 4965.65179 | 4750.14092 | 4893.71256 |
| RLTR26D_MM | 2384.88566 | 2423.30292 | 2405.99385 | 2655.01098 | 2631.3653 | 2656.96685 |
| RLTR27 | 5089.53105 | 5055.34146 | 5030.40269 | 5910.4051 | 5558.53313 | 5991.03508 |
| RLTR28 | 19119.2313 | 19244.9298 | 18918.1319 | 20257.0708 | 20748.3786 | 20658.4226 |
| RLTR28B | 3248.16528 | 3101.51435 | 3037.03922 | 3150.24472 | 3094.78712 | 3182.20047 |
| RLTR3_Mm | 152.683032 | 157.063483 | 173.499178 | 146.685818 | 155.093196 | 149.580122 |

| | | | | | | |
|---|---|---|---|---|---|---|
| RLTR30 | 2716.59023 | 2643.71075 | 2689.44691 | 2804.83437 | 3098.54581 | 2875.42996 |
| RLTR30B_MM | 2339.73498 | 2446.31977 | 2420.75231 | 2469.26733 | 2401.75493 | 2514.00289 |
| RLTR30C_MM | 973.92401 | 989.564939 | 1010.14868 | 962.843086 | 871.613904 | 1008.56369 |
| RLTR30D_MM | 2317.87001 | 2288.68885 | 2278.02735 | 2353.70573 | 2301.62773 | 2336.22933 |
| RLTR30D_RN | 457.393243 | 441.149689 | 440.875136 | 437.228959 | 443.907705 | 489.580919 |
| RLTR30D2_MM | 6244.75873 | 6095.02692 | 6156.47877 | 6552.51049 | 6250.47511 | 6487.16326 |
| RLTR30E_MM | 2741.83355 | 2747.38826 | 2723.60014 | 2815.39828 | 2622.67993 | 2824.46047 |
| RLTR31_Mm | 1296.18501 | 1323.44755 | 1298.2052 | 1318.7704 | 1356.12381 | 1319.95608 |
| RLTR31_Mur | 3166.61756 | 3090.33182 | 3142.07485 | 3214.4157 | 3096.8047 | 3147.13703 |
| RLTR31A_Mm | 1648.78623 | 1592.7577 | 1659.42942 | 1647.96082 | 1706.68632 | 1679.01293 |
| RLTR31B_Mm | 2965.68912 | 3001.34835 | 2956.83216 | 3101.83812 | 3044.04471 | 3165.72847 |
| RLTR31B2 | 9167.34966 | 8993.27086 | 9023.23642 | 8942.90024 | 8907.23411 | 8776.16286 |
| RLTR31C_MM | 4904.98796 | 4809.2938 | 4810.28857 | 4791.36237 | 4729.64294 | 4824.98765 |
| RLTR31D_MM | 23886.2927 | 23709.4744 | 23962.9752 | 23296.9107 | 23506.5956 | 23877.7837 |
| RLTR31M | 2325.48967 | 2341.83141 | 2340.14402 | 2299.44775 | 2316.5117 | 2386.92522 |
| RLTR33 | 16442.8584 | 16587.3455 | 16350.216 | 17376.6367 | 21424.7425 | 19164.0401 |
| RLTR34B_MM | 4853.587 | 4952.80631 | 4819.61108 | 5332.53512 | 5354.79776 | 5440.31317 |
| RLTR34C_MM | 3606.38085 | 3550.17195 | 3670.94247 | 3754.78386 | 3625.30842 | 3808.54669 |
| RLTR34D_MM | 3929.87544 | 4025.366 | 4035.54291 | 4253.89386 | 4168.33148 | 4368.79106 |
| RLTR35B_MM | 3243.84849 | 3365.92349 | 3352.64866 | 3409.67501 | 3357.1572 | 3504.56745 |
| RLTR4_Mm | 1220.6181 | 1254.79211 | 1238.82433 | 1427.28735 | 1404.70278 | 1423.12578 |
| RLTR4_MM-int | 11648.2496 | 11606.3751 | 11599.8424 | 14185.1676 | 13670.5209 | 14398.6698 |
| RLTR40 | 20888.3357 | 20780.3759 | 21090.8086 | 19218.3752 | 18443.5714 | 18715.1441 |
| RLTR41 | 8529.73693 | 8638.73853 | 8688.3893 | 9456.90035 | 9324.43058 | 9624.55947 |
| RLTR41A2 | 3516.48265 | 3637.27908 | 3693.20377 | 3977.63772 | 3917.19063 | 3943.98197 |
| RLTR41B | 701.646355 | 734.541578 | 739.407956 | 696.368146 | 691.190217 | 721.197825 |
| RLTR41C | 1119.93106 | 1101.89241 | 1131.46559 | 1173.33812 | 1097.88855 | 1150.48744 |
| RLTR42 | 1316.83945 | 1251.03922 | 1230.92892 | 1340.34058 | 1306.77547 | 1338.54448 |
| RLTR42-int | 17085.3306 | 17258.6965 | 17030.1124 | 16482.0133 | 16615.3786 | 16560.6257 |
| RLTR43A | 2.59720478 | 7.02037208 | 2.60443524 | 6.83558424 | 7.59513665 | 0 |
| RLTR43B | 5.22062278 | 4.24498908 | 8.81281828 | 3.86392068 | 5.73676514 | 3.04815745 |
| RLTR43C | 1763.79669 | 1713.25174 | 1761.69424 | 1834.38814 | 1852.46699 | 1862.12583 |
| RLTR44-int | 6110.97018 | 6248.4173 | 6337.43649 | 6354.77444 | 6168.82464 | 6434.57138 |
| RLTR44A | 1622.83358 | 1624.84036 | 1654.94804 | 1788.24581 | 1686.24301 | 1745.25918 |
| RLTR44B | 1457.90636 | 1391.69564 | 1461.94131 | 1634.11188 | 1598.59813 | 1645.89462 |
| RLTR44C | 1918.84101 | 1938.31403 | 1929.48976 | 2013.87841 | 1886.66516 | 1995.91896 |
| RLTR44D | 813.358142 | 818.787864 | 814.375466 | 863.766481 | 876.118236 | 873.875751 |
| RLTR44E | 985.55953 | 1003.6068 | 1010.94184 | 1159.27535 | 1080.66003 | 1120.74966 |
| RLTR45 | 11027.7279 | 11003.1062 | 11188.9632 | 12064.6393 | 11667.7538 | 12023.3647 |
| RLTR45-int | 10821.335 | 10667.4191 | 10883.2018 | 10160.47 | 9851.852 | 10061.9543 |
| RLTR46 | 274.47537 | 302.862212 | 285.10012 | 321.016446 | 288.067027 | 349.45789 |
| RLTR46A | 399.357197 | 431.517459 | 383.688178 | 463.399795 | 427.484998 | 474.346718 |

| | | | | | | |
|---|---|---|---|---|---|---|
| RLTR46A2 | 366.589637 | 365.474992 | 354.319034 | 344.246429 | 347.192171 | 356.1764 |
| RLTR46B | 490.120435 | 436.660271 | 461.529988 | 494.462397 | 473.803299 | 521.023657 |
| RLTR47 | 1642.26701 | 1643.45304 | 1636.73341 | 1496.71701 | 1335.09467 | 1481.73542 |
| RLTR47_MM | 122.395766 | 133.389295 | 125.7069 | 127.160451 | 103.926597 | 126.778222 |
| RLTR48A | 6098.45127 | 6173.72548 | 6321.54994 | 6074.46621 | 5788.89587 | 6045.59893 |
| RLTR48B | 863.452109 | 861.808122 | 816.299515 | 913.035016 | 832.346237 | 838.48602 |
| RLTR48C | 1458.93497 | 1457.1644 | 1503.84217 | 1460.62278 | 1552.38193 | 1466.05028 |
| RLTR49 | 6775.05195 | 6774.22113 | 6766.13468 | 6791.16615 | 6737.47054 | 6771.28995 |
| RLTR5_Mm | 1445.54316 | 1461.65372 | 1517.13141 | 1722.15949 | 1460.95832 | 1697.62767 |
| RLTR50A | 624.392882 | 630.622171 | 557.424378 | 588.426312 | 541.7949 | 651.718986 |
| RLTR50B | 2823.27799 | 2819.79446 | 2839.3748 | 3114.27519 | 2826.1309 | 3059.5927 |
| RLTR51A_Mm | 10773.6558 | 10769.7002 | 10911.8127 | 11189.9765 | 11225.0864 | 11447.0821 |
| RLTR51B_Mm | 4519.99823 | 4498.67652 | 4569.52737 | 4996.97371 | 6667.75606 | 6024.02726 |
| RLTR53_Mm | 3556.2009 | 3656.87019 | 3657.34492 | 4015.02155 | 4091.96143 | 4176.33707 |
| RLTR53B_Mm | 794.12184 | 787.848066 | 798.017041 | 855.619531 | 825.782996 | 801.325455 |
| RLTR6-int | 36059.1079 | 36391.9581 | 37128.6875 | 42182.3415 | 40214.782 | 44819.3678 |
| RLTR6_Mm | 1728.17032 | 1705.65787 | 1746.82796 | 1803.74416 | 1810.57342 | 1918.46709 |
| RLTR6B_Mm | 4810.52919 | 4779.17406 | 4811.63573 | 5110.35822 | 4899.1145 | 5349.05238 |
| RLTR6C_Mm | 524.729735 | 570.538423 | 533.573937 | 582.720047 | 500.949613 | 564.619504 |
| RLTR8 | 1936.87674 | 1912.02778 | 1856.8754 | 1886.29885 | 1828.83627 | 1898.55222 |
| RLTR9A | 4522.67407 | 4572.0658 | 4582.58888 | 5033.26992 | 4955.57179 | 5132.77202 |
| RLTR9A2 | 1742.51366 | 1811.53735 | 1742.40016 | 1806.3669 | 1802.35234 | 1840.15336 |
| RLTR9A3 | 5302.82741 | 5187.26459 | 5316.37808 | 5536.57859 | 5523.70781 | 5579.18522 |
| RLTR9A3A | 1459.33446 | 1513.90007 | 1493.10746 | 1556.68554 | 1419.04591 | 1497.50924 |
| RLTR9A3B | 2360.15874 | 2229.33758 | 2293.86744 | 2315.67196 | 2127.08179 | 2331.49901 |
| RLTR9A4 | 207.538367 | 208.656443 | 225.416545 | 206.471811 | 174.928787 | 198.715956 |
| RLTR9B | 1679.38701 | 1629.24877 | 1703.53348 | 1857.13312 | 1749.48902 | 1857.74107 |
| RLTR9B2 | 1153.82579 | 1139.60744 | 1143.99514 | 1259.96628 | 1141.56089 | 1328.85331 |
| RLTR9C | 1688.1957 | 1671.12647 | 1724.28686 | 1768.93 | 1576.88472 | 1813.19418 |
| RLTR9D | 5897.39858 | 5950.53478 | 6034.56358 | 6859.6088 | 6519.69467 | 6849.19482 |
| RLTR9D2 | 1554.015 | 1564.92125 | 1536.99805 | 1790.24795 | 1849.29658 | 1959.97949 |
| RLTR9E | 9384.36777 | 9498.02177 | 9330.8392 | 10550.8745 | 9943.50466 | 10364.7237 |
| RLTR9F | 3057.06889 | 3230.16889 | 3232.16705 | 3566.62703 | 3470.78283 | 3623.17858 |
| RLTRETN_Mm | 8852.44146 | 8888.45828 | 8920.99172 | 10055.371 | 10028.9959 | 10019.4205 |
| RMER10A | 30930.1548 | 31063.7885 | 30403.6794 | 34431.728 | 36041.4304 | 36314.42 |
| RMER10B | 18458.3711 | 18477.3242 | 18199.1966 | 21230.952 | 21864.1756 | 22031.9454 |
| RMER12 | 21496.0721 | 21509.8553 | 21486.4794 | 20386.7708 | 20077.1358 | 20096.8097 |
| RMER12B | 9093.6465 | 8943.14407 | 8873.0402 | 8595.04065 | 8258.98636 | 8492.54944 |
| RMER12C | 19240.8035 | 19372.5917 | 19201.7625 | 18731.4472 | 18388.6291 | 18317.738 |
| RMER13A | 12806.161 | 12746.557 | 13030.3542 | 14757.1455 | 14430.573 | 15080.6363 |
| RMER13A1 | 11793.1992 | 11796.3363 | 11900.2101 | 13918.3726 | 13617.3587 | 14166.2889 |
| RMER13A2 | 16115.0549 | 16033.9475 | 16506.0889 | 18174.8617 | 17613.6547 | 18465.0027 |

| | | | | | | |
|---|---|---|---|---|---|---|
| RMER13B | 27785.9711 | 28181.8892 | 28189.7179 | 33944.0525 | 32683.8161 | 34675.0802 |
| RMER15 | 114317.092 | 114033.247 | 114273.929 | 116397.64 | 115087.801 | 116699.318 |
| RMER15-int | 28240.6177 | 28147.5856 | 28106.5922 | 28707.78 | 28523.4463 | 29234.6636 |
| RMER16 | 3854.74681 | 3813.93205 | 3934.0248 | 3788.82522 | 3956.03717 | 3868.64443 |
| RMER16-int | 30654.0165 | 31089.8234 | 30893.1642 | 27239.7402 | 26233.3803 | 26107.4424 |
| RMER16_Mm | 3638.3206 | 3665.11391 | 3721.80826 | 3640.37762 | 3640.70045 | 3556.6729 |
| RMER16A2 | 1517.1954 | 1447.20391 | 1543.17143 | 1557.9035 | 1492.22916 | 1513.56019 |
| RMER16A3 | 2186.50776 | 2122.80439 | 2174.38534 | 2102.37545 | 1964.79637 | 2042.42465 |
| RMER16B | 761.381541 | 785.317921 | 755.862742 | 751.887247 | 822.468548 | 715.075163 |
| RMER16B2 | 2324.13917 | 2234.80825 | 2297.12654 | 2134.02881 | 2159.98431 | 2085.72763 |
| RMER16B3 | 652.35923 | 642.703339 | 661.458328 | 646.268974 | 580.65783 | 563.036145 |
| RMER16C | 1358.25632 | 1267.03842 | 1316.14067 | 1248.00566 | 1213.59559 | 1246.69846 |
| RMER17A | 17509.9437 | 17374.3666 | 17906.6274 | 20775.5295 | 19124.7676 | 21021.0794 |
| RMER17A-int | 3216.92124 | 3201.18928 | 3214.74776 | 3185.24948 | 2988.83867 | 3145.44828 |
| RMER17A2 | 20037.0664 | 19923.8766 | 20635.7149 | 24198.2598 | 23004.6717 | 25089.9543 |
| RMER17B | 37355.7952 | 37373.0273 | 37965.2577 | 46081.3317 | 44762.9007 | 47603.4245 |
| RMER17B2 | 9053.65353 | 9122.49455 | 9270.33394 | 10892.7898 | 10569.1092 | 11226.1469 |
| RMER17C | 39143.9917 | 39356.2197 | 39231.4788 | 45832.8573 | 47941.7455 | 49097.8633 |
| RMER17C-int | 15709.2944 | 15767.2382 | 16024.5222 | 15564.2203 | 15012.002 | 15360.6498 |
| RMER17C2 | 15740.9363 | 15603.2309 | 15817.645 | 17515.2957 | 17606.4283 | 17981.1057 |
| RMER17D | 8661.72988 | 8716.37017 | 8931.00607 | 10054.0312 | 9372.45894 | 10262.5055 |
| RMER17D2 | 14039.8982 | 13864.682 | 14131.5351 | 16451.0529 | 15390.7482 | 16829.0491 |
| RMER19A | 24501.7857 | 24522.471 | 24737.8209 | 25057.0229 | 24785.8286 | 25185.9722 |
| RMER19B | 74784.0028 | 75392.458 | 75011.1576 | 77595.4282 | 76775.015 | 78268.9908 |
| RMER19B2 | 26227.6681 | 26061.8427 | 25916.2407 | 24977.564 | 23706.2041 | 24031.1866 |
| RMER19C | 28044.3499 | 28030.607 | 27945.5656 | 26827.7923 | 25861.2563 | 26719.2273 |
| RMER1A | 50020.7314 | 50270.9272 | 50565.7936 | 60803.6326 | 60057.0198 | 62342.1898 |
| RMER1B | 80890.0108 | 81213.2563 | 82249.0352 | 94730.4047 | 93810.8618 | 96591.786 |
| RMER1C | 40985.3228 | 40696.7382 | 40862.6752 | 43152.111 | 41918.168 | 44083.069 |
| RMER2 | 19763.6768 | 19937.0233 | 19745.3731 | 21236.3865 | 20766.3302 | 21199.3657 |
| RMER20A | 10989.6312 | 11118.5328 | 11080.3286 | 11707.8395 | 11283.8998 | 11932.0649 |
| RMER20B | 15361.9342 | 15312.8761 | 15627.709 | 16304.7177 | 15429.9521 | 16188.6867 |
| RMER20C_Mm | 13411.4942 | 13346.4828 | 13266.6483 | 15143.4939 | 14554.681 | 14989.1024 |
| RMER21A | 5236.08544 | 5338.11906 | 5085.56924 | 6276.40326 | 6350.94383 | 6480.86629 |
| RMER21B | 7888.50049 | 7951.45977 | 8060.67305 | 8901.28656 | 9103.39135 | 9234.56352 |
| RMER30 | 15168.3067 | 15245.6111 | 15176.5497 | 15276.0856 | 15474.9681 | 15046.7651 |
| RMER3D-int | 22655.5233 | 22267.6567 | 22408.7441 | 19836.9254 | 19302.9874 | 19489.5337 |
| RMER3D1 | 3.96134022 | 3.10202958 | 0 | 7.47182782 | 2.86838257 | 9.14447234 |
| RMER3D2 | 12.3957025 | 10.2858261 | 6.17554117 | 9.9779235 | 23.795421 | 8.12402874 |
| RMER3D3 | 24.8700446 | 22.3677026 | 20.1643881 | 28.8708377 | 23.997179 | 18.2230778 |
| RMER3D4 | 3.96134022 | 6.85714996 | 6.20838304 | 4.58550211 | 4.40345286 | 9.14447234 |
| RMER4A | 18677.9865 | 18483.0336 | 18353.7537 | 18943.504 | 20762.7096 | 20102.1222 |

| | | | | | | |
|---|---|---|---|---|---|---|
| RMER4B | 39668.5468 | 39469.3878 | 38887.5964 | 39972.3932 | 42032.399 | 41377.6452 |
| RMER5 | 55137.9379 | 55526.5152 | 55160.0071 | 52248.9689 | 52828.9267 | 52665.8313 |
| RMER6-int | 5393.88214 | 5211.50429 | 5419.14266 | 4715.87557 | 4326.0974 | 4604.94826 |
| RMER6A | 43925.1678 | 43822.332 | 43790.9227 | 42870.6803 | 42916.8733 | 43248.6383 |
| RMER6B | 10344.5928 | 10090.8382 | 10155.6916 | 9831.14878 | 11213.2021 | 10629.6078 |
| RMER6BA | 5307.71513 | 5297.46188 | 5325.58067 | 5321.15475 | 5216.61649 | 5263.36245 |
| RMER6C | 46586.2138 | 45912.8938 | 46814.3236 | 42972.6347 | 39414.0787 | 41970.9263 |
| RMER6D | 18579.4657 | 18009.9699 | 18643.2343 | 16918.6724 | 15730.5653 | 16654.6048 |
| RNERVK23-int | 208.456878 | 232.1667 | 219.70079 | 217.931776 | 251.081912 | 240.448757 |
| RNLTR23 | 21.6038787 | 21.8778339 | 19.4932942 | 32.4787448 | 17.573704 | 28.3353003 |
| RodERV21-int | 22202.8621 | 22241.6327 | 22747.7001 | 23404.2922 | 23625.408 | 24011.3468 |
| RSINE1 | 483548.744 | 484870.253 | 477925.06 | 505027.472 | 535860.962 | 522925.523 |
| SRV_MM-int | 2127.21243 | 2113.91 | 2139.35683 | 2027.20395 | 1956.07212 | 2024.75131 |
| SSU-rRNA_Hsa | 4720.81343 | 4733.05855 | 4569.60668 | 6472.14047 | 11269.6271 | 8364.2375 |
| SUBTEL_sa | 87.7272237 | 90.7767895 | 66.4245558 | 92.8971915 | 90.0903004 | 91.2207761 |
| SYNREP_MM | 185154.106 | 185187.538 | 186757.029 | 180965.545 | 338365.794 | 271905.114 |
| T-rich | 51829.1933 | 51214.3216 | 51708.0294 | 46480.6777 | 47143.9642 | 46474.5556 |
| Tigger1 | 8781.51376 | 8681.43103 | 8820.54377 | 8153.75435 | 8165.3428 | 8056.24541 |
| Tigger10 | 1571.32305 | 1631.37045 | 1600.23198 | 1493.78422 | 1698.58862 | 1557.97229 |
| Tigger11a | 505.218719 | 492.007744 | 452.470855 | 460.11001 | 424.352865 | 422.439368 |
| Tigger12 | 874.733226 | 916.66603 | 898.930379 | 834.24375 | 866.806935 | 873.250016 |
| Tigger12A | 502.555457 | 463.191048 | 487.31377 | 435.824674 | 504.304168 | 483.734898 |
| Tigger12c | 580.595327 | 588.497308 | 561.044747 | 531.491556 | 609.949371 | 535.316454 |
| Tigger13a | 4240.31114 | 4281.93886 | 4336.0162 | 3989.69456 | 4101.31222 | 4004.76826 |
| Tigger14a | 1179.48276 | 1099.52488 | 1094.29523 | 1084.99514 | 1093.79323 | 1030.52957 |
| Tigger15a | 4078.51797 | 4084.7114 | 4041.90341 | 4063.28071 | 4205.48374 | 4037.93523 |
| Tigger16a | 1680.11678 | 1643.86008 | 1574.92161 | 1605.3259 | 1695.64185 | 1560.35873 |
| Tigger16b | 982.555496 | 945.647567 | 948.458947 | 948.497283 | 1044.82836 | 951.498379 |
| Tigger17 | 1515.27397 | 1538.22543 | 1552.78671 | 1555.12195 | 1608.61803 | 1510.58094 |
| Tigger17a | 7508.58091 | 7601.98813 | 7516.87066 | 7192.06663 | 7530.54423 | 7299.83264 |
| Tigger17b | 467.584939 | 408.170525 | 474.512194 | 403.116936 | 432.352739 | 406.773986 |
| Tigger17c | 3769.81497 | 3960.22085 | 3820.82543 | 3647.06148 | 3803.79474 | 3816.20355 |
| Tigger18a | 1646.41918 | 1713.41314 | 1645.18497 | 1526.13447 | 1564.34163 | 1449.71001 |
| Tigger19a | 3658.70556 | 3712.13655 | 3692.7254 | 3690.57827 | 3821.75496 | 3747.79785 |
| Tigger1a_Art | 167.689046 | 165.39034 | 143.168327 | 138.922888 | 140.468089 | 144.909086 |
| Tigger1a_Mars | 10.4674588 | 6.0410394 | 2.60443524 | 5.30708354 | 3.87839363 | 10.1385692 |
| Tigger20a | 1151.69413 | 1150.05882 | 1088.366 | 1095.07405 | 1138.2495 | 1029.24261 |
| Tigger2b | 1092.80406 | 1052.74738 | 998.672866 | 960.200915 | 1037.21195 | 984.527523 |
| Tigger5 | 11000.6061 | 10881.2981 | 10935.7732 | 10589.9825 | 10545.6918 | 10650.0897 |
| Tigger5b | 11143.595 | 11164.3209 | 11264.9488 | 11119.9394 | 11116.6968 | 10874.5366 |
| Tigger6a | 2937.13402 | 2860.4478 | 2970.28128 | 2750.22745 | 2846.39128 | 2746.94982 |
| Tigger6b | 1320.85217 | 1316.75281 | 1243.73049 | 1195.83894 | 1168.7127 | 1186.82821 |

| | | | | | | |
|---|---|---|---|---|---|---|
| Tigger7 | 30358.5386 | 30353.3232 | 30363.922 | 29101.8031 | 29940.9117 | 29607.0906 |
| Tigger8 | 811.954162 | 792.826746 | 795.27411 | 764.708339 | 719.551962 | 705.585136 |
| Tigger9a | 1180.33521 | 1158.38406 | 1099.91895 | 1117.74554 | 1140.02826 | 1116.60861 |
| Tigger9b | 2277.6988 | 2198.48051 | 2292.19683 | 2182.70328 | 2172.67328 | 2259.35051 |
| tRNA-Ala-GCA | 2057.34951 | 2085.33551 | 2068.76289 | 2203.11333 | 2268.016 | 2240.63697 |
| tRNA-Ala-GCG | 519.797458 | 500.08906 | 544.79414 | 506.640153 | 561.388626 | 526.675617 |
| tRNA-Ala-GCY | 32.7140853 | 40.6539417 | 36.1194182 | 42.0764383 | 36.1574191 | 37.5719862 |
| tRNA-Ala-GCY_ | 7160.35939 | 7228.10467 | 7183.07231 | 7431.31354 | 8215.65799 | 7862.0718 |
| tRNA-Arg-AGA | 74.0460254 | 76.8173529 | 67.5225939 | 74.3456287 | 59.6295389 | 72.0431215 |
| tRNA-Arg-AGG | 22.0432121 | 25.6331566 | 26.6355061 | 23.4784163 | 14.8870258 | 13.1735533 |
| tRNA-Arg-CGA | 9.85092411 | 10.122604 | 8.84566015 | 9.00032853 | 11.8369389 | 11.1853596 |
| tRNA-Arg-CGA_ | 4.65651455 | 9.79615978 | 9.68096336 | 2.25008213 | 5.57511453 | 5.08904466 |
| tRNA-Arg-CGG | 13.7860511 | 13.7147046 | 5.27455422 | 12.4840192 | 8.12019588 | 5.08904466 |
| tRNA-Arg-CGY_ | 55.5573246 | 50.7762422 | 53.9092641 | 52.4345918 | 54.9428923 | 54.7746204 |
| tRNA-Asn-AAC | 91.3934032 | 95.3479193 | 91.1267205 | 121.643813 | 75.5065221 | 98.2058008 |
| tRNA-Asp-GAY | 86.3169531 | 96.4912835 | 88.268843 | 83.5981805 | 77.3849473 | 78.5376845 |
| tRNA-Cys-TGY | 281.833942 | 251.922545 | 266.474971 | 279.405575 | 245.809434 | 221.909518 |
| tRNA-Gln-CAA | 28.9100244 | 33.9596091 | 29.2070994 | 37.8711662 | 23.8355284 | 17.7523761 |
| tRNA-Gln-CAA_ | 773.049041 | 790.544064 | 790.999069 | 180.98319 | 39.0659091 | 42.5819906 |
| tRNA-Gln-CAG | 69.5467902 | 80.1641144 | 57.4475282 | 68.992086 | 58.1345761 | 68.9686173 |
| tRNA-Glu-GAA | 15.1239733 | 12.2450986 | 20.7134071 | 23.7344298 | 15.8769831 | 12.166283 |
| tRNA-Glu-GAG | 15.6618683 | 20.0818847 | 13.2849112 | 8.72487566 | 14.0186116 | 11.1853596 |
| tRNA-Glu-GAG_ | 101.132659 | 111.348644 | 102.379765 | 74.6869801 | 99.1803999 | 81.1875939 |
| tRNA-Gly-GGA | 201.976973 | 218.452198 | 202.713406 | 231.346931 | 221.226425 | 199.762747 |
| tRNA-Gly-GGG | 51.0780113 | 63.5111082 | 42.4591687 | 42.8601279 | 50.196695 | 69.9758875 |
| tRNA-Gly-GGY | 149.180423 | 125.389692 | 130.521668 | 103.131129 | 130.934366 | 92.6921613 |
| tRNA-His-CAY | 5.91579711 | 1.30618162 | 5.24171235 | 5.39242139 | 7.11018482 | 3.04815745 |
| tRNA-His-CAY_ | 44.060211 | 57.47017 | 59.2166602 | 53.4121868 | 54.9027849 | 43.628781 |
| tRNA-Ile-ATA | 50.2187424 | 58.1232608 | 53.1068028 | 63.1340967 | 56.7611564 | 55.7555439 |
| tRNA-Ile-ATT | 95.0333694 | 99.4303945 | 86.4340272 | 108.694226 | 113.360662 | 105.892061 |
| tRNA-Leu-CTA | 5.91579711 | 7.67366524 | 8.91134388 | 3.86392068 | 8.4434971 | 8.09768199 |
| tRNA-Leu-CTA_ | 13.11709 | 16.6533097 | 10.5491084 | 9.34167991 | 16.2403918 | 11.1326661 |
| tRNA-Leu-CTG | 46.6574158 | 57.7966142 | 56.3823319 | 68.2705046 | 51.5494505 | 68.4452221 |
| tRNA-Leu-CTY | 27.545889 | 25.4697321 | 13.317753 | 20.6774284 | 19.2704249 | 22.2916789 |
| tRNA-Leu-TTA | 39.2202039 | 38.2049017 | 37.6679498 | 55.2355797 | 45.0858681 | 33.5165585 |
| tRNA-Leu-TTA(m) | 797.321949 | 797.237486 | 795.095644 | 180.890271 | 50.1347024 | 40.5147567 |
| tRNA-Leu-TTG | 63.5261402 | 55.1844534 | 67.4897521 | 51.751889 | 68.9213965 | 48.730999 |
| tRNA-Lys-AAA | 54.9869251 | 62.9398308 | 51.1406195 | 58.6339325 | 81.2232335 | 62.8327823 |
| tRNA-Lys-AAG | 446.84662 | 411.597481 | 422.749743 | 440.437196 | 479.580172 | 466.36101 |
| tRNA-Met | 45.1622142 | 45.7147378 | 46.8655779 | 78.9311308 | 43.7926632 | 58.790528 |
| tRNA-Met-i | 46.44771 | 45.3882936 | 48.4376587 | 52.481051 | 55.9127959 | 57.862298 |
| tRNA-Met_ | 102.903623 | 128.001954 | 118.925948 | 102.619101 | 96.9786735 | 113.565148 |

| | | | | | | |
|---|---|---|---|---|---|---|
| tRNA-Phe-TTY | 99.5257894 | 108.164498 | 72.6657807 | 105.718772 | 107.401475 | 95.8588791 |
| tRNA-Pro-CCA | 29.6576252 | 26.7759138 | 32.9095728 | 16.0065885 | 28.5622825 | 29.3689172 |
| tRNA-Pro-CCG | 18.363926 | 19.265673 | 16.8560171 | 9.17100422 | 17.573704 | 11.1458394 |
| tRNA-Pro-CCY | 77.6335653 | 91.5931024 | 81.1830222 | 77.7828602 | 118.672637 | 108.489277 |
| tRNA-SeC(e)-TGA | 7.22750611 | 9.4695132 | 10.6147922 | 6.7502464 | 6.42347498 | 11.1063193 |
| tRNA-Ser-AGY | 51.7207592 | 46.5312531 | 38.8223791 | 32.8200962 | 38.1373339 | 31.4493246 |
| tRNA-Ser-TCA | 4.57787489 | 1.95927243 | 3.50542219 | 0.80691928 | 1.85837151 | 0 |
| tRNA-Ser-TCA(m) | 9.15574978 | 6.20405915 | 2.20320457 | 3.69324499 | 6.94853421 | 8.09768199 |
| tRNA-Ser-TCA_ | 12.4743421 | 13.5512802 | 14.1858981 | 10.6995049 | 11.6752883 | 14.1939969 |
| tRNA-Ser-TCG | 10.5460984 | 12.4083207 | 9.7466471 | 18.3420084 | 13.856961 | 13.1999 |
| tRNA-Ser-TCY | 37.3181735 | 40.0005473 | 45.293497 | 62.1565018 | 32.6023267 | 31.4888447 |
| tRNA-Thr-ACA | 30.8120549 | 31.3475494 | 40.2395432 | 30.1044462 | 26.8655616 | 33.5165585 |
| tRNA-Thr-ACG | 9.87713733 | 7.02057443 | 14.1530562 | 10.4046127 | 3.55509241 | 4.06860106 |
| tRNA-Thr-ACG_ | 45.909815 | 47.5109905 | 32.7782054 | 46.2352511 | 46.3176909 | 33.4506917 |
| tRNA-Thr-ACY | 44.6505324 | 39.0213158 | 45.2278133 | 46.7083996 | 49.2875629 | 31.4756713 |
| tRNA-Thr-ACY_ | 31.4285896 | 33.1433974 | 38.2897809 | 44.8385475 | 59.831297 | 36.4725024 |
| tRNA-Trp-TGG | 38.039561 | 47.0211218 | 39.0194303 | 43.736736 | 35.5108167 | 39.5206598 |
| tRNA-Tyr-TAC | 52.5469997 | 64.3275223 | 43.2616301 | 51.2474425 | 47.3477556 | 38.63195 |
| tRNA-Tyr-TAT | 8.51300189 | 9.4695132 | 8.84566015 | 7.55716567 | 16.9271016 | 9.10495222 |
| tRNA-Val-GTA | 42.0009012 | 39.1842344 | 27.5693349 | 35.3650705 | 34.9456501 | 31.462498 |
| tRNA-Val-GTG | 72.8391693 | 97.3072929 | 89.9301567 | 82.3451327 | 82.7783575 | 75.0385855 |
| tRNA-Val-GTY | 60.4696801 | 45.8783646 | 49.6999062 | 33.6270155 | 51.1860418 | 39.5470066 |
| U1 | 1393.06799 | 1374.9595 | 1396.37777 | 1401.41043 | 1410.84428 | 1394.30356 |
| U13 | 52.5732129 | 47.3477684 | 54.8102511 | 35.9624355 | 42.5407867 | 40.5542768 |
| U13_ | 80.1715283 | 83.2663464 | 68.8319398 | 65.1281653 | 89.9687572 | 76.9740857 |
| U14 | 14.428799 | 14.3675931 | 14.1858981 | 16.0065885 | 10.4635192 | 8.13720211 |
| U17 | 124.533715 | 111.348644 | 111.126899 | 88.7772573 | 107.380811 | 148.06263 |
| U2 | 2130.00623 | 2213.82773 | 2154.19091 | 2103.13212 | 2044.87104 | 2017.4339 |
| U3 | 251.310233 | 237.553941 | 214.883858 | 223.712008 | 211.066153 | 211.418808 |
| U4 | 362.884138 | 375.67951 | 349.08445 | 376.445931 | 368.279639 | 343.914857 |
| U5 | 213.946448 | 235.921618 | 230.937414 | 190.496033 | 237.003139 | 214.141171 |
| U6 | 4132.85481 | 4075.73348 | 3964.48702 | 3996.4173 | 4253.90228 | 4047.69885 |
| U7 | 114.636656 | 120.083253 | 118.236269 | 95.1240441 | 92.5746101 | 107.992229 |
| U8 | 25.0273239 | 38.3679215 | 22.0648876 | 22.5472805 | 28.2389813 | 23.364816 |
| UCON1 | 90.4817078 | 74.4503287 | 77.1050317 | 66.3617738 | 52.7612195 | 76.0985492 |
| UCON10 | 300.460001 | 313.311362 | 307.369187 | 315.592726 | 313.539726 | 267.6714 |
| UCON11 | 274.389915 | 235.595275 | 282.02877 | 266.483008 | 232.901713 | 219.085308 |
| UCON12 | 110.367049 | 106.77711 | 120.596555 | 100.34579 | 117.521029 | 107.389793 |
| UCON12A | 110.000063 | 116.736795 | 128.344177 | 111.153862 | 122.692627 | 91.2998163 |
| UCON13 | 254.41912 | 260.656244 | 264.25751 | 228.638862 | 253.990402 | 234.220709 |
| UCON14 | 308.081229 | 296.330292 | 300.858099 | 300.377896 | 243.890291 | 295.012604 |
| UCON15 | 169.53865 | 174.206358 | 194.604523 | 158.964073 | 191.634077 | 156.120792 |

| | | | | | | |
|---|---|---|---|---|---|---|
| UCON16 | 104.929905 | 131.91979 | 128.114284 | 103.131129 | 105.664036 | 113.538802 |
| UCON17 | 152.827205 | 133.715942 | 148.344356 | 122.404273 | 116.673889 | 110.490644 |
| UCON18 | 102.523008 | 98.4504548 | 104.9842 | 107.107407 | 91.5852632 | 88.3175257 |
| UCON19 | 116.125566 | 136.654345 | 120.300978 | 115.444472 | 111.663941 | 121.636484 |
| UCON2 | 390.496084 | 423.026773 | 410.347234 | 360.594369 | 435.888388 | 405.700849 |
| UCON20 | 243.381261 | 235.921719 | 230.198472 | 217.846439 | 210.602476 | 186.987442 |
| UCON21 | 135.053601 | 111.675291 | 106.621964 | 101.175939 | 134.651109 | 113.578322 |
| UCON22 | 125.045397 | 136.737069 | 120.972072 | 144.548094 | 90.251951 | 115.434782 |
| UCON23 | 100.332632 | 88.0012042 | 105.148409 | 118.035906 | 133.560883 | 101.319825 |
| UCON24 | 107.455285 | 82.9401045 | 97.9405136 | 85.2082288 | 79.768378 | 67.921827 |
| UCON25 | 105.369762 | 88.4912752 | 103.346435 | 95.0619358 | 76.5165331 | 82.1026505 |
| UCON26 | 652.182553 | 597.07001 | 653.152394 | 567.919559 | 596.233397 | 637.907017 |
| UCON27 | 389.71493 | 418.536546 | 437.409684 | 402.391564 | 382.984961 | 375.140234 |
| UCON28a | 525.431201 | 542.048677 | 503.179567 | 473.552185 | 482.771856 | 520.664929 |
| UCON28b | 318.561794 | 314.289986 | 302.545126 | 305.614803 | 301.701566 | 299.054858 |
| UCON28c | 211.454096 | 211.920177 | 211.296331 | 187.361762 | 188.462447 | 196.661896 |
| UCON29 | 819.260308 | 841.889662 | 863.711948 | 884.354293 | 798.854221 | 844.486581 |
| UCON31 | 503.735576 | 516.252703 | 477.632806 | 537.1016 | 478.205531 | 482.727628 |
| UCON4 | 341.864289 | 367.270435 | 362.048071 | 355.070151 | 368.725705 | 343.704083 |
| UCON5 | 293.684935 | 274.451649 | 239.471076 | 256.407888 | 239.607161 | 320.181186 |
| UCON6 | 231.372466 | 252.411402 | 235.425927 | 269.311016 | 230.235088 | 249.316589 |
| UCON7 | 480.820512 | 448.331765 | 451.339975 | 432.076902 | 414.498283 | 378.346472 |
| UCON8 | 288.726444 | 343.189204 | 308.605721 | 301.960437 | 335.112146 | 302.965379 |
| UCON9 | 159.609086 | 163.512375 | 166.870979 | 150.902461 | 181.292101 | 156.186659 |
| URR1A | 97647.8755 | 97728.0106 | 97345.1284 | 96303.3365 | 100352.049 | 97787.7069 |
| URR1B | 91390.1779 | 91801.5494 | 90658.6313 | 91080.9653 | 94727.5503 | 92120.8299 |
| X1_LINE | 306.7302 | 272.493186 | 283.126809 | 254.421888 | 276.371075 | 262.503315 |
| X2_LINE | 174.968979 | 187.10475 | 180.418625 | 167.196361 | 162.485353 | 146.963147 |
| X3_LINE | 980.128677 | 1004.5033 | 980.033461 | 1062.9177 | 936.718289 | 967.413563 |
| X5A_LINE | 276.61332 | 298.044428 | 311.719205 | 229.10443 | 226.901808 | 267.645054 |
| X5B_LINE | 248.175133 | 257.472704 | 228.593549 | 259.294214 | 208.257931 | 236.222076 |
| X6A_LINE | 624.58319 | 535.682104 | 580.78219 | 500.141642 | 510.60671 | 541.830777 |
| X6B_LINE | 1029.09443 | 1020.26102 | 1069.35821 | 1037.78278 | 1069.06679 | 1013.63297 |
| X7A_LINE | 628.269815 | 614.537912 | 605.550039 | 634.917576 | 643.398838 | 605.516289 |
| X7B_LINE | 650.069769 | 653.886988 | 664.60249 | 630.626966 | 627.885263 | 617.014269 |
| X7C_LINE | 621.468012 | 593.478314 | 617.729783 | 640.279188 | 644.531613 | 662.30191 |
| X7D_LINE | 34.0520076 | 35.5921338 | 32.7125216 | 29.7242161 | 31.7539663 | 22.3180256 |
| X8_LINE | 659.566815 | 675.764518 | 686.808037 | 628.105222 | 613.745719 | 605.980404 |
| X9_LINE | 183.26546 | 194.942244 | 175.709511 | 148.749575 | 182.92744 | 174.515124 |
| YREP_Mm | 5093.94588 | 5193.45914 | 5126.41202 | 5271.40121 | 5259.88776 | 5535.93188 |
| Zaphod | 4783.01999 | 4907.98636 | 4745.88226 | 4343.52255 | 4408.93836 | 4365.89538 |
| Zaphod2 | 1011.92425 | 983.767519 | 943.005927 | 935.900418 | 884.502793 | 873.009848 |

| | | | | | | |
|---|---|---|---|---|---|---|
| Zaphod3 | 2766.61814 | 2735.54696 | 2766.35489 | 2459.42781 | 2603.89629 | 2348.1291 |
| ZP3AR | 12170.9504 | 12406.4664 | 12386.1723 | 16805.1535 | 17762.9958 | 18811.34 |
| (CTACG)n | 0.64274789 | 1.95927243 | 0 | 2.16474429 | 2.70673196 | 3.06133082 |
| CENSAT_MC | 1.28549578 | 0.65309081 | 0 | 0 | 0 | 1.02044361 |
| (CACGA)n | 0 | 0.81631292 | 0.90098695 | 1.44316286 | 0.84836045 | 0 |
| | #DIV/0! | #DIV/0! | #DIV/0! | #DIV/0! | #DIV/0! | #DIV/0! |
