## Supplementary material for "Inhibition of Topoisomerase 2 catalytic activity impacts the integrity of heterochromatin and repetitive DNA and leads to interlinks between clustered repeats": Table S2

### Table S2: Repetitive sequences from END-seq analysis

#### Normalized values of END-seq reads by DESeq2 analysis

| 1 Biological replicate | END-seq values | | |
|---|---|---|---|
| Repeat family name | DMSO | ICRF-193 0h release | ICRF-193 4h release |
| (A)n | 33347.32 | 28179.5 | 34707.8 |
| (AAATG)n | 192.0231 | 131.9989 | 267.2076 |
| (AACTG)n | 58.44181 | 91.03375 | 105.5358 |
| (AAGTG)n | 0 | 0 | 8.981768 |
| (AATAG)n | 617.8135 | 568.961 | 467.052 |
| (AATTG)n | 37.56974 | 106.9647 | 145.9537 |
| (ACATG)n | 48.00578 | 109.2405 | 121.2539 |
| (ACCG)n | 35.48253 | 0 | 20.20898 |
| (ACCTG)n | 43.83136 | 2.275844 | 58.38149 |
| (ACGTG)n | 0 | 13.65506 | 0 |
| (ACTAG)n | 93.92434 | 56.8961 | 116.763 |
| (ACTCG)n | 4.174415 | 0 | 0 |
| (ACTG)n | 292.2091 | 382.3418 | 350.289 |
| (ACTTG)n | 0 | 15.93091 | 0 |
| (AGATG)n | 89.74993 | 77.37869 | 65.11782 |
| (AGCTG)n | 198.2847 | 143.3782 | 224.5442 |
| (AGGGGG)n | 638.6855 | 302.6872 | 406.425 |
| (AGGTG)n | 25.04649 | 59.17194 | 285.1711 |
| (AGTAG)n | 123.1452 | 159.3091 | 51.64517 |
| (AGTTG)n | 41.74415 | 0 | 11.22721 |
| (ATAAG)n | 52.18019 | 81.93038 | 65.11782 |
| (ATAGG)n | 91.83714 | 111.5163 | 181.8808 |
| (ATATG)n | 1310.766 | 1081.026 | 844.2862 |
| (ATCTG)n | 52.18019 | 61.44778 | 20.20898 |
| (ATG)n | 4410.27 | 5660.024 | 4807.492 |
| (ATGGTG)n | 1561.231 | 2321.361 | 1852.49 |
| (ATGTG)n | 352.7381 | 391.4451 | 168.4082 |
| (ATTAG)n | 271.337 | 86.48207 | 85.3268 |
| (ATTG)n | 3702.706 | 3186.181 | 2914.584 |
| (ATTTG)n | 331.866 | 238.9636 | 476.0337 |
| (C)n | 233.7673 | 143.3782 | 206.5807 |
| (CA)n | 159364.6 | 164559.4 | 171938 |
| (CAA)n | 14128.31 | 13898.58 | 16153.71 |
| (CAAA)n | 41787.98 | 40742.16 | 44334.01 |
| (CAAAA)n | 37189.87 | 31447.61 | 36198.77 |
| (CAAAAA)n | 14871.35 | 14358.3 | 16178.41 |
| (CAAAC)n | 3807.067 | 3470.662 | 4124.877 |
| (CAAAG)n | 638.6855 | 591.7194 | 833.059 |

| | | | |
|---|---|---|---|
| (CAAAT)n | 498.8426 | 418.7553 | 368.2525 |
| (CAACC)n | 96.01155 | 63.72363 | 71.85415 |
| (CAACG)n | 0 | 0 | 0 |
| (CAACT)n | 0 | 6.827532 | 2.245442 |
| (CAAG)n | 1440.173 | 1793.365 | 1434.838 |
| (CAAGA)n | 116.8836 | 286.7563 | 222.2988 |
| (CAAGC)n | 66.79064 | 31.86181 | 67.36326 |
| (CAAGG)n | 29.22091 | 81.93038 | 49.39973 |
| (CAAGT)n | 0 | 0 | 0 |
| (CAAT)n | 2857.387 | 2162.052 | 2916.829 |
| (CAATA)n | 419.5287 | 450.6171 | 249.2441 |
| (CAATC)n | 0 | 4.551688 | 98.79945 |
| (CAATG)n | 39.65694 | 125.1714 | 74.09959 |
| (CAATT)n | 231.68 | 197.9984 | 47.15428 |
| (CACAA)n | 563.5461 | 343.6524 | 386.216 |
| (CACAC)n | 294.2963 | 234.4119 | 404.1796 |
| (CACAG)n | 509.2787 | 703.2358 | 442.3521 |
| (CACAT)n | 141.9301 | 77.37869 | 143.7083 |
| (CACCAT)n | 2145.649 | 2464.739 | 1937.817 |
| (CACCC)n | 262.9882 | 562.1334 | 720.7869 |
| (CACCG)n | 0 | 0 | 0 |
| (CACCT)n | 75.13947 | 52.34441 | 188.6171 |
| (CACG)n | 1559.144 | 1533.919 | 1533.637 |
| (CACGC)n | 0 | 11.37922 | 0 |
| (CACGT)n | 37.56974 | 0 | 0 |
| (CACTA)n | 206.6336 | 218.481 | 309.871 |
| (CACTC)n | 139.8429 | 54.62025 | 112.2721 |
| (CACTG)n | 346.4765 | 227.5844 | 296.3984 |
| (CACTT)n | 0 | 0 | 35.92707 |
| (CAG)n | 3391.712 | 3875.762 | 3197.51 |
| (CAGA)n | 12946.95 | 13843.96 | 14667.23 |
| (CAGAA)n | 256.7265 | 163.8608 | 278.4348 |
| (CAGAC)n | 108.5348 | 43.24103 | 116.763 |
| (CAGAG)n | 187.8487 | 293.5839 | 372.7434 |
| (CAGAGA)n | 8574.249 | 10846.67 | 9846.264 |
| (CAGAT)n | 0 | 52.34441 | 62.87238 |
| (CAGC)n | 162.8022 | 359.5833 | 509.7154 |
| (CAGCC)n | 152.3662 | 245.7911 | 206.5807 |
| (CAGCG)n | 0 | 0 | 0 |
| (CAGCT)n | 210.808 | 54.62025 | 123.4993 |
| (CAGG)n | 1135.441 | 1069.647 | 1351.756 |
| (CAGGA)n | 141.9301 | 227.5844 | 175.1445 |

| | | | |
|---|---|---|---|
| (CAGGC)n | 298.4707 | 131.9989 | 107.7812 |
| (CAGGG)n | 402.8311 | 398.2727 | 186.3717 |
| (CAGGT)n | 12.52325 | 0 | 49.39973 |
| (CAGT)n | 304.7323 | 257.1704 | 538.9061 |
| (CAGTA)n | 333.9532 | 209.3776 | 244.7532 |
| (CAGTC)n | 79.31389 | 52.34441 | 2.245442 |
| (CAGTG)n | 214.9824 | 364.135 | 251.4895 |
| (CAGTT)n | 123.1452 | 223.0327 | 42.6634 |
| (CAT)n | 4735.874 | 5125.2 | 4342.685 |
| (CATA)n | 30992.95 | 32166.78 | 29606.15 |
| (CATAA)n | 402.8311 | 177.5158 | 330.08 |
| (CATAC)n | 152.3662 | 38.68935 | 85.3268 |
| (CATAG)n | 162.8022 | 79.65453 | 136.972 |
| (CATAT)n | 1237.714 | 1429.23 | 851.0226 |
| (CATATA)n | 16653.83 | 21058.38 | 20027.1 |
| (CATCC)n | 156.5406 | 61.44778 | 141.4629 |
| (CATCG)n | 0 | 0 | 0 |
| (CATCT)n | 79.31389 | 27.31013 | 74.09959 |
| (CATG)n | 3673.485 | 2797.012 | 3226.7 |
| (CATGC)n | 33.39532 | 152.4815 | 107.7812 |
| (CATGT)n | 112.7092 | 125.1714 | 29.19075 |
| (CATTA)n | 160.715 | 268.5496 | 33.68163 |
| (CATTC)n | 93.92434 | 100.1371 | 161.6718 |
| (CATTG)n | 81.4011 | 0 | 0 |
| (CATTT)n | 244.2033 | 325.4457 | 406.425 |
| (CCA)n | 2458.731 | 2480.67 | 2674.322 |
| (CCAA)n | 3423.02 | 2583.083 | 3478.19 |
| (CCCA)n | 953.8539 | 1110.612 | 956.5583 |
| (CCCCAA)n | 198.2847 | 81.93038 | 166.1627 |
| (CCCCAG)n | 400.7439 | 960.4061 | 884.7042 |
| (CCCCCA)n | 958.0283 | 823.8555 | 1172.121 |
| (CCCCCG)n | 333.9532 | 109.2405 | 121.2539 |
| (CCCCCT)n | 452.9241 | 309.5148 | 401.9341 |
| (CCCCG)n | 225.4184 | 634.9604 | 590.5513 |
| (CCCG)n | 83.4883 | 129.7231 | 143.7083 |
| (CCCGA)n | 0 | 0 | 0 |
| (CCCGAA)n | 0 | 0 | 15.71809 |
| (CCCGG)n | 0 | 102.413 | 33.68163 |
| (CCCTA)n | 77.22668 | 0 | 24.69986 |
| (CCCTAA)n | 46240 | 14701.95 | 13479.39 |
| (CCCTAG)n | 48.00578 | 70.55116 | 172.899 |
| (CCCTG)n | 509.2787 | 273.1013 | 300.8892 |

| | | | |
|---|---|---|---|
| (CCG)n | 916.2841 | 1515.712 | 1394.42 |
| (CCGAA)n | 0 | 43.24103 | 29.19075 |
| (CCGAG)n | 0 | 47.79272 | 71.85415 |
| (CCGCG)n | 133.5813 | 120.6197 | 89.81768 |
| (CCGG)n | 0 | 0 | 0 |
| (CCGGA)n | 0 | 0 | 0 |
| (CCGGG)n | 156.5406 | 70.55116 | 35.92707 |
| (CCGTA)n | 0 | 0 | 0 |
| (CCTA)n | 803.5749 | 562.1334 | 538.9061 |
| (CCTAA)n | 131.4941 | 25.03428 | 65.11782 |
| (CCTAG)n | 175.3254 | 0 | 69.60871 |
| (CCTAT)n | 102.2732 | 9.103375 | 65.11782 |
| (CCTCG)n | 0 | 161.5849 | 150.4446 |
| (CCTG)n | 1248.15 | 1399.644 | 1428.101 |
| (CCTTG)n | 110.622 | 152.4815 | 29.19075 |
| (CG)n | 638.6855 | 1183.439 | 967.7855 |
| (CGA)n | 440.4008 | 93.3096 | 119.0084 |
| (CGAA)n | 0 | 50.06856 | 74.09959 |
| (CGAAA)n | 0 | 0 | 0 |
| (CGAAG)n | 0 | 0 | 0 |
| (CGAAT)n | 0 | 0 | 0 |
| (CGAG)n | 43.83136 | 195.7226 | 204.3352 |
| (CGAGA)n | 33.39532 | 22.75844 | 24.69986 |
| (CGAGG)n | 2.087208 | 0 | 26.94531 |
| (CGAT)n | 20.87208 | 15.93091 | 65.11782 |
| (CGATA)n | 0 | 0 | 15.71809 |
| (CGATG)n | 0 | 6.827532 | 8.981768 |
| (CGCGG)n | 83.4883 | 11.37922 | 78.59047 |
| (CGCTG)n | 0 | 0 | 53.89061 |
| (CGG)n | 1085.348 | 1183.439 | 1327.056 |
| (CGGA)n | 93.92434 | 45.51688 | 29.19075 |
| (CGGAA)n | 0 | 0 | 0 |
| (CGGAG)n | 43.83136 | 0 | 29.19075 |
| (CGGG)n | 271.337 | 0 | 134.7265 |
| (CGGGA)n | 0 | 0 | 0 |
| (CGGGG)n | 542.674 | 573.5127 | 913.8949 |
| (CGGGGG)n | 87.66272 | 109.2405 | 53.89061 |
| (CGGT)n | 16.69766 | 36.4135 | 15.71809 |
| (CGGTG)n | 87.66272 | 25.03428 | 0 |
| (CGTAG)n | 0 | 0 | 0 |
| (CGTG)n | 1592.539 | 1982.26 | 1908.626 |
| (CGTTG)n | 0 | 0 | 4.490884 |

| | | | |
|---|---|---|---|
| (CTA)n | 1054.04 | 1463.368 | 1255.202 |
| (CTAA)n | 1001.86 | 1044.612 | 758.9594 |
| (CTAAA)n | 396.5694 | 416.4794 | 119.0084 |
| (CTAAG)n | 48.00578 | 0 | 11.22721 |
| (CTAAT)n | 171.151 | 125.1714 | 186.3717 |
| (CTACT)n | 60.52902 | 184.3434 | 78.59047 |
| (CTAG)n | 118.9708 | 38.68935 | 60.62694 |
| (CTAGG)n | 35.48253 | 31.86181 | 74.09959 |
| (CTAGGG)n | 152.3662 | 84.20622 | 166.1627 |
| (CTAGT)n | 0 | 38.68935 | 24.69986 |
| (CTATA)n | 1191.796 | 794.2695 | 952.0675 |
| (CTATG)n | 106.4476 | 0 | 139.2174 |
| (CTATT)n | 688.7785 | 614.4778 | 653.4237 |
| (CTCA)n | 1223.104 | 976.337 | 1023.922 |
| (CTCCG)n | 45.91857 | 0 | 62.87238 |
| (CTCG)n | 102.2732 | 88.75791 | 139.2174 |
| (CTCGG)n | 114.7964 | 45.51688 | 44.90884 |
| (CTCTG)n | 346.4765 | 366.4109 | 253.735 |
| (CTG)n | 3230.997 | 3249.905 | 2988.683 |
| (CTGGGG)n | 463.3601 | 443.7896 | 615.2511 |
| (CTGTG)n | 1006.034 | 839.7864 | 1050.867 |
| (CTTA)n | 824.447 | 416.4794 | 559.1151 |
| (CTTAA)n | 0 | 0 | 24.69986 |
| (CTTAG)n | 0 | 0 | 58.38149 |
| (CTTAT)n | 75.13947 | 102.413 | 65.11782 |
| (CTTCG)n | 0 | 0 | 0 |
| (CTTG)n | 943.4178 | 1263.093 | 916.1404 |
| (CTTTA)n | 25.04649 | 20.48259 | 8.981768 |
| (CTTTG)n | 740.9587 | 587.1677 | 464.8065 |
| (G)n | 171.151 | 186.6192 | 408.6705 |
| (GA)n | 44309.33 | 52394.48 | 44367.69 |
| (GAA)n | 6278.321 | 9208.064 | 6565.673 |
| (GAAA)n | 23629.28 | 30976.51 | 24861.54 |
| (GAAAA)n | 7996.092 | 7316.838 | 7558.158 |
| (GAATG)n | 54.2674 | 143.3782 | 69.60871 |
| (GACTG)n | 0 | 20.48259 | 76.34503 |
| (GAGAA)n | 2496.3 | 2592.186 | 2842.73 |
| (GAGTG)n | 54.2674 | 43.24103 | 121.2539 |
| (GATTG)n | 123.1452 | 15.93091 | 134.7265 |
| (GCATG)n | 85.57551 | 0 | 53.89061 |
| (GCCTAA)n | 20.87208 | 6.827532 | 58.38149 |
| (GCCTG)n | 31.30811 | 86.48207 | 406.425 |

| | | | |
|---|---|---|---|
| (GCGTG)n | 0 | 0 | 0 |
| (GCTG)n | 73.05227 | 102.413 | 262.7167 |
| (GCTTG)n | 102.2732 | 6.827532 | 150.4446 |
| (GGA)n | 6518.349 | 6734.222 | 6853.089 |
| (GGAA)n | 10895.22 | 13971.41 | 10706.27 |
| (GGAAA)n | 1611.324 | 2164.328 | 1457.292 |
| (GGAGA)n | 1001.86 | 985.4404 | 821.8318 |
| (GGAGAA)n | 786.8773 | 1322.265 | 833.059 |
| (GGATG)n | 123.1452 | 61.44778 | 161.6718 |
| (GGCTG)n | 189.9359 | 229.8602 | 370.4979 |
| (GGGA)n | 2874.085 | 3495.696 | 2813.539 |
| (GGGAA)n | 275.5114 | 600.8228 | 435.6158 |
| (GGGAGA)n | 1333.726 | 1920.812 | 1257.448 |
| (GGGGA)n | 277.5986 | 498.4098 | 653.4237 |
| (GGGTG)n | 538.4996 | 457.4446 | 902.6677 |
| (GGTTG)n | 112.7092 | 161.5849 | 157.1809 |
| (GTATG)n | 214.9824 | 116.068 | 74.09959 |
| (GTCTG)n | 192.0231 | 63.72363 | 74.09959 |
| (GTGTG)n | 429.9648 | 277.653 | 433.3703 |
| (GTTTG)n | 3777.846 | 2988.183 | 3132.392 |
| (T)n | 32445.64 | 30528.17 | 32835.1 |
| (TA)n | 204051.7 | 174322.8 | 163933 |
| (TAA)n | 15520.48 | 17007.38 | 16584.84 |
| (TAAA)n | 31381.17 | 26499.93 | 31319.43 |
| (TAAAA)n | 7593.261 | 5837.539 | 7091.106 |
| (TAAAAA)n | 2638.23 | 1190.266 | 1603.246 |
| (TAAAG)n | 112.7092 | 18.20675 | 80.83592 |
| (TAAG)n | 818.1854 | 625.8571 | 601.7785 |
| (TAATG)n | 131.4941 | 152.4815 | 276.1894 |
| (TACGG)n | 0 | 0 | 0 |
| (TACTG)n | 298.4707 | 191.1709 | 80.83592 |
| (TAG)n | 1281.545 | 1634.056 | 1758.181 |
| (TAGA)n | 61221.97 | 61238.41 | 56751.3 |
| (TAGG)n | 434.1392 | 776.0628 | 808.3592 |
| (TAGGG)n | 18.78487 | 0 | 4.490884 |
| (TAGTG)n | 225.4184 | 361.8592 | 276.1894 |
| (TATAA)n | 13166.11 | 13036.03 | 11784.08 |
| (TATAG)n | 1534.098 | 1370.058 | 987.9945 |
| (TATATG)n | 19699.07 | 20223.15 | 18899.89 |
| (TATCG)n | 0 | 47.79272 | 0 |
| (TATG)n | 28957.92 | 31199.54 | 29738.64 |
| (TATTG)n | 642.8599 | 254.8945 | 312.1165 |

| | | | |
|---|---|---|---|
| (TC)n | 43499.49 | 56452.31 | 46741.12 |
| (TCC)n | 7117.378 | 7983.66 | 6621.809 |
| (TCCA)n | 9229.632 | 9406.063 | 8889.705 |
| (TCCC)n | 2911.655 | 3117.906 | 3713.961 |
| (TCCCC)n | 686.6913 | 512.0649 | 617.4966 |
| (TCCCG)n | 0 | 0 | 35.92707 |
| (TCCG)n | 246.2905 | 254.8945 | 159.4264 |
| (TCCGG)n | 0 | 0 | 0 |
| (TCCTG)n | 91.83714 | 311.7906 | 226.7897 |
| (TCG)n | 56.35461 | 207.1018 | 244.7532 |
| (TCGGG)n | 0 | 0 | 22.45442 |
| (TCGTG)n | 0 | 0 | 0 |
| (TCTA)n | 57654.94 | 60974.41 | 59713.04 |
| (TCTCC)n | 1018.557 | 1005.923 | 714.0506 |
| (TCTCCC)n | 1089.522 | 1465.643 | 1437.083 |
| (TCTCG)n | 0 | 0 | 0 |
| (TCTCTG)n | 9897.539 | 10994.6 | 9136.704 |
| (TCTG)n | 11387.8 | 14253.61 | 12646.33 |
| (TCTTG)n | 265.0754 | 261.722 | 199.8443 |
| (TG)n | 160015.8 | 166896.7 | 174527 |
| (TGAA)n | 7029.715 | 8454.76 | 6965.361 |
| (TGAG)n | 757.6564 | 1015.026 | 1185.593 |
| (TGG)n | 2494.213 | 3391.007 | 3139.128 |
| (TGGA)n | 9292.248 | 9695.095 | 8932.369 |
| (TGGG)n | 822.3598 | 719.1667 | 1037.394 |
| (TGGGGG)n | 509.2787 | 1051.44 | 943.0857 |
| (TTA)n | 14865.09 | 16629.59 | 15421.7 |
| (TTAA)n | 6831.431 | 6606.775 | 7427.922 |
| (TTAAA)n | 492.581 | 425.5828 | 303.1347 |
| (TTAAG)n | 0 | 31.86181 | 17.96354 |
| (TTACG)n | 0 | 29.58597 | 0 |
| (TTAG)n | 578.1565 | 823.8555 | 651.1782 |
| (TTAGG)n | 0 | 84.20622 | 20.20898 |
| (TTAGGC)n | 96.01155 | 25.03428 | 87.57224 |
| (TTAGGG)n | 52207.32 | 14988.71 | 18846 |
| (TTATA)n | 13740.09 | 14158.02 | 12825.97 |
| (TTATG)n | 235.8545 | 157.0332 | 161.6718 |
| (TTC)n | 6069.6 | 7610.422 | 6583.636 |
| (TTCA)n | 8338.394 | 6178.916 | 7484.059 |
| (TTCC)n | 11037.15 | 12357.83 | 10852.22 |
| (TTCCC)n | 601.1158 | 382.3418 | 390.7069 |
| (TTCCG)n | 0 | 0 | 0 |

| | | | |
|---|---|---|---|
| (TTCG)n | 37.56974 | 0 | 121.2539 |
| (TTCGG)n | 0 | 0 | 0 |
| (TTCGGG)n | 39.65694 | 0 | 11.22721 |
| (TTCTC)n | 2627.794 | 3174.802 | 2413.85 |
| (TTCTCC)n | 603.203 | 1083.302 | 817.3409 |
| (TTCTG)n | 367.3485 | 452.8929 | 285.1711 |
| (TTG)n | 15856.52 | 14756.57 | 16396.22 |
| (TTGG)n | 3481.462 | 3142.94 | 3592.707 |
| (TTGGGG)n | 79.31389 | 84.20622 | 130.2356 |
| (TTGTG)n | 258.8137 | 525.7199 | 480.5246 |
| (TTTA)n | 32520.78 | 27792.61 | 32657.71 |
| (TTTAA)n | 642.8599 | 514.3407 | 231.2805 |
| (TTTAG)n | 204.5463 | 566.6851 | 132.4811 |
| (TTTC)n | 25678.92 | 32328.36 | 27587.5 |
| (TTTCC)n | 2024.591 | 2166.603 | 1583.037 |
| (TTTCG)n | 33.39532 | 34.13766 | 11.22721 |
| (TTTG)n | 41151.39 | 38115.83 | 43000.22 |
| (TTTTA)n | 6898.221 | 6304.087 | 8680.879 |
| (TTTTC)n | 7470.116 | 7683.249 | 8085.837 |
| (TTTTG)n | 34242.73 | 30573.69 | 34290.15 |
| (TTTTTA)n | 1861.789 | 1381.437 | 1919.853 |
| (TTTTTG)n | 15497.52 | 13461.62 | 14723.36 |
| 4.5SRNA | 3132.899 | 2956.321 | 3076.256 |
| 5S | 2988.881 | 3006.39 | 3440.017 |
| 7SK | 3679.747 | 2562.6 | 3271.609 |
| 7SLRNA | 2270.882 | 1725.09 | 2306.069 |
| A-rich | 72186.08 | 68739.59 | 70753.88 |
| AmnSINE1 | 5297.333 | 4947.685 | 5436.215 |
| AmnSINE2 | 1039.429 | 1454.264 | 1488.728 |
| Arthur1 | 9052.219 | 6461.121 | 8694.352 |
| Arthur1A | 446.6624 | 741.9251 | 678.1235 |
| Arthur1B | 4093.014 | 4581.274 | 4053.023 |
| Arthur1C | 922.5458 | 523.4441 | 866.7407 |
| AT_rich | 641584.7 | 560483.4 | 628077.1 |
| B1_Mm | 159652.6 | 146755.5 | 175434.1 |
| B1_Mur1 | 164142.2 | 163974.5 | 172483.6 |
| B1_Mur2 | 136735.1 | 131350.3 | 146663.3 |
| B1_Mur3 | 99186.19 | 101268.2 | 112483.2 |
| B1_Mur4 | 158437.8 | 156912.6 | 186349.2 |
| B1_Mus1 | 361423 | 345327.4 | 411519.9 |
| B1_Mus2 | 257306.8 | 249594.1 | 301632.5 |
| B1F | 166454.8 | 172436.1 | 177367.5 |

| | | | |
|---|---|---|---|
| B1F1 | 76391.8 | 74649.95 | 77200.55 |
| B1F2 | 63962.48 | 62349.02 | 67017.47 |
| B2_Mm1a | 89434.76 | 80259.91 | 95038.34 |
| B2_Mm1t | 106526.9 | 99961.89 | 123692.4 |
| B2_Mm2 | 470492.1 | 447622.1 | 511327.6 |
| B3 | 823075.7 | 820726.2 | 879016.5 |
| B3A | 429561.9 | 437399 | 454322.5 |
| B4 | 289220.2 | 292170.6 | 299088.4 |
| B4A | 518785.9 | 534852.9 | 536245.3 |
| BC1_Mm | 18818.26 | 17583.17 | 18744.95 |
| BGLII | 18455.09 | 16900.42 | 18316.07 |
| BGLII_A | 2097.644 | 1497.505 | 2027.634 |
| BGLII_B | 7906.342 | 8336.416 | 7807.402 |
| BGLII_B2 | 2212.44 | 1804.744 | 2317.296 |
| BGLII_C | 488.4066 | 566.6851 | 485.0155 |
| BGLII_Mur | 5618.763 | 4988.65 | 4282.058 |
| BGLII_Mus | 4097.189 | 4460.654 | 4621.12 |
| BLACKJACK | 8252.819 | 7046.013 | 7969.074 |
| C-rich | 26705.82 | 27542.26 | 27133.92 |
| C573_MM | 392.395 | 320.894 | 833.059 |
| Chap1_Mam | 676.2553 | 764.6835 | 976.7673 |
| Charlie1 | 9423.742 | 9781.577 | 9936.081 |
| Charlie10 | 2727.98 | 2842.529 | 2398.132 |
| Charlie10a | 336.0404 | 254.8945 | 419.8977 |
| Charlie10b | 336.0404 | 304.9631 | 103.2903 |
| Charlie11 | 482.145 | 464.2721 | 386.216 |
| Charlie12 | 5545.711 | 5025.063 | 6121.075 |
| Charlie13a | 1623.848 | 1279.024 | 1205.802 |
| Charlie13b | 421.6159 | 1158.405 | 1362.983 |
| Charlie14a | 628.2495 | 398.2727 | 642.1964 |
| Charlie15a | 884.976 | 1069.647 | 1044.131 |
| Charlie15b | 1966.15 | 2137.017 | 1975.989 |
| Charlie16a | 1344.162 | 942.1994 | 1172.121 |
| Charlie17 | 651.2088 | 1222.128 | 965.5401 |
| Charlie17a | 294.2963 | 348.2041 | 240.2623 |
| Charlie17b | 1619.673 | 830.683 | 1277.657 |
| Charlie18a | 1619.673 | 1397.368 | 1497.71 |
| Charlie19a | 1532.01 | 1638.608 | 1625.7 |
| Charlie1a | 24720.89 | 24362.91 | 24464.09 |
| Charlie1b | 6161.437 | 6882.152 | 7115.806 |
| Charlie1b_Mars | 27.1337 | 0 | 24.69986 |
| Charlie20a | 1490.266 | 1247.162 | 1327.056 |

| | | | |
|---|---|---|---|
| Charlie21a | 2001.632 | 1631.78 | 1823.299 |
| Charlie22a | 1529.923 | 1297.231 | 1062.094 |
| Charlie23a | 1085.348 | 869.3724 | 1005.958 |
| Charlie24 | 1792.911 | 2298.602 | 2959.493 |
| Charlie25 | 2523.434 | 2826.598 | 2452.023 |
| Charlie26a | 292.2091 | 516.6166 | 556.8696 |
| Charlie29a | 2719.632 | 2025.501 | 2166.852 |
| Charlie2a | 3919.776 | 4346.862 | 4520.075 |
| Charlie2b | 7263.483 | 7077.874 | 7171.942 |
| Charlie4 | 1158.4 | 978.6129 | 1082.303 |
| Charlie4a | 9066.83 | 7910.833 | 8364.272 |
| Charlie4z | 2153.998 | 2278.12 | 1841.263 |
| Charlie5 | 11191.61 | 10241.3 | 11714.47 |
| Charlie6 | 1231.452 | 1668.194 | 1650.4 |
| Charlie7 | 7708.058 | 8090.625 | 6536.482 |
| Charlie7a | 496.7554 | 559.8576 | 637.7056 |
| Charlie8 | 4752.572 | 4000.933 | 3900.333 |
| Charlie9 | 949.6795 | 1345.024 | 1356.247 |
| Cheshire | 6994.233 | 7335.045 | 7210.115 |
| Cheshire_Mars | 0 | 0 | 69.60871 |
| Cheshire_Mars_ | 20.87208 | 0 | 31.43619 |
| CR1_Mam | 1185.534 | 1606.746 | 1726.745 |
| CT-rich | 60414.22 | 64053.63 | 57108.33 |
| CYRA11_Mm | 3266.48 | 3017.769 | 3695.998 |
| DNA1_Mam | 237.9417 | 421.0311 | 211.0716 |
| ERV3-16A3_I-int | 7288.529 | 6012.779 | 6253.556 |
| ERV3-16A3_LTR | 1365.034 | 1638.608 | 1511.183 |
| ERVB2_1-I_MM-int | 1302.418 | 1110.612 | 1502.201 |
| ERVB2_1A-I_MM-int | 3850.898 | 2976.804 | 3076.256 |
| ERVB3_1-I_MM-int | 1888.923 | 2002.743 | 2719.23 |
| ERVB3_1-LTR_MM | 1625.935 | 1310.886 | 1104.758 |
| ERVB4_1-I_MM-int | 6756.291 | 5787.471 | 5934.703 |
| ERVB4_1-LTR_MM | 4.174415 | 109.2405 | 132.4811 |
| ERVB4_1B-I_MM-int | 17154.76 | 14986.43 | 17552.62 |
| ERVB4_1B-LTR_MM | 6524.611 | 4854.375 | 5393.552 |
| ERVB4_1C-LTR_Mm | 3389.625 | 3181.63 | 3217.719 |
| ERVB4_2-I_MM-int | 8513.72 | 7883.523 | 7960.092 |
| ERVB4_2-LTR_MM | 490.4938 | 480.2031 | 664.6509 |
| ERVB4_3-I_MM-int | 2487.951 | 3752.867 | 3597.198 |
| ERVB4_3-LTR_MM | 1003.947 | 1071.922 | 1239.484 |
| ERVB5_1-I_MM-int | 3454.329 | 4255.828 | 3752.134 |
| ERVB5_1-LTR_MM | 684.6041 | 1417.851 | 882.4587 |

| | | | |
|---|---|---|---|
| ERVB5_2-LTR_MM | 1863.876 | 1827.503 | 1942.307 |
| ERVB7_1-LTR_MM | 3815.416 | 4431.068 | 4008.114 |
| ERVB7_2-LTR_MM | 2913.742 | 2052.811 | 2519.386 |
| ERVB7_2B-LTR_MM | 2650.754 | 2535.29 | 2788.839 |
| ERVB7_3-LTR_MM | 2141.475 | 1734.193 | 1836.772 |
| ERVB7_4-LTR_MM | 1573.755 | 1153.853 | 1084.549 |
| ERVL-B4-int | 12567.08 | 11982.32 | 13515.32 |
| ERVL-E-int | 6956.663 | 8773.378 | 7643.485 |
| ERVL-int | 5781.565 | 6800.221 | 6507.291 |
| ETnERV-int | 22059.7 | 19192.19 | 19865.43 |
| ETnERV2-int | 26868.62 | 24030.64 | 27627.92 |
| ETnERV3-int | 18826.61 | 16686.49 | 18244.22 |
| Eulor1 | 187.8487 | 304.9631 | 381.7252 |
| Eulor10 | 118.9708 | 268.5496 | 139.2174 |
| Eulor11 | 373.6102 | 275.3771 | 235.7714 |
| Eulor12 | 12.52325 | 145.654 | 177.3899 |
| Eulor2A | 87.66272 | 248.067 | 107.7812 |
| Eulor2B | 162.8022 | 259.4462 | 325.5891 |
| Eulor2C | 106.4476 | 111.5163 | 110.0267 |
| Eulor3 | 37.56974 | 143.3782 | 136.972 |
| Eulor4 | 283.8602 | 236.6878 | 231.2805 |
| Eulor5A | 605.2902 | 391.4451 | 664.6509 |
| Eulor5B | 93.92434 | 127.4473 | 184.1263 |
| Eulor6A | 4.174415 | 56.8961 | 103.2903 |
| Eulor6B | 632.4239 | 248.067 | 397.4433 |
| Eulor6C | 68.87785 | 61.44778 | 47.15428 |
| Eulor6D | 73.05227 | 382.3418 | 229.0351 |
| Eulor6E | 6.261623 | 113.7922 | 51.64517 |
| Eulor7 | 0 | 0 | 33.68163 |
| Eulor8 | 778.5284 | 896.6825 | 790.3956 |
| Eulor9A | 114.7964 | 327.7215 | 437.8612 |
| Eulor9B | 10.43604 | 0 | 33.68163 |
| Eulor9C | 52.18019 | 236.6878 | 85.3268 |
| FAM | 283.8602 | 223.0327 | 294.1529 |
| FordPrefect | 3176.73 | 2560.324 | 2988.683 |
| FordPrefect_a | 421.6159 | 318.6181 | 395.1978 |
| G-rich | 28354.72 | 28391.15 | 27446.04 |
| GA-rich | 57515.09 | 60425.93 | 56140.54 |
| GC_rich | 9311.033 | 9626.82 | 9978.745 |
| GSAT_MM | 3809765 | 4397558 | 6012369 |
| HAL1 | 25687.26 | 23780.29 | 23763.51 |
| HAL1_SS | 1070.738 | 1281.3 | 1286.638 |

| | | | |
|---|---|---|---|
| HAL1b | 5305.682 | 4833.892 | 4731.147 |
| HAL1M8 | 5036.432 | 4208.035 | 4782.792 |
| HAL1ME | 6497.477 | 5141.131 | 6792.462 |
| hAT-N1_Mam | 1870.138 | 1363.23 | 1461.783 |
| Helitron1Na_Mam | 321.43 | 421.0311 | 496.2427 |
| Helitron1Nb_Mam | 699.2146 | 730.5459 | 633.2147 |
| Helitron2Na_Mam | 809.8366 | 418.7553 | 444.5975 |
| Helitron3Na_Mam | 1273.197 | 1017.302 | 1493.219 |
| HERV16-int | 2022.504 | 1543.022 | 1407.892 |
| HERVL32-int | 0 | 0 | 0 |
| HERVL40-int | 1498.615 | 1026.406 | 1059.849 |
| HERVL74-int | 1584.191 | 1388.265 | 1077.812 |
| HUERS-P3b-int | 0 | 0 | 8.981768 |
| HY1 | 70.96506 | 343.6524 | 80.83592 |
| HY3 | 22.95928 | 0 | 24.69986 |
| HY4 | 60.52902 | 0 | 22.45442 |
| HY5 | 0 | 0 | 0 |
| IAP-d-int | 9770.219 | 9986.403 | 10196.55 |
| IAP1-MM_I-int | 8643.127 | 7237.183 | 7798.42 |
| IAP1-MM_LTR | 2373.155 | 1756.951 | 1899.644 |
| IAPA_MM-int | 235.8545 | 393.721 | 220.0533 |
| IAPEy-int | 9430.004 | 11643.22 | 10021.41 |
| IAPEY_LTR | 3994.915 | 3099.699 | 3022.365 |
| IAPEY2_LTR | 6545.483 | 5587.197 | 6381.546 |
| IAPEY3-int | 19891.09 | 19820.32 | 18091.53 |
| IAPEY3_LTR | 3009.753 | 2569.428 | 2986.438 |
| IAPEY3C_LTR | 1966.15 | 1599.918 | 2187.061 |
| IAPEY4_I-int | 8922.813 | 9287.719 | 9327.567 |
| IAPEY4_LTR | 442.488 | 755.5802 | 705.0688 |
| IAPEY5_I-int | 1817.958 | 1891.226 | 1648.155 |
| IAPEY5_LTR | 1448.522 | 1345.024 | 1354.002 |
| IAPEz-int | 154943.9 | 167299.6 | 154490.9 |
| IAPLTR1_Mm | 5848.356 | 5830.712 | 6563.427 |
| IAPLTR1a_Mm | 8649.388 | 8602.69 | 8909.914 |
| IAPLTR2_Mm | 7219.651 | 8254.486 | 7569.385 |
| IAPLTR2a | 3627.567 | 3035.976 | 3377.145 |
| IAPLTR2a2_Mm | 9768.132 | 11244.94 | 10436.81 |
| IAPLTR2b | 5837.92 | 5072.856 | 5752.823 |
| IAPLTR3 | 394.4822 | 1370.058 | 1252.957 |
| IAPLTR3-int | 10371.33 | 9533.51 | 10254.93 |
| IAPLTR4 | 2101.818 | 2546.669 | 1839.017 |
| IAPLTR4_I | 1943.19 | 2371.429 | 1749.199 |

| | | | |
|---|---|---|---|
| ID | 19333.8 | 17951.86 | 17545.88 |
| ID_B1 | 444132.7 | 444604.3 | 460286.4 |
| ID2 | 16273.96 | 16306.42 | 16367.03 |
| ID4 | 69956.94 | 75002.71 | 76293.39 |
| ID4_ | 72526.29 | 79329.09 | 80447.45 |
| IMPB_01 | 63983.35 | 76602.63 | 70003.9 |
| Kanga1 | 2972.184 | 2407.843 | 1892.908 |
| Kanga11a | 999.7724 | 1470.195 | 1277.657 |
| Kanga1a | 1277.371 | 1283.576 | 1120.476 |
| Kanga1b | 945.5051 | 780.6144 | 918.3858 |
| Kanga1c | 1676.028 | 1192.542 | 1250.711 |
| Kanga1d | 953.8539 | 700.9599 | 583.8149 |
| Kanga2_a | 2865.736 | 3352.318 | 3015.629 |
| L1_Mm | 45807.95 | 42323.87 | 45638.61 |
| L1_Mur1 | 111914 | 96327.37 | 105302.3 |
| L1_Mur2 | 281032.1 | 254616.9 | 269450.8 |
| L1_Mur3 | 362723.3 | 318083.3 | 348427.5 |
| L1_Mus1 | 435297.6 | 408691.5 | 431733.4 |
| L1_Mus2 | 336674.9 | 302061.4 | 330565 |
| L1_Mus3 | 707951.6 | 664815 | 708243.9 |
| L1_Mus4 | 200134 | 181191.3 | 190153 |
| L1_Rod | 130949.3 | 112119.4 | 124922.9 |
| L1M | 2423.248 | 1950.398 | 1466.274 |
| L1M1 | 3681.834 | 3980.451 | 4021.587 |
| L1M2 | 172589.1 | 155904.4 | 169984.5 |
| L1M2a | 1617.586 | 2039.156 | 1594.264 |
| L1M2a1 | 0 | 0 | 0 |
| L1M2b | 35.48253 | 11.37922 | 47.15428 |
| L1M2c | 2984.707 | 3566.247 | 3251.4 |
| L1M3 | 54670.23 | 51597.93 | 56998.3 |
| L1M3a | 891.2377 | 587.1677 | 992.4854 |
| L1M3b | 1423.476 | 1078.75 | 1163.139 |
| L1M3c | 2911.655 | 2385.084 | 3704.979 |
| L1M3d | 1730.295 | 1121.991 | 1131.703 |
| L1M3de | 1279.458 | 1392.816 | 1212.539 |
| L1M3e | 4944.595 | 4874.858 | 4630.102 |
| L1M3f | 862.0167 | 748.7526 | 929.613 |
| L1M4 | 116422.4 | 106812.2 | 112386.6 |
| L1M4a1 | 5514.403 | 5257.199 | 4035.059 |
| L1M4a2 | 5051.042 | 5689.61 | 5770.786 |
| L1M4b | 19231.53 | 18539.02 | 21210.45 |
| L1M4c | 19206.48 | 18543.58 | 19683.55 |

| | | | |
|---|---|---|---|
| L1M5 | 130690.5 | 120424 | 128618.9 |
| L1M6 | 2179.045 | 2724.185 | 2092.752 |
| L1M6B | 210.808 | 184.3434 | 388.4615 |
| L1M7 | 1221.016 | 2173.431 | 1398.91 |
| L1M8 | 2352.283 | 1779.71 | 1702.045 |
| L1MA10 | 6267.884 | 5858.022 | 5719.141 |
| L1MA4 | 69547.85 | 68209.32 | 69294.34 |
| L1MA4A | 26540.93 | 23350.16 | 25297.15 |
| L1MA5 | 43724.91 | 40341.61 | 40554.93 |
| L1MA5A | 12832.15 | 11322.32 | 12096.2 |
| L1MA6 | 62002.59 | 60177.86 | 61511.64 |
| L1MA7 | 28116.77 | 28020.19 | 28966.2 |
| L1MA8 | 36793.3 | 34347.04 | 33171.92 |
| L1MA9 | 40299.8 | 37323.84 | 37220.45 |
| L1MB1 | 22059.7 | 19360.6 | 20920.78 |
| L1MB2 | 24092.64 | 21643.28 | 22869.83 |
| L1MB3 | 40907.18 | 41832.29 | 41983.03 |
| L1MB4 | 23082.43 | 20644.18 | 21792.02 |
| L1MB5 | 29421.28 | 29287.83 | 31119.58 |
| L1MB7 | 60090.71 | 61582.06 | 59746.72 |
| L1MB8 | 52712.43 | 48773.61 | 47830.16 |
| L1MC | 46062.58 | 41566.01 | 44311.55 |
| L1MC1 | 52457.79 | 47763.13 | 49628.76 |
| L1MC2 | 21272.82 | 19724.74 | 19941.77 |
| L1MC3 | 48118.48 | 47241.97 | 47163.27 |
| L1MC4 | 43593.42 | 43782.68 | 42528.67 |
| L1MC4a | 25000.57 | 24287.81 | 23485.08 |
| L1MC5 | 27081.52 | 27180.4 | 26727.5 |
| L1MC5a | 19064.55 | 17624.13 | 18673.1 |
| L1MCa | 29686.35 | 31179.06 | 29576.96 |
| L1MCb | 6866.913 | 5623.61 | 5631.569 |
| L1MCc | 3938.561 | 2863.012 | 4578.456 |
| L1MD | 45916.48 | 43844.13 | 42349.04 |
| L1Md_A | 168093.3 | 155139.7 | 165704.6 |
| L1Md_F | 89056.97 | 81391 | 86894.12 |
| L1Md_F2 | 1172124 | 1083634 | 1151602 |
| L1Md_F3 | 199232.3 | 188797.2 | 199640 |
| L1Md_Gf | 14954.84 | 12772.04 | 12513.85 |
| L1Md_T | 306942.7 | 291704 | 306529.8 |
| L1MD1 | 20323.14 | 19649.64 | 21540.53 |
| L1MD2 | 26883.23 | 25348.35 | 24744.77 |
| L1MD3 | 16369.97 | 15407.46 | 16142.48 |

| | | | |
|---|---|---|---|
| L1MDa | 28511.26 | 28243.22 | 29035.81 |
| L1MDb | 3871.77 | 4922.65 | 4508.848 |
| L1ME1 | 50287.09 | 49226.5 | 49927.41 |
| L1ME2 | 24384.85 | 22018.79 | 23718.6 |
| L1ME2z | 11185.35 | 11745.63 | 12253.38 |
| L1ME3 | 11542.26 | 11308.67 | 13158.29 |
| L1ME3A | 16457.63 | 16101.6 | 16200.86 |
| L1ME3B | 4998.862 | 5377.819 | 5914.495 |
| L1ME3C | 2467.079 | 3299.974 | 2732.703 |
| L1ME3Cz | 6445.297 | 7594.491 | 7286.46 |
| L1ME3D | 2807.294 | 2410.119 | 2649.622 |
| L1ME3E | 2988.881 | 2325.912 | 2382.414 |
| L1ME3F | 3095.329 | 2548.945 | 2928.057 |
| L1ME3G | 8966.644 | 7592.215 | 7719.83 |
| L1ME4a | 12107.89 | 13024.65 | 13079.7 |
| L1ME4b | 12969.91 | 13764.3 | 12253.38 |
| L1ME4c | 6436.948 | 6313.191 | 6410.737 |
| L1ME5 | 1433.912 | 1335.92 | 1041.885 |
| L1MEa | 1425.563 | 908.0617 | 1129.457 |
| L1MEb | 2805.207 | 2410.119 | 2184.815 |
| L1MEc | 47542.42 | 44567.85 | 46339.19 |
| L1MEd | 24756.37 | 21782.1 | 25467.8 |
| L1MEf | 29156.2 | 24640.56 | 27201.29 |
| L1MEg | 28617.7 | 27341.99 | 27735.7 |
| L1MEg1 | 615.7262 | 605.3745 | 758.9594 |
| L1MEg2 | 703.389 | 782.8903 | 651.1782 |
| L1MEh | 1498.615 | 1160.68 | 1295.62 |
| L1MEi | 3404.236 | 3206.664 | 3343.463 |
| L1MEj | 1047.778 | 1208.473 | 1446.065 |
| L1P5 | 795.2261 | 653.1672 | 601.7785 |
| L1PB4 | 1206.406 | 1665.918 | 1585.282 |
| L1VL1 | 42128.2 | 38759.9 | 39818.42 |
| L1VL2 | 112819.8 | 97783.91 | 109440.6 |
| L1VL4 | 167354.4 | 137187.9 | 155880.8 |
| L2 | 103448.3 | 103498.6 | 104038.1 |
| L2a | 133894.4 | 134568.4 | 137311 |
| L2b | 59481.24 | 62501.5 | 63988.36 |
| L2c | 59464.55 | 62342.19 | 58262.49 |
| L3 | 44634.93 | 47000.73 | 49754.51 |
| L3b | 9907.975 | 8928.135 | 9442.084 |
| L4_A_Mam | 4324.694 | 3980.451 | 3862.16 |
| L4_B_Mam | 2312.626 | 1763.779 | 2362.205 |

| | | | |
|---|---|---|---|
| L4_C_Mam | 3627.567 | 3450.179 | 4019.341 |
| L5 | 626.1623 | 516.6166 | 893.686 |
| LFSINE_Vert | 5230.542 | 4253.552 | 4270.831 |
| Looper | 1780.388 | 2080.121 | 1490.974 |
| LSU-rRNA_Hsa | 17772.57 | 14164.85 | 10117.96 |
| LTR10_RN | 4337.217 | 3040.527 | 3473.699 |
| LTR101_Mam | 960.1155 | 1670.469 | 999.2217 |
| LTR102_Mam | 1655.156 | 812.4763 | 1940.062 |
| LTR103_Mam | 202.4591 | 375.5142 | 159.4264 |
| LTR103b_Mam | 323.5172 | 348.2041 | 289.662 |
| LTR104_Mam | 1110.394 | 1791.089 | 1320.32 |
| LTR105_Mam | 392.395 | 976.337 | 604.0239 |
| LTR106_Mam | 1179.272 | 1226.68 | 1203.557 |
| LTR107_Mam | 521.8019 | 741.9251 | 572.5877 |
| LTR108a_Mam | 175.3254 | 159.3091 | 166.1627 |
| LTR108b_Mam | 75.13947 | 109.2405 | 148.1992 |
| LTR108c_Mam | 114.7964 | 270.8254 | 33.68163 |
| LTR108d_Mam | 198.2847 | 163.8608 | 240.2623 |
| LTR108e_Mam | 185.7615 | 200.2743 | 422.1431 |
| LTR109A2 | 158.6278 | 172.9641 | 251.4895 |
| LTR16A | 8974.993 | 8695.999 | 8348.554 |
| LTR16A1 | 3775.759 | 3750.591 | 3660.071 |
| LTR16A2 | 2594.399 | 1975.432 | 2265.651 |
| LTR16B | 1911.882 | 1313.162 | 1394.42 |
| LTR16B1 | 1619.673 | 1918.536 | 1659.382 |
| LTR16B2 | 2469.167 | 2059.639 | 2297.087 |
| LTR16C | 7973.133 | 7721.938 | 7861.293 |
| LTR16D | 507.1915 | 1258.542 | 736.505 |
| LTR16D1 | 519.7147 | 446.0654 | 666.8963 |
| LTR16D2 | 381.959 | 79.65453 | 294.1529 |
| LTR16E1 | 2792.684 | 2960.873 | 2663.094 |
| LTR16E2 | 2162.347 | 2824.322 | 2191.551 |
| LTR23-int | 8.34883 | 43.24103 | 103.2903 |
| LTR27C | 0 | 31.86181 | 17.96354 |
| LTR28 | 18.78487 | 0 | 26.94531 |
| LTR28B | 423.7031 | 334.549 | 383.9706 |
| LTR29 | 262.9882 | 177.5158 | 105.5358 |
| LTR31 | 1680.202 | 814.7521 | 1028.412 |
| LTR32 | 0 | 61.44778 | 2.245442 |
| LTR33 | 7603.697 | 7660.49 | 8278.945 |
| LTR33A | 3362.491 | 3466.11 | 2795.575 |
| LTR33A_ | 2506.736 | 2753.771 | 2254.424 |

| | | | |
|---|---|---|---|
| LTR33B | 1195.97 | 1929.916 | 1129.457 |
| LTR33C | 1588.365 | 1065.095 | 1486.483 |
| LTR34 | 0 | 104.6888 | 78.59047 |
| LTR36 | 18.78487 | 56.8961 | 17.96354 |
| LTR37-int | 2398.202 | 2467.015 | 1978.234 |
| LTR37A | 3711.055 | 4192.104 | 3321.009 |
| LTR37B | 1824.219 | 2496.601 | 2613.695 |
| LTR39 | 4.174415 | 63.72363 | 15.71809 |
| LTR40a | 4763.008 | 5022.787 | 5063.472 |
| LTR40A1 | 949.6795 | 1046.888 | 574.8332 |
| LTR40b | 3775.759 | 3566.247 | 3691.507 |
| LTR40c | 1534.098 | 2068.742 | 1852.49 |
| LTR41 | 3032.713 | 2831.15 | 3067.274 |
| LTR41B | 1924.405 | 1811.572 | 1571.809 |
| LTR44 | 0 | 0 | 17.96354 |
| LTR45 | 4.174415 | 175.24 | 168.4082 |
| LTR45C | 2.087208 | 97.86129 | 87.57224 |
| LTR48 | 2890.783 | 3161.147 | 3273.855 |
| LTR48B | 1803.347 | 1820.675 | 1978.234 |
| LTR50 | 2677.887 | 3202.112 | 2557.559 |
| LTR52 | 1636.371 | 1570.332 | 1261.938 |
| LTR52-int | 70.96506 | 762.4077 | 415.4068 |
| LTR53 | 1206.406 | 1158.405 | 1396.665 |
| LTR53B | 703.389 | 771.5111 | 525.4335 |
| LTR55 | 1248.15 | 987.7162 | 965.5401 |
| LTR58 | 45.91857 | 2.275844 | 83.08136 |
| LTR59 | 0 | 0 | 0 |
| LTR64 | 54.2674 | 81.93038 | 40.41796 |
| LTR65 | 711.7378 | 696.4082 | 1084.549 |
| LTR67B | 3036.887 | 3416.042 | 3285.082 |
| LTR68 | 573.9821 | 685.029 | 502.979 |
| LTR69 | 774.354 | 539.375 | 509.7154 |
| LTR72_RN | 5113.659 | 6315.467 | 6222.12 |
| LTR73 | 872.4528 | 935.3718 | 815.0955 |
| LTR75 | 384.0462 | 225.3085 | 709.5597 |
| LTR75_1 | 6.261623 | 0 | 44.90884 |
| LTR75B | 538.4996 | 241.2394 | 240.2623 |
| LTR77 | 102.2732 | 195.7226 | 103.2903 |
| LTR78 | 2692.498 | 3329.56 | 3202 |
| LTR78B | 2416.986 | 2023.225 | 2683.303 |
| LTR79 | 2611.097 | 2216.672 | 2119.697 |
| LTR80A | 576.0693 | 832.9589 | 866.7407 |

| | | | |
|---|---|---|---|
| LTR80B | 467.5345 | 539.375 | 725.2778 |
| LTR81 | 325.6044 | 316.3423 | 330.08 |
| LTR81A | 1154.226 | 992.2679 | 1261.938 |
| LTR81B | 1304.505 | 1474.747 | 1205.802 |
| LTR81C | 411.1799 | 366.4109 | 671.3872 |
| LTR82A | 2594.399 | 1895.778 | 2139.906 |
| LTR82B | 1164.662 | 1752.4 | 1827.79 |
| LTR83 | 1028.993 | 1201.646 | 909.4041 |
| LTR84a | 1039.429 | 1033.233 | 857.7589 |
| LTR84b | 1615.499 | 1572.608 | 1255.202 |
| LTR85a | 1605.063 | 896.6825 | 1320.32 |
| LTR85b | 425.7904 | 960.4061 | 705.0688 |
| LTR85c | 903.7609 | 327.7215 | 332.3254 |
| LTR86A1 | 592.767 | 721.4425 | 471.5428 |
| LTR86A2 | 242.1161 | 452.8929 | 273.9439 |
| LTR86B1 | 298.4707 | 145.654 | 112.2721 |
| LTR86B2 | 233.7673 | 220.7569 | 161.6718 |
| LTR86C | 333.9532 | 482.4789 | 388.4615 |
| LTR87 | 1279.458 | 914.8892 | 1232.748 |
| LTR88a | 271.337 | 166.1366 | 170.6536 |
| LTR88b | 932.9818 | 1012.751 | 808.3592 |
| LTR88c | 569.8077 | 746.4768 | 660.16 |
| LTR89 | 507.1915 | 714.615 | 669.1417 |
| LTR9 | 0 | 27.31013 | 166.1627 |
| LTR90A | 429.9648 | 461.9963 | 487.2609 |
| LTR90B | 599.0286 | 562.1334 | 410.9159 |
| LTR91 | 269.2498 | 334.549 | 215.5624 |
| LTR9A1 | 4.174415 | 40.96519 | 26.94531 |
| LTR9D | 73.05227 | 25.03428 | 20.20898 |
| LTRIS_Mm | 4333.043 | 4201.208 | 3994.642 |
| LTRIS_Mus | 7340.709 | 8072.418 | 7205.624 |
| LTRIS2 | 12251.91 | 10298.19 | 11602.2 |
| LTRIS3 | 1794.999 | 1360.955 | 1708.781 |
| LTRIS4 | 995.598 | 666.8222 | 1010.449 |
| LTRIS4A | 1630.109 | 949.0269 | 1385.438 |
| LTRIS4B | 484.2322 | 248.067 | 514.2062 |
| LTRIS5 | 1106.22 | 923.9926 | 792.6411 |
| LTRIS6 | 521.8019 | 1099.233 | 794.8865 |
| Lx | 356747.6 | 331706.5 | 352031.4 |
| Lx10 | 84477.64 | 84208.5 | 84114.26 |
| Lx2 | 217336.8 | 206933.4 | 223841.4 |
| Lx2A | 20204.17 | 18127.1 | 16932.88 |

| | | | |
|---|---|---|---|
| Lx2A1 | 27830.83 | 26502.2 | 27293.35 |
| Lx2B | 84429.64 | 76670.9 | 79533.56 |
| Lx2B2 | 138901.6 | 129821 | 138094.7 |
| Lx3_Mus | 185189.6 | 170094.3 | 182132.3 |
| Lx3A | 113598.4 | 104927.8 | 111299.8 |
| Lx3B | 71088.2 | 65328.1 | 68750.95 |
| Lx3C | 84400.41 | 77770.14 | 81599.37 |
| Lx4A | 133322.5 | 126361.7 | 131679.5 |
| Lx4B | 65143.84 | 63402.73 | 65912.71 |
| Lx5 | 211686.7 | 214357.2 | 213925.5 |
| Lx5b | 92550.96 | 94713.79 | 95817.51 |
| Lx5c | 181918.9 | 173230.4 | 182511.8 |
| Lx6 | 225299.5 | 216717.2 | 230400.3 |
| Lx7 | 439639 | 413634.6 | 415007.1 |
| Lx8 | 414999.5 | 414067 | 420903.6 |
| Lx8b | 221141.7 | 215750 | 224346.6 |
| Lx9 | 457791.4 | 449215.2 | 451232.8 |
| MADE2 | 749.3075 | 596.2711 | 675.8781 |
| Mam_R4 | 862.0167 | 782.8903 | 608.5148 |
| MamGyp-int | 221.244 | 213.9293 | 370.4979 |
| MamGypLTR1a | 855.7551 | 762.4077 | 606.2694 |
| MamGypLTR1b | 640.7727 | 769.2352 | 480.5246 |
| MamGypLTR1c | 1185.534 | 694.1324 | 747.7322 |
| MamGypLTR1d | 471.7089 | 571.2368 | 752.2231 |
| MamGypLTR2b | 110.622 | 448.3412 | 669.1417 |
| MamGypLTR2c | 603.203 | 655.443 | 651.1782 |
| MamGypLTR3 | 156.5406 | 477.9272 | 334.5709 |
| MamGypLTR3a | 1552.882 | 985.4404 | 669.1417 |
| MamRep1151 | 2458.731 | 3475.214 | 3094.219 |
| MamRep1161 | 31.30811 | 34.13766 | 17.96354 |
| MamRep137 | 1961.975 | 2100.604 | 1504.446 |
| MamRep1527 | 2385.678 | 2034.604 | 1821.054 |
| MamRep1879 | 434.1392 | 339.1007 | 473.7883 |
| MamRep1894 | 102.2732 | 179.7917 | 175.1445 |
| MamRep38 | 951.7667 | 857.9931 | 635.4601 |
| MamRep4096 | 920.4586 | 1684.124 | 1145.175 |
| MamRep434 | 3788.282 | 3934.934 | 3329.991 |
| MamRep488 | 43.83136 | 134.2748 | 211.0716 |
| MamRep564 | 749.3075 | 741.9251 | 586.0604 |
| MamRep605 | 4266.252 | 3429.697 | 3770.097 |
| MamSINE1 | 1959.888 | 2200.741 | 1623.455 |
| MamTip1 | 192.0231 | 546.2025 | 374.9888 |

| | | | |
|---|---|---|---|
| MamTip2 | 2742.591 | 3864.383 | 3592.707 |
| MamTip3 | 960.1155 | 1331.369 | 830.8136 |
| MARINER1_EC | 2.087208 | 0 | 0 |
| MARNA | 3239.346 | 3083.768 | 3332.236 |
| MER101B | 12.52325 | 157.0332 | 62.87238 |
| MER102a | 2464.992 | 1986.812 | 1996.198 |
| MER102b | 2581.876 | 3356.87 | 2791.085 |
| MER102c | 1974.498 | 2159.776 | 1973.744 |
| MER103C | 5090.699 | 6411.052 | 6199.666 |
| MER104 | 2262.533 | 2134.742 | 1641.418 |
| MER105 | 870.3656 | 901.2342 | 595.0422 |
| MER106A | 594.8542 | 760.1318 | 833.059 |
| MER106B | 1060.301 | 700.9599 | 1071.076 |
| MER110 | 1287.807 | 778.3386 | 886.9496 |
| MER110-int | 123.1452 | 234.4119 | 368.2525 |
| MER110A | 820.2726 | 992.2679 | 1396.665 |
| MER112 | 2423.248 | 2118.811 | 1821.054 |
| MER113 | 3581.648 | 5204.855 | 4708.692 |
| MER113A | 924.633 | 873.924 | 873.477 |
| MER113B | 571.8949 | 343.6524 | 583.8149 |
| MER115 | 3109.939 | 2837.977 | 3206.491 |
| MER117 | 2932.527 | 3174.802 | 3305.291 |
| MER119 | 3842.549 | 3350.042 | 2970.72 |
| MER121 | 4226.595 | 3391.007 | 4365.139 |
| MER123 | 377.7846 | 279.9288 | 202.0898 |
| MER124 | 1682.289 | 1390.541 | 1717.763 |
| MER125 | 473.7961 | 523.4441 | 543.397 |
| MER126 | 557.2844 | 855.7173 | 624.2329 |
| MER127 | 978.9004 | 748.7526 | 716.296 |
| MER129 | 484.2322 | 341.3766 | 381.7252 |
| MER130 | 348.5637 | 131.9989 | 532.1698 |
| MER131 | 1181.36 | 1244.887 | 873.477 |
| MER132 | 0 | 143.3782 | 67.36326 |
| MER133A | 35.48253 | 106.9647 | 125.7448 |
| MER133B | 20.87208 | 50.06856 | 6.736326 |
| MER134 | 283.8602 | 323.1698 | 327.8345 |
| MER135 | 1277.371 | 755.5802 | 1120.476 |
| MER136 | 231.68 | 79.65453 | 213.317 |
| MER2 | 61176.06 | 61666.27 | 60049.86 |
| MER20 | 27311.11 | 30152.66 | 29731.9 |
| MER20B | 5821.222 | 6083.331 | 5018.563 |
| MER21-int | 381.959 | 712.3391 | 763.4503 |

| | | | |
|---|---|---|---|
| MER21B | 11830.29 | 12535.35 | 11952.49 |
| MER21C | 15908.7 | 16461.18 | 16717.32 |
| MER2B | 4896.589 | 4881.685 | 4472.921 |
| MER3 | 15528.82 | 15232.22 | 16173.92 |
| MER31-int | 1131.267 | 908.0617 | 1517.919 |
| MER31A | 4464.537 | 4576.722 | 4798.51 |
| MER31B | 4356.002 | 3570.799 | 3621.898 |
| MER33 | 17200.68 | 18523.09 | 17846.77 |
| MER34 | 4009.526 | 4105.622 | 4385.348 |
| MER34-int | 824.447 | 452.8929 | 343.5526 |
| MER34A | 4322.607 | 4462.93 | 4008.114 |
| MER34A1 | 3487.724 | 3695.97 | 3114.428 |
| MER34B-int | 467.5345 | 710.0633 | 502.979 |
| MER44A | 5435.089 | 4808.858 | 5146.553 |
| MER44B | 11897.08 | 10710.12 | 11004.91 |
| MER44C | 1989.109 | 2075.57 | 1562.828 |
| MER44D | 5885.925 | 5680.506 | 4901.8 |
| MER45A | 3623.392 | 3682.315 | 3008.892 |
| MER45B | 4656.56 | 5070.58 | 4578.456 |
| MER45C | 1893.097 | 1847.985 | 1601 |
| MER45R | 2122.69 | 2007.294 | 2047.843 |
| MER46C | 4036.66 | 5072.856 | 5341.907 |
| MER47A | 749.3075 | 805.6487 | 790.3956 |
| MER47B | 2947.137 | 3404.662 | 2950.511 |
| MER47C | 62.61623 | 13.65506 | 47.15428 |
| MER49 | 0 | 0 | 6.736326 |
| MER4B-int | 417.4415 | 225.3085 | 298.6438 |
| MER4CL34 | 0 | 2.275844 | 0 |
| MER50 | 118.9708 | 77.37869 | 56.13605 |
| MER50-int | 52.18019 | 47.79272 | 71.85415 |
| MER50B | 22.95928 | 91.03375 | 150.4446 |
| MER51-int | 81.4011 | 45.51688 | 94.30857 |
| MER52-int | 839.0575 | 953.5786 | 1329.302 |
| MER53 | 7532.732 | 8513.932 | 8099.31 |
| MER54A | 1415.127 | 1970.881 | 2101.734 |
| MER54B | 592.767 | 955.8544 | 900.4223 |
| MER57-int | 1183.447 | 846.6139 | 788.1502 |
| MER57C2 | 2563.091 | 2974.528 | 2984.193 |
| MER57D | 1630.109 | 2014.122 | 2267.897 |
| MER57E1 | 1767.865 | 1142.474 | 1113.739 |
| MER57E2 | 467.5345 | 600.8228 | 572.5877 |
| MER57E3 | 0 | 9.103375 | 42.6634 |

| | | | |
|---|---|---|---|
| MER58A | 18693.03 | 19276.4 | 22225.39 |
| MER58B | 17135.97 | 15892.22 | 17220.3 |
| MER58C | 2057.987 | 2030.053 | 2142.152 |
| MER58D | 1202.232 | 1395.092 | 1639.173 |
| MER5A | 33787.72 | 34868.2 | 36052.82 |
| MER5A1 | 15639.45 | 17578.62 | 16001.02 |
| MER5B | 21366.74 | 18928.19 | 20049.55 |
| MER5C | 1740.731 | 1404.196 | 1398.91 |
| MER5C1 | 252.5521 | 461.9963 | 350.289 |
| MER63A | 2949.224 | 2642.255 | 3419.808 |
| MER63B | 4007.439 | 3525.282 | 3298.554 |
| MER63C | 3807.067 | 3880.314 | 3902.578 |
| MER63D | 5975.675 | 5498.439 | 5636.06 |
| MER65-int | 300.5579 | 145.654 | 282.9257 |
| MER65D | 753.482 | 484.7547 | 630.9692 |
| MER66-int | 37.56974 | 45.51688 | 154.9355 |
| MER66A | 60.52902 | 72.827 | 26.94531 |
| MER66B | 62.61623 | 0 | 17.96354 |
| MER66C | 62.61623 | 47.79272 | 38.17252 |
| MER67A | 845.3191 | 1028.681 | 767.9412 |
| MER67B | 1277.371 | 1213.025 | 1845.753 |
| MER67C | 4846.496 | 4137.484 | 4791.773 |
| MER67D | 2602.748 | 3431.973 | 2786.594 |
| MER68 | 4754.659 | 4524.378 | 4771.564 |
| MER68-int | 563.5461 | 455.1688 | 354.7799 |
| MER68B | 1465.22 | 1791.089 | 1556.091 |
| MER68C | 137.7557 | 111.5163 | 505.2245 |
| MER70-int | 419.5287 | 370.9625 | 487.2609 |
| MER70A | 1125.005 | 1627.228 | 1362.983 |
| MER70B | 2014.155 | 1820.675 | 1434.838 |
| MER72 | 0 | 27.31013 | 58.38149 |
| MER72B | 296.3835 | 268.5496 | 114.5175 |
| MER73 | 1022.732 | 835.2347 | 1138.439 |
| MER74A | 4404.008 | 5195.752 | 4589.684 |
| MER74B | 1590.452 | 2164.328 | 2101.734 |
| MER74C | 768.0924 | 555.3059 | 844.2862 |
| MER76 | 1686.464 | 1217.576 | 1439.328 |
| MER76-int | 137.7557 | 34.13766 | 71.85415 |
| MER77 | 3496.073 | 4264.931 | 4091.196 |
| MER77B | 3742.363 | 3623.143 | 4070.987 |
| MER81 | 1797.086 | 2107.431 | 2016.407 |
| MER83A-int | 0 | 0 | 0 |

| | | | |
|---|---|---|---|
| MER83B-int | 0 | 34.13766 | 13.47265 |
| MER87 | 144.0173 | 275.3771 | 127.9902 |
| MER88 | 473.7961 | 395.9968 | 253.735 |
| MER89 | 4159.805 | 4690.514 | 3850.933 |
| MER89-int | 503.017 | 607.6503 | 586.0604 |
| MER90 | 1874.312 | 1483.85 | 1493.219 |
| MER90a | 3456.416 | 3325.008 | 3307.536 |
| MER91A | 1838.83 | 1408.747 | 1412.383 |
| MER91B | 968.4643 | 780.6144 | 736.505 |
| MER91C | 279.6858 | 225.3085 | 455.8247 |
| MER92-int | 2323.062 | 2567.152 | 2121.943 |
| MER92A | 294.2963 | 68.27532 | 161.6718 |
| MER92B | 1598.801 | 1624.953 | 1627.946 |
| MER92C | 457.0985 | 427.8586 | 325.5891 |
| MER92D | 400.7439 | 405.1002 | 684.8598 |
| MER94 | 2172.783 | 2198.465 | 2092.752 |
| MER94B | 590.6798 | 443.7896 | 345.7981 |
| MER95 | 171.151 | 234.4119 | 161.6718 |
| MER96 | 262.9882 | 245.7911 | 422.1431 |
| MER96B | 2567.265 | 1615.849 | 2276.878 |
| MER97a | 388.2206 | 694.1324 | 464.8065 |
| MER97b | 404.9183 | 355.0316 | 559.1151 |
| MER97c | 782.7029 | 634.9604 | 489.5064 |
| MER97d | 204.5463 | 159.3091 | 246.9986 |
| MER99 | 1897.272 | 1247.162 | 1735.727 |
| MERV1_I-int | 17645.25 | 18554.95 | 16618.52 |
| MERV1_LTR | 4293.386 | 4378.724 | 4506.602 |
| MERVK26-int | 14322.42 | 11112.95 | 12394.84 |
| MERVL-int | 82369.56 | 86743.79 | 82358.33 |
| MERVL_2A-int | 47498.58 | 48662.09 | 47742.59 |
| MERX | 262.9882 | 350.48 | 419.8977 |
| MIR | 191883.3 | 194862.3 | 191118.6 |
| MIR3 | 62695.54 | 64645.34 | 61664.33 |
| MIRb | 198777.3 | 211346.2 | 204788.8 |
| MIRc | 95166.23 | 100521.7 | 94723.98 |
| MLT-int | 6883.611 | 7057.392 | 6291.729 |
| MLT1-int | 2473.341 | 2444.256 | 1603.246 |
| MLT1A | 28202.35 | 30198.17 | 28196.02 |
| MLT1A-int | 882.8888 | 901.2342 | 646.6873 |
| MLT1A0 | 63325.88 | 66063.2 | 61873.16 |
| MLT1A0-int | 1527.836 | 2068.742 | 1708.781 |
| MLT1A1 | 45753.68 | 46950.66 | 44504.66 |

| | | | |
|---|---|---|---|
| MLT1A1-int | 972.6388 | 1015.026 | 1306.847 |
| MLT1B | 57999.33 | 57674.43 | 57588.85 |
| MLT1B-int | 1648.894 | 1260.817 | 1282.147 |
| MLT1C | 49105.73 | 51395.38 | 48225.36 |
| MLT1C-int | 755.5692 | 732.8217 | 788.1502 |
| MLT1D | 49896.79 | 55264.32 | 55298.5 |
| MLT1D-int | 1797.086 | 1174.335 | 1248.466 |
| MLT1E | 3777.846 | 3495.696 | 3370.409 |
| MLT1E-int | 79.31389 | 31.86181 | 22.45442 |
| MLT1E1 | 3723.578 | 3447.903 | 3590.462 |
| MLT1E1-int | 25.04649 | 86.48207 | 40.41796 |
| MLT1E1A | 7111.116 | 7455.664 | 7479.568 |
| MLT1E1A-int | 404.9183 | 350.48 | 179.6354 |
| MLT1E2 | 12477.33 | 11982.32 | 12221.94 |
| MLT1E2-int | 546.8484 | 489.3064 | 339.0618 |
| MLT1E3 | 5837.92 | 7851.661 | 6630.791 |
| MLT1E3-int | 292.2091 | 304.9631 | 132.4811 |
| MLT1F | 10772.08 | 11060.6 | 10836.5 |
| MLT1F-int | 3201.776 | 2853.908 | 3300.8 |
| MLT1F1 | 9544.8 | 8978.204 | 9839.527 |
| MLT1F1-int | 674.1681 | 619.0295 | 749.9777 |
| MLT1F2 | 15484.99 | 16190.35 | 15763 |
| MLT1F2-int | 682.5169 | 691.8565 | 545.6424 |
| MLT1G | 5599.978 | 6704.636 | 5465.406 |
| MLT1G-int | 619.9007 | 277.653 | 366.0071 |
| MLT1G1 | 6109.257 | 6149.33 | 6592.618 |
| MLT1G1-int | 644.9472 | 407.376 | 401.9341 |
| MLT1G3 | 6042.466 | 7348.7 | 7614.294 |
| MLT1G3-int | 707.5634 | 894.4066 | 758.9594 |
| MLT1H | 14199.27 | 13582.24 | 12724.92 |
| MLT1H-int | 3406.323 | 2744.668 | 2306.069 |
| MLT1H1 | 6290.844 | 6461.121 | 6154.757 |
| MLT1H1-int | 705.4762 | 591.7194 | 655.6691 |
| MLT1H2 | 5149.141 | 5407.405 | 4886.082 |
| MLT1H2-int | 732.6099 | 421.0311 | 410.9159 |
| MLT1I | 8599.295 | 9560.82 | 9594.774 |
| MLT1I-int | 463.3601 | 318.6181 | 433.3703 |
| MLT1J | 17486.63 | 18559.51 | 17139.46 |
| MLT1J-int | 3719.404 | 3873.486 | 3554.535 |
| MLT1J1 | 4710.828 | 7255.39 | 6219.875 |
| MLT1J1-int | 429.9648 | 550.7542 | 377.2343 |
| MLT1J2 | 5591.629 | 5634.989 | 5618.096 |

| | | | |
|---|---|---|---|
| MLT1J2-int | 85.57551 | 227.5844 | 350.289 |
| MLT1K | 13773.48 | 15687.39 | 15897.73 |
| MLT1K-int | 31.30811 | 34.13766 | 62.87238 |
| MLT1L | 8198.552 | 9476.614 | 7930.902 |
| MLT1L-int | 118.9708 | 193.4467 | 119.0084 |
| MLT1M | 2003.719 | 1490.678 | 927.3676 |
| MLT1N2 | 3160.032 | 3340.939 | 3484.926 |
| MLT1N2-int | 0 | 18.20675 | 0 |
| MLT1O | 2771.812 | 2685.496 | 3237.928 |
| MLT1O-int | 12.52325 | 0 | 26.94531 |
| MLT2B1 | 10210.62 | 10962.74 | 10151.64 |
| MLT2B2 | 8828.888 | 9094.272 | 8741.506 |
| MLT2B3 | 9734.736 | 9767.922 | 8907.669 |
| MLT2B4 | 14568.71 | 11504.39 | 12264.6 |
| MLT2B5 | 1939.016 | 2751.495 | 2310.56 |
| MLT2C1 | 4639.863 | 5619.058 | 5743.841 |
| MLT2C2 | 4099.276 | 3971.348 | 4106.914 |
| MLT2D | 9672.12 | 8912.205 | 9410.648 |
| MLT2E | 872.4528 | 785.1661 | 314.3619 |
| MLT2F | 3009.753 | 4091.967 | 2986.438 |
| MLTR11A | 15610.23 | 16508.97 | 16519.72 |
| MLTR11B | 20356.54 | 20858.11 | 21113.89 |
| MLTR12 | 3589.997 | 3404.662 | 3828.479 |
| MLTR13 | 1594.627 | 2075.57 | 1688.572 |
| MLTR14 | 45787.07 | 45277.91 | 41935.88 |
| MLTR18_MM | 3963.607 | 3759.694 | 3568.008 |
| MLTR18A_MM | 3485.637 | 3133.837 | 3089.728 |
| MLTR18B_MM | 3892.642 | 4642.721 | 4724.41 |
| MLTR18C_MM | 5251.414 | 5589.473 | 6183.948 |
| MLTR18D_MM | 6357.634 | 7198.494 | 6491.573 |
| MLTR25A | 21379.27 | 20553.15 | 22227.63 |
| MLTR25C | 8050.36 | 6593.12 | 7533.458 |
| MLTR31A_MM | 2984.707 | 3787.004 | 3094.219 |
| MLTR31C_MM | 2323.062 | 1939.019 | 2247.688 |
| MLTR31D_MM | 2586.05 | 3680.04 | 3408.581 |
| MLTR31E_MM | 1632.196 | 1342.748 | 1338.283 |
| MLTR31F_MM | 3875.945 | 4178.449 | 3565.762 |
| MLTR31FA_MM | 5407.955 | 5468.853 | 5256.58 |
| MLTR32C_MM | 1214.755 | 1144.749 | 837.5499 |
| MLTR73 | 5151.228 | 4940.857 | 4951.2 |
| MMAR1 | 3080.718 | 3345.49 | 3695.998 |
| MMERGLN-int | 17238.25 | 18866.75 | 17117.01 |

| | | | |
|---|---|---|---|
| MMERGLN_LTR | 3360.404 | 2960.873 | 3507.381 |
| MMERVK10C-int | 57179.05 | 54538.32 | 53145.12 |
| MMERVK10D3_I-int | 8970.818 | 11115.22 | 10234.73 |
| MMERVK10D3_LTR | 1223.104 | 1137.922 | 958.8038 |
| MMERVK9C_I-int | 34741.57 | 31260.99 | 34847.02 |
| MMERVK9E_I-int | 31516.84 | 31099.41 | 29960.93 |
| MMETn-int | 31320.64 | 29508.59 | 29776.81 |
| MMSAT4 | 11072.64 | 11370.12 | 12477.92 |
| MMTV-int | 515.5403 | 509.789 | 700.5779 |
| MMVL30-int | 12882.25 | 10835.29 | 11494.42 |
| MRLTR33 | 3487.724 | 3429.697 | 3127.901 |
| MT-int | 3214.3 | 2751.495 | 3098.71 |
| MT2_Mm | 19335.89 | 18432.06 | 18453.04 |
| MT2A | 63100.46 | 62071.37 | 61628.4 |
| MT2B | 39393.96 | 39474.51 | 40285.48 |
| MT2B1 | 47319.08 | 47471.83 | 48607.09 |
| MT2B2 | 14771.17 | 18441.16 | 16688.13 |
| MT2C_Mm | 16509.81 | 15029.67 | 16295.17 |
| MTA_Mm | 97238.83 | 99964.17 | 99275.49 |
| MTA_Mm-int | 56413.05 | 49169.61 | 54732.65 |
| MTB | 44215.41 | 47569.69 | 46404.31 |
| MTB-int | 9267.202 | 8459.312 | 8305.89 |
| MTB_Mm | 33712.58 | 34979.72 | 36953.24 |
| MTB_Mm-int | 5867.141 | 5912.642 | 5687.705 |
| MTC | 173478.3 | 182088 | 182195.2 |
| MTC-int | 33176.17 | 34547.31 | 35866.45 |
| MTD | 339457.2 | 341114.9 | 343577.3 |
| MTD-int | 28070.86 | 25635.11 | 27542.59 |
| MTE-int | 45762.03 | 45760.39 | 45452.24 |
| MTE2a | 106810.8 | 108885.5 | 104179.5 |
| MTE2a-int | 4297.56 | 4647.273 | 4362.894 |
| MTE2b | 90879.11 | 93100.22 | 93890.92 |
| MTE2b-int | 8073.319 | 6720.567 | 8301.399 |
| MTEa | 117100.7 | 123239.2 | 118990.5 |
| MTEa-int | 14956.93 | 14492.57 | 16250.26 |
| MTEb | 63267.44 | 60125.52 | 60927.83 |
| MTEb-int | 6599.75 | 6481.603 | 6064.939 |
| MuLV-int | 3721.491 | 4246.725 | 3839.706 |
| MurERV4-int | 18375.78 | 18657.37 | 20426.79 |
| MurERV4_19-int | 17774.66 | 16606.83 | 16075.12 |
| MuRRS-int | 8924.9 | 6643.188 | 7084.37 |
| MuRRS4-int | 11984.75 | 11383.77 | 10809.56 |

| | | | |
|---|---|---|---|
| MurSAT1 | 809.8366 | 550.7542 | 819.5864 |
| MurSatRep1 | 17290.43 | 16736.56 | 15974.08 |
| MURVY-int | 4570.985 | 4087.416 | 3298.554 |
| MURVY-LTR | 4677.432 | 4640.446 | 4371.876 |
| MusHAL1 | 54565.87 | 46454.52 | 56879.29 |
| MYSERV-int | 18797.39 | 18181.72 | 18789.86 |
| MYSERV16_I-int | 40976.06 | 40737.6 | 42227.78 |
| MYSERV6-int | 49521.09 | 49069.47 | 49752.26 |
| nhAT5a_ML | 0 | 31.86181 | 26.94531 |
| ORR1A0 | 13299.69 | 13038.31 | 12751.87 |
| ORR1A0-int | 10110.43 | 9053.307 | 9352.266 |
| ORR1A1 | 24065.5 | 23623.26 | 24980.54 |
| ORR1A1-int | 16405.45 | 14488.02 | 16355.8 |
| ORR1A2 | 93567.43 | 94638.69 | 94463.5 |
| ORR1A2-int | 35405.3 | 35127.65 | 33131.5 |
| ORR1A3 | 26346.82 | 28586.87 | 28999.88 |
| ORR1A3-int | 16983.61 | 16798 | 16164.94 |
| ORR1A4 | 51282.69 | 51641.17 | 53351.7 |
| ORR1A4-int | 23476.91 | 20999.21 | 23480.59 |
| ORR1B1 | 95994.85 | 102822.6 | 105728.9 |
| ORR1B1-int | 34547.46 | 30004.73 | 30798.48 |
| ORR1B2 | 50092.98 | 50548.77 | 51274.67 |
| ORR1B2-int | 11692.54 | 10009.16 | 11575.25 |
| ORR1C1 | 47913.94 | 48511.89 | 49395.24 |
| ORR1C1-int | 9974.765 | 9062.41 | 10865.69 |
| ORR1C2 | 63359.27 | 63277.56 | 62364.91 |
| ORR1C2-int | 11936.74 | 10578.12 | 11799.8 |
| ORR1D-int | 3527.381 | 4578.998 | 4365.139 |
| ORR1D1 | 121844.9 | 120678.9 | 121406.6 |
| ORR1D1-int | 20024.67 | 18994.19 | 19432.06 |
| ORR1D2 | 94091.32 | 92836.22 | 92527.93 |
| ORR1D2-int | 13533.45 | 12146.18 | 12253.38 |
| ORR1E | 142153.4 | 138146 | 138058.8 |
| ORR1E-int | 14458.09 | 16625.04 | 16140.24 |
| ORR1F | 136745.5 | 140974.9 | 135308.1 |
| ORR1F-int | 3471.026 | 2696.875 | 4017.096 |
| ORR1G | 26678.69 | 25179.94 | 26231.25 |
| ORR1G-int | 732.6099 | 728.27 | 518.6971 |
| ORSL | 1031.081 | 1422.402 | 1266.429 |
| ORSL-2a | 511.3659 | 332.2732 | 258.2258 |
| ORSL-2b | 1440.173 | 1934.467 | 1616.718 |
| PB1 | 41143.04 | 42269.25 | 44210.51 |

| | | | |
|---|---|---|---|
| PB1D10 | 239027 | 233365 | 242420.2 |
| PB1D11 | 62975.23 | 58871.53 | 62748.88 |
| PB1D7 | 85099.63 | 81964.52 | 91492.78 |
| PB1D9 | 104304 | 104454.4 | 118739 |
| Plat_L3 | 3003.492 | 3045.079 | 3734.17 |
| polypurine | 3032.713 | 2751.495 | 2593.486 |
| polypyrimidine | 2667.451 | 2680.944 | 2905.602 |
| RCHARR1 | 69643.86 | 67715.46 | 68384.94 |
| Ricksha | 859.9295 | 949.0269 | 1212.539 |
| Ricksha_0 | 699.2146 | 659.9947 | 770.1866 |
| Ricksha_a | 561.4588 | 170.6883 | 366.0071 |
| Ricksha_b | 0 | 0 | 103.2903 |
| Ricksha_c | 2598.573 | 2269.016 | 2016.407 |
| RLTR1 | 269.2498 | 475.6514 | 489.5064 |
| RLTR10 | 13806.88 | 12644.59 | 14307.96 |
| RLTR10-int | 11642.44 | 12341.9 | 13043.77 |
| RLTR10A | 4621.078 | 4808.858 | 4715.428 |
| RLTR10B | 1168.836 | 1631.78 | 1643.664 |
| RLTR10B2 | 6382.681 | 5477.956 | 5375.588 |
| RLTR10C | 12955.3 | 15441.6 | 15006.29 |
| RLTR10D | 3746.538 | 3461.559 | 3035.838 |
| RLTR10D2 | 1502.789 | 1404.196 | 1342.774 |
| RLTR10E | 1400.516 | 1577.16 | 1331.547 |
| RLTR10F | 3514.858 | 2819.771 | 2692.285 |
| RLTR11A | 22863.27 | 24444.84 | 25885.46 |
| RLTR11A2 | 24977.61 | 24572.29 | 24960.33 |
| RLTR11B | 9079.353 | 8422.898 | 7953.356 |
| RLTR11C_MM | 7029.715 | 7193.942 | 7425.677 |
| RLTR11D | 3825.852 | 4528.929 | 4785.037 |
| RLTR12A | 7175.82 | 9353.718 | 8007.247 |
| RLTR12B | 8968.731 | 8632.276 | 8180.146 |
| RLTR12B2 | 1264.848 | 1137.922 | 1199.066 |
| RLTR12BD_Mm | 8075.406 | 7869.868 | 6630.791 |
| RLTR12C | 2880.347 | 2266.74 | 2267.897 |
| RLTR12D | 5170.013 | 5521.197 | 5797.732 |
| RLTR12E | 4965.467 | 5302.716 | 5842.64 |
| RLTR12F | 803.5749 | 475.6514 | 507.4699 |
| RLTR12G | 3022.277 | 2403.291 | 2205.024 |
| RLTR12H | 1515.313 | 1665.918 | 1663.873 |
| RLTR13A | 2352.283 | 2146.121 | 2330.769 |
| RLTR13A1 | 2339.76 | 2335.016 | 2324.033 |
| RLTR13A2 | 2373.155 | 2325.912 | 2431.814 |

| | | | |
|---|---|---|---|
| RLTR13A3 | 3022.277 | 3921.279 | 3390.618 |
| RLTR13B1 | 8895.679 | 9513.027 | 8754.979 |
| RLTR13B2 | 4026.223 | 4736.031 | 4888.327 |
| RLTR13B3 | 3640.09 | 4132.932 | 3891.351 |
| RLTR13B4 | 3197.602 | 4294.517 | 4010.36 |
| RLTR13C1 | 5817.048 | 6008.228 | 5310.471 |
| RLTR13C2 | 11999.36 | 11508.94 | 11494.42 |
| RLTR13C3 | 3913.514 | 3850.728 | 3902.578 |
| RLTR13D | 308.9067 | 537.0991 | 462.5611 |
| RLTR13D1 | 3485.637 | 3807.487 | 3898.088 |
| RLTR13D2 | 4591.857 | 5270.854 | 4582.947 |
| RLTR13D3 | 5608.327 | 5036.442 | 5344.152 |
| RLTR13D3A | 1607.15 | 2608.117 | 2265.651 |
| RLTR13D3A1 | 7250.959 | 7733.317 | 8395.708 |
| RLTR13D4 | 3164.207 | 3527.558 | 3024.611 |
| RLTR13D5 | 9409.132 | 10798.88 | 8849.287 |
| RLTR13D6 | 11003.76 | 12191.7 | 13876.83 |
| RLTR13E | 6906.57 | 7983.66 | 7996.019 |
| RLTR13F | 1657.243 | 1706.883 | 1589.773 |
| RLTR13G | 7282.267 | 6406.5 | 7068.652 |
| RLTR14 | 14071.95 | 16142.56 | 16564.63 |
| RLTR14-int | 11556.87 | 13356.93 | 12379.12 |
| RLTR14_RN | 6823.082 | 7180.287 | 7998.265 |
| RLTR15 | 19304.58 | 19016.95 | 20837.7 |
| RLTR16 | 9311.033 | 9795.232 | 10111.23 |
| RLTR16B_MM | 5510.228 | 5011.408 | 4906.291 |
| RLTR16C_MM | 3903.078 | 2824.322 | 2959.493 |
| RLTR17 | 15034.16 | 16242.7 | 19470.23 |
| RLTR17B_Mm | 144749.9 | 148542.1 | 135838 |
| RLTR17C_Mm | 1803.347 | 2357.774 | 2243.197 |
| RLTR17D_Mm | 16090.28 | 15159.4 | 17649.17 |
| RLTR18 | 5203.409 | 6497.534 | 5310.471 |
| RLTR18-int | 4253.729 | 3595.833 | 5270.053 |
| RLTR18B | 8054.534 | 8236.279 | 8034.192 |
| RLTR19 | 7013.018 | 7455.664 | 7937.638 |
| RLTR19-int | 22489.66 | 22444.37 | 20541.3 |
| RLTR19A | 1099.958 | 1101.508 | 1129.457 |
| RLTR19A2 | 573.9821 | 532.5475 | 451.3339 |
| RLTR19B | 2229.138 | 2284.947 | 2202.779 |
| RLTR19C | 1874.312 | 1913.985 | 1830.035 |
| RLTR19D | 415.3543 | 250.3428 | 330.08 |
| RLTR1A2_MM | 3473.113 | 2783.357 | 2847.221 |

| | | | |
|---|---|---|---|
| RLTR1B | 7639.18 | 6909.462 | 7742.284 |
| RLTR1B-int | 8194.377 | 8370.554 | 7744.53 |
| RLTR1C | 5679.292 | 5056.925 | 6174.966 |
| RLTR1D | 2423.248 | 2564.876 | 2831.502 |
| RLTR1D2_MM | 1968.237 | 1665.918 | 1526.901 |
| RLTR1E_MM | 724.261 | 625.8571 | 561.3605 |
| RLTR1F_Mm | 559.3716 | 541.6508 | 516.4517 |
| RLTR20A | 2669.539 | 1497.505 | 2023.143 |
| RLTR20A1 | 807.7493 | 612.202 | 502.979 |
| RLTR20A2 | 1913.969 | 2353.223 | 2425.077 |
| RLTR20A2B_MM | 3187.166 | 2585.359 | 3134.637 |
| RLTR20A3_MM | 6130.129 | 6254.019 | 5898.776 |
| RLTR20A4 | 25683.09 | 25250.49 | 26628.7 |
| RLTR20B1 | 4608.554 | 4467.481 | 4632.347 |
| RLTR20B2 | 6449.472 | 5639.541 | 6150.266 |
| RLTR20B3 | 7019.279 | 6347.329 | 6922.698 |
| RLTR20B3A_MM | 4385.223 | 4342.31 | 4701.956 |
| RLTR20B4_MM | 3660.962 | 2530.738 | 3107.692 |
| RLTR20B5_MM | 6461.995 | 6260.846 | 6543.218 |
| RLTR20C | 1179.272 | 1285.852 | 1567.319 |
| RLTR20C1_MM | 12427.23 | 13484.37 | 12576.72 |
| RLTR20C2_MM | 10669.81 | 11875.35 | 11285.59 |
| RLTR20D | 4320.52 | 3953.141 | 4895.064 |
| RLTR21 | 25756.14 | 23960.08 | 23103.35 |
| RLTR22_Mur | 32550 | 28614.18 | 29056.02 |
| RLTR22_Mus | 9457.138 | 8416.071 | 8121.764 |
| RLTR23 | 10168.88 | 12030.11 | 10879.17 |
| RLTR24 | 9651.248 | 12351 | 10052.84 |
| RLTR24B_MM | 3665.137 | 2849.357 | 3042.574 |
| RLTR25A | 13610.68 | 13174.86 | 14114.85 |
| RLTR25B | 21034.88 | 20498.53 | 22054.73 |
| RLTR26 | 2210.353 | 2710.53 | 3080.747 |
| RLTR26_Mus | 7987.744 | 9417.442 | 8072.364 |
| RLTR26B_MM | 3798.718 | 4212.587 | 3940.751 |
| RLTR26C_MM | 3341.619 | 3454.731 | 3648.843 |
| RLTR26D_MM | 1955.714 | 2020.949 | 2175.833 |
| RLTR27 | 4090.927 | 4121.553 | 3653.334 |
| RLTR28 | 15528.82 | 16640.97 | 17831.06 |
| RLTR28B | 3208.038 | 2626.324 | 2779.857 |
| RLTR3_Mm | 29.22091 | 209.3776 | 105.5358 |
| RLTR30 | 2717.544 | 3070.113 | 2858.448 |
| RLTR30B_MM | 1909.795 | 2469.291 | 2483.459 |

| | | | |
|---|---|---|---|
| RLTR30C_MM | 1193.883 | 1267.645 | 1122.721 |
| RLTR30D_MM | 3164.207 | 2273.568 | 2478.968 |
| RLTR30D_RN | 373.6102 | 250.3428 | 538.9061 |
| RLTR30D2_MM | 5572.844 | 5746.506 | 6778.99 |
| RLTR30E_MM | 1790.824 | 2330.464 | 2663.094 |
| RLTR31_Mm | 989.3364 | 1281.3 | 1497.71 |
| RLTR31_Mur | 2876.172 | 3468.386 | 2660.849 |
| RLTR31A_Mm | 1847.179 | 2002.743 | 1910.871 |
| RLTR31B_Mm | 2473.341 | 3115.63 | 2775.366 |
| RLTR31B2 | 8693.22 | 10311.85 | 9963.027 |
| RLTR31C_MM | 4493.758 | 4508.447 | 4836.682 |
| RLTR31D_MM | 20410.8 | 23969.19 | 22883.3 |
| RLTR31M | 2575.614 | 2309.982 | 2189.306 |
| RLTR33 | 15848.17 | 17694.69 | 17732.26 |
| RLTR34B_MM | 4708.74 | 4954.512 | 4434.748 |
| RLTR34C_MM | 4470.799 | 2867.563 | 3911.56 |
| RLTR34D_MM | 3694.357 | 3591.282 | 4055.268 |
| RLTR35B_MM | 2320.975 | 3773.349 | 3175.055 |
| RLTR4_Mm | 1014.383 | 903.51 | 1109.248 |
| RLTR4_MM-int | 9377.824 | 11005.98 | 9823.809 |
| RLTR40 | 25674.74 | 25059.32 | 26536.63 |
| RLTR41 | 6925.355 | 7184.839 | 7445.886 |
| RLTR41A2 | 3245.608 | 3534.386 | 3294.064 |
| RLTR41B | 782.7029 | 974.0612 | 419.8977 |
| RLTR41C | 678.3425 | 1115.163 | 1138.439 |
| RLTR42 | 1102.046 | 1315.438 | 1181.103 |
| RLTR42-int | 20254.26 | 20000.12 | 19308.56 |
| RLTR43A | 0 | 6.827532 | 17.96354 |
| RLTR43B | 31.30811 | 0 | 0 |
| RLTR43C | 1650.981 | 1750.124 | 1724.5 |
| RLTR44-int | 6570.53 | 5047.822 | 5319.452 |
| RLTR44A | 1281.545 | 2093.776 | 1226.011 |
| RLTR44B | 1567.493 | 955.8544 | 1185.593 |
| RLTR44C | 1239.801 | 1574.884 | 1302.356 |
| RLTR44D | 713.825 | 566.6851 | 532.1698 |
| RLTR44E | 941.3306 | 839.7864 | 904.9132 |
| RLTR45 | 8678.609 | 9599.509 | 8094.819 |
| RLTR45-int | 12767.45 | 11925.42 | 13257.09 |
| RLTR46 | 375.6974 | 323.1698 | 345.7981 |
| RLTR46A | 154.4534 | 300.4114 | 168.4082 |
| RLTR46A2 | 344.3893 | 134.2748 | 352.5344 |
| RLTR46B | 438.3136 | 325.4457 | 381.7252 |

| | | | |
|---|---|---|---|
| RLTR47 | 1880.574 | 1752.4 | 1758.181 |
| RLTR47_MM | 260.901 | 129.7231 | 184.1263 |
| RLTR48A | 6034.117 | 4563.067 | 5014.072 |
| RLTR48B | 841.1447 | 926.2684 | 1140.685 |
| RLTR48C | 1435.999 | 1615.849 | 1549.355 |
| RLTR49 | 5554.059 | 5739.678 | 6296.22 |
| RLTR5_Mm | 1006.034 | 1137.922 | 1210.293 |
| RLTR50A | 782.7029 | 446.0654 | 678.1235 |
| RLTR50B | 1918.144 | 2480.67 | 2449.777 |
| RLTR51A_Mm | 8407.272 | 9777.025 | 9424.121 |
| RLTR51B_Mm | 4007.439 | 3732.384 | 4102.423 |
| RLTR53_Mm | 3754.887 | 3108.803 | 2254.424 |
| RLTR53B_Mm | 991.4236 | 896.6825 | 664.6509 |
| RLTR6-int | 32191 | 33413.94 | 31032.01 |
| RLTR6_Mm | 1980.76 | 1652.263 | 1470.765 |
| RLTR6B_Mm | 4508.368 | 4014.589 | 4524.566 |
| RLTR6C_Mm | 434.1392 | 587.1677 | 428.8794 |
| RLTR8 | 2074.684 | 1488.402 | 1367.474 |
| RLTR9A | 3711.055 | 4055.554 | 3893.597 |
| RLTR9A2 | 1580.016 | 1388.265 | 1605.491 |
| RLTR9A3 | 6409.815 | 4173.898 | 4457.203 |
| RLTR9A3A | 947.5923 | 1119.715 | 1219.275 |
| RLTR9A3B | 2425.335 | 1793.365 | 2449.777 |
| RLTR9A4 | 39.65694 | 31.86181 | 136.972 |
| RLTR9B | 1306.592 | 1424.678 | 1466.274 |
| RLTR9B2 | 930.8946 | 1028.681 | 985.7491 |
| RLTR9C | 1721.946 | 1415.575 | 1634.682 |
| RLTR9D | 4383.136 | 5147.959 | 5160.026 |
| RLTR9D2 | 989.3364 | 1547.574 | 1286.638 |
| RLTR9E | 9060.568 | 7489.802 | 7578.367 |
| RLTR9F | 2619.446 | 3086.044 | 2699.021 |
| RLTRETN_Mm | 8309.174 | 6873.048 | 7095.597 |
| RMER10A | 27607.5 | 27278.26 | 28562.02 |
| RMER10B | 16770.71 | 16934.55 | 17431.37 |
| RMER12 | 26184.02 | 23730.22 | 26669.12 |
| RMER12B | 9148.231 | 9599.509 | 11081.26 |
| RMER12C | 25501.5 | 25100.28 | 24657.2 |
| RMER13A | 11078.9 | 11793.42 | 9632.947 |
| RMER13A1 | 9803.614 | 10889.91 | 9745.219 |
| RMER13A2 | 13109.75 | 14704.23 | 14153.02 |
| RMER13B | 22178.67 | 22865.4 | 23348.11 |
| RMER15 | 99202.89 | 107756.7 | 103660.8 |

| | | | |
|---|---|---|---|
| RMER15-int | 29229.26 | 27244.13 | 28530.59 |
| RMER16 | 4109.712 | 4205.759 | 3664.562 |
| RMER16-int | 40103.61 | 38402.59 | 39874.56 |
| RMER16_Mm | 3754.887 | 4169.346 | 3345.709 |
| RMER16A2 | 1481.917 | 1672.745 | 1874.944 |
| RMER16A3 | 2748.852 | 2560.324 | 2609.204 |
| RMER16B | 905.8481 | 737.3734 | 716.296 |
| RMER16B2 | 2517.172 | 2030.053 | 2445.286 |
| RMER16B3 | 678.3425 | 512.0649 | 767.9412 |
| RMER16C | 1260.673 | 1099.233 | 1686.327 |
| RMER17A | 13143.15 | 15555.39 | 12695.73 |
| RMER17A-int | 3375.015 | 3233.974 | 3305.291 |
| RMER17A2 | 14364.16 | 15771.6 | 16607.29 |
| RMER17B | 30370.96 | 31829.95 | 31191.44 |
| RMER17B2 | 7073.547 | 7731.042 | 7715.339 |
| RMER17C | 33745.97 | 34929.65 | 35799.08 |
| RMER17C-int | 17678.65 | 16495.32 | 16191.88 |
| RMER17C2 | 14929.8 | 13711.96 | 12783.3 |
| RMER17D | 6178.135 | 8136.142 | 6983.325 |
| RMER17D2 | 10926.53 | 11850.32 | 11757.13 |
| RMER19A | 21869.76 | 21156.24 | 22613.85 |
| RMER19B | 67690.23 | 72185.22 | 69503.17 |
| RMER19B2 | 26895.76 | 25908.21 | 25887.7 |
| RMER19C | 25107.02 | 28898.67 | 28142.13 |
| RMER1A | 40114.04 | 42451.32 | 40536.97 |
| RMER1B | 69698.12 | 68138.76 | 72197.7 |
| RMER1C | 39815.57 | 37078.05 | 41435.14 |
| RMER2 | 18413.35 | 17446.62 | 17355.02 |
| RMER20A | 8567.987 | 9283.167 | 9583.547 |
| RMER20B | 13944.63 | 12221.28 | 14027.28 |
| RMER20C_Mm | 11940.91 | 11062.88 | 10960 |
| RMER21A | 4178.59 | 4178.449 | 4782.792 |
| RMER21B | 6578.878 | 6044.641 | 6756.535 |
| RMER30 | 14754.47 | 16497.59 | 16804.89 |
| RMER3D-int | 29014.27 | 26834.47 | 27946.77 |
| RMER3D1 | 0 | 0 | 0 |
| RMER3D2 | 0 | 0 | 2.245442 |
| RMER3D3 | 43.83136 | 0 | 0 |
| RMER3D4 | 16.69766 | 0 | 11.22721 |
| RMER4A | 18872.53 | 19358.33 | 20292.06 |
| RMER4B | 38124.93 | 40052.58 | 40467.36 |
| RMER5 | 60476.84 | 60407.72 | 58720.56 |

| | | | |
|---|---|---|---|
| RMER6-int | 7292.703 | 6932.22 | 8155.446 |
| RMER6A | 48371.04 | 46682.11 | 46954.44 |
| RMER6B | 10344.2 | 9308.201 | 11128.41 |
| RMER6BA | 5355.775 | 4578.998 | 6139.039 |
| RMER6C | 48709.16 | 48291.13 | 51871.96 |
| RMER6D | 18567.8 | 20079.77 | 19560.05 |
| RNERVK23-int | 302.6451 | 279.9288 | 303.1347 |
| RNLTR23 | 8.34883 | 0 | 0 |
| RodERV21-int | 20070.59 | 21370.17 | 20673.79 |
| RSINE1 | 497579.9 | 485157.6 | 505426.6 |
| SRV_MM-int | 1648.894 | 1847.985 | 2508.159 |
| SSU-rRNA_Hsa | 4115.973 | 5004.581 | 4091.196 |
| SUBTEL_sa | 58.44181 | 0 | 38.17252 |
| SYNREP_MM | 357002.3 | 394053.3 | 308948.1 |
| T-rich | 70361.86 | 66682.22 | 71043.54 |
| Tigger1 | 10379.68 | 11008.26 | 10425.59 |
| Tigger10 | 1659.33 | 1918.536 | 1347.265 |
| Tigger11a | 421.6159 | 407.376 | 473.7883 |
| Tigger12 | 1104.133 | 1258.542 | 770.1866 |
| Tigger12A | 262.9882 | 550.7542 | 581.5695 |
| Tigger12c | 436.2264 | 593.9952 | 696.0871 |
| Tigger13a | 6495.39 | 5933.125 | 4695.219 |
| Tigger14a | 1611.324 | 1408.747 | 1131.703 |
| Tigger15a | 4153.543 | 4458.378 | 4165.295 |
| Tigger16a | 2272.969 | 1176.611 | 1697.554 |
| Tigger16b | 711.7378 | 578.0643 | 877.9679 |
| Tigger17 | 1506.964 | 1857.089 | 1493.219 |
| Tigger17a | 8459.452 | 8798.412 | 8344.063 |
| Tigger17b | 640.7727 | 384.6176 | 570.3423 |
| Tigger17c | 5094.874 | 3541.213 | 4710.938 |
| Tigger18a | 1943.19 | 2073.294 | 2229.724 |
| Tigger19a | 3441.805 | 4303.621 | 4405.557 |
| Tigger1a_Art | 54.2674 | 141.1023 | 114.5175 |
| Tigger1a_Mars | 0 | 0 | 0 |
| Tigger20a | 1125.005 | 1326.817 | 1473.01 |
| Tigger2b | 1028.993 | 1381.437 | 1405.647 |
| Tigger5 | 11826.12 | 12608.17 | 11436.04 |
| Tigger5b | 11231.26 | 13327.34 | 11590.97 |
| Tigger6a | 4080.491 | 3780.177 | 3837.461 |
| Tigger6b | 1688.551 | 1695.504 | 1767.163 |
| Tigger7 | 35931.28 | 36495.43 | 35448.79 |
| Tigger8 | 1369.208 | 723.7183 | 1230.502 |

| | | | |
|---|---|---|---|
| Tigger9a | 1402.604 | 1226.68 | 1571.809 |
| Tigger9b | 2892.87 | 2542.118 | 2400.378 |
| tRNA-Ala-GCA | 2076.772 | 2307.706 | 1609.982 |
| tRNA-Ala-GCG | 684.6041 | 259.4462 | 597.2876 |
| tRNA-Ala-GCY | 14.61045 | 207.1018 | 0 |
| tRNA-Ala-GCY_ | 6324.239 | 6843.462 | 6987.816 |
| tRNA-Arg-AGA | 185.7615 | 113.7922 | 49.39973 |
| tRNA-Arg-AGG | 0 | 97.86129 | 15.71809 |
| tRNA-Arg-CGA | 41.74415 | 27.31013 | 0 |
| tRNA-Arg-CGA_ | 0 | 0 | 42.6634 |
| tRNA-Arg-CGG | 0 | 0 | 6.736326 |
| tRNA-Arg-CGY_ | 127.3197 | 207.1018 | 89.81768 |
| tRNA-Asn-AAC | 81.4011 | 136.5506 | 78.59047 |
| tRNA-Asp-GAY | 137.7557 | 106.9647 | 56.13605 |
| tRNA-Cys-TGY | 237.9417 | 70.55116 | 339.0618 |
| tRNA-Gln-CAA | 37.56974 | 43.24103 | 0 |
| tRNA-Gln-CAA_ | 13247.51 | 8750.62 | 7230.324 |
| tRNA-Gln-CAG | 60.52902 | 91.03375 | 80.83592 |
| tRNA-Glu-GAA | 20.87208 | 0 | 0 |
| tRNA-Glu-GAG | 41.74415 | 0 | 0 |
| tRNA-Glu-GAG_ | 91.83714 | 97.86129 | 139.2174 |
| tRNA-Gly-GGA | 196.1975 | 63.72363 | 199.8443 |
| tRNA-Gly-GGG | 70.96506 | 11.37922 | 51.64517 |
| tRNA-Gly-GGY | 196.1975 | 88.75791 | 305.3801 |
| tRNA-His-CAY | 0 | 0 | 0 |
| tRNA-His-CAY_ | 0 | 184.3434 | 110.0267 |
| tRNA-Ile-ATA | 87.66272 | 248.067 | 143.7083 |
| tRNA-Ile-ATT | 285.9474 | 36.4135 | 217.8079 |
| tRNA-Leu-CTA | 4.174415 | 0 | 0 |
| tRNA-Leu-CTA_ | 0 | 0 | 15.71809 |
| tRNA-Leu-CTG | 0 | 145.654 | 65.11782 |
| tRNA-Leu-CTY | 0 | 77.37869 | 13.47265 |
| tRNA-Leu-TTA | 75.13947 | 213.9293 | 78.59047 |
| tRNA-Leu-TTA(m) | 20456.72 | 10443.85 | 9451.066 |
| tRNA-Leu-TTG | 39.65694 | 102.413 | 74.09959 |
| tRNA-Lys-AAA | 108.5348 | 186.6192 | 240.2623 |
| tRNA-Lys-AAG | 350.6509 | 828.4072 | 339.0618 |
| tRNA-Met | 0 | 61.44778 | 53.89061 |
| tRNA-Met-i | 171.151 | 38.68935 | 31.43619 |
| tRNA-Met_ | 152.3662 | 161.5849 | 103.2903 |
| tRNA-Phe-TTY | 85.57551 | 56.8961 | 246.9986 |
| tRNA-Pro-CCA | 31.30811 | 79.65453 | 20.20898 |

| | | | |
|---|---|---|---|
| tRNA-Pro-CCG | 0 | 79.65453 | 4.490884 |
| tRNA-Pro-CCY | 108.5348 | 68.27532 | 78.59047 |
| tRNA-SeC(e)-TGA | 14.61045 | 95.58544 | 47.15428 |
| tRNA-Ser-AGY | 102.2732 | 172.9641 | 83.08136 |
| tRNA-Ser-TCA | 0 | 0 | 0 |
| tRNA-Ser-TCA(m) | 0 | 0 | 15.71809 |
| tRNA-Ser-TCA_ | 52.18019 | 0 | 0 |
| tRNA-Ser-TCG | 79.31389 | 0 | 0 |
| tRNA-Ser-TCY | 62.61623 | 195.7226 | 80.83592 |
| tRNA-Thr-ACA | 20.87208 | 68.27532 | 4.490884 |
| tRNA-Thr-ACG | 0 | 47.79272 | 0 |
| tRNA-Thr-ACG_ | 0 | 0 | 80.83592 |
| tRNA-Thr-ACY | 52.18019 | 0 | 60.62694 |
| tRNA-Thr-ACY_ | 154.4534 | 104.6888 | 42.6634 |
| tRNA-Trp-TGG | 6.261623 | 43.24103 | 38.17252 |
| tRNA-Tyr-TAC | 123.1452 | 34.13766 | 121.2539 |
| tRNA-Tyr-TAT | 66.79064 | 0 | 0 |
| tRNA-Val-GTA | 125.2325 | 40.96519 | 107.7812 |
| tRNA-Val-GTG | 73.05227 | 143.3782 | 211.0716 |
| tRNA-Val-GTY | 73.05227 | 25.03428 | 101.0449 |
| U1 | 1079.086 | 1392.816 | 1403.401 |
| U13 | 0 | 65.99947 | 62.87238 |
| U13_ | 144.0173 | 141.1023 | 38.17252 |
| U14 | 0 | 29.58597 | 44.90884 |
| U17 | 150.2789 | 127.4473 | 161.6718 |
| U2 | 2389.853 | 2733.288 | 2175.833 |
| U3 | 181.5871 | 304.9631 | 139.2174 |
| U4 | 267.1626 | 461.9963 | 374.9888 |
| U5 | 229.5928 | 270.8254 | 314.3619 |
| U6 | 4241.206 | 4260.38 | 4495.375 |
| U7 | 306.8195 | 195.7226 | 112.2721 |
| U8 | 2.087208 | 13.65506 | 38.17252 |
| UCON1 | 87.66272 | 50.06856 | 2.245442 |
| UCON10 | 158.6278 | 259.4462 | 154.9355 |
| UCON11 | 599.0286 | 327.7215 | 345.7981 |
| UCON12 | 283.8602 | 120.6197 | 159.4264 |
| UCON12A | 58.44181 | 86.48207 | 163.9173 |
| UCON13 | 336.0404 | 22.75844 | 211.0716 |
| UCON14 | 308.9067 | 332.2732 | 312.1165 |
| UCON15 | 146.1045 | 70.55116 | 141.4629 |
| UCON16 | 121.058 | 125.1714 | 132.4811 |
| UCON17 | 131.4941 | 141.1023 | 159.4264 |

| | | | |
|---|---|---|---|
| UCON18 | 338.1276 | 125.1714 | 150.4446 |
| UCON19 | 133.5813 | 188.895 | 132.4811 |
| UCON2 | 569.8077 | 939.9235 | 392.9524 |
| UCON20 | 269.2498 | 534.8233 | 276.1894 |
| UCON21 | 177.4126 | 63.72363 | 145.9537 |
| UCON22 | 108.5348 | 152.4815 | 197.5989 |
| UCON23 | 6.261623 | 0 | 136.972 |
| UCON24 | 0 | 70.55116 | 92.06313 |
| UCON25 | 48.00578 | 52.34441 | 89.81768 |
| UCON26 | 505.1042 | 589.4436 | 693.8416 |
| UCON27 | 601.1158 | 316.3423 | 451.3339 |
| UCON28a | 369.4357 | 887.5791 | 406.425 |
| UCON28b | 500.9298 | 443.7896 | 276.1894 |
| UCON28c | 338.1276 | 40.96519 | 229.0351 |
| UCON29 | 855.7551 | 764.6835 | 529.9243 |
| UCON31 | 613.639 | 507.5132 | 507.4699 |
| UCON4 | 296.3835 | 339.1007 | 303.1347 |
| UCON5 | 304.7323 | 316.3423 | 348.0435 |
| UCON6 | 219.1568 | 534.8233 | 258.2258 |
| UCON7 | 732.6099 | 794.2695 | 666.8963 |
| UCON8 | 137.7557 | 637.2363 | 314.3619 |
| UCON9 | 139.8429 | 257.1704 | 204.3352 |
| URR1A | 106291 | 109399.8 | 108740 |
| URR1B | 97069.76 | 101493.5 | 95887.11 |
| X1_LINE | 98.09876 | 238.9636 | 260.4713 |
| X2_LINE | 164.8894 | 380.0659 | 150.4446 |
| X3_LINE | 1035.255 | 1008.199 | 651.1782 |
| X5A_LINE | 505.1042 | 614.4778 | 639.951 |
| X5B_LINE | 256.7265 | 327.7215 | 287.4166 |
| X6A_LINE | 644.9472 | 423.307 | 639.951 |
| X6B_LINE | 1870.138 | 1074.198 | 821.8318 |
| X7A_LINE | 661.6448 | 671.3739 | 482.7701 |
| X7B_LINE | 377.7846 | 314.0665 | 738.7505 |
| X7C_LINE | 540.5868 | 639.5121 | 577.0786 |
| X7D_LINE | 0 | 0 | 11.22721 |
| X8_LINE | 795.2261 | 910.3375 | 929.613 |
| X9_LINE | 227.5056 | 84.20622 | 123.4993 |
| YREP_Mm | 6833.518 | 5302.716 | 6341.129 |
| Zaphod | 7390.802 | 5830.712 | 6558.936 |
| Zaphod2 | 1120.83 | 1156.129 | 1464.028 |
| Zaphod3 | 3084.893 | 3734.66 | 3008.892 |
| ZP3AR | 9004.214 | 10109.3 | 9350.021 |

| | | | |
|---|---|---|---|
| (CTACG)n | 0 | 0 | 0 |
| CENSAT_MC | 0 | 0 | 0 |
| (CACGA)n | 0 | 0 | 0 |
